# Synthetic ratio computation for programming population composition and multicellular morphology

**DOI:** 10.1101/2024.11.26.624747

**Authors:** Bolin An, Tzu-Chieh Tang, Qian Zhang, Teng Wang, Yanyi Wang, Kesheng Gan, Kun Liu, Daniel L Zhang, Yuzhu Liu, Yu Kui Pan, Min Yu, William M. Shaw, Qianyi Liang, Yaomin Wang, Chunbo Lou, Timothy K. Lu, George M. Church, Chao Zhong

**Author notes:** Correspondence should be addressed to T.-C.T., G.M.C, and C.Z. B.A, T.-C.T and Q.Z. contributed equally.

## Abstract

Recent advancements in genetic engineering have provided diverse tools for artificially synthesizing population diversity in both prokaryotic and eukaryotic systems. However, achieving precise control over the ratios of multiple cell types within a population derived from a single founder remains a significant challenge. In this study, we introduce a suite of recombinase-mediated genetic devices designed to achieve accurate population ratio control, enabling the distribution of distinct functionalities across multiple cell types. We systematically evaluated key parameters influencing recombination efficiency and developed data-driven models to reliably predict binary differentiation outcomes. Using these devices, we implemented parallel and series circuit topologies to create user-defined, complex cell fate branching programs. These branching devices facilitated the autonomous differentiation of precision fermentation consortia from a single founder strain, optimizing cell-type ratios for applications such as pigmentation and cellulose degradation. Beyond biomanufacturing, we engineered multicellular aggregates with genetically encoded morphologies by coordinating self-organization through cell adhesion molecules (CAMs). Our work provides a comprehensive characterization of recombinase-based cell fate branching mechanisms and introduces a novel approach for the bottom-up, high-resolution construction of synthetic consortia and multicellular assemblies.

## Main

In multicellular organisms, pattern formation is woven through asymmetric cell divisions, where stem cells give rise to progeny with distinct phenotypes^1, 2^. Through the orchestration of complex gene regulatory networks, eukaryotic stem cells undergo differentiation into specialized cell lineages that integrate seamlessly to form functional tissues or organs. This sophisticated differentiation machinery reflects the genetic intricacy of multicellular eukaryotes and has significantly contributed to their evolutionary triumph^2^. Parallel yet more streamlined phenomena are observed in microorganisms. For example, *Caulobacter crescentus* undergoes asymmetric division to produce motile cells lacking reproductive capabilities and sessile cells capable of division but not motility^3^. Similarly, under nitrogen-limiting conditions, *Anabaena* filaments’ central cells asymmetrically divide to generate progeny specialized in nitrogen fixation or photosynthesis, ensuring the survival and proliferation of the colony^4^. Natural selection has consistently favored programmed differentiation and cooperative division of labor, maximizing population fitness. These cellular communities accomplish sophisticated tasks by efficiently allocating resources and exploiting complementary strengths, achieving benefits unattainable by a single cell type^5^.

Synthetic biology has unlocked the reprogramming of cellular functions, facilitating the emergence of diverse artificial cell states^6–12^, even in simple unicellular systems like *Escherichia coli*. In 2000, A seminal achievement was the development of a genetic toggle switch in *E. coli*, enabling reversible switching between two distinct states^13^. Advances in genetic tools have since enabled engineered bacterial stem cells through mechanisms such as the bacterial chromosome partitioning system (par), inducing asymmetric cell division, and generating progeny with distinct genetic content^7^. Epigenetic tools, including phase-separated and scaffolding proteins like PopZ, have further advanced microbial cell differentiation^8, 9^. Beyond bacteria, mammalian systems have also been engineered; for example, the MultiFate circuit generates multiple cellular phenotypes in Chinese hamster ovary (CHO) cells^14^. These examples highlight the potential of genetic circuits to program and rewrite cell fate. However, scaling genetic toggle switches and DNA partitioning systems with increasing cell states and quantitatively controlling the ratio of specific cell types within offspring populations remain challenges. Moreover, most artificial systems are incapable of autonomously generating complex multicellular systems from a single founder cell type. These limitations impede the capacity of synthetic communities to differentiate autonomously and accurately perform complex tasks, thereby constraining their potential in biomanufacturing, regenerative medicine, and therapeutic applications.

Recombinases, enzymes that catalyze recombination between specific DNA sequences, offer a promising solution. These enzymes target *attB* (bacterial) and *attP* (phage) sites, with recombination outcome—excision or inversion— determined by the number and orientation of these sites^15^. Serine recombinases, favored for their irreversible actions and independence from additional cofactors, are widely used in constructing Boolean logic gates^16^, amplification circuits^17^, and information storage systems^18, 19^. Strategic positioning of multiple *att* sites enables programmable rearrangement of gene elements, expanding the toolbox for cell state programming^18, 20^.

Here, we present a suite of recombinase-based cell fate branching devices for generating diverse progeny cell types with user-defined ratios and quantitatively managing multicellular morphology. By manipulating DNA sequences between *att* sites and employing mutated *att* variants, we achieved high-precision control over the ratios (0.1%-99.9%) of two progeny cell types bifurcated from a single founder. Our branching devices can be implemented in parallel or series configurations, enabling scalability and complex outputs, including transcription factors, secreted enzymes, and biosynthesis pathways. Additionally, we used cell surface display^21^ and synthetic adhesion molecules^22–24^ to engineer self-organizing cellular architectures with highly controllable geometries, all differentiated from a single founder. Unlike traditional co-culture strategies that rely on manually mixing multiple cell types, our approach enables autonomous, programmable diversification across bacteria, yeast, and mammalian cells. This platform lays the groundwork for bottom-up construction of synthetic consortia and the rational design of coordinated, complex multicellular systems.

## Results

### Design and characterization of cell fate branching devices

Binary cell fate branching devices were developed using a Bxb1 recombinase-based framework. The device contained a single Bxb1 *attP* site flanked by two identical Bxb1 *attB* sites in the configuration *attB*-*attP*-*attB* (**Fig. 1a**). Recombination between the *attP* site and one of the *attB* sites was triggered by the recombinase, resulting in the excision of either the left or right DNA segment and the generation of two distinct recombination products (**Fig. S1, Fig. S2**). To ensure the differentiation of each parent cell into a single progeny type, the system was constructed to allow only one copy of the device per genome (**Fig. S3**). We assumed that such a mechanism would be host-agnostic and set out to test our hypotheses in model bacterial (*E. coli*), yeast (*Saccharomyces cerevisiae*), and mammalian (HEK293FT) systems.

**Fig. 1.**
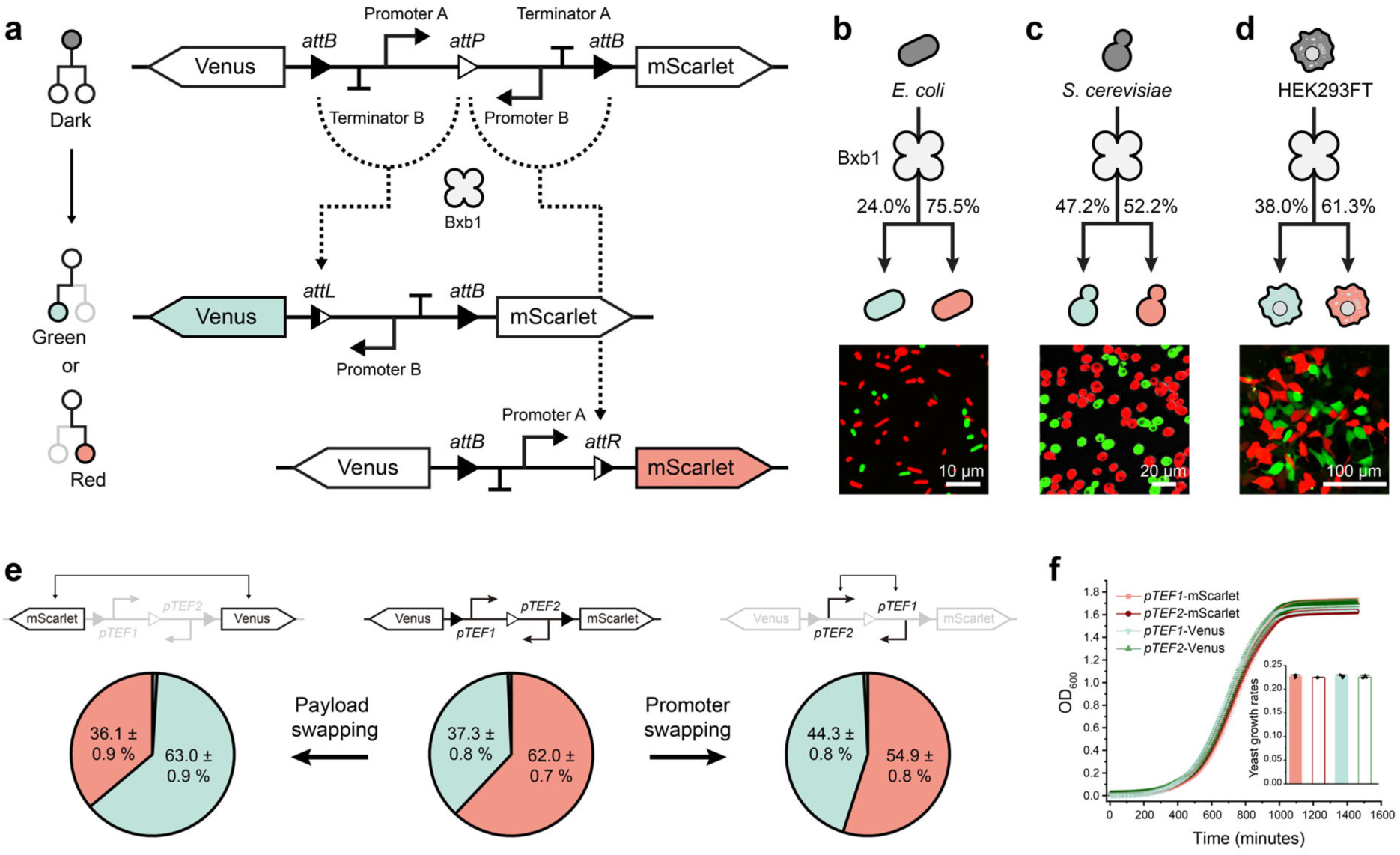
Design and characterization of gene devices for cellular differentiation control. **a**, Diagram illustrating the design of general branching devices that probabilistically generate two types of fluorescent cells. **b-d**, Schematics and fluorescence data demonstrating cellular differentiation in engineered cells: *E. coli* (**b**), *S. cerevisiae* (**c**), and HEK293FT cells (**d**). Progeny cell type ratios were determined via flow cytometry and confocal fluorescence microscopy analysis, and are reported as mean values. **e**, Factors influencing yeast cellular differentiation driven by the designed gene devices. Progeny cell ratios were measured by flow cytometry (data are mean ± s.d. of *n* = 8 replicates). **f**, Growth curves of four selected yeast strains, with their respective growth rates shown in the inset for comparison (data are mean ± s.d. of *n* = 4 replicates).

To track differentiation, two constitutive promoters with comparable transcription strengths were incorporated into the branching device. The promoters were arranged to face opposite directions and separated by *att* sites, creating a genetic device in the order *attB*-*attP*-*attB* (**Fig. 1a**). Using identical promoters on both left and right arms failed due to homologous recombination in *E. coli* (**Fig. S4**). To minimize leaky expression caused by transcriptional read-through, we placed terminators between the promoters and their downstream coding sequences. Reporter genes encoding Venus (green fluorescence) and mScarlet (red fluorescence) were positioned on opposite sides. Upon induction with small molecules such as aTc or β-estradiol (**Fig. S5**), Bxb1 recombinase excised one of the DNA segments flanked by *att* sites, leading to the expression of either Venus or mScarlet. This recombination-mediated differentiation produced two distinct cell populations in *E. coli*, *S. cerevisiae*, and HEK293FT cells, as confirmed by flow cytometry and confocal microscopy (**Fig. 1b-d**, **Extended Data** Fig.1). Despite these successes, leaky recombinase activity remained a challenge, with spontaneous recombination detectable even without induction (**Extended Data** Fig.2). Using yeast as an example, we fused degradation tags (e.g., UbiM, UbiY, UbiR)^25^ to Bxb1 and found that they effectively reduced leakiness. Particularly, the UbiY tag achieved <1% spontaneous recombination over 15 passages without impairing inducibility (**Extended Data** Fig.2), indicating that protein destabilization provides a robust strategy for minimizing background activity.

To systematically investigate the branching mechanism, we use the model unicellular eukaryote *S. cerevisiae* as the chassis for detailed characterization, owing to its extensive genetic tools and highly efficient homologous recombination that expedites rapid, iterative genome engineering. We swapped the two fluorescent reporters and promoters within the genetic circuit to probe the underlying factors leading to the observed constant progeny cell ratios. Notably, in *S. cerevisiae*, when fluorescent proteins are used as reporters, we found that terminators between the *att* sites are not required to prevent leaky expression of the output (**Fig. S6-S7**). Therefore, to simplify the design, the branching devices in the following investigation only separate the promoter and coding sequence of fluorescent proteins from another promoter in the reverse complement strand to inhibit output gene expression.

As depicted in **Fig. 1e**, swapping the two fluorescent reporters did not appreciably change the outcome of the recombination event. Progeny with the reporter on the left side consistently represented 36.1 ± 0.9% and 37.3 ± 0.8% of the population in the two designs (**Fig. 1e**). However, swapping the positions of the promoters led to altered ratios, with green cells increasing from 37.3 ± 0.8% to 44.3 ± 0.8% and red cells decreasing from 62.0 ± 0.7% to 54.9 ± 0.8% (**Fig. 1e**). We conducted a growth rate analysis of the four yeast progenies (labeled *pTEF1*-mScarlet, *pTEF2*-mScarlet, *pTEF1*-Venus, and *pTEF2*-Venus, according to their respective promoter-reporter pairings) differentiated from the branching devices shown in **Fig. 1e** and found they share indistinguishable growth rates (0.22∼0.23 OD_600_/hour, **Fig. 1f)**. Furthermore, competition assays based on passaging every 24 hours at 1000-fold dilution for seven days show that the red-to-green ratio remains essentially constant (**Fig. S8**). These findings collectively suggest that the observed variation in progeny ratio is not a consequence of differential growth caused by metabolic burden associated with protein overexpression, but rather the result of recombinase action on different device topologies.

### Exploration of critical factors influencing cell differentiation devices

#### Sequence between att sites and steric hindrance

We hypothesized that promoter strength could influence differential excision and tested this by systematically varying promoters positioned between *att* sites. Ten promoters, ranging from the strongest (*pTDH3*) to the weakest (*pREV1*), were incorporated into the left and right arms of our branching device design (**Fig. 2a, Extended Data** Fig. 3a). A library of 90 unique designs, combining these promoters, revealed distinct progeny cell ratios upon recombinase induction (**Fig. 2b, Extended Data** Fig. 3a**, Fig. S9-S19**). For example, a device with *pTDH3* on the left arm and *pREV1* on the right arm resulted in 84.5 ± 1.0% red fluorescent cells (**Fig. S16a**). In contrast, a device featuring *pTEF2* and *pRPL18B* yielded only 15.0 ± 1.3% red fluorescent cells (**Fig. S18d**). Growth assays indicated no significant metabolic burden from expressing fluorescent proteins driven by strong or weak promoters over a week of competition (**Fig. S20-S21**). The closer analysis demonstrated that strong promoters, such as *pTDH3*, reduced Bxb1 recombination efficiency when located between *att* sites (**Fig. 2b**). For instance, devices with *pTDH3* on the left arm and *pREV1* on the right arm showed only 14.9 ± 0.9% green fluorescent cells, suggesting a bias toward right-arm excision (**Fig. S16a**). Substituting *pTDH3* with the weaker *pSAC6* promoter increased the proportion of green cells to 60.9 ± 3.6%, favoring left-arm excision (**Fig. S15a**). We propose that steric hindrance from transcription machinery recruited by strong promoters may obstruct recombinase binding and DNA bending, reducing recombination efficiency.

**Fig. 2.**
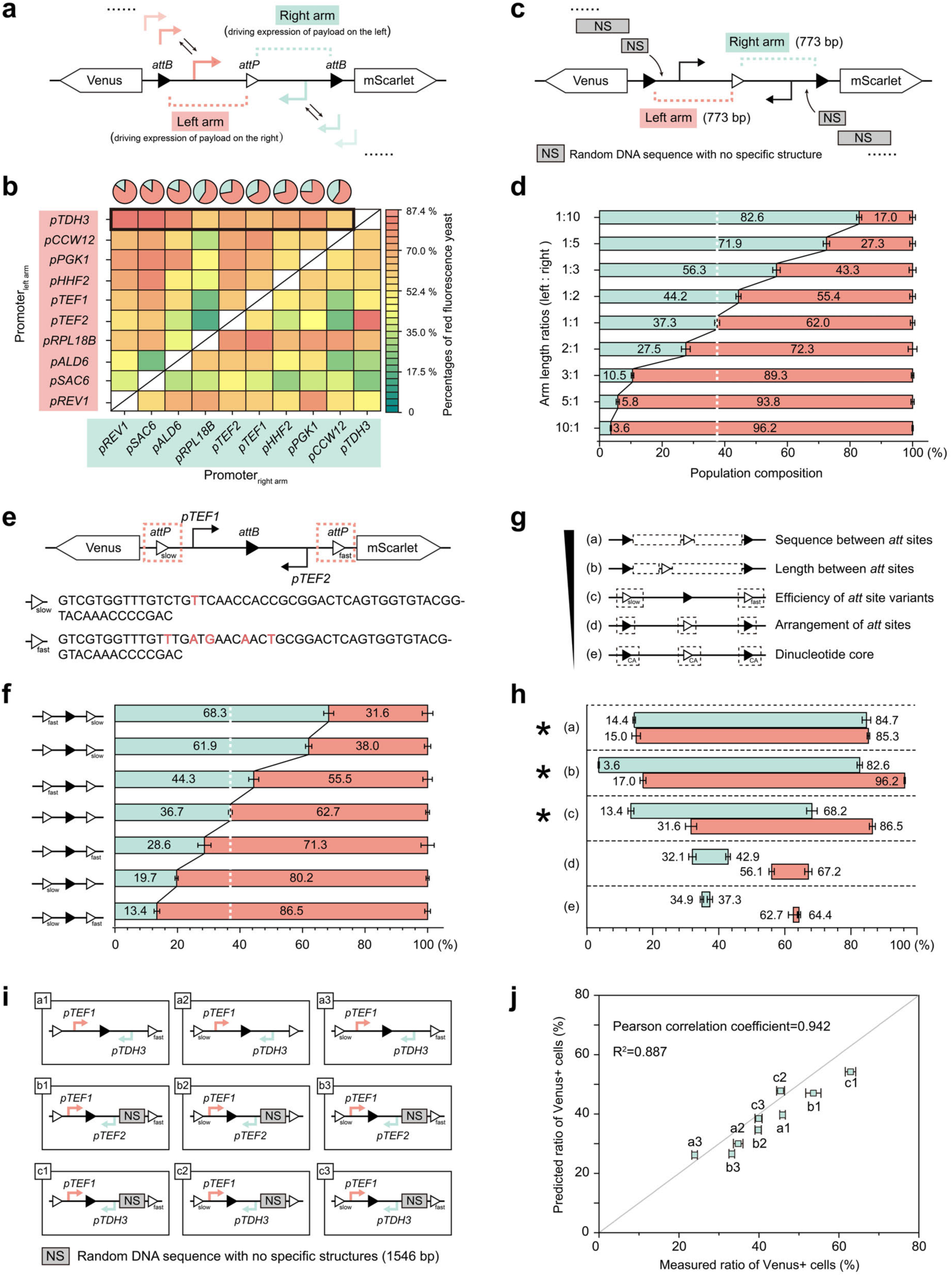
Identifying factors influencing the performance of recombinase-based cellular differentiation circuits. **a**, Schematic of branching devices using different yeast promoters placed on the left or right arm. **b**, Heat map comparing the performance of 90 constructed strains, each with two different constitutive yeast promoters on the arms. The transcriptional strength of the promoters decreases from the strongest *pTDH3* to the weakest *pREV1*. Red intensity indicates the percentage of red fluorescence cells. The pie chart above shows the statistics of progeny cells for the representative data points marked by the thick black borders. **c**, Branching device design with added nonsense (NS) DNA sequences to adjust promoter arm lengths. **d**, Flow cytometry data showing differentiated cell distributions for circuits with left-to-right arm length ratios ranging from 1:10 to 10:1 (data are mean ± s.d. of *n* = 8 replicates). **e**, Schematic of cell fate branching device design using two *attP* site mutants with different recombination efficiencies. Mutated bases in *attP*_slow_ and *attP*_fast_ are shown in red. **f**, Flow cytometry data showing branched cell distributions for seven circuits with different *attP* mutants (data are mean ± s.d. of *n* = 8 replicates). **g**, Summary of five factors influencing the output cell differentiation devices. **h**, Comparison of progeny cell differentiation ratios from circuits incorporating each of the 5 selected factors, as described in Fig. 2g, showing that the first three factors significantly expand progeny cell distributions. **i**, Nine representative circuit designs used to validate the predictive accuracy of the formulaic model. **j**, Comparison between model-predicted Venus cell ratios (lines) and flow cytometry data (dots) from designs in Figure 2i.

To investigate steric hindrance further, we inserted Tet operator (*TetO*) sequences, specific binding sites for the tetracycline repressor (TetR), between *att* sites in *attB*-*attP* or *attP*-*attB* configurations (**Fig. S22a-b**). Next-generation sequencing (NGS) revealed that TetR binding decreased recombination efficiency, with the rate for *attB*-*attP* sites dropping from 61.0 ± 4.3% in the absence of TetR to 39.1% in its presence (**Fig. S22a**). Similar results were obtained with dCas9 and guide RNA targeting sequences between *att* sites (**Fig. S22c**). These findings indicate that steric constraints imposed by sequences between *att* sites affect recombination efficiency across various yeast promoters (**Fig. 2a**). Additionally, replacing promoters with 100 bp non-functional random DNA sequences of varying GC content revealed substantial variations in recombination frequencies, independent of GC content (**Fig. S23**). These results suggest the involvement of other unknown factors influencing Bxb1 recombination. As functional promoters are necessary for gene expression outputs, subsequent investigations focused on branching devices with one promoter on each arm.

#### Length of promoter arms between att sites

We next evaluated how the length of promoter-containing DNA sequence between *att* sites impacts recombination efficiency (**Fig. 2c**). We hypothesized that increasing DNA length would decrease recombinase tetramerization likelihood, influencing branching outcomes. By varying the length of DNA on one arm while maintaining a fixed length on the other, we observed a consistent impact on progeny cell proportions (**Fig. 2d**). Increasing the length of the right arm relative to the left (1:1 to 1:10) reduced the proportion of red fluorescent cells from 62.0 ± 0.7% to 17.0 ± 0.9% (**Fig. 2d, Fig. S24**). Conversely, elongating the left arm (1:1 to 10:1) increased red fluorescent cells to 96.2 ± 0.2% (**Fig. 2d, Fig. S25**). Fixed-length, nonfunctional DNA sequences showed relatively mild effects compared to DNA length differences (**Fig. S26**). These results confirm that DNA sequence length flanked by *att* sites modulates recombinase efficiency and progeny cell distribution.

#### att site mutants and recombination efficiency

Variants of Bxb1 *att* sites can directly impact the recognition and binding of the Bxb1 recombinase to the DNA molecule^26^. By exploiting the differential affinities of recombinases to distinct *att* sites with varying recombination kinetics, we hoped to selectively modulate the enzyme’s catalytic preference within branching devices, thus enabling tuning of sub-population ratios (as depicted in **Fig. 2e**). Our study employed an *attP*_slow_ variant, which contains a C14A mutation, and *attP*_fast_, featuring multiple substitutions (5A, 8T, 12C, 14T, and 17A) (**Fig. 2e**). Since *attP*_slow_ demonstrates reduced recombination efficiency compared to the more efficiently processed *attP*_fast_, we designed six distinct branching device configurations by enlisting wild-type and variant *att* sites to modulate recombination preferences (**Fig. 2f**, **Fig. S27**). The results verified that the configuration of *attP* variants effectively influenced progeny cell fluorescence ratios. For instance, the *attP*_fast_-*attB*-*attP*_slow_ arrangement yielded 68.3 ± 1.6% green fluorescent cells, while the *attP*_fast_-*attB*-*attP* configuration resulted in 44.3 ± 1.6%. The *attP*_slow_-*attB*-*attP*_fast_ setup produced 13.4 ± 0.9% green fluorescent cells (**Fig. 2f, Fig. S27**). These findings demonstrate the potential for *att* site variants to selectively program recombinase-based cell differentiation outcomes.

### **M**inor factors influencing the output of branching devices

#### Arrangement of att sites in the branching device

The general architecture of our branching devices included three *att* sites (**Fig. 1a**), resulting in 16 possible permutations of *att* type (*attB* or *attP*) and orientation (**Extended Data** Fig. 3b). Each configuration was tested in yeast (**Fig. S28-S29**), and all designs, including the prototype in **Fig. 1a**, consistently generated two distinct progeny cell types with minor variations in ratios, generally not exceeding 10% (**Extended Data** Fig. 3b). The proportion of green fluorescent cells fluctuated between 32.1 ± 1.1% and 42.9 ± 0.7% across all configurations. These results suggest that the specific arrangement and orientation of *att* sites only mildly impact recombination and differentiation outcomes. Notably, Bxb1-mediated DNA inversion was preferred over DNA excision events when both competed against each other (**Extended Data** Fig. 3b).

#### Orthogonal dinucleotides in att sites

The central dinucleotide of *att* sites generates a 2-bp overhang that mediates ligation between cleaved *attB* and *attP* sites with complementary overhangs^27^. While all devices in this study used the GA dinucleotide^28^, six alternative dinucleotides (GT, TT, CA, CC, CT, and TC as controls) were tested to assess their potential effects (**Extended Data** Fig. 3c). Each variant recombined exclusively with *att* sites containing complementary dinucleotides, allowing up to six independent recombination events with a single recombinase. However, when these dinucleotides were applied in the *attB-attP-attB* branching device, the distribution of red and green fluorescent cell populations remained consistent. For instance, the green fluorescent cell fraction ranged from 34.9 ± 0.5% and 37.3 ± 0.8% (**Extended Data** Fig. 3c, **Fig. S30**). These findings indicate that orthogonal *att* variants do not appreciably alter cell state ratios, supporting the integration of multiple orthogonal devices within a single genome to enable complex circuit designs.

#### Differential cell growth rates

To examine the impact of protein payloads and differential cell growth rates on progeny cell ratios, the Venus reporter was replaced with several recombinases (e.g., No67^29^, Si74^29^, TP901^30^, R4^30^, PhiC31^30^) (**Extended Data** Fig. 3d**, Fig. S31**). These replacements were incorporated into branching devices designed to produce a 1:1 ratio of Venus-to mScarlet-expressing cells, with full differentiation achieved after 8 hours (**Extended Data** Fig. 3d**, Fig. S32**). Following an 18-hour induction period, limited variation in progeny cell-type distribution was observed with R4 recombinase, resulting in 51.8 ± 0.6% R4-expressing and 48.2 ± 0.6% mScarlet-expressing progeny cells (**Fig. S33**). Conversely, devices containing No67 and mScarlet exhibited an increase in red fluorescent cells from 52.3 ± 0.5% at 8 hours to 57.6 ± 0.5% at 18 hours, suggesting minor toxicity of the No67 recombinase (**Fig. S34**). This effect was more pronounced with PhiC31 recombinase, where red fluorescent cells increased from 57.9 ± 1.5% at 8 hours to 94.3 ± 0.6% at 18 hours (**Fig. S35**). These observations highlight the potential to utilize differential growth rates to dynamically modulate progeny cell ratios over time (**Fig. S36**).

#### Choice of recombinases

To evaluate the generality of these observations with Bxb1-based devices, additional serine recombinases (e.g., TP901, R4, A118) were tested (**Extended Data** Fig. 3e). Flow cytometry analysis confirmed that all serine recombinases tested effectively induced differentiation, generating progeny cells in two distinct states (**Extended Data** Fig. 3e). However, significant disparities in progeny cell distributions were noted. For instance, the TP901-based system produced approximately 16.5 ± 0.2% green fluorescent cells and 79.4 ± 0.6% red fluorescent cells, with the remainder showing no fluorescence (**Fig. S37**). By contrast, the A118-based system yielded approximately 38% green, 39% red, and 23% undifferentiated cells (**Fig. S37**). When undifferentiated cells sorted from the A118-based system were re-induced in fresh YPD medium, a similar differentiation pattern re-emerged: 38% green, 39% red, and 23% undifferentiated cells (**Fig. S38a-c**). Similarly, partially induced Bxb1-based systems displayed repeatable differentiation patterns (**Fig. S38d-f**). These results indicate that potential construct asymmetric differentiation systems where specific founder cell ratios retain pluripotency, thereby prolonging system stemness.

### Developing a formulaic model for predicting cell differentiation outcomes

Our experimental measurements revealed the complex, multifaceted nature of cell differentiation outcomes (**Fig. 2g-h**). To achieve predictive control over cell differentiation ratios, we developed a simple, formula-based white-box model comprising five ordinary differential equations that capture the essential dynamics of Bxb1-mediated computation^26, 31, 32^. This model incorporates: (1) the differential cleavage of recombinases on DNA fragments, (2) transcription machinery recruitment, and (3) steric constraints (**Fig. S39**). Key parameters include: functional DNA elements flanked by *att* sites, the length of DNA sequences flanked by *att* sites, and the recombination efficiencies of *att* site variants. We parameterized the model by fitting the simulation results with experimental measurements (**Fig. S39-S40**, **Tab. S1-S3**). The model demonstrated strong predictive accuracy for novel experimental configurations (**Fig. 2i, Fig. S41**), as evidenced by the agreement between predicted and actual differentiation outcomes (R^2^ = 0.887, **Fig. 2j**). These results suggest that cellular differentiation behavior can be quantitatively programmed through the rational design of the recombinase circuit architecture.

### Precise ratio multiplication through parallel genetic devices

Having established that our branching devices can quantitatively control the ratio between two progeny cell types, we next asked whether multiple devices could be combined to enable complex, multi-lineage differentiation. This requires parallel branching circuits that are functionally orthogonal within the same genome by incorporating dinucleotide variants into the Bxb1 *att* sites (**Fig. S30**), thereby creating insulated recombination units^28^. Each device undergoes independent Bxb1-mediated recombination, producing progeny cells with distinct traits. The probability of generating each cell type is determined by multiplying the probabilities of individual states in the branching devices. For *n* devices, this approach yields *2^n^*unique cell types, each defined by specific trait combinations (**Fig. S42**). For example, with two devices producing outputs for green and red fluorescence, the probability of a progeny cell being yellow (expressing both green and red fluorescence) is calculated as:

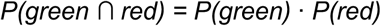

To validate the rule of multiplication, we integrated two orthogonal branching circuits into the yeast genomes: one using ShadowG and Venus, and another using ShadowG and mScarlet (**Fig. 3a**). Following Bxb1 induction, four progeny cell types emerged, corresponding to distinct fluorescence states: dark (no fluorescence), green, red, and yellow (both green and red fluorescence, **Fig. 3b**). Utilizing the probabilities measured from strains hosting only one of the two branching devices (**Fig. S43-S45**), we computed the theoretical distribution of the four progeny cell types in the final mixed population (**Fig. 3b**). Flow cytometry confirmed a strong correlation between the predicted probabilities and the observed cell type proportions (**Fig. 3c**). We also inoculated ∼1000 founder cells on an agar plate supplemented with the inducer and observed colonies with four-color stripes under fluorescence microscopy (**Fig. 3d**). Varying individual branching device designs yielded results reflecting the modified ratios, with the calculated progeny cell distribution closely resembling the experimental outcomes, highlighting the robustness of the rule of multiplication (**Fig. S46-S57**).

**Fig. 3.**
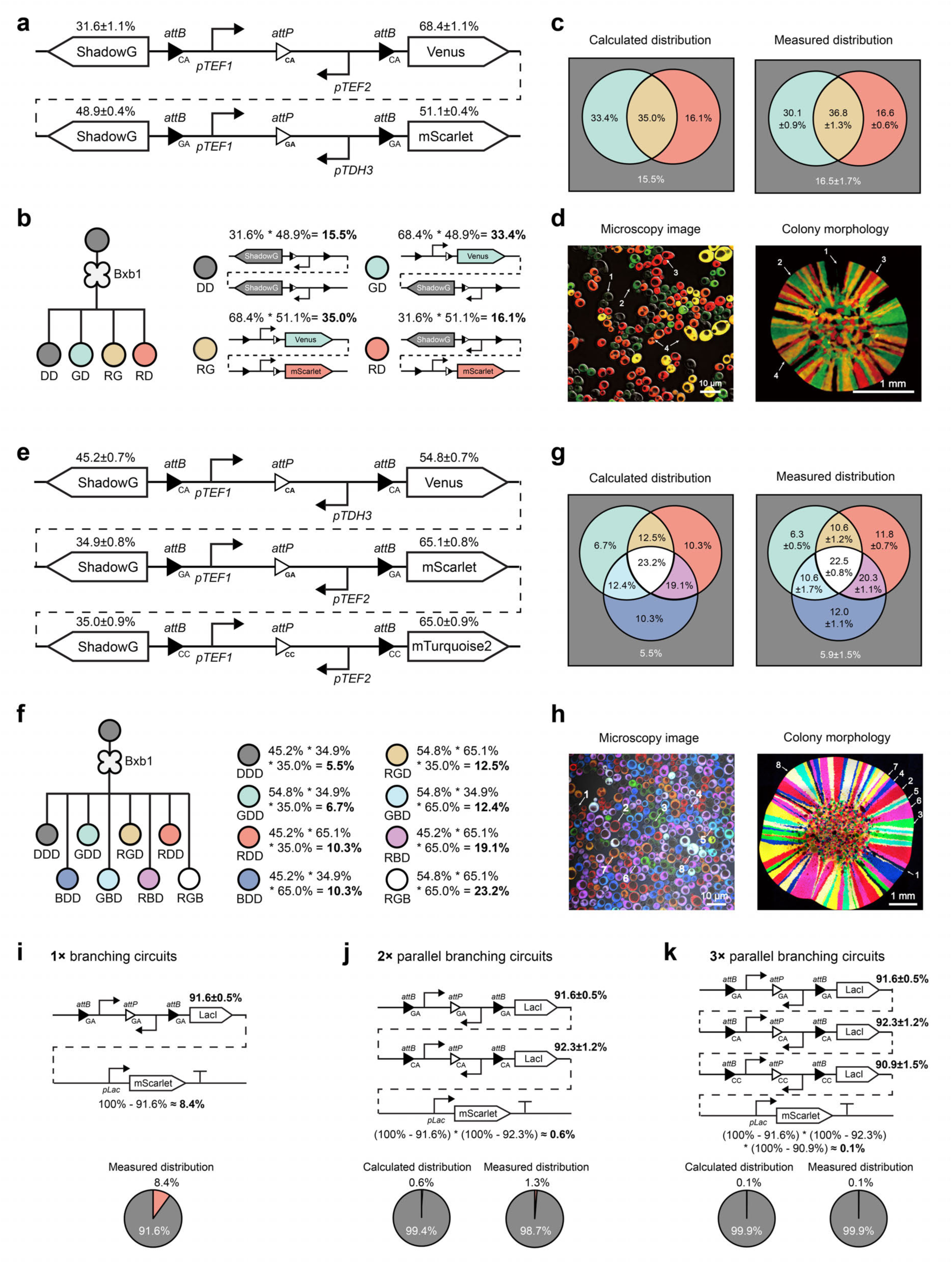
Designing parallel orthogonal genetic circuits for quantitatively controlled cell states. **a**, Schematic of a 2× orthogonal branching circuit design. The first circuit uses CA *att* sites with *pTEF1* and *pTEF2* promoters, and ShadowG and Venus as reporters. The second circuit uses GA *att* sites with *pTEF1* and *pTDH3* promoters, and ShadowG and mScarlet as reporters. Progeny proportions represent the probability of reporter expression. **b**, Theoretical progeny types and proportions from the 2× orthogonal circuit: **DD** (ShadowG only, non-fluorescent), **GD** (Venus + ShadowG, green fluorescence), **RG** (mScarlet + Venus, red and green fluorescence), and **RD** (mScarlet + ShadowG, red fluorescence). **c**, Comparison of calculated progeny distributions with flow cytometry results (data are mean ± s.d. of *n* = 8 replicates). **d**, Fluorescence microscopy images showing four distinct cell populations in solution (left) and colonies (right). **e**, Schematic of a 3× orthogonal branching circuit. **f**, Theoretical progeny types and proportions from the 3× orthogonal circuit. **g**, Comparison of calculated distributions with flow cytometry data (data are mean ± s.d. of *n* = 8 replicates). **h**, Fluorescence microscopy images showing eight distinct cell populations in solution (left) and colonies (right). **i**, Schematic design and flow cytometry data of a 1× cell branching circuit. **j**, Schematic design, calculated distribution, and flow cytometry data for a 2× parallel cell branching circuit. **k**, Schematic design, calculated distribution, and flow cytometry data for a 3× parallel cell branching circuit.

Expanding the system to three orthogonal branching devices (**Fig. 3e**) yielded eight progeny types. For instance, the probability of generating white cells expressing all three fluorescent proteins (green, red, and blue) was computed as:

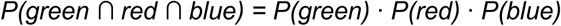

Both fluorescence microscopy and flow cytometry confirmed that the experimental distributions of eight progeny cell types matched their theoretical predictions (**Fig. 3f-g**, **Fig. S58**). Fluorescence microscopy further demonstrated all eight fluorescent states in both liquid and solid media (**Fig. 3h**). By modifying branching device parameters, various cell-type distribution ratios were achieved (**Fig. S59-S62**). In all configurations, calculated distributions aligned closely with experimental findings, confirming the robustness of the rule of multiplication and its broad applicability.

### Stacking parallel branching devices to achieve extreme ratios

Thus far, the lowest ratio achieved with a single branching device was 3.6%, modulated by arm length (**Fig. 2d**). Given that parallel branching devices enable robust multiplication computation, we hypothesize that one can achieve <3.6% during cell differentiation using the previously demonstrated rule of multiplication. By deploying analogous branching devices in parallel within a single founder strain, we implemented exponential functions to generate progeny populations with extremely low ratios.

We designed a branching device utilizing the transcriptional repressor LacI. In this system, LacI expression occurs ∼91.6% of the time post-differentiation. When mScarlet was placed under the control of *pLac* promoter (which activates only in the absence of the LacI repressor), the probability of progeny cells expressing red fluorescence was calculated as:

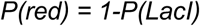

Experimentally, this yielded about 8-9% red fluorescent progeny, validating the model (**Fig. 3i, Fig. S63a-c**).

When two orthogonal repressor-based branching devices were integrated into the same genome, the probability of progeny cells with no inhibiting mScarlet expression was:

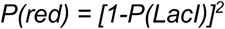

This computation predicted a 1% red fluorescent progeny ratio, confirmed experimentally (**Fig. 3j, Fig. S63d)**. Adding a third analogous device further reduced the likelihood of red fluorescent progeny to:

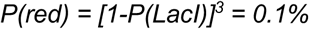

This prediction also matched experimental results (**Fig. 3k, Fig. S63e**).

We envision that this “exponential function” can be easily expanded by installing up to six analogous, orthogonal Bxb1-based branching devices with different dinucleotides across various chromosomes to avoid homology-induced looping out, achieving ratios in the 10^-6^ range. Notably, replacing the LacI repressor with the reverse tetracycline transactivator rtTA^33^ also enables comparable multiplicative operations (**Fig. S64**), indicating the universality of such designs. In scenarios where a very small subpopulation begins multicellular programs, tight regulation of its initiation frequency is essential. Conventional genetic circuits, which depend on a few layers of gene expression events or protein-protein interactions, often lack the precision needed to trigger program onset at rates of the thousandth or millionth^34^. Our approach thus enables robust control over low-frequency events like pattern formation, morphogen gradient initiation, or fail-safe population control mechanisms^35^.

### Branching devices in series for programming sequential differentiation

Multicellular organism development is a hierarchical process in which development involves stem cells undergoing sequential differentiation into specialized progeny, guided by external cues that enable adaptation to environmental conditions. To replicate this process, we engineered branching devices that allow cells to differentiate sequentially into predefined populations in response to specific signals. These devices use cascading recombinase-based circuits, each inducible by small molecules, to program sequential cellular functions.

**Fig. 4a** and **Fig. 4b** illustrate our design, featuring two branching devices in series activated by distinct recombinases, Bxb1 and TP901. To minimize leaky expression of sensitive regulatory proteins (e.g., recombinases, transcription factors), which can disrupt branching circuits even at low levels (**Fig. S65**), terminators were placed between *att* sites. The first branching device uses β-estradiol-inducible Bxb1 to direct founder cells into two progeny types: those constitutively expressing mTurquoise2 (blue) and those retaining an inducible TP901 cassette (dark). TP901 subsequently governs the differentiation of dark cells into green or red fluorescent cells upon aTc induction.

**Fig. 4.**
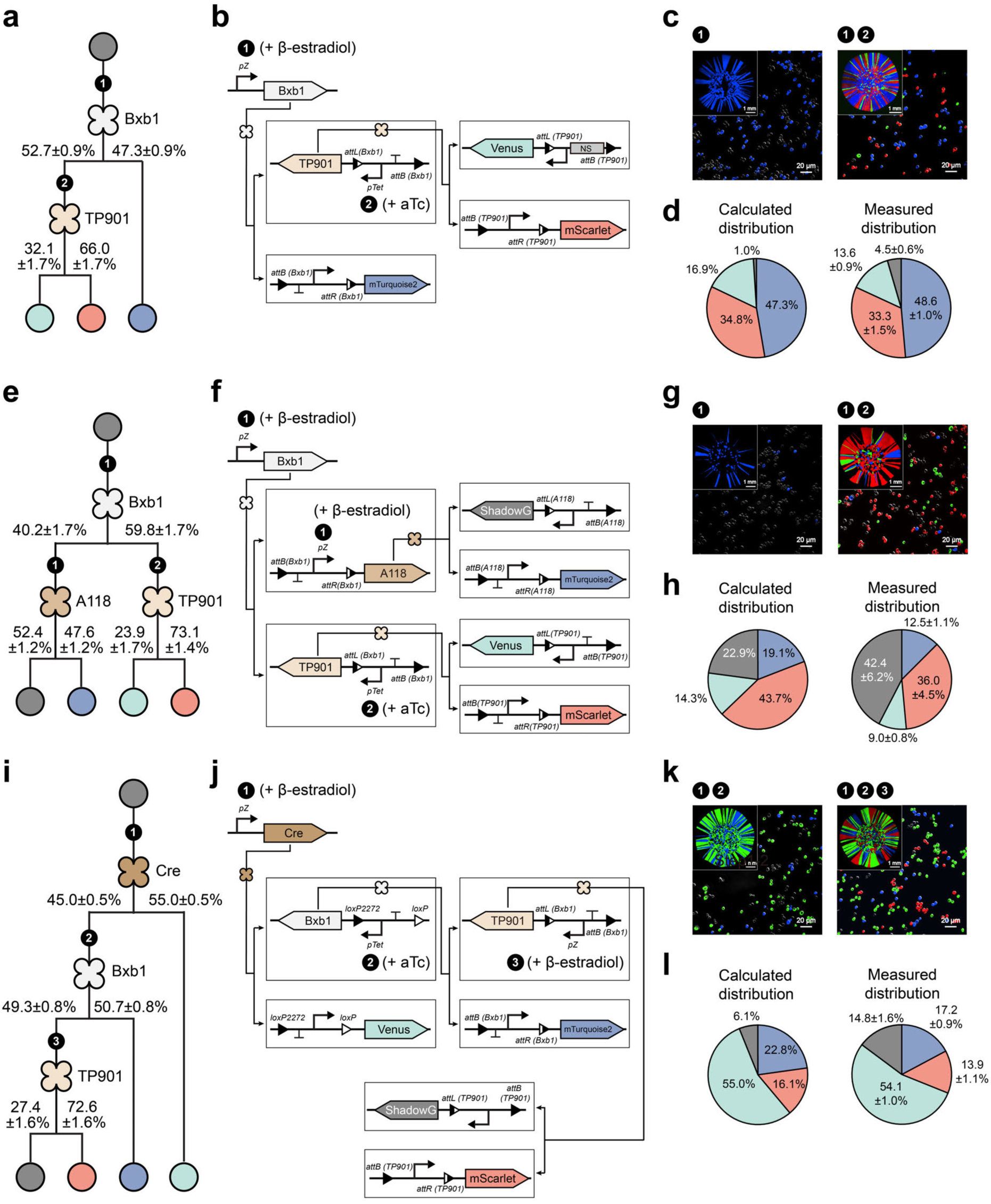
Sequential genetic circuits for controlled creation of diverse cell states. **a**, Schematic of a 2-layer branching device using 2 recombinases to sequentially differentiate cells into 3 terminal cell types. Experimentally determined progeny proportions were used to calculate expected circuit outputs. **b**, Circuit changes during 2-step sequential differentiation. **c**, Fluorescence microscopy images displaying cell population changes at each step. **d**, Comparison of calculated progeny distributions with flow cytometry data (data are mean ± s.d. of *n* = 8 replicates). **e**, Schematic of a 2-layer branching device using 3 recombinases to produce 4 progeny types. **f**, Circuit changes during 2-step differentiation. **g**, Fluorescence microscopy images displaying cell population changes at each step. **h**, Comparison of calculated distributions with flow cytometry data (data are mean ± s.d. of *n* = 8 replicates). **i**, Schematic of a 3-layer branching device using 3 recombinases to differentiate into 4 progeny types. **j**, Circuit changes during 3-step sequential differentiation. **k**, Fluorescence microscopy images displaying cell population changes at each step. **l**, Comparison of calculated distributions with flow cytometry data (data are mean ± s.d. of *n* = 8 replicates).

In the presence of β-estradiol alone, 52.7% of the progeny remained dark, while 47.3% fully differentiated into blue cells (**Fig. 4c, left**). Dark progenitors retained multipotency, differentiating into green and red cells only after aTc induction (**Fig. 4c, right**). The final proportions of green and red cells were determined by multiplying the ratio of dark cells at the first layer by their respective differentiation ratios at the second layer:

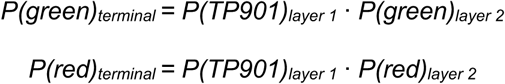

The calculated ratios, 16.9% green and 34.8% red, closely matched the experimental results (**Fig. 4d, Fig. S66-S67**). Minor discrepancies in the dark cell population were attributed to incomplete differentiation, leaving residual stem cells (**Fig. S37**). This validated system enabled the engineering of a founder strain capable of stepwise differentiation into three cell types secreting laccase, Mel1, and lipase, forming a yeast consortium for the degradation of multiple water contaminants (**Fig. S68**). Furthermore, transcription factors were shown to function with these devices, facilitating the sequential execution of genetic programs with multiple transcriptional units (**Fig. S69**).

We evaluated the scalability of these designs in both horizontal (output diversity) and vertical (circuit depth) expansions. A two-layer binary branching circuit was constructed, resulting in four cell types upon full differentiation (**Fig. 4e**). The first branching device, activated by β-estradiol-inducible Bxb1, generated two intermediate progenitor types. One type expressed A118, directing differentiation into dark (ShadowG) or blue (mTurquoise2) cells via second branching devices. The other type expressed TP901 in the presence of aTc, driving differentiation into green (Venus) and red (mScarlet) cells (**Fig. 4f-h**). Although recombination inefficiencies in TP901 and A118 led to an increased number of dark cells (**Fig. S37-38**), the observed distributions aligned with predictions (**Fig. 4h, Fig. S70**).

To mitigate inefficiency propagation in deeper circuits, we incorporated the highly efficient Cre recombinase into a three-layer series design (**Fig. 4i**). Each layer contained one branching device responsive to sequential inputs (β-estradiol → aTc → β-estradiol), producing four terminal cell types (**Fig. 4j, Fig. S71**). The ratio of red cells in the final cell population was defined as:

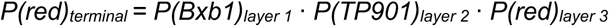

**Fig. 4k** and **Fig. 4l** show that replacing A118 with Cre greatly reduced recombination inefficiencies. No observable leakiness occurred across layers, and reusing β-estradiol as an input greatly expands the versatility of this series design under a limited number of inducer species.

In summary, we demonstrated that sequential differentiation through branching devices arranged in series provides high programmability and control, akin to natural morphogen-driven processes. This platform offers a robust framework for implementing complex genetic circuits with applications in synthetic biology and cellular engineering.

### From a single strain to a consortium of synergistic members

So far, we have shown that inducible cellular differentiation programs can produce multiple cell types at specific ratios from a single founder strain (**Fig.1-4, Movie S1-S3**). While manually growing and mixing multiple strains into artificial co-cultures may seem convenient in the lab, it also demands professional personnel and equipment^36^. Our *in situ* differentiation approach eliminates such needs, streamlining workflows for austere environments or use cases inside the human body, where manual mixing is not easily achievable. To illustrate this, two multi-gene biosynthetic pathways were selected to produce pigments: violacein and β-carotene. Violacein, a purple bis-indole pigment, is synthesized by five enzymes: VioA, VioB, VioC, VioD, and VioE^37^ (**Fig. 5a**). β-carotene, a vibrant orange pigment, is produced by four enzymes: CrtI, CrtE, CrtYB, and tHGM1^37^ (**Fig. 5a**). By fine-tuning the ratios of the two cell types synthesizing these pigments, a spectrum of colors was created by blending these colors.

**Fig. 5.**
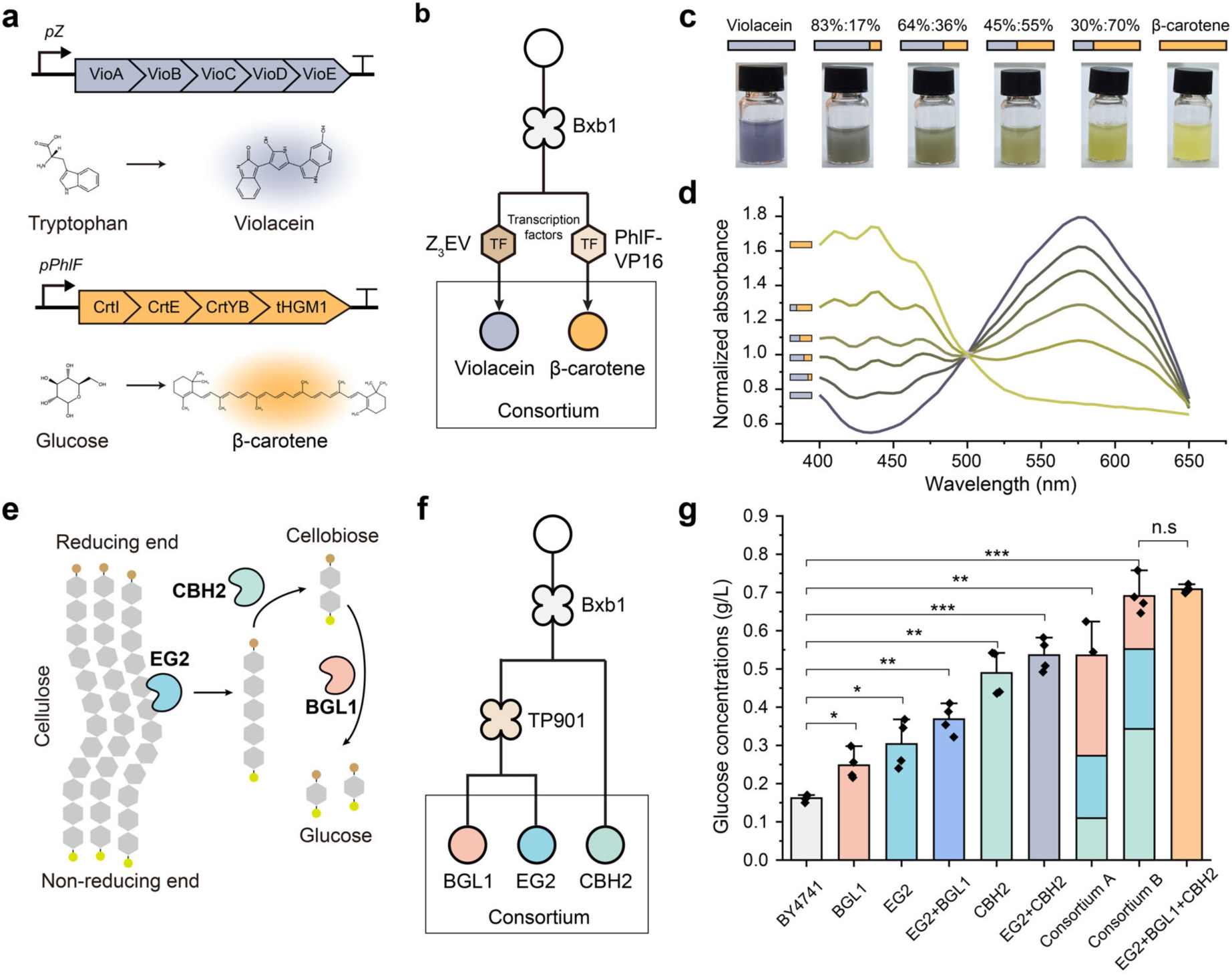
Engineering single-cell-controlled consortia for metabolic engineering. **a**, Diagram of enzymes involved in violacein (from tryptophan) and β-carotene (from glucose) synthesis. **b**, Founder cells differentiate into two progeny cell types producing either violacein or β-carotene enzymes, controlled by Z_3_EV or PhIF-VP16 transcription factors, respectively. **c**, Photographs of yeast cultures producing violacein, β-carotene, or mixtures at varying ratios. **d**, Absorbance spectra of extracted pigments, with values normalized to the absorbance at 500 nm. **e**, Diagram of cellulase enzymes (CBH2, EG2, and BGL1) breaking down cellulose into glucose. **f**, Controlled generation of yeast consortia secreting three cellulases using branching devices. **g**, Comparison of reduced sugar production from cellulose by single-strain monocultures versus consortia (data are mean ± s.d. of *n* = 4 replicates). Consortium A and Consortium B featured differentiated populations producing BGL1, EG2, and CBH2 in ratios of 5:3:2 and 2:3:5, respectively. Statistical analysis was performed using unpaired two-tailed Student’s t-tests; n.s., not significant, *\*P* < 0.05, *\*\*P* < 0.01, *\*\*\*P* < 0.001.

To control these pathways, two synthetic transcription factors, Z_3_EV and PhIF-VP16, were used to regulate violacein and β-carotene biosynthesis, respectively (**Fig. 5b, Fig. S72a**). A single branching device differentiated the founder strain into two specialized strains: one using Z_3_EV to drive violacein production and the other using PhlF-VP16 to trigger β-carotene biosynthesis. Adjusting the proportions of these two “pixel” cell types resulted in cultures that displayed colors ranging from pure purple to pure orange, with four intermediate colors (**Fig. 5c, Fig. S73**). Extracted dyes showed absorbance profiles matching their genetically encoded pigments (**Fig. 5d, Fig. S72b**). This “genetic color palette” approach reduces metabolic stress on individual cells through the division of labor (**Fig. S74**) and demonstrates its potential for designing self-pigmenting living materials^38^. Moreover, we rapidly constructed a small library of 25 synthetic differentiation circuits, each harboring a distinct engineered *attP* variant that modulates the recombination efficiency of Bxb1 recombinase, using one-pot Golden Gate assembly^25,39^ (**Fig. S75a-b**). Next, we randomly selected 96 yeast colonies, induced their differentiation, and observed that the circuits produced varied ratios of violacein-and β-carotene-producing cells, leading to visibly different pigmentation (**Fig. S75**). We could determine population composition by analyzing the RGB values of images in a high-throughput way (**Fig. S75**), which shows the possibility of using pooled branching circuits to find the best population ratio for user-defined functions.

Branching devices were also applied to cooperative enzymatic reactions relevant to industry. A cellulose-degrading system was chosen as a model, involving three cellulases—EG2, CBH2, and BGL1^40^, each performing a step in breaking down cellulose into glucose (**Fig. 5e**). These enzymes were integrated into a two-layer sequential genetic circuit controlled by recombinases Bxb1 and TP901, directing the founder strain to differentiate into three distinct cellulase-secreting progeny cell types (**Fig. 5f**). The performance of these engineered consortia was evaluated by measuring glucose production from engineered yeast supernatants degrading Avicel-101 using the DNS method^41^ (**Fig. S76**). Two consortia with different cellulase-secreting cell ratios were tested: Consortium A (BGL1: EG2: CBH2 at 5:3:2) and Consortium B (2:3:5). Both consortia showed significantly higher glucose production compared to the wild-type BY4741 strain (**Fig. 5g**). Consortium B, with optimized enzyme ratios distributed across specialized cell types, achieved similar cellulose degradation efficiency to a control strain that constitutively co-expressed all three cellulases but showed slow growth (**Fig. 5g** and **Fig. S77**). These results highlight how fine-tuning cell-type ratios can enhance consortium efficiency. The universality of branching devices across different genetic systems makes them a powerful tool for engineering functionally diverse consortia. These consortia, derived from a single founder strain, can perform complex tasks and processes, offering new possibilities for synthetic biology and industrial biotechnology.

### Controllable self-organizing morphogenesis programmed by branching devices

The relative ratios and spatial arrangement of cell types, shaped by differentiation and cell-cell interactions, play a key role in the development of multicellular structures and morphogenesis^23, 24, 42^. We hypothesized that branching devices could be used to program multicellular morphologies beyond simple co-cultures created by manual mixing. To validate this, we combined yeast surface display systems (e.g., SED1 and GPI-anchored proteins)^21, 43^ and customized CAMs, such as antigen-antibody complexes^22^, as final outputs for the branching devices.

As depicted in **Fig. 6a** and **Fig. 6b**, synthetic transcription factors Z_3_EV and PhlF-VP16 were used to drive the expression of seven highly specific, orthogonal CAM pairs. Upon induction, this architecture enables engineered yeast to differentiate into two unique cell types, each displaying one of the CAM pairs, which facilitates the self-assembly of cell aggregates. To visualize these structures, green and red fluorescent proteins were co-expressed with the CAMs (**Extended Data** Fig. 4).

**Fig. 6.**
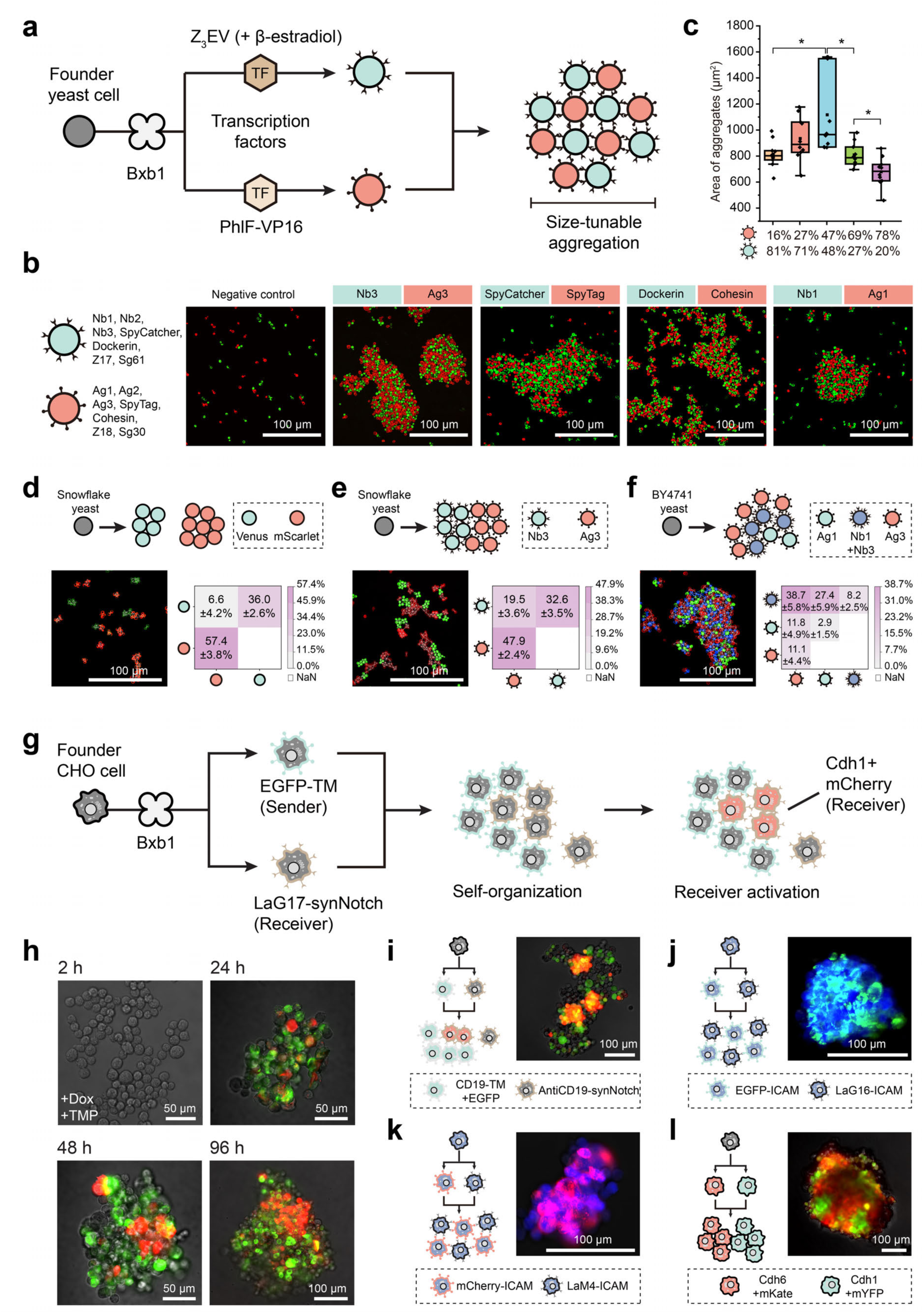
Engineering self-organized cellular morphogenesis via designed branching devices. **a,** Schematic of founder yeast cells differentiating into two progeny types displaying specific CAM, driving cellular pattern formation. **b**, CAM selection and fluorescence microscopy images of yeast consortia forming aggregates facilitated by CAM pairs (Nb3-Ag3, Nb1-Ag1, SpyTag-SpyCatcher, and Dockerin-Cohesin); negative control is the fluorescent BY4741 yeast cells lacking CAM-expression. **c**, Comparison of aggregate sizes with different ratios of Nb3-and Ag3-displaying cells (data are mean ± s.d. of 10 independent images from 3 independent experiments). **d**, Phase-separated cellular morphologies formed by snowflake yeast. Heatmaps depict the frequency of direct contacts between red- and green-labeled cells. Data represent mean ± s.d. from *n* = 4 biologically independent replicates. **e**, Engineered snowflake yeast expressing Ag3 (red) and Nb3 (green) surface adhesion proteins form heterotypic aggregates through Ag3-Nb3 interactions. Heterotypic interactions promote increased red-green cell associations. Data represent mean ± s.d. from *n* = 4 biologically independent replicates. **f**, A layered morphology emerges when the blue progeny (Nb1+Nb3) simultaneously adheres to both red (Ag3) and green (Ag1) cell types, creating a structured tri-lineage assembly. Quantitative image analysis reveals specific interaction patterns based on orthogonal adhesion pairing. Data represent mean ± s.d. from *n* = 4 biologically independent replicates. **g**, Schematic showing founder CHO-K1 cell differentiation into two progeny types driving cellular pattern formation.TM: PDGFRβ transmembrane domain. **h**, In situ cellular differentiation and pattern formation initiated from 100 founder cells upon induction with Dox and TMP. mCherry expression remains spatially confined to cell-cell interfaces. **i**, Bxb1-mediated differentiation programs multicellular patterning in CHO-K1 cells via CD19-antiCD19 synNotch contact-dependent signaling. **j**, Differentiated progeny expressing EGFP or LaG16 on the cell surface self-organize into multicellular aggregates through heterophilic adhesion between EGFP and LaG16. **k**, Differentiated progeny expressing mCherry or LaM4 form multicellular aggregates via heterophilic adhesion between mCherry and LaM4. **l**, Differentiated progeny expressing Cdh6 or Cdh1 assemble into structured multicellular aggregates via homophilic cadherin-mediated adhesion. Statistical analysis was performed using unpaired two-tailed Student’s t-tests; *\*P* < 0.05.

Microscopy confirmed the successful display of the CAM pairs on yeast cell surfaces, with four pairs— Nb3-Ag3, Nb1-Ag1, Spytag-SpyCatcher, and Dockerin-Cohesin—showing strong aggregation-promoting properties (**Fig. 6b, Extended Data** Fig. 4). Flow cytometry further revealed elevated doublet ratios (exceeding 5%) for these four pairs, indicating a higher aggregation tendency in solution (**Extended Data** Fig. 4). Using Nb3-Ag3 as a model, we investigated how altering cell-type ratios affected aggregate size. Adjusting the ratio of Nb3-to Ag3-expressing cells from 1:4 to 4:1 significantly influenced aggregate size, with more balanced ratios (e.g., 1:1) producing larger aggregates compared to imbalanced ones (e.g., 1:4 or 4:1) (**Fig. 6c, Fig. S78**). Additionally, sedimentation rates were higher for aggregates with balanced progeny ratios, indicating distinct physical properties (**Fig. S79**).

To further explore programmed spatial self-organization, we leveraged the intrinsic homo-adhesion property of snowflake yeast, which spontaneously forms multicellular aggregates^44^. Here, a single founder differentiated into two distinct lineages (red and green), which then self-organized into aggregates with minimal heterotypic mixing (**Fig. 6d, Extended Data** Fig. 5a-d). Engineering the differentiated snowflake progenies to display either Ag3 or Nb3 on their surfaces introduced hetero-adhesion in addition to homo-adhesion (**Fig. 6e, Extended Data** Fig. 5e-h). Although snowflake-based self-sorting remained the dominant morphological driver, enhanced heterotypic associations were observed, rising from 6.6 ± 4.2% to 19.5 ± 3.6% (**Fig. 6d-e**). Furthermore, we built a two-layered sequential branching device in BY4741 yeast cells, generating three progeny cell types: red (displaying Ag3), green (Ag1), and blue (Nb1+Nb3) (**Fig. 6f, Extended Data** Fig. 5i). The blue progeny, designed to act as a “double-sided tape”, bound to both red and green cells, as mediated by the Ag-Nb pairs. This strategy promoted heterotypic clustering, with red-blue interactions reaching 38.7 ± 5.8% and green-blue interactions at 27.0 ± 5.9% (**Fig. 6f, Extended Data** Fig. 5i-l). These findings validate the expected adhesion logic and further support the concept that spatial cell-cell organization can be precisely directed via recombinase-mediated branching devices.

To demonstrate the applicability of the developed recombinase-based differentiation system beyond yeast, we engineered analogous branching devices in CHO-K1 mammalian cells using the PhiC31-assisted landing pad integration strategy, digital PCR-based copy number validation, and lentiviral delivery of doxycycline (Dox)-inducible recombinase cassettes (**Extended Data** Fig. 6). To minimize background recombinase activity, we fused Bxb1 to a dihydrofolate reductase (DHFR) degron^45^, achieving tight control with background recombination below 1% after five passages spanning 15 days (**Fig. S80**). We next imported the synNotch signaling system, enabling programmable, contact-dependent multicellular patterning^23, 46^ (**Fig. 6g, Extended Data** Fig. 7). Specifically, Bxb1-mediated in situ differentiation produced two distinct progeny cell types: “sender” cells expressing membrane-tethered EGFP (EGFP-TM) via the PDGFRβ transmembrane domain^23, 46^, and “receiver” cells expressing the cognate LaG17-SynNotch receptor fused to a Gal4-VP64 transcriptional activator. All cells additionally contained a Gal4-responsive cassette driving co-expression of Cadherin-1 (E-cadherin) and mCherry (**Fig. 6g, Extended Data** Fig. 7a). In the absence of induction, the system remained inactive with no detectable fluorescence (**Extended Data** Fig. 7b). Upon induction with Dox and trimethoprim (TMP), which stabilizes the DHFR degron to prevent Bxb1 degradation, EGFP-positive sender cells began to emerge. This led to selective mCherry expression in receiver cells upon direct contact with sender cells, resulting in localized cell aggregation over 96 hours (**Fig. 6h, Extended Data** Fig. 7b). Control experiments showed that sender cells expressing cytoplasmic EGFP failed to activate synNotch signaling, confirming that membrane localization and direct cell-cell contact were both necessary and sufficient to trigger this circuit^46^ (**Extended Data** Fig. 7c-f). Moreover, the generality of this approach was validated using an alternative synNotch pair (CD19-antiCD19)^23^, which also successfully mediated contact-dependent activation and multicellular assembly (**Fig. 6i, Extended Data** Fig. 7g-j). Finally, to demonstrate the versatility of our platform, we validated additional adhesion-based morphogenetic circuits. For example, recombinase-mediated differentiation generated complementary progeny expressing matched adhesion pairs, such as LaG17-EGFP^47^ (**Fig. 6j, Extended Data** Fig. 8) and LaM4-mCherry^47^(**Fig. 6k, Extended Data** Fig. 9), enabling the formation of mixed, tightly integrated cell aggregates. In another case, branching device driving expression of classical cadherins like Cdh6 and Cdh1 resulted in self-sorting cell clusters that organized into distinct spatial patterns (**Fig. 6l**, **Extended Data** Fig. 10).

In summary, we demonstrated that branching devices combined with CAMs can generate complex, tunable morphologies in both yeast and mammalian cells. Most importantly, these structures were assembled autonomously through cell differentiation from one founder strain, without external intervention such as manual mixing. With advancements in CAM technology across species, branching devices hold great promise for tissue engineering and organoid development.

## Discussion and Conclusion

Synthetic biology offers genetic tools with greater flexibility and predictability in controlling cell fate compared to natural systems, enabling the design of tailored cellular behaviors and more intricate functional networks that may not arise in nature. As a result, recent research has focused on using genetic circuits to emulate stem cells’ asymmetric division and guide the differentiation of microbial communities or diverse eukaryotic tissues and organs^6–10, 48^. Among these efforts, recombinase-based systems have played a prominent role. Seminal work by Roquet et al., for example, demonstrated how recombinase logic can be used to construct genetic state machines that record the identity and temporal order of input signals, thereby driving irreversible transitions to terminal cell states^18^. However, such systems primarily function at the single-cell level and are optimized for lineage tracing or memory storage. A key unresolved challenge is how to autonomously establish a balanced, quantitative relationship between functionally diverse progeny cell types within a consortium^49^. To surmount this challenge, we have developed highly programmable and robust cell fate branching devices using recombinases. By exploiting the differential recombination efficiencies of these enzymes on user-defined DNA sequences, we have achieved precise quantitative control over binary cell fate branching and population composition. Integrating these orthogonal branching devices into various circuit topologies, we have crafted an expandable platform for parallel and series gene circuits, enabling precise ratio manipulation and effective management of progenitor and progeny diversity. Moreover, a data-driven computational model provides predictive insights, guiding the construction of cell differentiation programs that generate multiple cell types from a single founder strain in user-defined ratios.

To showcase the significance of ratio distribution, we applied branching device outputs to scenarios requiring specific consortia ratios, such as self-pigmentation, cellulose degradation, and autonomous cell assemblies. Previously, these applications relied on manual mixing of cell types for optimal performance^22, 36, 40, 50^. Our approach eliminates this manual step by condensing multiple genetic programs into a single founder strain capable of autonomous or conditional differentiation in response to external cues (**Mov. S1-S5**). Although our demonstrations featured two or three progeny cell types (**Fig. 5 and Fig. 6**), the platform is highly modular and expandable. For example, in **Figs. 3f** and **3g**, we demonstrated that a single founder strain could differentiate into eight progeny types, each expressing a unique combination of three fluorescent proteins. By replacing these fluorescent proteins with transcription factors and using them to activate genetic programs controlled by 3-input AND gates^51, 52^, one could create eight progeny cell types, each performing pathway-level tasks, as seen in **Fig. 5a**. The exponential scalability (*2^n^*) of our parallel branching framework (**Fig. 3**), combined with advances in synthetic promoter design, suggests that systems with sixteen or more cell types are achievable.

Another promising direction involves engineering dependencies between progenitor and progeny cell types to create functionally and morphologically diverse cell assemblies^53, 54^. In this study, we demonstrated that direct-contact signaling (e.g., synNotch)^46, 55, 56^ can effectively trigger conditional differentiation. Building on this foundation, future integration of additional signaling modalities, such as soluble communication^57^, could further enable sequential, context-dependent differentiation for applications in organoids, tissue engineering, and metabolic engineering. Additionally, our framework can complement other artificial differentiation strategies utilizing asymmetric cell division^7, 9^, or quorum sensing^58–60^, to further enhance consortium stability. Notably, our system reserves a subpopulation of undifferentiated or partially differentiated progenitors due to incomplete recombination, which can be re-exposed to inducers to restore differentiation potential. Such a feature could be further optimized through programmed asymmetric division, as demonstrated in bacterial systems^7–9^.

One challenge in our system is the high sensitivity of population ratio to growth rate variations among progeny cell types with various metabolic burdens (**Fig. S36**). Incorporating feedback mechanisms or quorum-sensing-based devices could help maintain ratios over longer timescales. Less toxic recombinases and leakiness-free expression control can also significantly boost the robustness of the branching devices. In conclusion, our branching device offers a modular, expandable, and rapid approach to programmable ratio-specific cell differentiation and pattern formation. Their universality across diverse organisms and independence from pre-established morphogen or quorum presence make them a versatile tool for applications ranging from metabolic engineering to tissue development, both in liquid and solid phases.

## Methods

### Bacterial strains and culture medium

All bacterial cloning experiments were performed using *E. coli* DH5α chemically competent cells (Tsingke). For genetic circuit demonstration in *E. coli*, the DH10B strain (Tsingke) was used. *E. coli* cultures were grown in lysogeny broth (LB) medium at 37 °C with shaking at 220 rpm. To maintain plasmids, LB medium was supplemented with antibiotics at the following final concentrations: chloramphenicol at 34 μg/mL, carbenicillin at 100 μg/mL, kanamycin at 50 μg/mL, spectinomycin at 100 μg/mL, and zeocin at 100 μg/mL.

### Yeast strains and growth media

The *S. cerevisiae* strain BY4741 (MAT*a his3*Δ*1 leu2*Δ*0 met17*Δ*0 ura3*Δ*0 sed1*Δ) purchased from Horizon Discovery served as the default chassis for all experiments. Cultures were grown in yeast extract peptone dextrose (YPD) medium, consisting of 1.5% (w/v) yeast extract (Angel Yeast), 2.0% (w/v) peptone (Angel Yeast), and 2% (w/v) glucose (Sangon). Circuit induction and growth curve measurements used the YPD medium. For auxotrophy selection, SD minimal media (SD-Ura, SD-Leu, SD-His, and SD-Ura-Leu-His) containing 2% agar (Coolaber) were employed. Antibiotic selection used SD medium supplemented with 500 μg/mL hygromycin or 300 μg/mL zeocin. Snowflake yeast was created by deleting *ACE2* using a CRISPR-Cas9 system, as previously reported^44^.

### Mammalian cell strains and culture medium

Human embryonic kidney 293FT cells (Invitrogen, R70007, Thermo Fisher Scientific) were cultured in high-glucose Dulbecco’s Modified Eagle Medium (DMEM, Gibco), supplemented with 10% fetal bovine serum (FBS, Gibco) and 1% (v/v) penicillin-streptomycin (10,000 U/mL penicillin, 10 mg/mL streptomycin, Gibco), at 37 °C with 5% CO₂. Cells were maintained in 10 cm cell culture dishes and passaged at a 1:3 ratio every 2 days. Chinese Hamster Ovary (CHO-K1) cells (National Infrastructure Cell Line Resource, NICR) were cultured in F-12K medium (Kaighn’s modification of Ham’s F-12, M&C GENE TECHNOLOGY LTD), supplemented with 10% FBS (Gibco) and 2% (v/v) penicillin-streptomycin under the same incubation conditions. CHO-K1 cells were passaged at a 1:6 ratio every 3 days. Depending on the experimental purpose, the medium was supplemented with antibiotics at the following final concentrations to facilitate cell selection: 800 μg/mL hygromycin, 400-1000 μg/mL zeocin, 20 μg/mL blasticidin S, and 4 μg/mL puromycin.

### Plasmid construction and single-copy maintenance in E. coli

To ensure that the branching device remained present as a single copy in *E. coli*, we utilized a bacterial artificial chromosome (BAC) vector backbone^18^. The inherently low copy number of BAC plasmids minimizes potential interference caused by multiple plasmid copies, thereby providing a more stable and controlled environment for the genetic circuit. All genetic elements, including fluorescent protein genes, insulators, terminators, and promoters, were efficiently assembled with the BAC backbone using the one-pot Gibson assembly method. To achieve precise control over recombinase expression, we employed a tightly regulated anhydrotetracycline (aTc)-inducible system for regulated expression of the Bxb1 recombinase^61^. Sanger sequencing (Tsingke), and whole-plasmid sequencing (KBSeq, Sangon Biotech) were employed for plasmid sequence verification. Detailed information on specific genetic parts, protein sequences, plasmid backbones, and representative constructs can be found in **Tables S4-S7**.

### Plasmid construction and single-copy integration in S. cerevisiae BY4741

For *S. cerevisiae* (BY4741, haploid), we integrated the genetic circuit directly into the yeast genome to ensure single-copy presence in each cell. The plasmids were constructed using two modular cloning systems: the YTK system^25^ (Dueber lab) and the MYT system^62^ (Tom Ellis lab), both employing Golden Gate assembly. Assembly reactions were prepared as follows: 6.5 μL of DNA inserts, 1.0 μL of plasmid backbone, 1.0 μL of T4 DNA Ligase buffer (New England Biolabs), 0.5 μL of T4 DNA Ligase (New England Biolabs), 1.0 μL of a restriction enzyme, with the total volume adjusted to 10 μL. Restriction enzymes *Bsa* I (BsaI-HF v2, NEB) or *BsmB* I (*BsmB* I v2, NEB) were used. Thermocycling conditions consisted of 80 cycles of digestion (*Bsa* I at 37 °C or *BsmB* I at 42 °C for 5 minutes) and ligation (16 °C for 5 minutes), followed by digestion at 55 °C for 1 hour and heat inactivation at 80 °C for 10 minutes. Assembled plasmids were transformed into *E. coli* and cultured on LB agar with the appropriate antibiotic. Verification of plasmid DNA was performed via Sanger sequencing (Tsingke) and whole plasmid sequencing (KBSeq, Sangon Biotech). Genetic parts and protein sequences are listed in the **Table. S4 and Table. S5,** while plasmid backbones and representative constructions are listed in **Table. S6 and Table. S7**.

### Plasmid construction and single-copy integration in HEK293FT cells

To engineer HEK293FT cells for genomic integration of synthetic circuits, we first constructed a “landing pad” plasmid containing a single-copy PhiC31 *attB* site (cgcgcccggggagcccaagggcacgccctggcac) flanked by Rogi1-targeting homology arms and a puromycin resistance cassette. A sgRNA targeting the Rogi1 locus (sequence: TTAGTCCTAGTGCCATGAAG) was cloned into the pSpCas9-(BB)-2A-GFP vector under the control of the human U6 promoter to enable CRISPR-Cas9-mediated targeting^63^. Following co-transfection of the Cas9-sgRNA and landing pad plasmids, cells were cultured in puromycin-containing medium for initial selection of integrants. Surviving cells were then subjected to single-cell dilution into 96-well plates (**Extended Data** Fig. 6). Clonal populations were expanded and screened by digital PCR to identify lines harboring a single-copy insertion of the *attB* landing pad. The selected clones were used for downstream integration of the branching circuit.

For site-specific genomic integration of the branching circuit, we constructed a donor plasmid carrying the complete genetic device, a PhiC31 *attP* site (cccccaactgagagaactcaaaggttaccccagttgggg), and a bleomycin resistance gene (BleoR). Co-transfection of this plasmid with a PhiC31 integrase expression vector enabled recombination between the genomic *attB* site and the plasmid-borne *attP* site. Following transfection, cells were continuously cultured and passaged in zeocin-containing DMEM medium for 2-4 weeks to select for stable integrants and to allow dilution of residual episomal plasmids through cell division. Then, single-cell dilution and digital PCR screening were again performed to isolate clones harboring a single-copy insertion of the circuit. Notably, single-cell isolation and digital PCR were essential, as PhiC31-mediated integration in mammalian cells is prone to off-target events that can result in multiple gene copies being inserted into the genome. Such multicopy insertions would compromise the experimental design, which requires single-copy integration to ensure reliable and interpretable differentiation outcomes.

To achieve doxycycline-inducible expression of the Bxb1 recombinase, the Bxb1 or DHFR-Bxb1 gene was cloned into a lentiviral vector (LV-Tet-Bxb1) under a Tet-On promoter, along with a blasticidin (BSD) resistance gene and an iRFP-expressing cassette. Successfully transfected cells were enriched via antibiotic selection and flow cytometry sorting, and subsequently used for downstream differentiation experiments. All plasmids were validated by antibiotic selection, Sanger sequencing (Tsingke), and full plasmid sequencing (KBSeq, Sangon Biotech). Detailed component descriptions, protein sequences, and plasmid information are provided in **Tables S4-S8**.

### Plasmid construction and single-copy integration in CHO-K1 cells

As shown in **Extended Data** Fig. 6, the PhiC31 *attB*-containing landing pad was first integrated into the CHO-K1 genome using CRISPR-Cas9. Single-cell dilution and digital PCR screening were then performed to isolate clones harboring a single-copy insertion. We then co-transfected these cells with a plasmid containing the designed branching circuit (genetic device), a BleoR gene, a TagBFP expression cassette, and the PhiC31 *attP* site, along with a PhiC31 integrase expression vector. This enabled site-specific recombination between the genomic *attB* site and the plasmid-borne *attP* site, resulting in precise genomic integration of the branching circuit. Following transfection, cells were cultured and serially passaged in zeocin-containing F-12K medium for approximately one month to enrich stable integrants and allow for dilution of residual episomal plasmids through cell division. Subsequently, fluorescence-activated cell sorting (FACS) was performed to isolate TagBFP-positive cells (excitation at 405 nm), followed by single-cell dilution and digital PCR to identify clones harboring a single-copy insertion of the branching circuit. To enable doxycycline-inducible expression of Bxb1 recombinase, the Bxb1 or DHFR-Bxb1 gene was cloned into a lentiviral vector (LV-Tet-Bxb1) under the control of a Tet-On promoter, along with a BSD resistance gene and an iRFP expression cassette. Successfully transduced cells were selected using blasticidin and sorted via flow cytometry based on iRFP fluorescence (excitation at 640 nm), and then used for downstream differentiation experiments. All plasmids were validated through antibiotic selection, Sanger sequencing (Tsingke), and full-plasmid sequencing (KBSeq, Sangon Biotech). Detailed descriptions of genetic components, protein sequences, and plasmid maps are provided in **Tables S4-S8**.

### DNA transformation into E. coli

DNA (1-10 μL) was mixed with 200 μL of freshly thawed competent *E. coli* and incubated on ice for 30 minutes. A 45-second heat shock at 42 °C was performed, followed by a 2-minute cooldown. The mixture was recovered in 500 μL antibiotic-free LB medium at 37 °C for 1 hour before plating on LB agar containing the relevant antibiotic.

### Yeast transformation

Plasmid DNA was pre-digested with the *Not* I enzyme in a reaction containing 1 μL of CutSmart buffer (NEB), 0.5 μL of NotI-HF enzyme (NEB), and 8.5 μL of plasmid DNA. Following digestion at 37 °C for 1 hour, 5 μL of the digested plasmid was mixed with boiled salmon sperm DNA (ssDNA) to a final volume of 10 μL. *S. cerevisiae* was cultured in YPD medium at 30 °C for 1-2 days. After culturing, 1 mL of the yeast solution was centrifuged at 5,000 rpm for 30 seconds at room temperature, and the cell pellet was washed three times with 1 mL of 0.1 M lithium acetate (LiOAc)^64^. After the final wash, the cells were resuspended in 50 μL of 0.1M LiOAc, and 2 μL was transferred to PCR tubes to serve as competent cells for each transformation. For the transformation, 10 μL plasmid-ssDNA was added to 2 μL of competent yeast cells. The mixture also included 22.4 μL of 50% PEG-3350 and 3.6 μL of 1 M LiOAc. After mixing well by gentle vortexing, the reaction was incubated at 42 °C and 30 °C for 40 minutes each. After the incubation, the yeast cells in the PCR tubes were centrifuged at 8,000 rpm for 20 seconds at room temperature. The supernatant was removed, and the cells were resuspended in 0.1 mL of sterile water. Transformed cells were plated on SD agar plates for auxotrophic or antibiotic selection.

### Lentivirus packaging and transfection of engineered HEK293FT and CHO-K1 cells

Lentiviral particles (LV-Tet-Bxb1) were generated by transient transfection of HEK293FT cells using a three-plasmid system^65^. The transfer plasmid encoding Bxb1 or DHFR-Bxb1 was co-transfected with the packaging plasmids psPAX2 (which encodes gag, pol, and rev) and pMD2.G (which encodes the VSV-G envelope protein) using Lipofectamine™ 3000 (Invitrogen, L3000015) according to the manufacturer’s instructions. After 48 and 72 hours, the lentiviral supernatant was collected, filtered through a 0.45 μm PES filter (Millipore, SLHP033RB), and concentrated by ultracentrifugation.

For lentiviral transduction, about 1×10^6^ HEK293FT or 8×10^5^ CHO-K1 cells were seeded into each well of a 6-well plate and cultured for 24 hours. The culture medium was then removed and replaced with 1 mL of the collected lentiviral suspension. After an 18-hour incubation, 2 mL of fresh medium (without selection antibiotics) was added, and the cells were incubated for an additional 24 hours. Subsequently, all contents from the 6-well plate were transferred into a 10 cm culture dish. Cells were dissociated using 0.25% trypsin and pooled into the dish. An additional 7 mL of complete medium containing the appropriate selection antibiotics was added. Cells were cultured under standard conditions (37°C, 5% CO_2_), with fresh medium (containing antibiotics) replaced every 4 days, until they reached confluence in the 10 cm dish.

### Digital PCR-based absolute quantification of gene copy number

To determine the copy number of integrated genetic constructs in engineered *E. coli*, *S. cerevisiae* BY4741, and mammalian cells, we used a ratio-based digital PCR (dPCR) approach. Genomic DNA was extracted from the host cells and submitted to Tsingke Biotechnology Co., Ltd. (Beijing, China) for digital PCR analysis. The copy number of the target construct was calculated by comparing its absolute concentration to that of endogenous reference genes with known copy numbers in each chassis.

Digital PCR (dPCR) is an absolute quantification method for nucleic acids that offers high sensitivity and accuracy^66^. The technique partitions a conventional PCR reaction into tens of thousands of nanoliter-sized droplets, each of which may contain zero, one, or a few copies of the target DNA molecule. Each droplet serves as an independent PCR microreactor. After amplification, fluorescence detection is performed for each droplet to determine the presence or absence of the target sequence. The proportion of fluorescent (positive) versus non-fluorescent (negative) droplets is then used, based on Poisson distribution, to calculate the absolute concentration of the nucleic acid targets (in copies per microliter). In this work, fluorescence detection relies on TaqMan probes. FAM-BHQ1 was used for target sequence detection, and VIC-MGB was used for the internal reference. FAM and VIC serve as the fluorophores, while BHQ1 and MGB act as the respective 3’quencher moieties. For sample preparation, DNA was extracted from bacteria, yeast, or mammalian cells and used directly as a template. dPCR reactions were performed using the 2× T5 Fast qPCR Mix (probe-based, Tsingke), and all analyses followed the manufacturer’s protocol.

The copy number (N) was calculated using the formula:

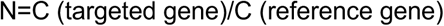

Here, C represents the absolute concentration of DNA molecules in solution, expressed in copies per microliter (copies/μL). This value was calculated using the Poisson-based digital PCR equation:

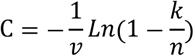

where n is the total number of partitions (i.e., Nb Droplets), k is the number of positive partitions (i.e., Nb pos), and v is the volume of each partition. For our system, which was based on the digital PCR instrument (SinoGene DQ24 Digital PCR Instrument) from Tsingke Biotechnology Co., Ltd (Beijing, China), v=0.6 ×10^-3^ μL. The obtained dPCR results are shown in **Fig. S3**.

### Flow cytometry

For most flow cytometry analyses of yeast, the following protocol was used: On Day 1, yeast strains were inoculated into 1 mL of YPD medium supplemented with inducers (1 mM aTc or 1 μM β-estradiol) and cultured overnight in 2.2 mL deep-well plates at 30 °C with shaking at 800 rpm. On Day 2, cells were diluted 1:50 into 200 μL of ddH₂O in a 96-well flat-bottom plate and analyzed using an Attune NxT Flow Cytometer (Thermo Scientific). A total of 10,000 events were collected per sample, and data were analyzed with FlowJo software. For doublet ratio assessments, yeast cultures were diluted 1:5 in ddH₂O prior to measurement.

For mammalian cell sorting experiments, cells grown to confluency in 10 cm petri dishes were first digested using 0.25% trypsin, centrifuged, and resuspended in 2 mL of F-12K medium supplemented with 4% penicillin-streptomycin. The cell suspension was then filtered through a 75 μm cell strainer and loaded into sample tubes for sorting. Target populations were sorted using a FACSAria Fusion flow cytometer (BD Biosciences), and approximately 2 × 10^5^ sorted cells were transferred into 6-well plates for further culture.

### Growth curve measurements

Single yeast colonies were grown overnight in 1 mL of YPD medium at 30 °C with shaking at 800 rpm in a 2.2 mL 96 deep-well plate. The following day, cultures were diluted to an OD_600_ of approximately 0.06 in 200 μL of fresh YPD medium in a clear, flat-bottom 96-well microplate. OD_600_ readings were recorded every 5 minutes for 30 hours at 30 °C using a microplate reader (Biotek, SYNERGY H1), which included a 10-second shake before each measurement. The maximum growth rate during the exponential phase was calculated using the formula: LN(OD_600_(*t* + 6)/OD_600_(*t*))/6, where ’*t*’ is the time in hours.

### Co-culture passaging

To evaluate the long-term stability of differentiated progeny cell consortia, yeast cells were cultured in predetermined proportions in 5 mL of YPD medium at 30 °C for one day. After this initial growth period, the proportions of each type of yeast within the consortia were analyzed using flow cytometry. For each subsequent round of growth, the yeast culture was diluted 1:1000 into 5 mL of fresh YPD medium and allowed to grow again for one day. This cycle of growth, dilution, and flow cytometry analysis was repeated for a total of 7 rounds to monitor changes in the consortia composition over time.

To assess the long-term stability of the branching circuits in the absence of inducers, yeast cells were serially passaged at a 1:1000 dilution for 15 consecutive rounds (∼15 days), and monitored for spontaneous differentiation events caused by recombinase leakiness via flow cytometry (Attune NxT Flow Cytometer, Thermo Scientific). Similarly, in CHO-K1 cells, stability was assessed over five passages (∼15 days) at a 1:6 dilution, by monitoring for unintended differentiation events that could result from leaky recombinase expression.

### NGS and analysis

Next-generation sequencing (NGS) via amplicon sequencing was performed on yeast cells encoding differentiation circuits. Yeast cultures were grown in 5 mL YPD and induced with either 1 mM aTc or 1 μM β-estradiol. Genomic DNA was extracted using a TIANamp Yeast DNA Kit (TIANGEN). A 300 bp DNA fragment undergoing recombinase-catalyzed rearrangement was amplified via PCR using genomic DNA as a template. The primers used for PCR were: forward primer (5’-cgacggcgatcacagacattaa-3’) and reverse primer (5’-ctcacactgacgaatcatgt-3’). The amplified PCR products were purified and sent to Tsingke for NGS and data analysis.

### Confocal fluorescence microscopy

For liquid-cultured yeast characterization, 5 μL of the yeast suspension was placed onto the center of a microscope slide. A clean 1 cm × 1 cm coverslip was gently placed over the droplet to ensure the even spreading of the solution. Once prepared, the sample was observed under a confocal fluorescence microscope (AX, Nikon). To characterize colonies grown on solid media, an SD medium containing 0.7% agar and appropriate inducers or antibiotics was prepared, allowed to cool, and solidified. A 2 μL aliquot of freshly cultured yeast solution (diluted 1:200 to an OD_600_ of approximately 0.06) was pipetted onto the agar surface. Plates were incubated at 30 °C for 3-5 days. After incubation, the colonies were imaged using the confocal fluorescence microscope.

For time-dependent imaging of cellular differentiation (**Mov. S1-S3**), uninduced yeast cells were diluted 1:1000 to an initial OD_600_ of ∼0.01 and seeded into glass-bottom confocal imaging dishes containing YPD medium supplemented with chemical inducers. Cultures were incubated at 30 °C on a temperature-controlled microscope stage, and fluorescence microscopy (Nikon AX) was performed at 10-minute intervals to monitor the emergence of multicellular aggregates.

For imaging of differentiation-driven pattern formation in yeast (**Mov. S4-S5**), engineered yeast progenitors were diluted to approximately 100 cells and seeded into 96-well round-bottom ultra-low attachment plates (Corning, #7007) containing YPD medium supplemented with aTc. Plates were centrifuged at 100 × g for 10 minutes to facilitate cell settling, then incubated at 30 °C on a temperature-controlled microscope stage. Time-lapse fluorescence microscopy (Nikon AX) was performed at 10-minute intervals to monitor the development of multicellular aggregates.

For time-lapse imaging of CHO-K1 cells, 100 or 1000 cells were seeded into 96-well round-bottom ultra-low attachment plates (Corning, #7007) containing F-12K medium supplemented with antibiotics, and, when necessary, Dox and TMP. Plates were incubated in a standard cell culture incubator at 37 °C with 5% CO₂. Confocal fluorescence imaging was performed every 24 hours to monitor cell differentiation and aggregate formation over time.

### Calculation of the area of yeast aggregates

Ten images of each engineered yeast aggregate depicted in **Fig. S78** were captured using the confocal fluorescence microscope (AX, Nikon). The area of each yeast aggregate in the fluorescence images was quantified using ImageJ software. Areas smaller than 400 μm^2^ were excluded from the analysis. The remaining data were compiled to create a graphical representation of yeast aggregate size distribution.

### Sedimentation rate measurement

Yeast cells containing genetic circuits were cultured in 1.5 mL of induced YPD medium in 96 deep-well microplates at 30 °C and 800 rpm overnight. The yeast suspension was then adjusted to an OD_600_ of 1.7. One milliliter of yeast solution was transferred to a cuvette, and OD_600_ measurements were taken every 30 minutes for 210 minutes using a UV spectrophotometer. The collected data were analyzed to determine sedimentation rates.

### Characterization of yeast protein secretion

The Nano-Glo® HiBiT Extracellular Detection System (Promega) was utilized to confirm the secretion of enzymes by engineered yeast. This system relies on the interaction between the HiBiT tag, an 11-amino-acid epitope (sequence: VSGWRLFKKIS), and LgBiT, which generates bioluminescence upon binding^67^. For the assay, HiBiT was genetically fused to target proteins or enzymes. Supernatants (100 µL) from cultured yeast were combined with 1 µL of LgBiT protein, 97 µL of reaction buffer, and 2 µL of bioluminescent substrate, yielding a total reaction volume of 200 µL. The mixture was incubated for 5 minutes at room temperature, and the bioluminescence was measured using a microplate reader (BioTek). The BY4741 strain, which does not secrete engineered proteins, served as a negative control.

### Enzyme activity of secreted α-Galactosidase (Mel1)

A stock solution of X-α-Gal (40 mg/mL, Sigma-Aldrich, 16555) was prepared in dimethyl sulfoxide (DMSO). For activity assessment, 100 µL of this solution was added to 3 mL of YPD medium inoculated with yeast cells at a 1:1000 dilution. After 24 hours of incubation at 30 °C, cultures were collected and imaged.

### Enzyme activity of secreted Laccase

As copper-dependent enzymes, laccases require copper supplementation^68^. Stock solutions of 2,2’-azino-bis (3-ethylbenzothiazoline-6-sulphonic acid, ABTS) (Sigma-Aldrich, A1888) and copper sulfate were prepared at final concentrations of 0.1 M and 1 M, respectively. For activity evaluation of the secreted laccase from *Coriolopsis trogii*, 125 µL of ABTS solution and 25 µL of CuSO₄ solution were added to 3 mL of YPD medium inoculated with yeast cells (1:1000 inoculation). Cultures were incubated at 30 °C for 24 hours and then imaged.

### Enzyme activity of secreted Lipase

Yeast strains were cultured in YPD medium supplemented with 1% soybean oil for 48 hours. Fermentation broth (5 mL) was harvested by centrifugation (12,000 rpm, 15 minutes). Lipids were extracted from the supernatant by mixing with 500 μL acetic acid, 500 μL of 12% (w/v) sodium chloride solution, and 2 mL ethyl acetate. The mixture was vortexed for 15 minutes, centrifuged (12,000 rpm, 15 minutes), and the organic phase was collected, dried under nitrogen, and resuspended in chloroform. Lipid separation was performed via thin layer chromatography (TLC) using a solvent system (hexane: diethyl ester: acetic acid = 70:30:1). Lipids were visualized using iodine vapor.

### Extraction and characterization of Violacein and β-carotene

To extract Violacein and β-carotene from cultured yeast, yeast cells (1 mL, OD_600_ = 2.0) were harvested by centrifugation (4,000 rpm, 1 minute). After discarding the supernatant, pellets were resuspended in 500 μL of DMSO and incubated at 37 °C for 30-60 minutes with grinding to accelerate cell lysis and improve extraction efficiency. Following incubation, the mixture was centrifuged (4,000 rpm, 1 minute), and the supernatant containing extracted Violacein and β-carotene, was analyzed via full-spectrum scanning (400-650 nm, 5 nm steps) using a BioTek microplate reader.

### Counting ratios of pigment-producing yeast generated by branching devices

Yeast strains with *pZ*-driven violacein and *pPhIF*-regulated β-carotene synthesis pathways were transformed with a designed branching circuit using Z_3_EV and PhIF-VP16 as reporters. Colonies were grown on agar plates for three days, inoculated into 5 mL of YPD medium supplemented with 1 mM aTc and 1 nM β-estradiol, and cultured at 30 °C with shaking (1,000 rpm) for 8 hours. A low concentration of β-estradiol (1 nM) achieved effective induction, remarkably exceeding the commonly used 1 μM concentration. Cultures were diluted 10,000×, spread onto YPD agar plates (supplemented with 1 nM β-estradiol), and incubated for 3 days. Plates were imaged, and colonies producing orange (β-carotene) and purple (violacein) pigments were counted.

### Construction of a recombinase-guided pigment-producing yeast library

A small library of 25 synthetic differentiation circuits was constructed using the Golden Gate assembly method^25^. Each construct contained the unique engineered *attP* variants bearing specific point mutations that modulate its recombination efficiency with the Bxb1 recombinase, thereby enabling a tunable range of catalytic activity, from “fast” to “slow” (**Fig. S75**). This diversity in recombination rates allows differential activation of downstream genetic modules (e.g., Z_3_EV or PhlF-VP16), resulting in variable proportions of violacein- and β-carotene-producing subpopulations. The assembled library containing 25 different plasmids was transformed into *S. cerevisiae* BY4741, which had been engineered to harbor both violacein and β-carotene biosynthetic pathways, each regulated by a different synthetic transcription factor (Z_3_EV or PhlF-VP16, respectively). A total of 96 individual colonies were randomly selected and cultured in 2.2 mL deep-well plates at 30 °C with shaking at 800 rpm, in the presence of inducers (1 mM aTc and 1 nM β-estradiol) to initiate differentiation.

### DNS method for measuring cellulose degradation by cellulase

Glucose concentration was determined using a modified DNS method^69^. Samples (50 µL) were mixed with 100 µL of DNS buffer (Macklin) in 1.5 mL tubes, heated at 100 °C for 5 minutes, and cooled on ice for 1 minute. After cooling, 150 µL of ddH₂O was added, and 200 µL of the solution was transferred to a microplate. Absorbance at 540 nm was measured, and glucose concentrations were determined using a standard curve (0.125-5 g/L) (**Fig. S76**). Cellulase activity was assessed using 1 mL supernatant of yeast culture (OD_600_=2.0) with 2% Avicel PH-101 (Sigma) in sealed deep-well microtiter plates. Samples were incubated at 35°C with shaking (∼300 rpm) for 24 hours, and 50 µL aliquots were analyzed for glucose content using the DNS method.

### Construction of the snowflake yeast

Snowflake yeast strains were constructed by deleting ACE2 using the CRISPR-Cas9 system, as previously reported^44^. The commercially available plasmid pMYT095, which contains Cas9 and gRNA cassettes, was employed for this purpose^62^. The guide RNA sequence (TTATTCAAAATATAATTGTCGGG) was selected using the Yeastriction online tool (http://yeastriction.tnw.tudelft.nl/). Plasmids designed for ACE2 deletion were constructed and validated by Sanger sequencing (Sangon Biotech). Donor DNA fragments were synthesized and purified using a DNA Extraction Mini Kit (Vazyme Biotech Co., Ltd) prior to transformation. Both the plasmids and donor DNA were introduced into yeast cells via the lithium acetate transformation method. Transformant colonies were subsequently confirmed by PCR and Sanger sequencing (Sangon Biotech).

### Image-processing workflow for quantifying cell-cell interaction frequencies

To quantify cell-cell interactions in engineered yeast populations, we developed a five-step image processing pipeline based on fluorescent microscopy data (**Fig. S81**). First, raw images underwent manual enhancement using Adobe Photoshop to improve contrast and delineate boundaries between touching or dividing cells, thereby facilitating accurate downstream analysis. Second, enhanced images were segmented using CellProfiler (v4.2.1). Fluorescent channels (R, G, B) were split and processed using the IdentifyPrimaryObjects module with typical object diameters set to 10-160 pixels. Adaptive thresholding was performed using the Otsu method in three-class mode to ensure robust cell identification across varying fluorescence intensities. Third, segmented images and corresponding object data were imported into MATLAB for further analysis. In the fourth step, individual cells were classified into distinct types based on their dominant RGB fluorescence intensities, which served as proxies for specific cell types or engineered markers. Lastly, cell-cell interactions were identified through spatial proximity analysis: pairwise distances between all classified cells were computed, and interactions were defined for cell pairs located within a 35-pixel threshold. Interaction networks were visualized by drawing connection lines between spatially adjacent cells, and interaction frequency heatmaps were generated to represent normalized contact frequencies between all cell type combinations. These matrices provided a quantitative view of interaction preferences, enabling identification of preferential self-or heterotypic associations within the population.

### In vitro formation of cellular morphologies in engineered mammalian cells

To investigate and engineer cellular morphologies in CHO-K1 cells, two primary in vitro culture strategies were employed. In the first approach, CHO-K1 cells that had undergone lineage differentiation via doxycycline and TMP induction were collected and diluted to a final density of approximately 100 or 1000 cells per well. The cells were seeded into 96-well ultra-low attachment plates in 100 μL of F-12K medium supplemented with 10% FBS, 2% PS, 4 μg/mL puromycin, 20 μg/mL BSD, and 400 μg/mL zeocin. Cells were incubated at 37 °C with 5% CO₂. Cellular morphologies were monitored over time using confocal fluorescence microscopy at 2, 12, 24, 48, 72, and 96 hours post-seeding.

In the second approach, approximately 100 CHO-K1 cells genetically engineered to contain a single-copy differentiation circuit and an inducible Bxb1 recombinase system were seeded into 96-well ultra-low attachment plates under the same medium conditions. To induce in situ differentiation, 1 μg/mL doxycycline and 0.1 mM TMP were added at the time of seeding. Morphological changes and cell structure formation were similarly monitored using confocal fluorescence microscopy at 2, 12, 24, 48, 72, and 96 hours post-induction. These time-course analyses enabled the observation of dynamic morphological changes driven by genetic differentiation programs and chemical inducers in a controlled 3D culture environment.

### Development of a mathematical model for predictive circuit design

To design cell differentiation patterns predictively, a simplified model was constructed to describe the Bxb1-mediated excision of the engineered circuit. The excision process was considered both with and without steric hindrance imposed by the transcription machinery. We hypothesized that compared to the weak promoter, a strong promoter on the differentiation circuit would attract more transcription factors, potentially reducing the cleavage efficiency of Bxb1 recombinase. In the absence of steric hindrance, the left and right arm could be cleaved at the rates *k*_*A*_ and *k*_*B*_, respectively. The values of *k*_*A*_ and *k*_*B*_ were determined by the lengths of the left and right arms. A power function was used to describe the relationship between the arm length and the excision rate: 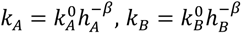. Here, *h*_*A*_ And *h*_*B*_ represent the arm lengths, and 𝛽 is a positive constant.

The promoters on both arms recruit transcription machinery, creating a physical obstruction that reduces the recombination efficiency. The recruitment of transcription machinery on the left arm reduces the cleavage rate of the left arm, changing it from _𝑘!_ to _𝑣!_. Similarly, recruitment on the right arm changes *k*_*B*_ to *v*_*B*_. We assumed that the steric hindrance of one arm does not affect the recombination of the other arm. We used a variable 𝛼 to describe the strength of the steric hindrance: 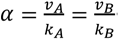.

Therefore, the cell differentiation dynamics mediated by the engineered circuit can be described with a group of ordinary differential equations (ODEs):

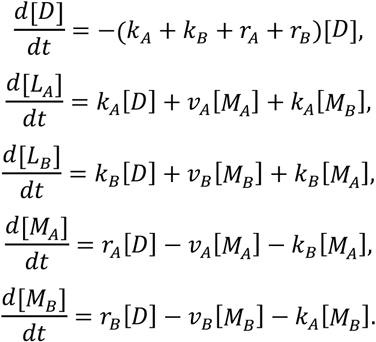

Here, _[_𝐷_]_ presents the population-level concentration of the ‘naked’ genetic device that has been neither cleaved nor bound with transcription machinery _[𝐿_*_A_*_]_ and _[𝐿_*_B_*_]_ are the concentrations of the recombination products. Excision of the left arm produces _𝐿_*_A_*, while cutting off the right arm produces _𝐿_*_B_*. _𝑀_*_A_* and _𝑀_*_B_* describe the concentrations of the circuit bound with transcription machineries. *r*_A_ And *r*_B_ are the binding rates of the transcription machineries.

If all the parameters were given, this model would allow us to predict the ratios of left or right arm excision. However, we didn’t have the luxury since many of the parameters were challenging to directly measure. Therefore, we sought to estimate these parameters by fitting our model with a finite number of experimental measurements. We hypothesized that the optimal parameters should be those minimizing the discrepancy between the predicted and measured ratios. To this end, we defined an objective function 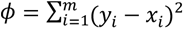 where _𝑥_*_i_* is experimentally determined ratio, _𝑦_*_i_* the predicted value and 𝑖 the index of the data. The parameters can then be estimated by searching for the minimum of the objective function in the defined parameter space.

In experiments, we varied the lengths of left and right arms and measured the ratio of Venus and mScarlet cells resulting from different arm length combinations (**Fig. 2D**). We used these data to fit our model (**Fig. S39-S40**). In particular, we screened the parameters in the following ranges: 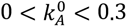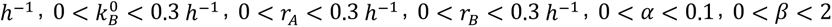. For each chosen parameter combination, we calculated the corresponding 𝜙 value. We then obtained the optimal parameter combination that best fit the experimental measures: 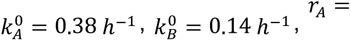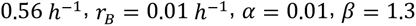.

These parameters can be further applied to determine the 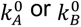 of the *attP* variant. For instance, for an *attP* variant on the left arm, the new 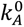 can be determined by fitting our model with the new measurements while keeping all other parameters unchanged. The new 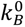 for the *attP* variant on the right arm can be estimated similarly (**Table. S1**). The same approach can also be used to determine the _𝑟_*_A_* and _𝑟_*_B_* values corresponding to different promoters (**Table. S2**). After the parameter associated with each condition (arm’s length, *attP* variant, and promoter type) was estimated, the further enables the prediction of green-to-red ratios under different condition combinations (**Table. S3**).

## Statistics and Reproducibility

All data was processed using Excel (Microsoft) and Origin (OriginLab) software. Error bars in the figures represent standard deviation, as specified in the figure legends. The number of replicates for each data set is also detailed in the figure legends, and all replicates have been included in the manuscript for comprehensive analysis.

## Acknowledgments

This work was partially sponsored by the National Key R&D Program of China (2025YFA0921400 to C.Z.), the National Science Fund for Distinguished Young Scholars (32125023 to C.Z.), the National Natural Science Foundation of China (32201105 to B.A., 32201202 to Y.W., and 32401222 to Q.Z.), the Shenzhen Medical Research Fund (A2303072 to Q.Z.), and the Shenzhen Science and Technology Program (ZDSYS20220606100606013 to C.Z. and JCYJ20240813154818024 to Q.Z.). This work was also supported by the US Defense Advanced Research Projects Agency (DARPA) Engineered Living Materials award (W911NF-17-2-0077 to T.K.L.) and the US Department of Energy (DOE) (DE-FG02-02ER63445 to G.M.C.).

## Author contributions

T.-C.T., C.Z., G.M.C., and B.A. conceptualized and directed the research. T.-C.T. built the branching device prototype in bacteria and yeast. B.A. designed and carried out most of the gene construction and flow cytometry experiments, with assistance from Q.Z., Y.L., K.G., Y.W., Q.L., and Y.W. Q.Z. contributed to the construction of snowflake yeast and pigment-producing strains. T.W. developed the mathematical model for progeny cell prediction. K.L., Y.P., M.Y., and C.L. assisted with culturing HEK293FT cells and characterizing the branching devices in mammalian cells. D.Z. assisted with image processing for quantifying cell-cell interaction frequencies. W.S. contributed to optimizing the yeast inducible systems. T.-C.T., T.K.L., G.M.C., B.A., and C.Z. contributed to funding acquisition. B.A., T.-C.T., C.Z., and G.M.C. wrote the manuscript, with input from all authors.

---

**Extended Data Fig. 1.**
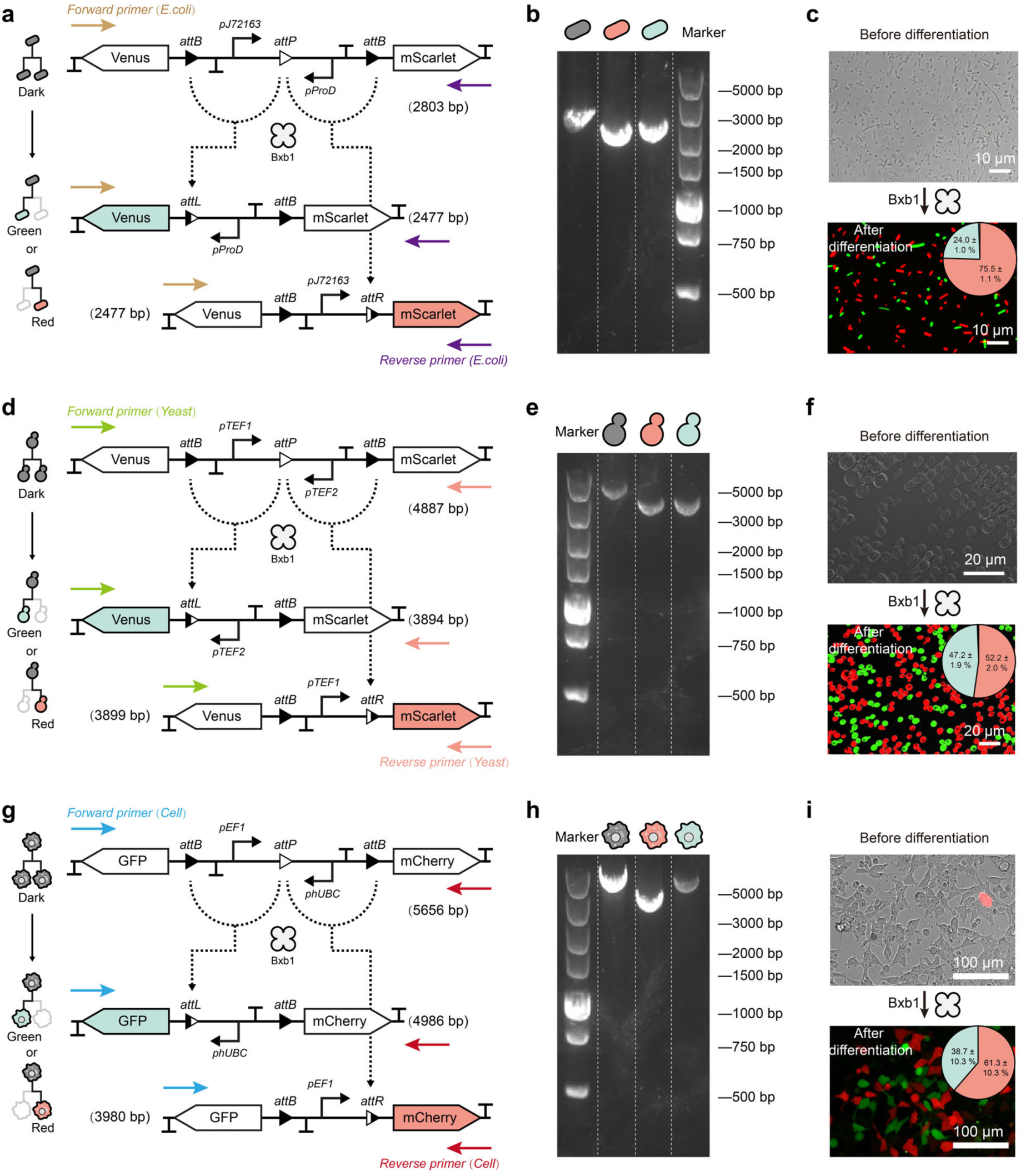
Genotypic validation of Bxb1-mediated recombination in bacteria, yeast, and mammalian cells. **a-c,** In *E. coli*, a recombinase-recognized genetic circuit containing *attB* and *attP* sites flanking fluorescent reporters (Venus and mScarlet) was used to assess Bxb1-mediated differentiation. **a**, Schematic of expected recombination products and PCR primer locations. The undifferentiated state yields a 2803 bp product, while recombined states yield 2477 bp bands corresponding to the two possible differentiation outcomes. **b**, PCR analysis of individual bacterial clones exhibiting dark, green, or red fluorescence shows band sizes consistent with their recombination status. **c**, Microscopy before and after induction shows robust green and red fluorescent proteins’ expression, with quantification confirming recombinase-mediated diversification (data are mean ± s.d. of *n* = 8 samples). **d-f**, Equivalent strategy applied in *S. cerevisiae* (BY4741) using yeast-specific promoters. **d**, Circuit diagram with predicted PCR products: 4887 bp for the undifferentiated state and 3894/3899 bp for the recombined states. **e**, PCR analysis of sorted yeast colonies reveals genotype-specific bands matching fluorescence phenotypes. **f**, Microscopy before and after Bxb1 recombinase induction confirms expected differentiation profiles, with proportions summarized in pie chart (data are mean ± s.d. of *n* = 8 samples). **g-i**, Validation of recombinase function in mammalian cells. **g**, Schematic of the circuit and locations of PCR primer pairs. The undifferentiated state yields a 5565 bp band; recombined products yield 3980 bp and 4986 bp, depending on the recombination outcome. **h**, PCR from HEK293FT cells sorted by fluorescence confirms genotypic identities consistent with observed phenotypes. **i**, Fluorescence microscopy before and after induction shows effective circuit activation and recombinase-dependent diversification into mCherry+ and GFP+ populations, with quantification of cell-type frequencies (data are mean ± s.d. of *n* = 8 samples).

**Extended Data Fig. 2.**
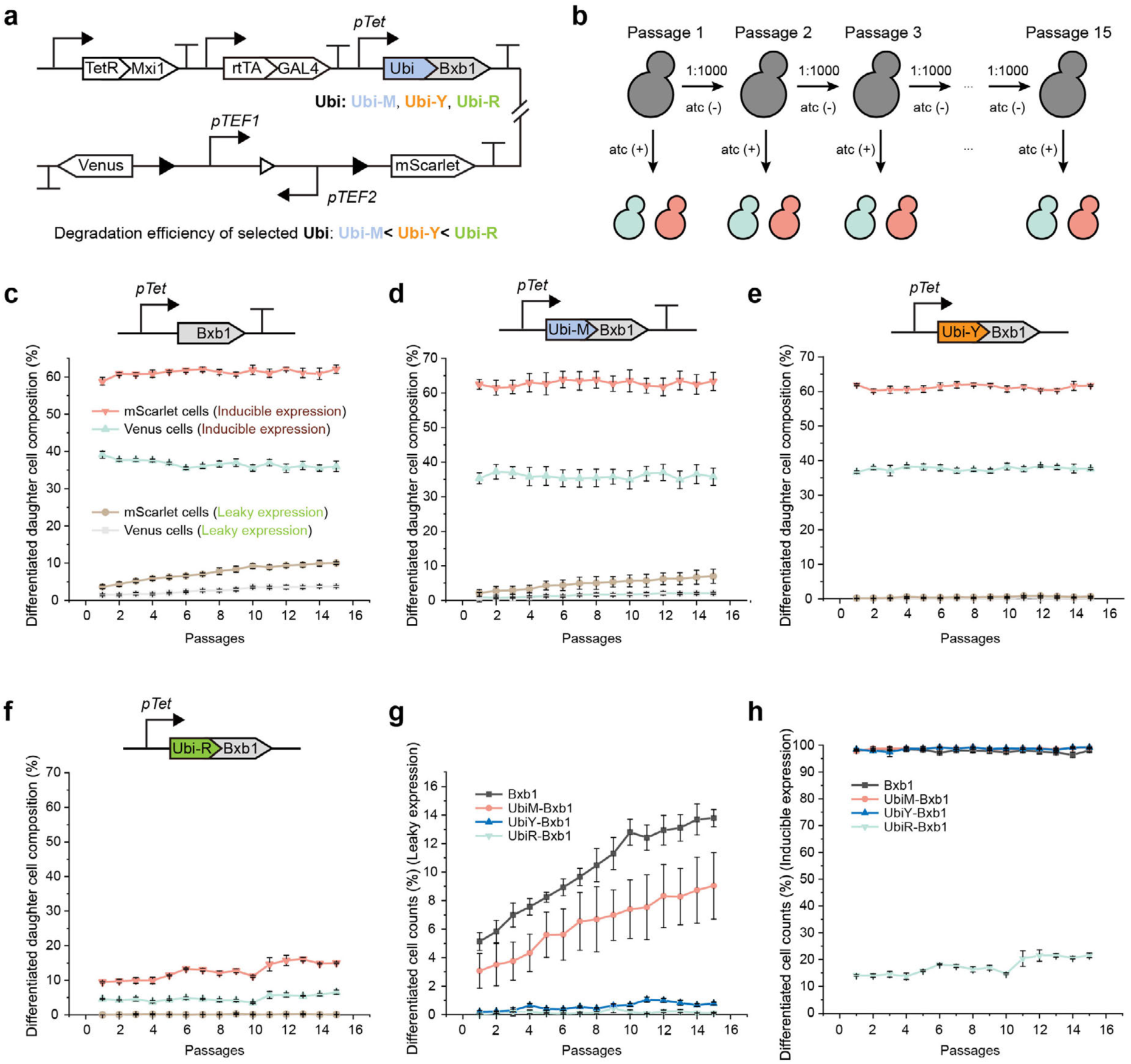
Long-term assessment of recombinase leakiness and stability in yeast. **a**, Schematic of the genetic circuit used in this study. Bxb1 recombinase was placed under control of an aTc-inducible promoter (*pTet*) and fused to one of three degradation tags (Ubi-M, Ubi-Y, or Ubi-R), each conferring different levels of proteolytic degradation. Venus and mScarlet served as differentiation markers. **b**, Overview of the experimental setup. Yeast strains carrying engineered genetic circuits were serially diluted and passaged 15 times (1:1000 dilution daily at 30 °C) with or without the aTc inducer. **c,** Fraction of Venus- and mScarlet-positive cells over 15 days of serial passage, using Bxb1 under the control of the aTc-inducible promoter to trigger circuit recombination. Brown and gray lines indicate leaky expression of mScarlet and Venus, respectively, in the absence of aTc induction. Red and green lines represent the proportion of mScarlet- and Venus-positive cells after aTc induction. **d**, Fraction of Venus- and mScarlet-positive cells over 15 days of serial passage, using UbiM-Bxb1 variant under the control of the aTc-inducible promoter to trigger circuit recombination. **e**, Fraction of Venus- and mScarlet-positive cells over 15 days of serial passage, using UbiY-Bxb1 variant under the control of the aTc-inducible promoter to trigger circuit recombination. **f**, Fraction of Venus- and mScarlet-positive cells over 15 days of serial passage, using UbiR-Bxb1 variant under the control of the aTc-inducible promoter to trigger circuit recombination. **g**, Summary of leakiness under non-induced conditions. Wild-type Bxb1 showed increasing spontaneous recombination (∼14% by passage 15), while UbiY- and UbiR-tagged variants maintained leakiness below 1%. **h**, Induced differentiation after 15 passages. UbiR-Bxb1 exhibited reduced induction efficiency (∼20%), demonstrating that while strong degradation can effectively suppress leakiness, it may also compromise overall recombinase activity. All data represent mean ± SD from 6 biological replicates.

**Extended Data Fig. 3.**
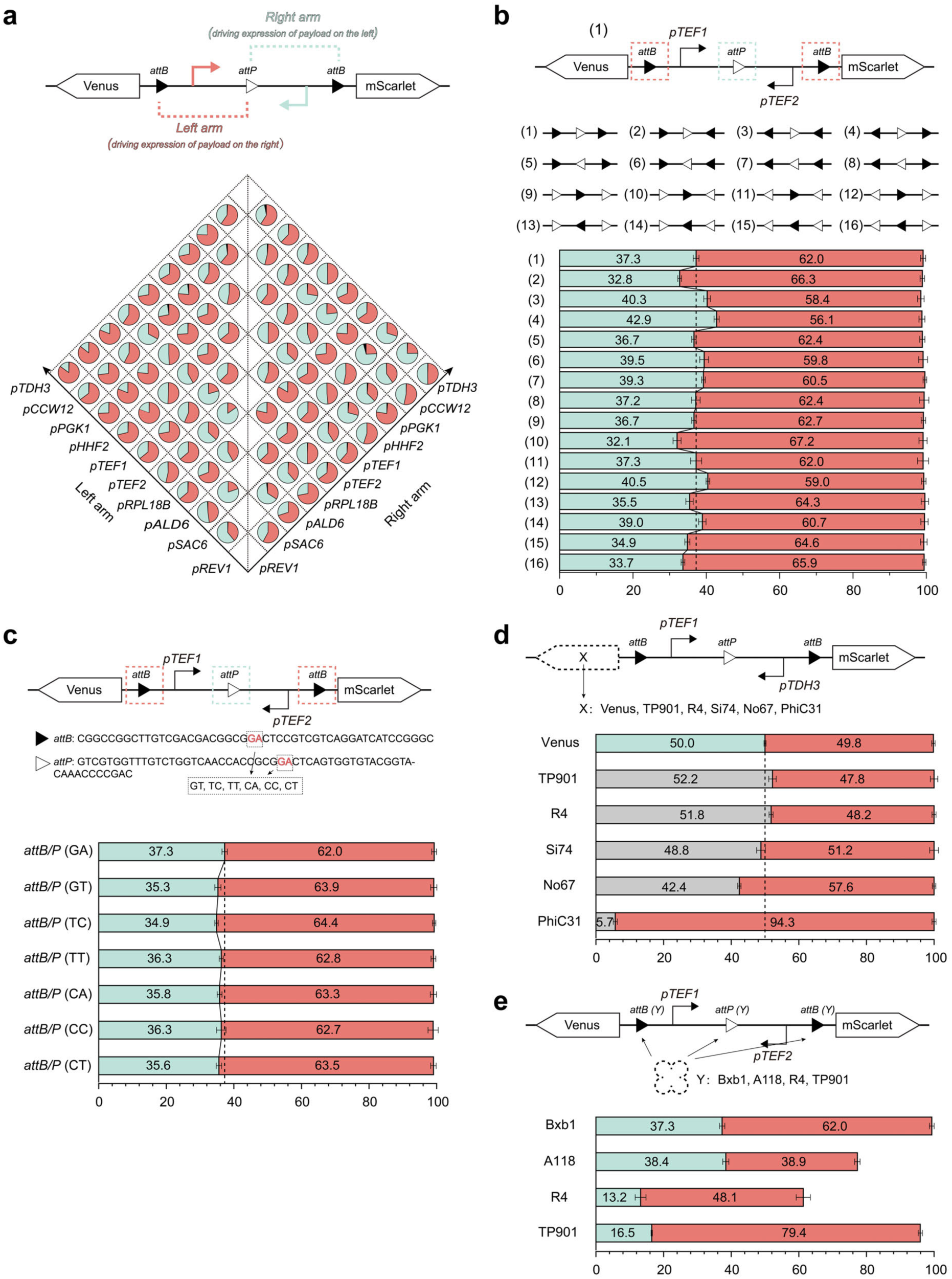
Factors determining the performance of recombinase-based branching devices. **a**, Schematic of a branching genetic circuit in which ten constitutive yeast promoters were systematically positioned in the left and right arms flanking a central recombination module. This combinatorial design generated 90 unique circuit architectures with varied Venus-to-mScarlet progeny ratios. Data are mean ± s.d. of *n* = 8 samples. **b**, 16 circuit configurations obtained by permuting the orientation of three recombination sites, with identical flanking promoters (*pTEF1* on the left, *pTEF2* on the right). Data are mean ± s.d. of *n* = 8 samples. **c**, Design and performance of Bxb1 *att* sites carrying orthogonal central dinucleotide variants (GA, GT, TC, TT, CA, CC, CT) in otherwise identical recombination cassettes. Data are mean ± s.d. of *n* = 8 samples. **d**, Effect of alternative payloads on expression balance (data are mean ± s.d. of *n* = 8 samples). The Venus gene was replaced with one of several recombinases (e.g., TP901, R4, Si74, No67, or PhiC31), positioned upstream of the *pTDH3* promoter driving mScarlet. **e**, Comparative analysis of branching circuits catalyzed by four different serine integrases (Bxb1, A118, R4, TP901) using identical flanking promoters. Data are mean ± s.d. of *n* = 8 samples.

**Extended Data Fig. 4.**
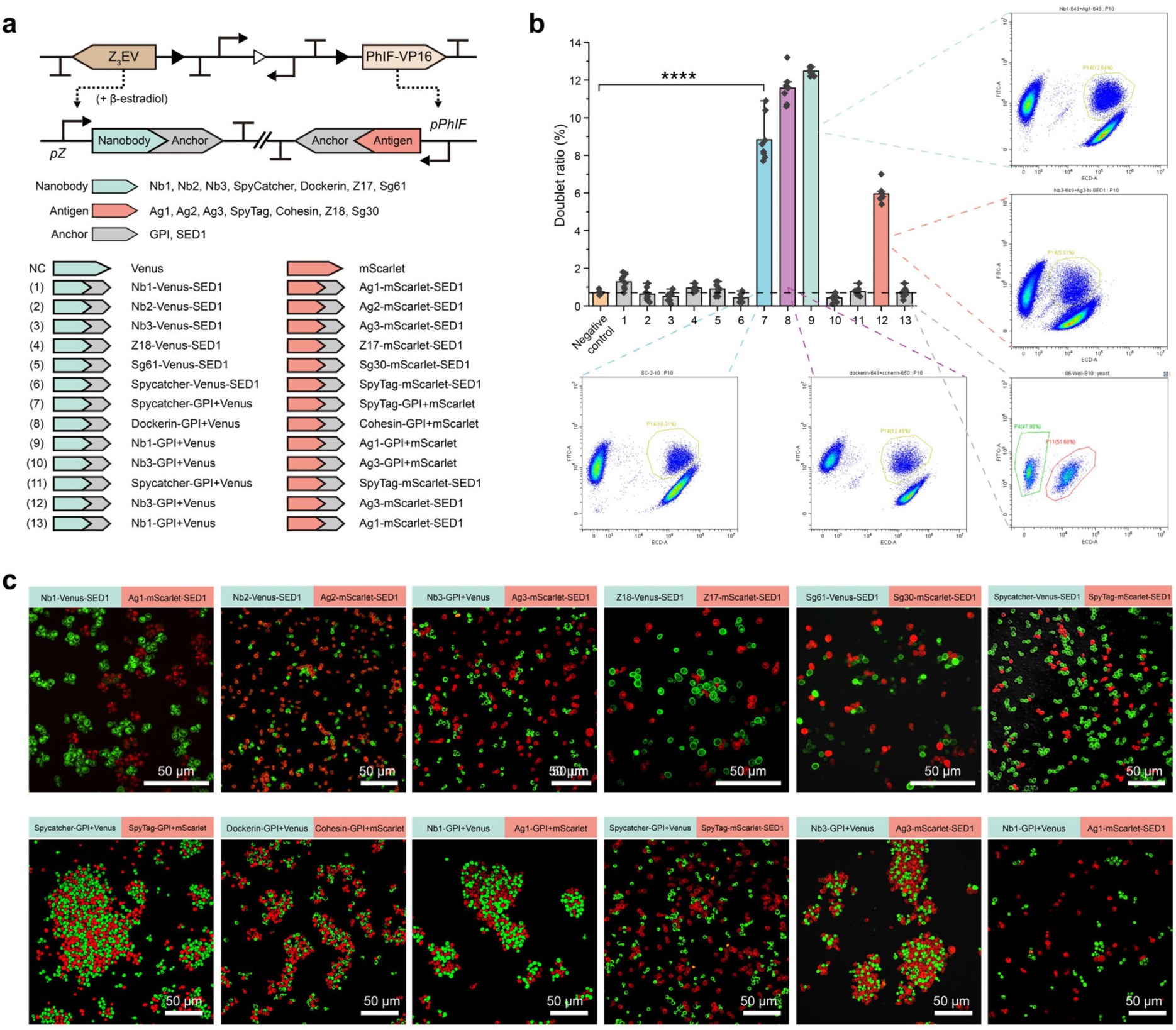
Screening protein pairs for yeast surface display and cellular morphogenesis. **a**, Schematic of the branching circuit and 13 protein pair candidates. “NC” denotes the negative control, consisting of fluorescent BY4741 yeast cells lacking CAM expression. **b**, Flow cytometry analysis identified four designs with increased doublet ratios, indicating successful protein display and strong binding interactions between the two progeny cell types. Data are mean ± s.d. of *n* = 8 samples. **c**, Representative fluorescence images of cellular aggregates formed by yeast displaying different protein pairs. Protein sequences are listed in the **Table. S5**.

**Extended Data Fig. 5.**
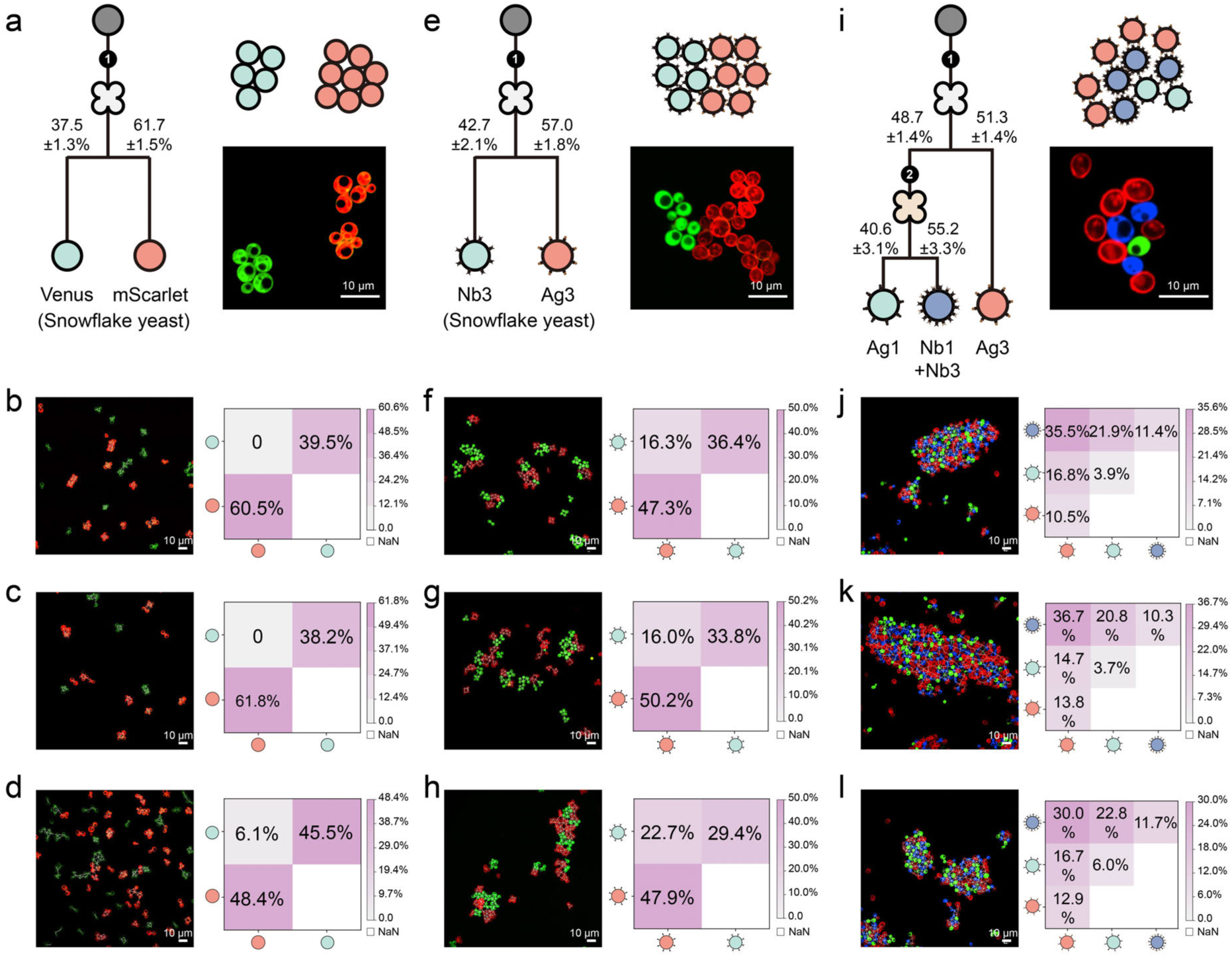
Representative quantification of cell-cell interaction patterns in differentiated yeast. **a**, Schematic of a recombinase-based differentiation pattern in engineered snowflake yeast. Upon recombination, initially non-fluorescent cells differentiate into red (mScarlet) and green (Venus) fluorescent subpopulations at defined ratios. **b-d**, Representative fluorescence microscopy images of differentiated yeast aggregates, along with corresponding heatmaps quantifying pairwise interaction frequencies between red and green cells. Each image represents an independent experimental replication. Heatmap values reflect the relative frequency of self- (same color) and heterotypic (different color) interactions, normalized to row and column totals. **e-h**, Similar analysis for snowflake yeast displaying surface adhesion proteins Ag3 (red) and Nb3 (green) after differentiation, enabling heterotypic adhesion via Ag3-Nb3 binding. **e**, Circuit schematic showing fluorescent and adhesion markers. **f-h**, Representative images and interaction frequency heatmaps from three independent samples. **i-l**, Three-color BY4741 yeast system with differentiation into red (Ag3), blue (Nb1+Nb3), and green (Ag1) subpopulations. **i**, Circuit schematic showing differentiation into three adhesive cell types. **j-l**, Representative images and corresponding heatmaps quantifying all pairwise interaction types in three biological replicates.

**Extended Data Fig. 6.**
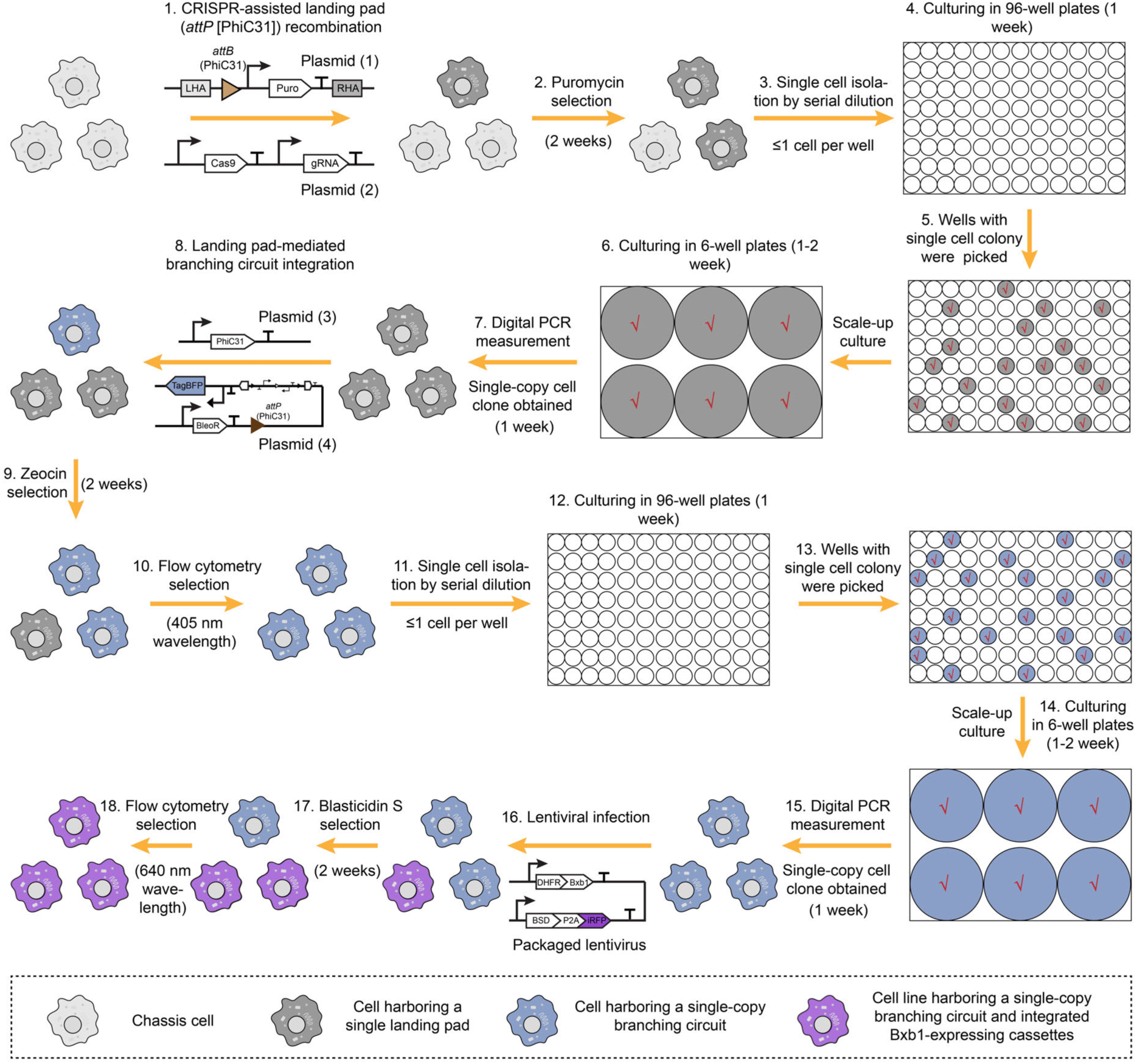
Stepwise workflow for constructing mammalian cell lines with single-copy, recombinase-catalytic branching circuits. The figure illustrates the complete workflow for engineering CHO-K1 or HEK293FT cell lines that stably carry single-copy synthetic differentiation circuits under the control of inducible recombinases. The process begins with CRISPR-Cas9-assisted knock-in of a landing pad cassette into a defined safe harbor locus. The landing pad contains a PhiC31 *attB* site and a puromycin resistance gene (PuroR). Following 2 weeks of puromycin selection, single-cell clones are isolated via limiting dilution into 96-well plates. Clones that form visible colonies are further expanded in 6-well plates and screened by digital PCR to identify those harboring a single landing pad copy. These validated clones are then subjected to PhiC31-mediated site-specific integration of the synthetic branching circuit, which contains a PhiC31 *attP* site, a bleomycin resistance gene (BleoR), and a TagBFP reporter. Zeocin selection enriches successfully recombined cells, and the brightest TagBFP-positive population is isolated by flow cytometry. Due to possible off-target insertions and sorting bias, an additional round of clonal isolation and digital PCR is performed to recover cells with a single integrated copy of the branching circuit. Finally, lentiviral vectors encoding an inducible recombinase (e.g., Bxb1 or DHFR-Bxb1) are delivered to these cells. After blasticidin (BSD) selection and flow cytometric enrichment of iRFP-positive strains, the resulting cell lines stably express a single-copy differentiation circuit along with tightly regulated recombinase expression.

**Extended Data Fig. 7.**
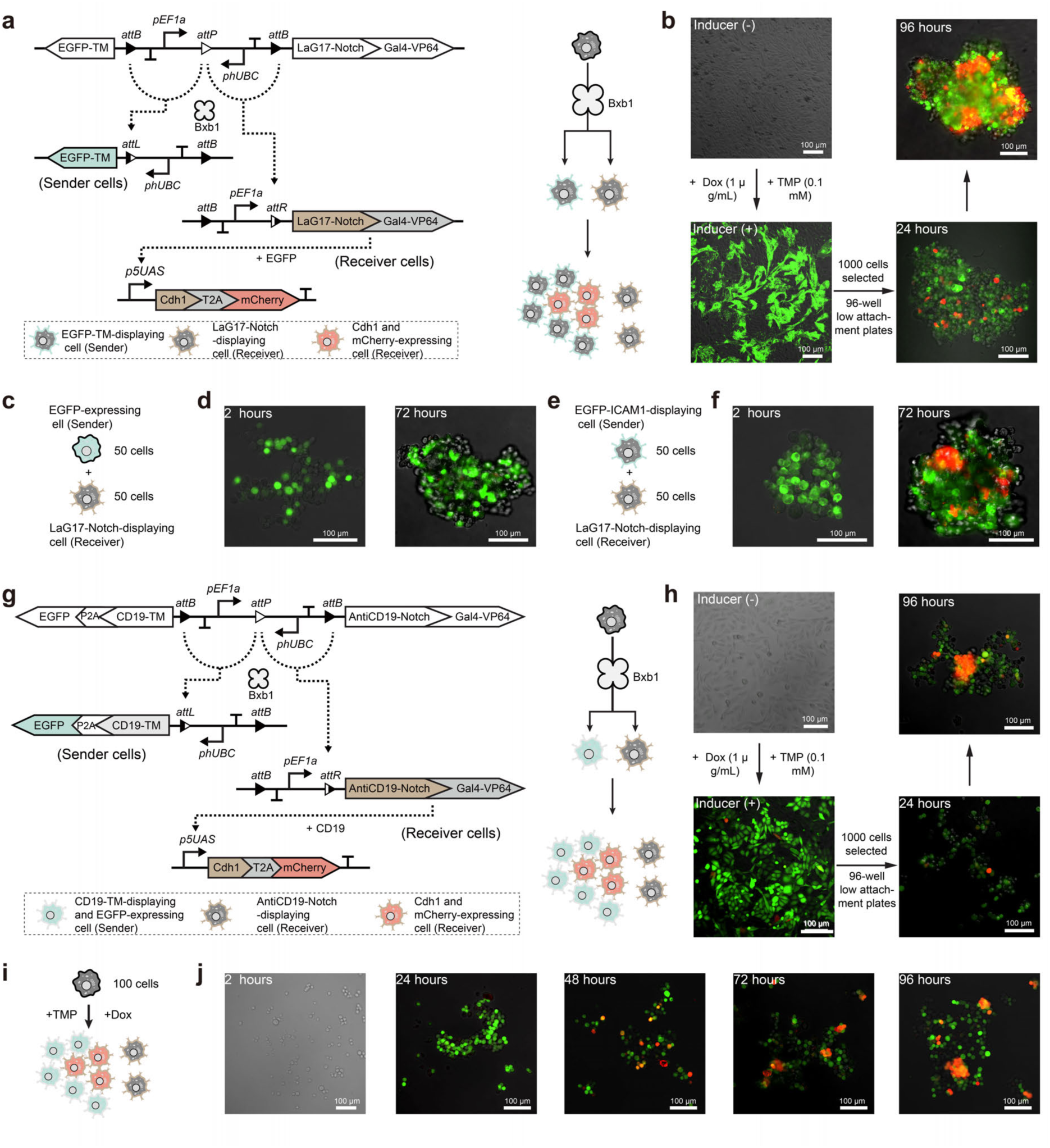
Recombinase-mediated differentiation programs spatial patterning through synNotch-based contact signaling in CHO-K1 cells. **a**, Schematic of a synthetic gene circuit combining Bxb1-mediated differentiation with synNotch signaling. Upon recombination, CHO-K1 founder cells produce two populations: sender cells expressing membrane-bound EGFP, and receiver cells displaying LaG17-Notch fused to Gal4-VP64. All cells carry a Cdh1-T2A-mCherry cassette under a Gal4-responsive promoter *p5UAS*, activated upon synNotch signaling. **b**, Circuit validation and aggregation assay. Without inducers, no fluorescence was observed, indicating tight regulation and minimal leakiness. Upon induction, EGFP localized to the membrane of partial progenies. Cells were dissociated and seeded at low density into ultra-low-attachment plates. Aggregates formed within 24- 96 hours, exhibiting contact-dependent mCherry expression and self-organized patterning. **c-d**, Control co-culture of cytoplasmic EGFP-expressing sender cells and LaG17-Notch receiver cells. No mCherry expression or aggregate formation was observed, confirming that synNotch activation strictly requires membrane-tethered ligand presentation and direct cell-cell contact. **e-f**, Co-culture with sender cells displaying membrane-anchored EGFP-ICAM1 successfully induced mCherry expression in receiver cells, confirming that both physical contact and surface localization of the ligand are sufficient to activate the synNotch pathway and drive downstream cell aggregation. **g-h**, Schematic of a synthetic gene circuit integrating recombinase-based differentiation and synNotch signaling. Sender cells express CD19-TM with EGFP; receivers express antiCD19-Notch. Contact-dependent mCherry and Cdh1 expression was observed. **i-j**, Time course of synNotch activation and spatial pattern formation in 100- cell cultures.

**Extended Data Fig. 8.**
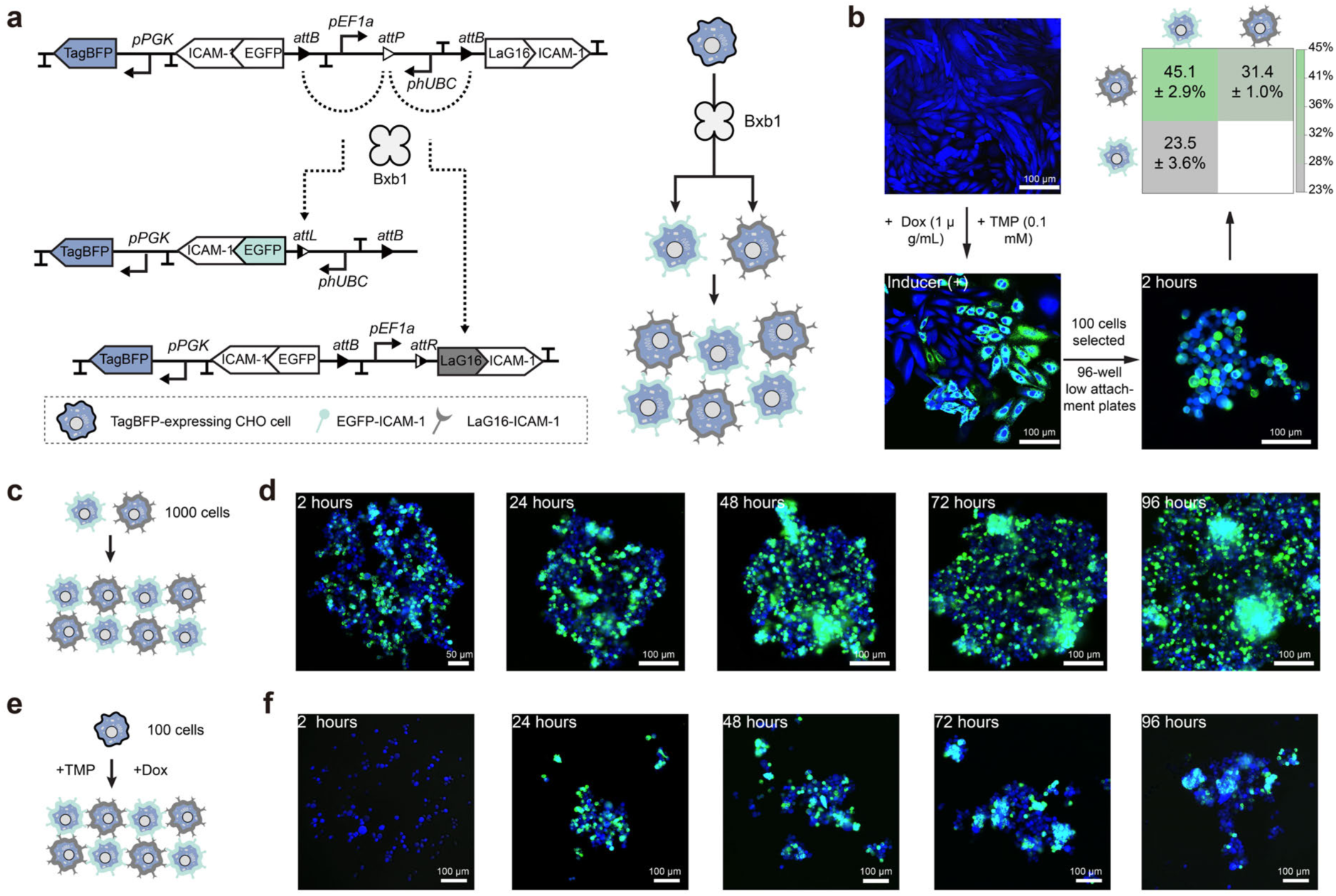
Recombinase-mediated differentiation and EGFP-LaG16-driven multicellular assembly in CHO-K1 cells. **a**, Schematic of the genetic circuit used to generate cell-type-specific adhesion. TagBFP-positive CHO-K1 founder cells undergo Bxb1-mediated recombination, producing two differentiated populations: one expressing EGFP fused to ICAM-1, and the other expressing LaG16 (a designed anti-EGFP nanobody) fused to ICAM-1. These engineered surface proteins enable specific heterotypic adhesion via EGFP-LaG16 interactions. **b**, Experimental validation of circuit behavior and aggregation. In the absence of inducers (1 μg/mL doxycycline and 0.1 mM trimethoprim), only TagBFP fluorescence was observed (top left), confirming minimal background recombination. Upon induction, membrane-localized EGFP fluorescence emerged (bottom left), indicating successful recombinase activation and protein display. Differentiated cells (n = 100) were seeded into ultra-low-attachment 96-well plates. Cell aggregation was observed 2 hours after seeding, and heatmaps quantifying cell-cell interactions revealed a higher frequency of contacts between the two differentiated cell types. Data are mean ± s.d. from three independent biological replicates. **c-d**, Aggregation kinetics of 1,000 pre-induced differentiated cells in culture, imaged at multiple time points over 96 hours. EGFP-positive and LaG16-positive cells progressively assembled into stable multicellular structures. **e-f**, In situ differentiation and aggregation from 100 founder cells following induction with doxycycline and TMP. Multicellular assemblies resembling those formed in pre-induced populations were observed within 96 hours, validating inducibility and functional adhesion of newly differentiated cells.

**Extended Data Fig. 9.**
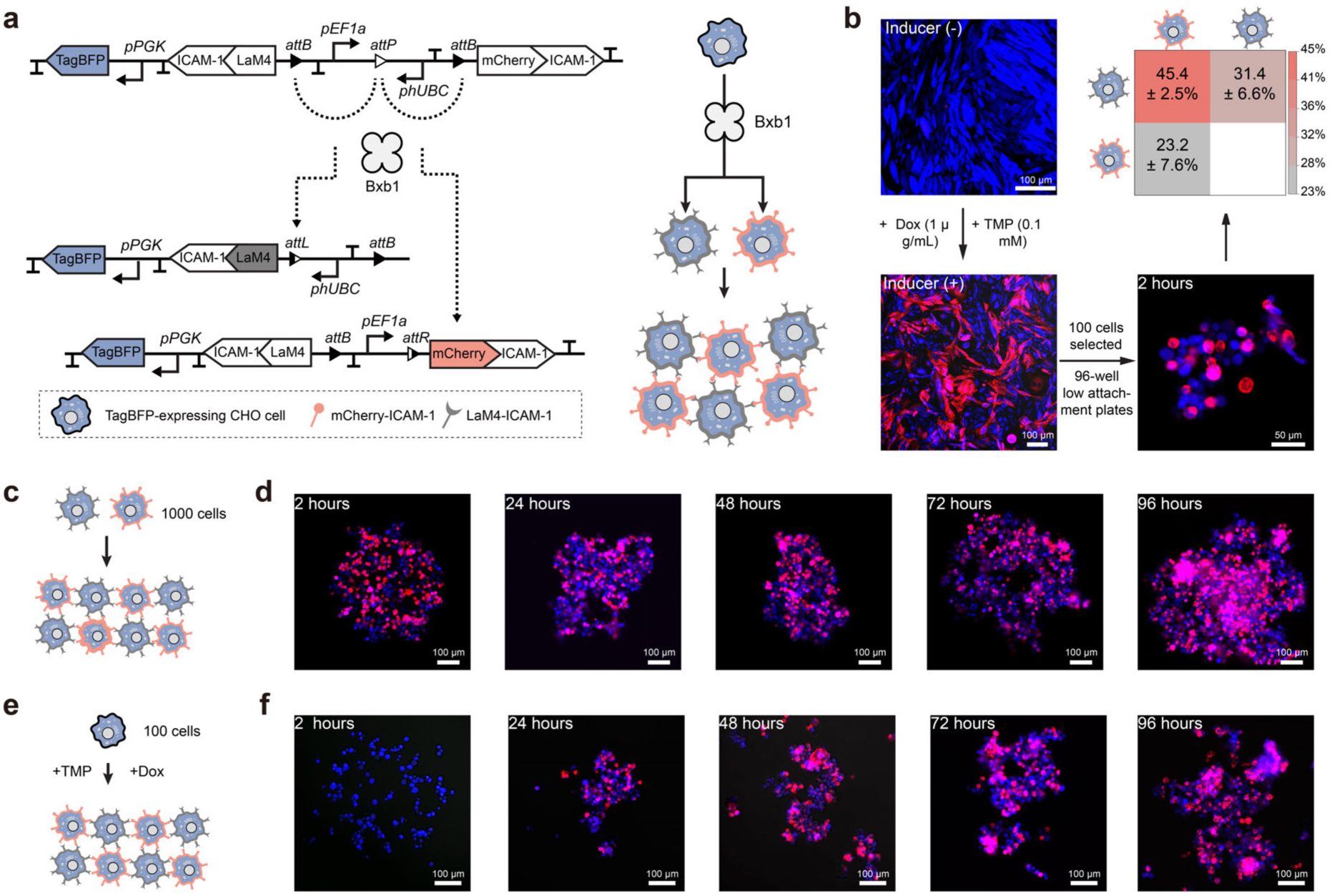
Recombinase-mediated differentiation drives programmable multicellular assembly via mCherry-LaM4 interactions in CHO-K1 cells. **a**, Schematic of the engineered genetic circuit enabling cell-type-specific adhesion. CHO-K1 founder cells stably expressing TagBFP were engineered to undergo Bxb1-mediated recombination, generating two differentiated populations: one expressing mCherry fused to ICAM-1 and the other expressing LaM4 (an anti-mCherry nanobody) fused to ICAM-1. The two cell types are designed to selectively bind through mCherry-LaM4 interactions on the cell surface. **b**, Experimental validation of circuit function and cell aggregation. In the absence of inducers (1 μg/mL Dox and 0.1 mM TMP), only TagBFP fluorescence was detected, indicating tight transcriptional control and negligible recombinase leakiness (top left). Upon induction, red fluorescence localized to the cell membrane was observed, confirming successful circuit activation and surface expression of mCherry (bottom left). Induced cells were enzymatically dissociated and seeded at low density (100 cells per well) into ultra-low-attachment 96-well plates. Cell aggregation was observed 2 hours after seeding, and heatmaps quantifying cell-cell interactions revealed a higher frequency of contacts between the two differentiated cell types. Data are mean±s.d. from 3 independent biological replicates. **c-d,** Aggregation dynamics of 1,000 pre-induced differentiated cells cultured in medium. Red and blue fluorescent cells progressively coalesced into structured clusters over 96 hours. **e-f**, In situ induction of differentiation and aggregation from 100 founder cells exposed to Dox and TMP. Mixed aggregates formed de novo over 96 hours, recapitulating the differentiation and adhesion-driven assembly observed with pre-induced populations.

**Extended Data Fig. 10.**
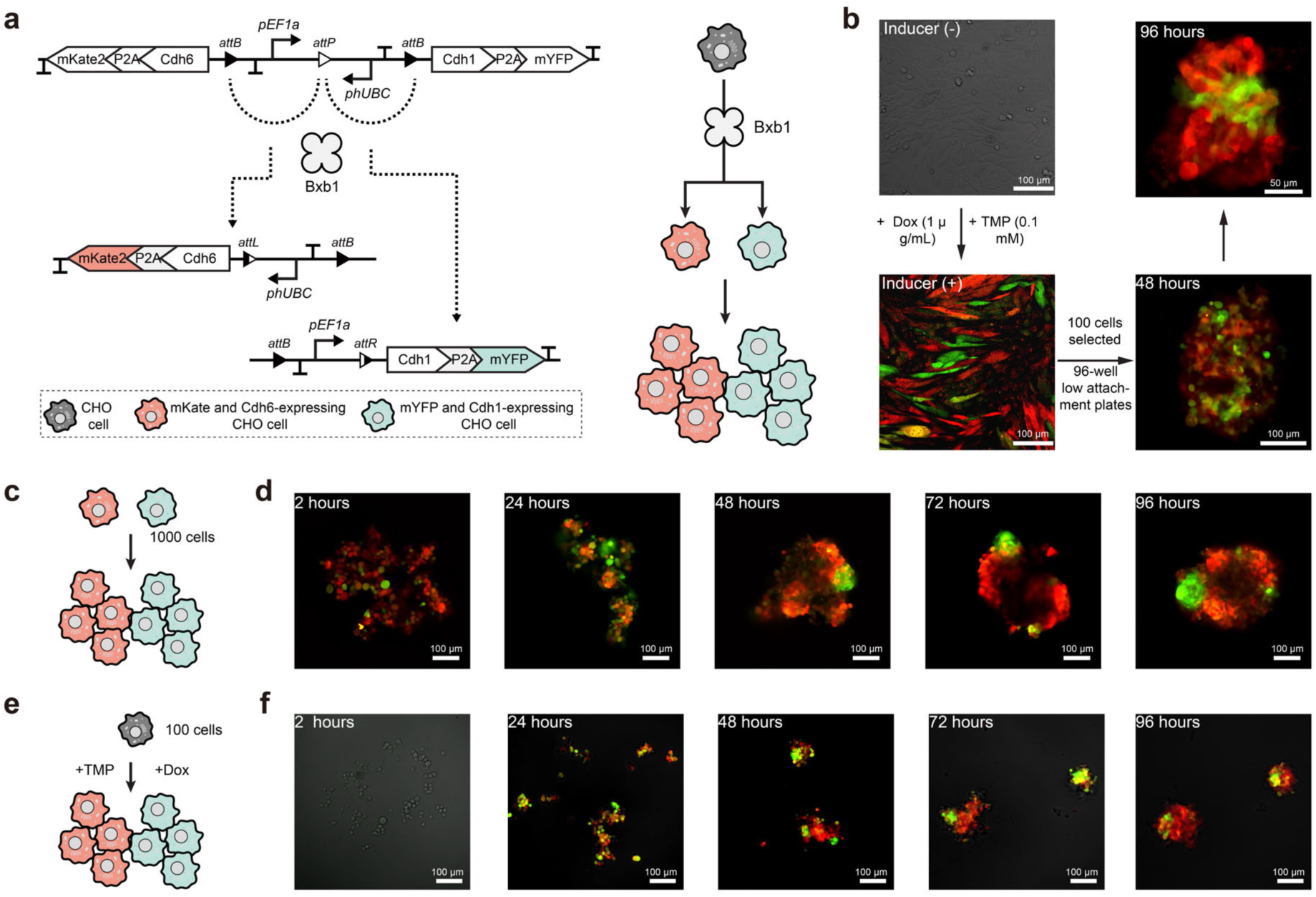
Recombinase-directed expression of cadherin adhesion proteins enables multicellular assembly in CHO-K1 cells. **a**, Schematic of the genetic circuit for recombinase-mediated differentiation in CHO-K1 cells. Upon Bxb1-mediated recombination, founder cells differentiate into two populations: one expressing mKate2 and Cadherin-6 (Cdh6), and the other expressing mYFP and Cadherin-1 (Cdh1). These classical cadherins mediate homophilic adhesion, facilitating specific cell-cell binding. **b**, Experimental validation of circuit function and induced aggregation. Top left: without inducers (1 μg/mL doxycycline and 0.1 mM TMP), no fluorescence is observed, indicating minimal recombinase leakiness. Bottom left: upon induction, both mKate2- and mYFP-expressing cells emerge. Right: 100 differentiated cells were transferred to ultra-low-attachment 96-well plates, with multicellular clusters forming and increasing in size and complexity over time. **c-d**, Suspension culture of 1,000 pre-induced differentiated cells monitored at 2-96 hours. Red and green cells progressively aggregated, forming dense multicellular assemblies. **e-f**, In situ induction of 100 founder cells followed by imaging over 96 hours. Gradual expression of Cdh6 and Cdh1 promoted cell-cell adhesion and the emergence of organized cluster morphologies.

