## Supplementary Information for "Synthetic ratio computation for programming population composition and multicellular morphology"

**Supplementary figures**

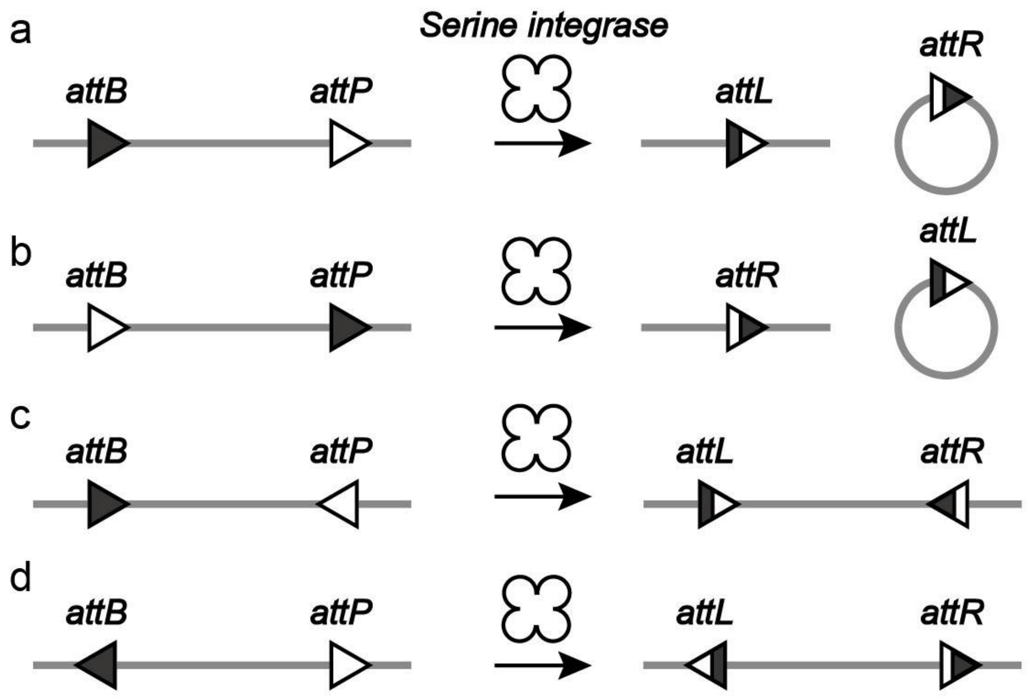

**Fig. S1 | Recombination patterns of serine recombinase**. Serine integrase/recombinase specifically recognizes two types of *att* sites, *attB* and *attP*. When aligned in the same orientation (**Fig. S1a** and **Fig. S1b**), the serine recombinase facilitates DNA excision by removing sequences between the *attB* and *attP* sites, producing new *attL* and *attR* sites. Conversely, oppositely oriented *attB* and *attP* sites (**Fig. S1c** and **Fig. S1d**) led to DNA inversion, flipping the DNA between *att* sites.

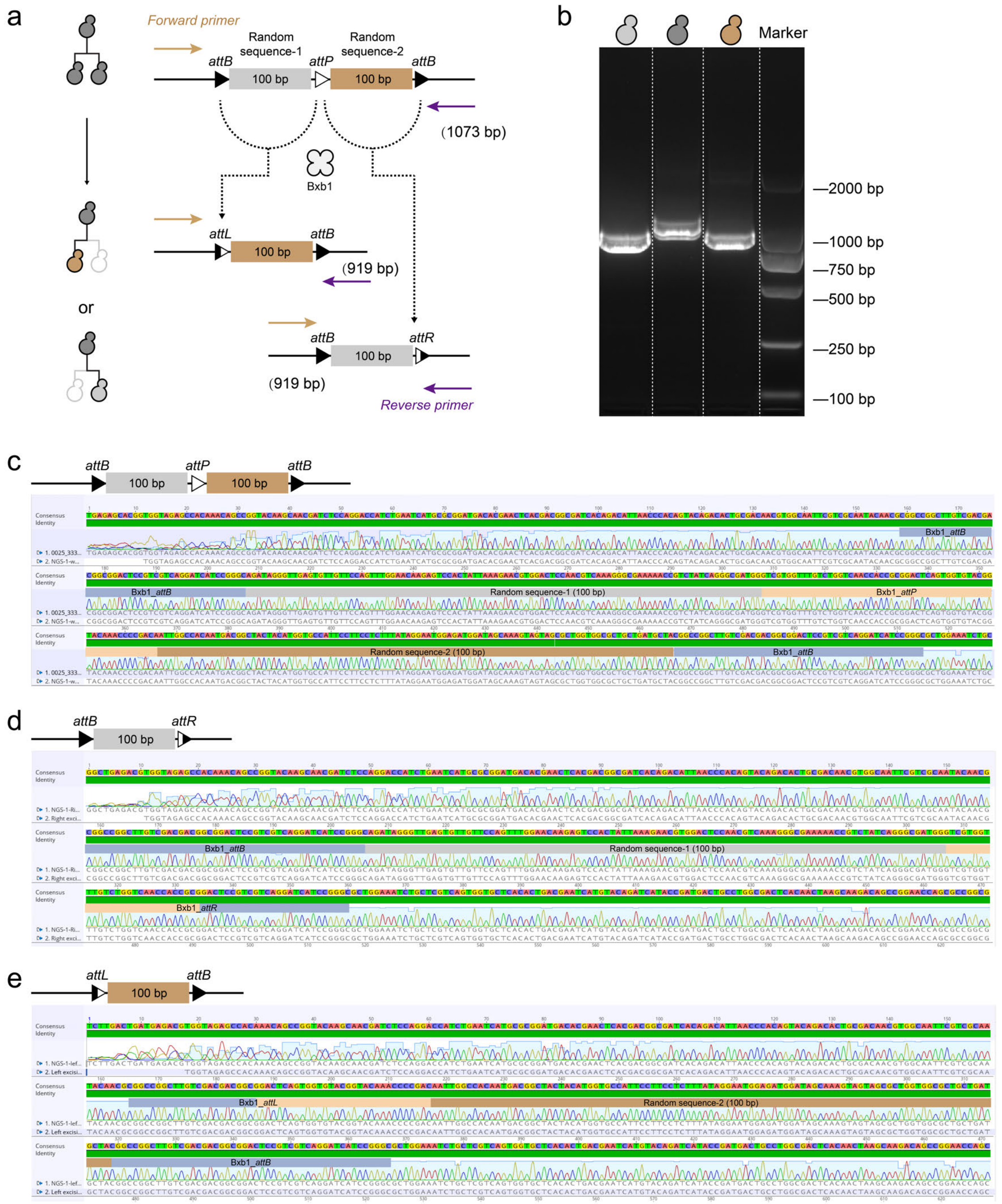

**Fig. S2 | Validation of Bxb1-mediated site-specific recombination by Sanger sequencing.** **a**, Schematic of the synthetic recombination substrate used for sequencing analysis. The initial construct contained two 100 bp non-coding spacers separating one *attP* site and two *attB* sites arranged in the configuration *attB-attP-attB*. Bxb1-mediated recombination between *attP* and either *attB* site generates two distinct products: one yielding *attL-attB* and the other yielding *attB-attR* junctions. **b**, Agarose gel electrophoresis showing distinct PCR products. Band sizes match the expected lengths of parental and recombined products. **c-e**, Aligned Sanger sequencing results before and after Bxb1 catalysis.

a

| | | FAM(pBranch-E. coli) | | | | | | VIC(RecA) | | | | | |
|---|---|---|---|---|---|---|---|---|---|---|---|---|---|
| Sample Name | Nb Droplets | C(cp/ul) | Nb Pos | Nb Neg | C_min(cp/ul) | C_max(cp/ul) | Relative Uncertainly(%) | C(cp/ul) | Nb Pos | Nb Neg | C_min(cp/ul) | C_max(cp/ul) | Relative Uncertainly(%) |
| E.coli-1 | 21241 | 138.77 | 1697 | 19544 | 132.18 | 145.39 | 4.76 | 217.97 | 2604 | 18637 | 209.62 | 226.37 | 3.84 |
| E.coli-2 | 21374 | 214.4 | 2580 | 18794 | 206.14 | 222.7 | 3.86 | 189.22 | 2294 | 19080 | 181.49 | 196.99 | 4.09 |
| E.coli-3 | 21612 | 209.1 | 2411 | 18318 | 203.56 | 211.9 | 4.02 | 193.32 | 2336 | 19314 | 183.63 | 198.19 | 4.21 |
| E.coli-4 | 20821 | 156.09 | 2689 | 20132 | 151.9 | 160.28 | 4.48 | 186.87 | 2140 | 17080 | 134.8 | 178.9 | 3.71 |

| PCR reaction system(μL) | Detection volume | Dilution factor | The copy number of the target gene pBranch per microliter of extracted DNA | The copy number of the target gene RecA (Haploid) per microliter of extracted DNA | pBranch/RecA (Haploid) | pBranch copy number |
|---|---|---|---|---|---|---|
| 20 | 2 | 1000 | 1.39E+06 | 2.18E+06 | 0.64 | 0.64 |
| 20 | 2 | 1000 | 2.14E+06 | 1.89E+06 | 1.13 | 1.13 |
| 20 | 2 | 1000 | 2.09E+06 | 1.93E+06 | 1.08 | 1.08 |
| 20 | 2 | 1000 | 1.56E+06 | 1.87E+06 | 0.84 | 0.84 |

b

| | | FAM(pBranch-Yeast) | | | | | | VIC(RPB2) | | | | | |
|---|---|---|---|---|---|---|---|---|---|---|---|---|---|
| Sample Name | Nb Droplets | C(cp/ul) | Nb Pos | Nb Neg | C_min(cp/ul) | C_max(cp/ul) | Relative Uncertainly(%) | C(cp/ul) | Nb Pos | Nb Neg | C_min(cp/ul) | C_max(cp/ul) | Relative Uncertainly(%) |
| Yeast-1 | 20927 | 15.6 | 195 | 20732 | 13.41 | 17.79 | 14.04 | 15.68 | 196 | 20731 | 13.49 | 17.88 | 14 |
| Yeast-2 | 20521 | 15.01 | 184 | 20337 | 12.84 | 17.18 | 14.45 | 15.67 | 192 | 20329 | 13.45 | 17.88 | 14.15 |
| Yeast-3 | 20817 | 14.64 | 182 | 20635 | 12.51 | 16.76 | 14.53 | 15.2 | 189 | 20628 | 13.04 | 17.37 | 14.26 |
| Yeast-4 | 20216 | 17.4 | 210 | 20006 | 15.05 | 19.76 | 13.53 | 16.24 | 196 | 20020 | 13.97 | 18.51 | 14 |

| PCR reaction system(μL) | Detection volume | Dilution factor | The copy number of the target gene pBranch per microliter of extracted DNA | The copy number of the target gene RPB2 (Haploid) per microliter of extracted DNA | pBranch/RPB2 (Haploid) | pBranch copy number |
|---|---|---|---|---|---|---|
| 20 | 2 | 100 | 1.56E+04 | 1.57E+04 | 0.99 | 0.99 |
| 20 | 2 | 100 | 1.50E+04 | 1.57E+04 | 0.96 | 0.96 |
| 20 | 2 | 100 | 1.46E+04 | 1.52E+04 | 0.96 | 0.96 |
| 20 | 2 | 100 | 1.74E+04 | 1.62E+04 | 1.07 | 1.07 |

c

| | | FAM(pBranch-HEK293FT) | | | | | | VIC(RPPH1) | | | | | |
|---|---|---|---|---|---|---|---|---|---|---|---|---|---|
| Sample Name | Nb Droplets | C(cp/ul) | Nb Pos | Nb Neg | C_min(cp/ul) | C_max(cp/ul) | Relative Uncertainly(%) | C(cp/ul) | Nb Pos | Nb Neg | C_min(cp/ul) | C_max(cp/ul) | Relative Uncertainly(%) |
| HEK293FT-1 | 20924 | 99.7 | 1215 | 19709 | 94.1 | 105.32 | 5.62 | 174.86 | 2084 | 18840 | 167.36 | 182.39 | 4.3 |
| HEK293FT-2 | 21104 | 86.05 | 1062 | 20042 | 80.89 | 91.24 | 6.02 | 146.34 | 1774 | 19330 | 139.54 | 153.17 | 4.66 |
| HEK293FT-3 | 20643 | 58.5 | 712 | 19931 | 54.21 | 62.8 | 7.35 | 104.45 | 1254 | 19389 | 98.68 | 110.24 | 5.54 |
| HEK293FT-4 | 20948 | 128.12 | 1550 | 19398 | 121.75 | 134.51 | 4.98 | 220.78 | 2599 | 18349 | 212.31 | 229.3 | 3.85 |

| PCR reaction system(μL) | Detection volume | Dilution factor | The copy number of the target gene pBranch per microliter of extracted DNA | The copy number of the target gene RPPH1 (Diploid) per microliter of extracted DNA | pBranch/RPPH1 (Diploid) | pBranch copy number |
|---|---|---|---|---|---|---|
| 20 | 2 | 1000 | 9.97E+05 | 1.75E+06 | 0.57 | 1.14 |
| 20 | 2 | 1000 | 8.61E+05 | 1.46E+06 | 0.59 | 1.18 |
| 20 | 2 | 1000 | 5.85E+05 | 1.04E+06 | 0.56 | 1.12 |
| 20 | 2 | 1000 | 1.28E+06 | 2.21E+06 | 0.58 | 1.16 |

d

| | | FAM(pBranch-CHO cells) | | | | | | VIC(GAPDH) | | | | | |
|---|---|---|---|---|---|---|---|---|---|---|---|---|---|
| Sample Name | Nb Droplets | C(cp/ul) | Nb Pos | Nb Neg | C_min(cp/ul) | C_max(cp/ul) | Relative Uncertainly(%) | C(cp/ul) | Nb Pos | Nb Neg | C_min(cp/ul) | C_max(cp/ul) | Relative Uncertainly(%) |
| CHO-1 | 20459 | 566.56 | 5896 | 14563 | 552.1 | 581.16 | 2.56 | 1072.68 | 9710 | 10749 | 1051.12 | 1094.53 | 2.02 |
| CHO-2 | 20797 | 594.22 | 6237 | 14560 | 579.46 | 609.11 | 2.5 | 1248.61 | 10965 | 9832 | 1224.86 | 1272.71 | 1.92 |
| CHO-3 | 20904 | 586.25 | 6199 | 14705 | 571.65 | 600.99 | 2.5 | 1226.43 | 10889 | 10015 | 1203.03 | 1250.15 | 1.92 |
| CHO-4 | 20791 | 584.72 | 6152 | 14639 | 570.1 | 599.47 | 2.51 | 1201.33 | 10679 | 10112 | 1178.21 | 1224.77 | 1.94 |

| PCR reaction system(μL) | Detection volume | Dilution factor | The copy number of the target gene pBranch per microliter of extracted DNA | The copy number of the target gene GAPDH(Diploid) per microliter of extracted DNA | pBranch/GAPDH (Diploid) | pBranch copy number |
|---|---|---|---|---|---|---|
| 20 | 2 | 10 | 5.67E+04 | 1.07E+05 | 0.53 | 1.06 |
| 20 | 2 | 10 | 5.94E+04 | 1.25E+05 | 0.48 | 0.95 |
| 20 | 2 | 10 | 5.86E+04 | 1.23E+05 | 0.48 | 0.96 |
| 20 | 2 | 10 | 5.85E+04 | 1.20E+05 | 0.49 | 0.97 |

**Fig. S3 | Copy number determination of the branching circuit in *E. coli,* haploid yeast BY4741, and mammalian cells (HEK293FT and CHO-K1). a**, Using the *E. coli* chromosomal *recA* gene as an internal reference, digital PCR analysis showed a circuit copy number of 0.92 ± 0.20 across all biological replicates (mean ± s.d., *n* = 4), consistent with single-copy plasmid maintenance. **b**, Using the haploid yeast chromosomal *RPB2* gene as an internal reference, digital PCR analysis showed a circuit copy number of 1.00 ± 0.05 across all biological replicates (mean ± s.d., *n* = 4), also consistent with single-copy plasmid maintenance. **c**, Using the mammalian chromosomal RPPH1 gene as an internal reference, digital PCR analysis showed that the branching circuit copy number relative to *RPPH1* was 0.57 ± 0.01 (mean ± s.d., *n* = 4). Since *RPPH1* is diploid in HEK293FT cells, the circuit copy number across replicates was consistent with single-copy plasmid maintenance. **d**, Using the CHO-K1 chromosomal *GAPDH* gene as an internal reference, digital PCR analysis showed that the branching circuit copy number relative to *GAPDH* was 0.49 ± 0.02 (mean ± s.d., *n* = 4). As *GAPDH* is diploid in CHO-K1 cells, the circuit copy number across replicates was consistent with single-copy plasmid maintenance.

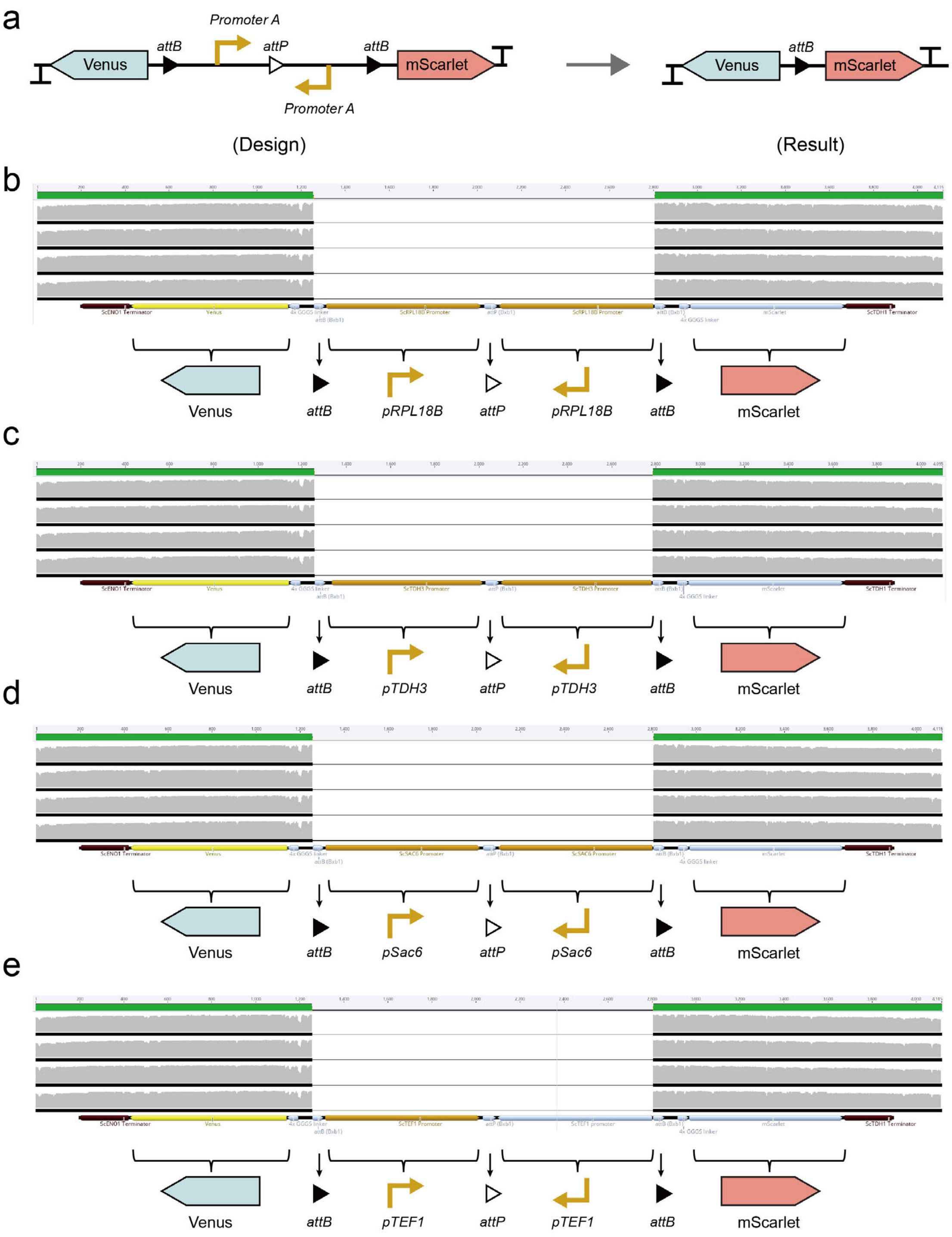

**Fig. S4 | Homologous recombination disrupts circuit design when two identical promoters are used. a**, Schematic representation of the intended genetic circuit (left), in which two identical promoters flank the recombinase site (*attP*). The actual construct obtained in *E. coli* DH5α after plasmid assembly (right) shows deletion of the intermediate region between the promoters, resulting in direct fusion of the upstream and downstream modules (Venus and mScarlet). **b-e**, Representative Sanger sequencing results from four designs using pairs of identical promoters: **b**, *pRPL18B*; **c**, *pTDH3*; **d**, *pSac6*; **e**, *pTEF1*.

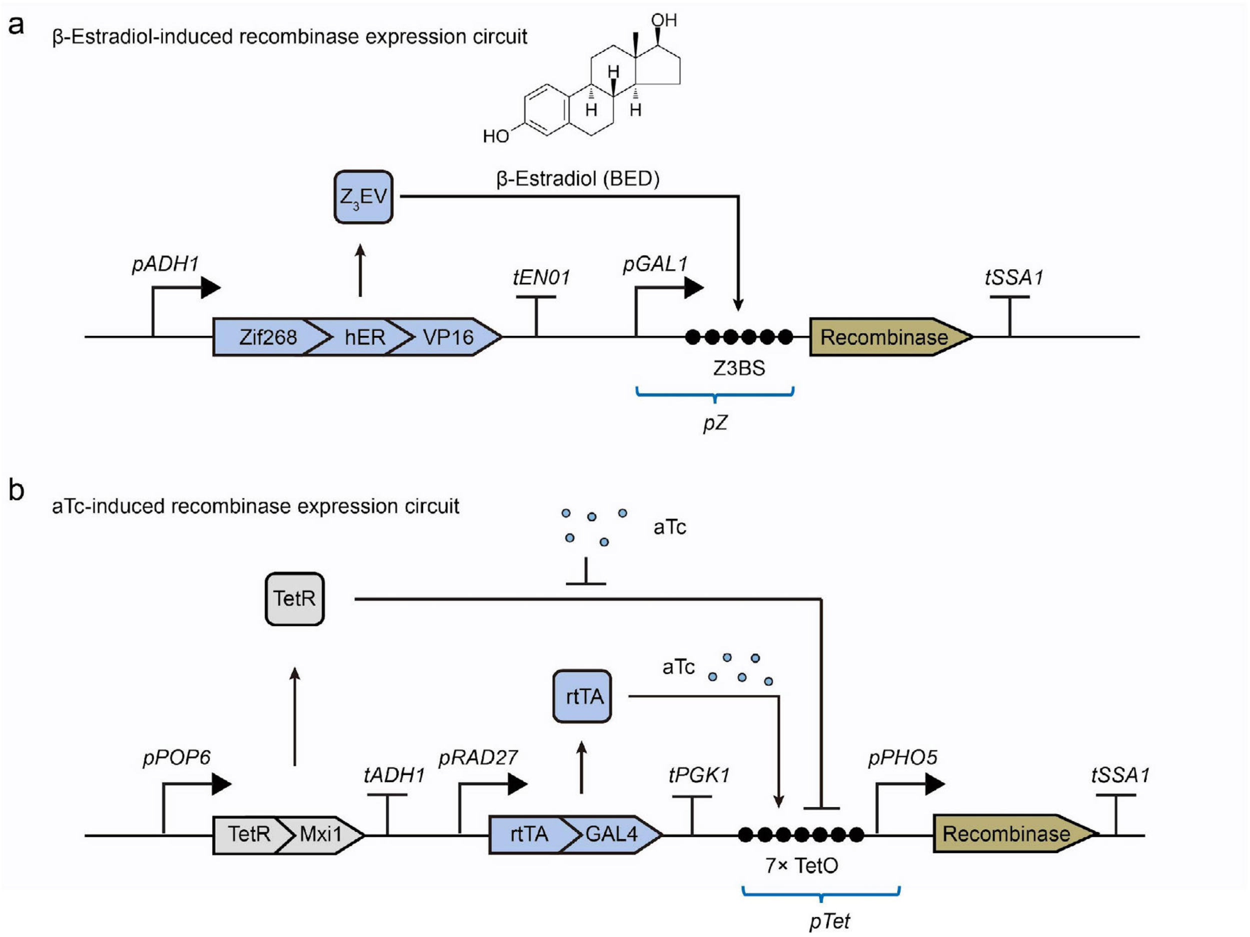

**Fig. S5 | Inducible gene circuits for recombinase expression control in yeast. a.** β-estradiol-induced recombinase expression system[1]. The circuit utilizes the constitutively expressed transcription factor $Z_3EV$ transcription factor in yeast. Upon binding β-estradiol, $Z_3EV$ attaches to the $Z_3BS$ site on the *pZ* promoter, initiating recombinase production. **b.** Anhydrotetracycline (aTc)-induced recombinase expression system[2]. In the absence of aTc, TetR-Mxi1 fusion proteins bind to the TetO operator within the *pTet* promoter, repressing transcription. When aTc is present, it disrupts TetR-TetO binding, allowing the reverse tetracycline transactivator (rtTA) to bind to TetO. The rtTA-GAL4 fusion protein subsequently activates the downstream *pTet* promoter.

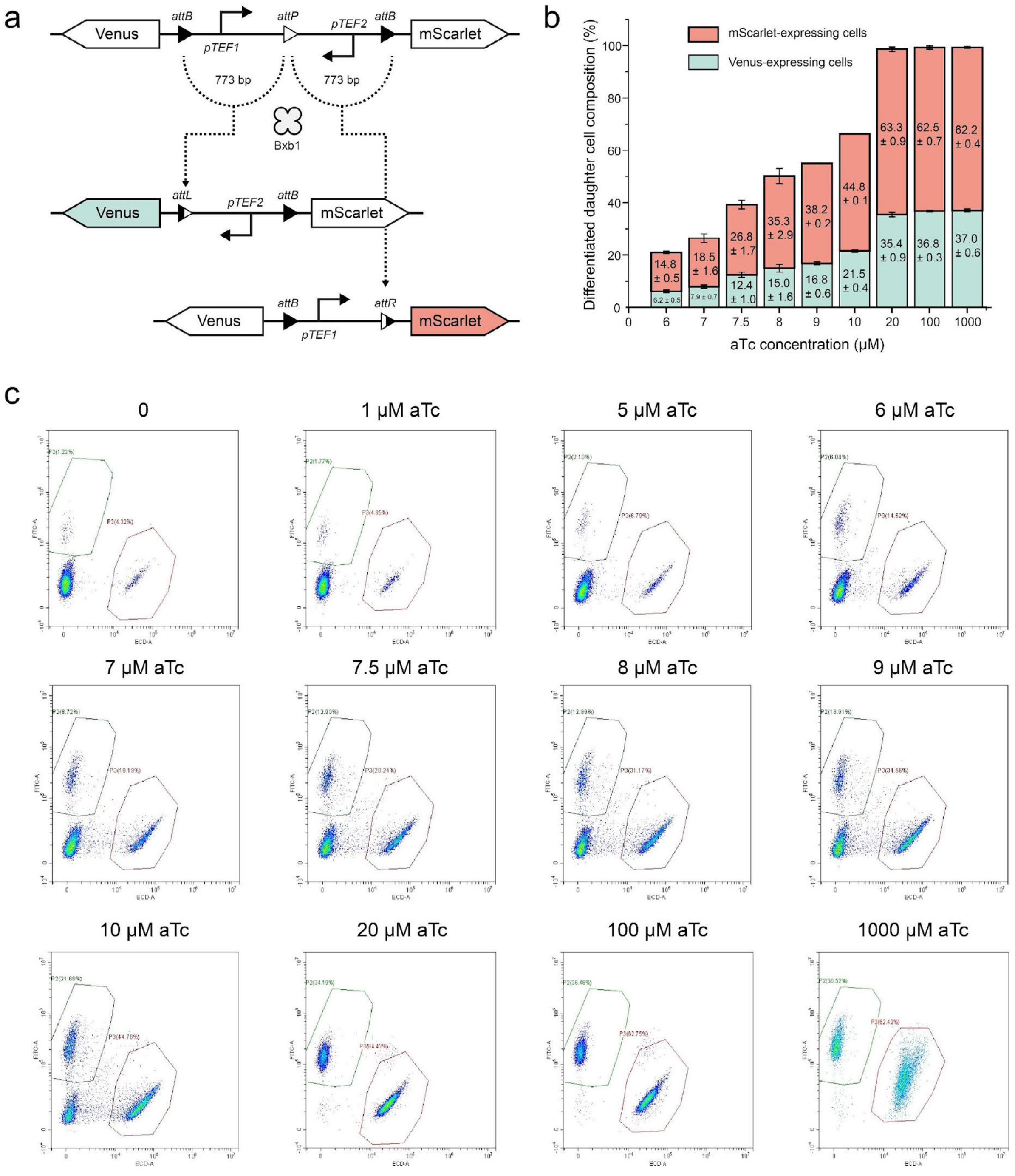

**Fig. S6 | Effect of aTc concentrations on binary cell differentiation. a.** Schematic of the gene circuit. **b.** Statistical analysis of differentiated progeny populations exposed to varying aTc concentrations (data are mean ± s.d. of $n = 8$ replicates). Low aTc concentrations (e.g., 6 μM) resulted in most yeast cells remaining undifferentiated. **c.** Representative flow cytometry data illustrate the correlation between aTc concentrations and differentiation efficiency.

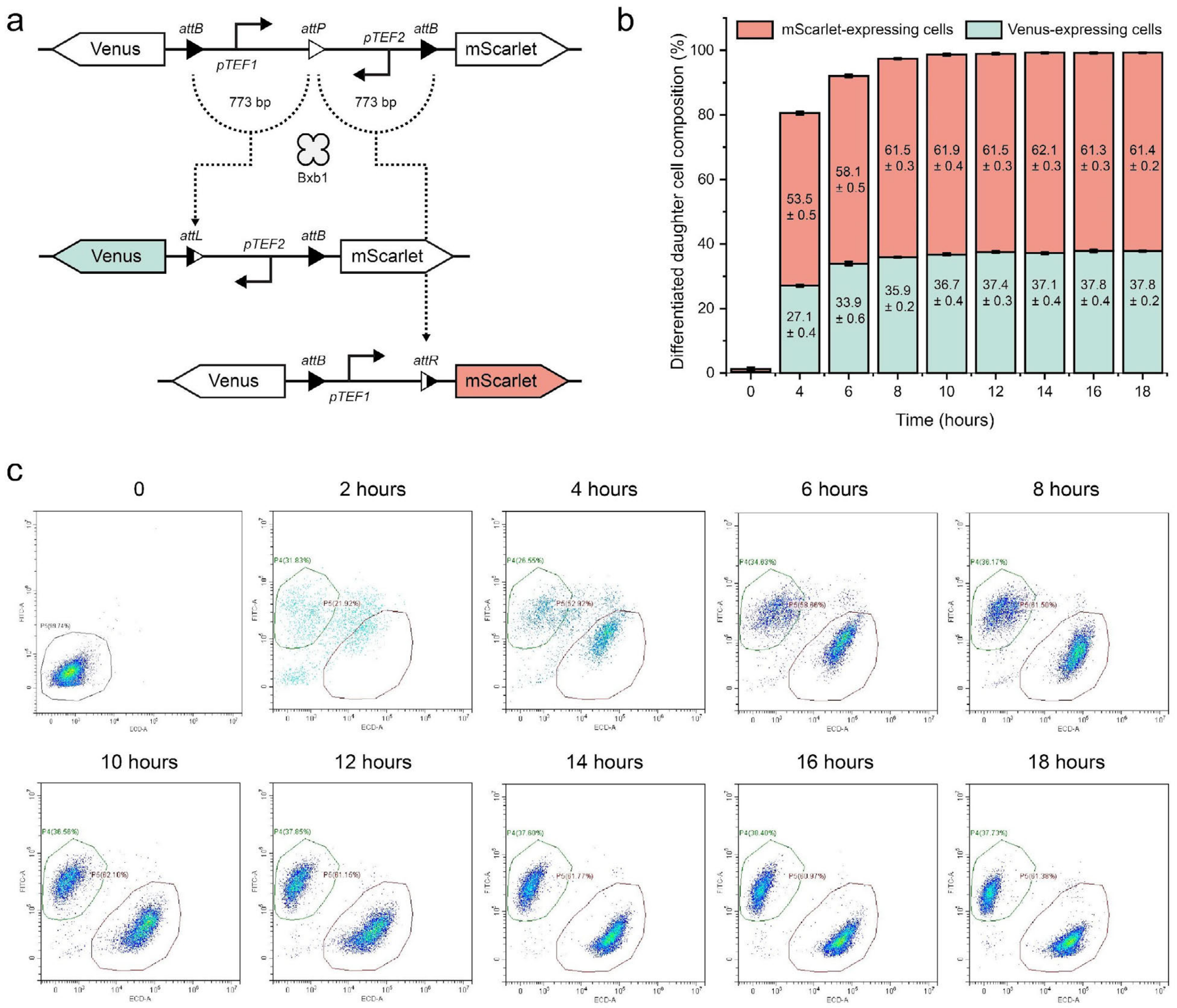

**Fig. S7 | Stable yeast progeny cell populations are generated within 8-10 hours of Bxb1 induction. a.** Gene circuit schematic. **b.** Statistical data showing progeny cell population dynamics over 18 hours with a final aTc inducer concentration of 1 mM (data are mean ± s.d. of *n* = 8 replicates). **c.** Flow cytometry data collected at 2-hour intervals.

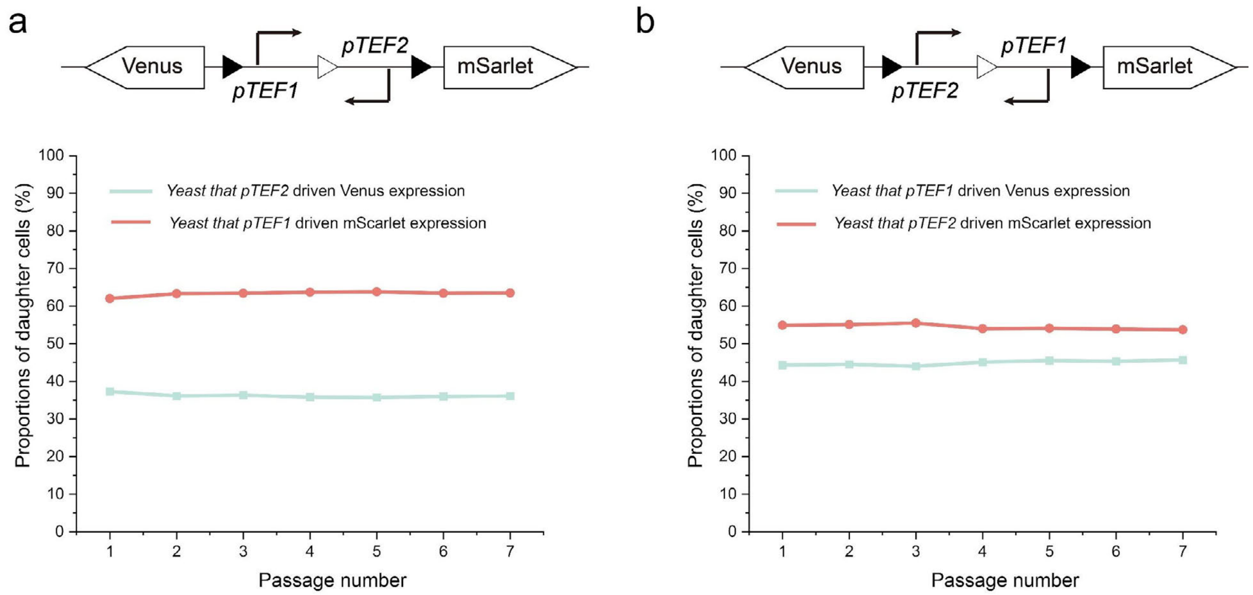

**Fig. S8 | Stability of progeny cell populations in microbial consortia.** Stability progeny ratios were maintained over multiple generations of co-culture: **a**. Red fluorescent yeast (63%) and green fluorescent yeast (37%) remained consistent over seven passages. **b**. Red (55%) and green (45%) fluorescent yeast showed stable ratios over seven days. Data are mean ± s.d. of $n = 4$ replicates.

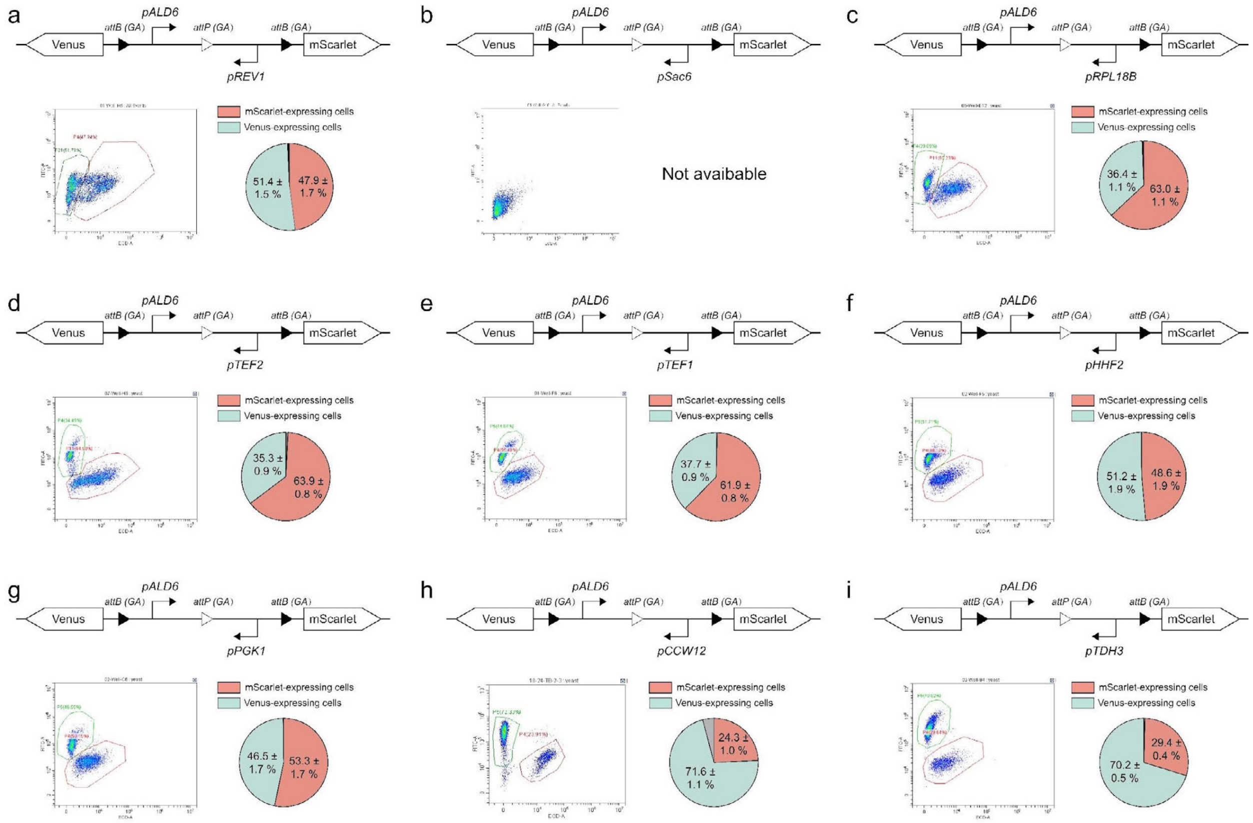

**Fig. S9 | Circuit design with a fixed *pALD6* promoter on the left arm and varying right arms.** Each panel shows the gene circuit design, flow cytometry results, and data on progeny cell distributions (data are mean ± s.d. of $n = 8$ replicates).

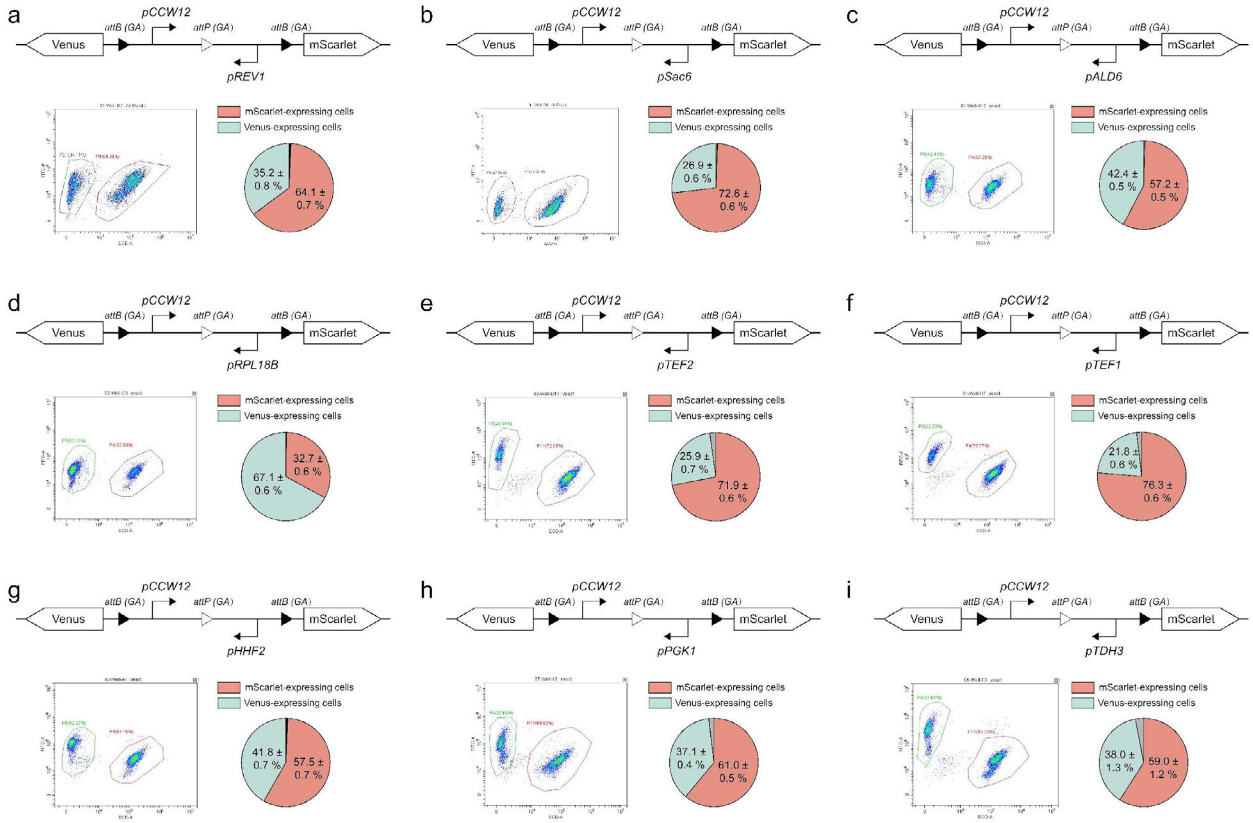

**Fig. S10 | Circuit design with a fixed *pCCW12* promoter on the left arm and varying right arms.** Each panel shows the gene circuit design, flow cytometry results, and data on progeny cell distributions (data are mean ± s.d. of $n$ = 8 replicates).

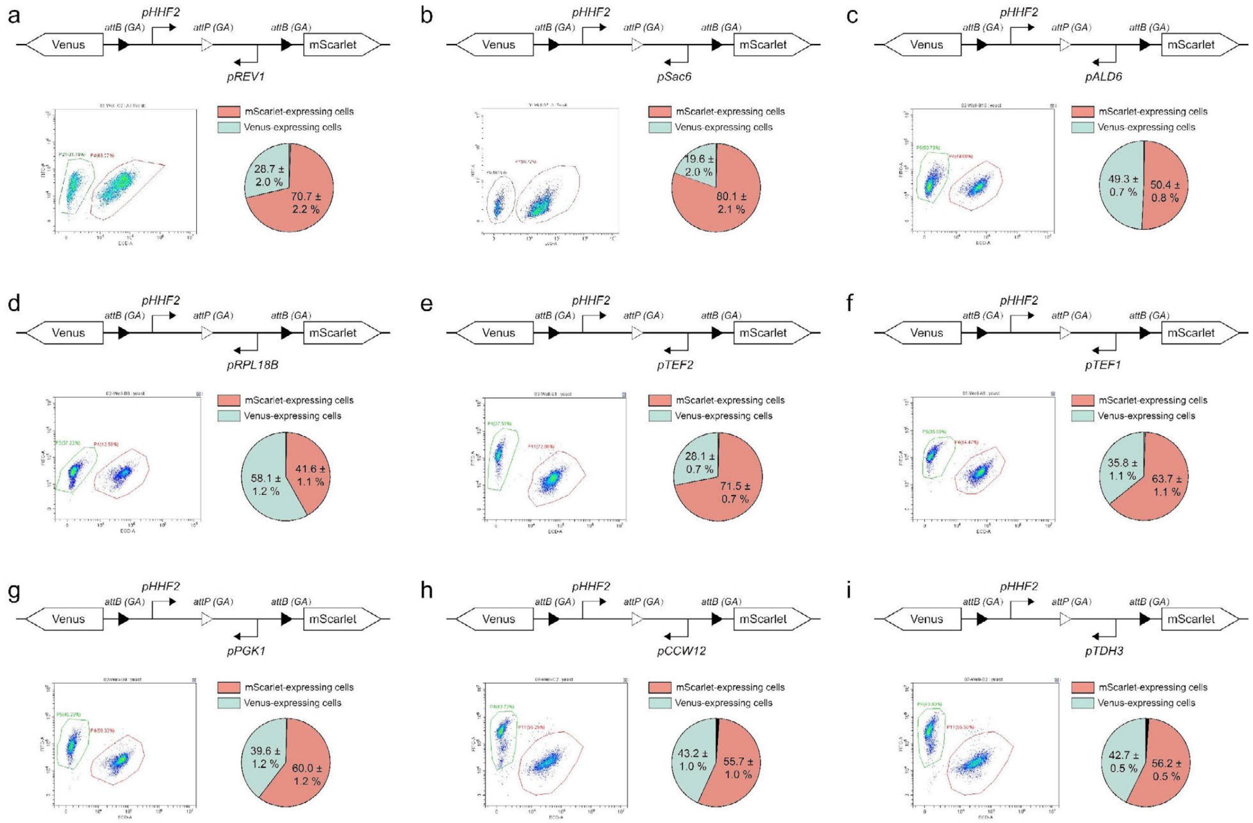

**Fig. S11 | Circuit design with a fixed *pHHF2* promoter on the left arm and varying right arms.** Each panel shows the gene circuit design, flow cytometry results, and data on progeny cell distributions (data are mean ± s.d. of *n* = 8 replicates).

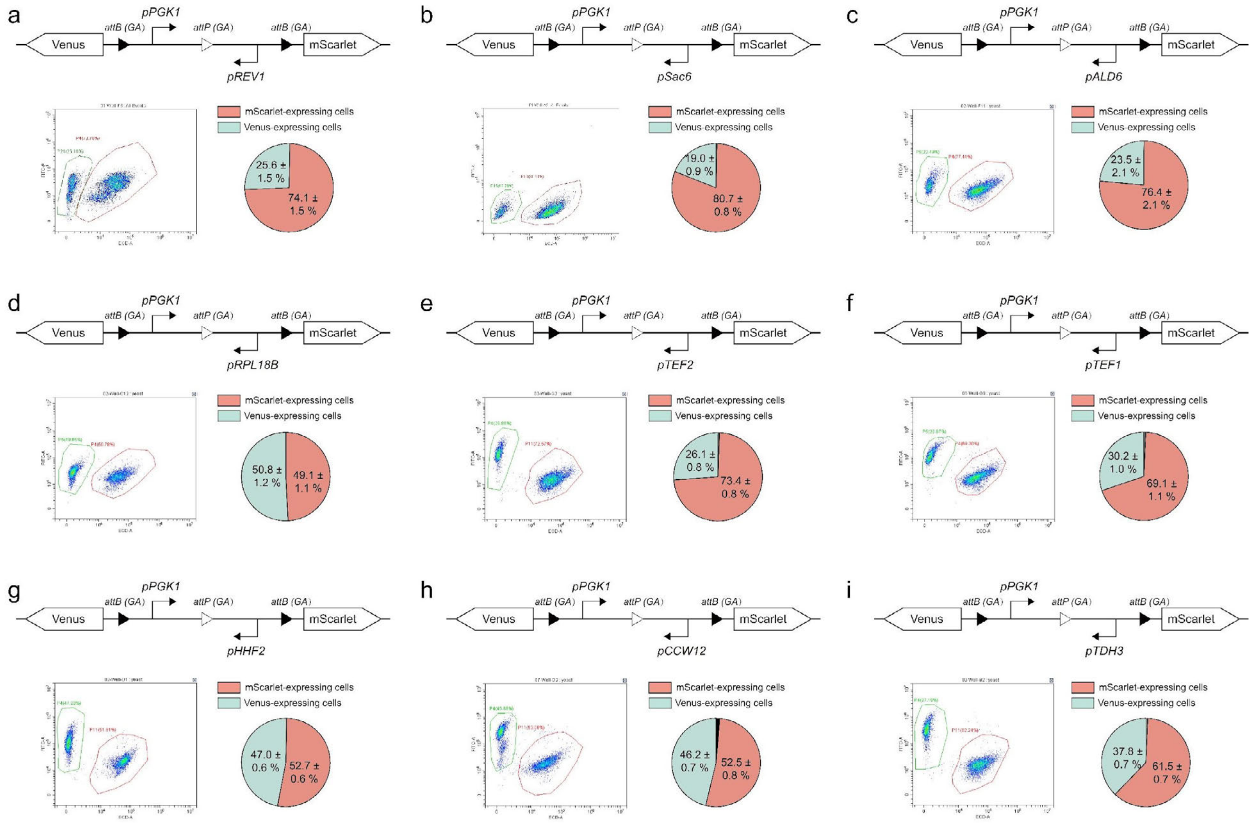

**Fig. S12 | Circuit design with a fixed *pPGK1* promoter on the left arm and varying right arms.** Each panel shows the gene circuit design, flow cytometry results, and data on progeny cell distributions (data are mean ± s.d. of $n$ = 8 replicates).

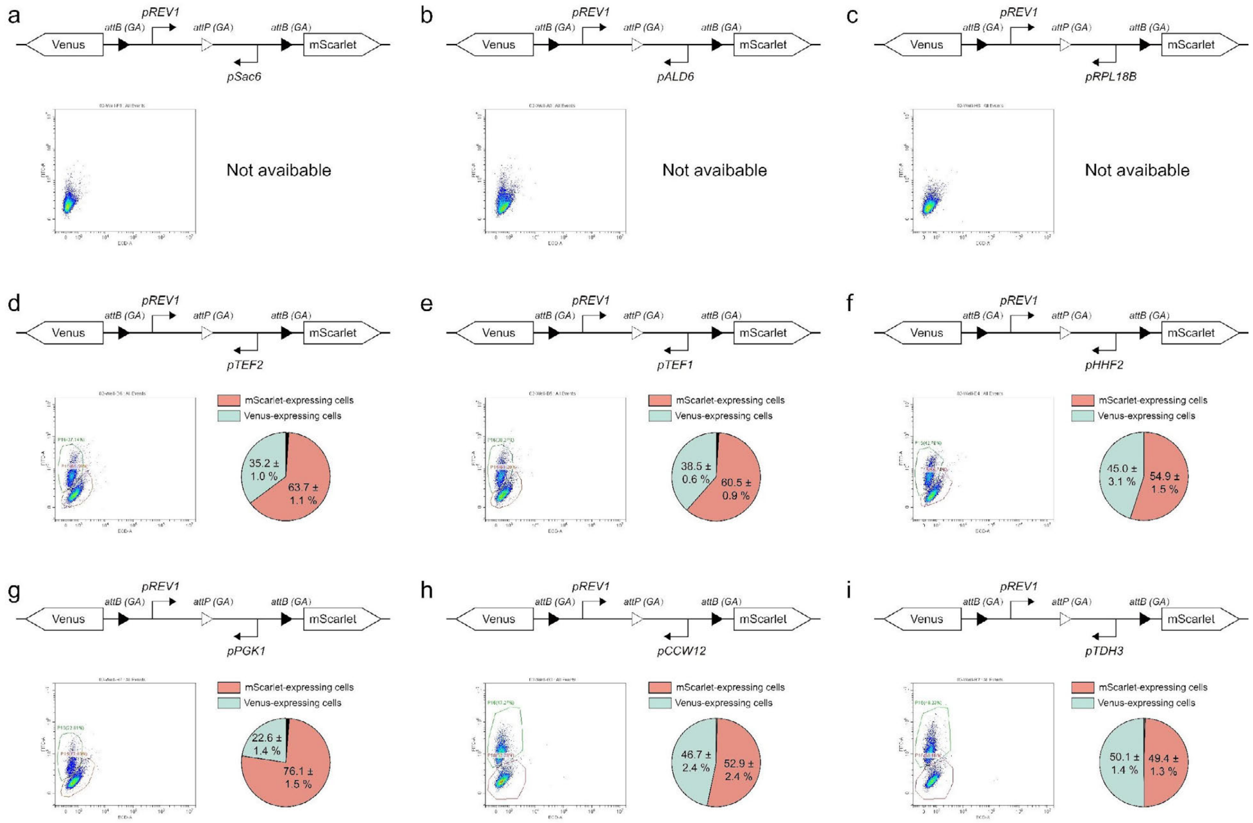

**Fig. S13 | Circuit design with a fixed *pREV1* promoter on the left arm and varying right arms.** Each panel shows the gene circuit design, flow cytometry results, and data on progeny cell distributions (data are mean ± s.d. of *n* = *8* replicates).

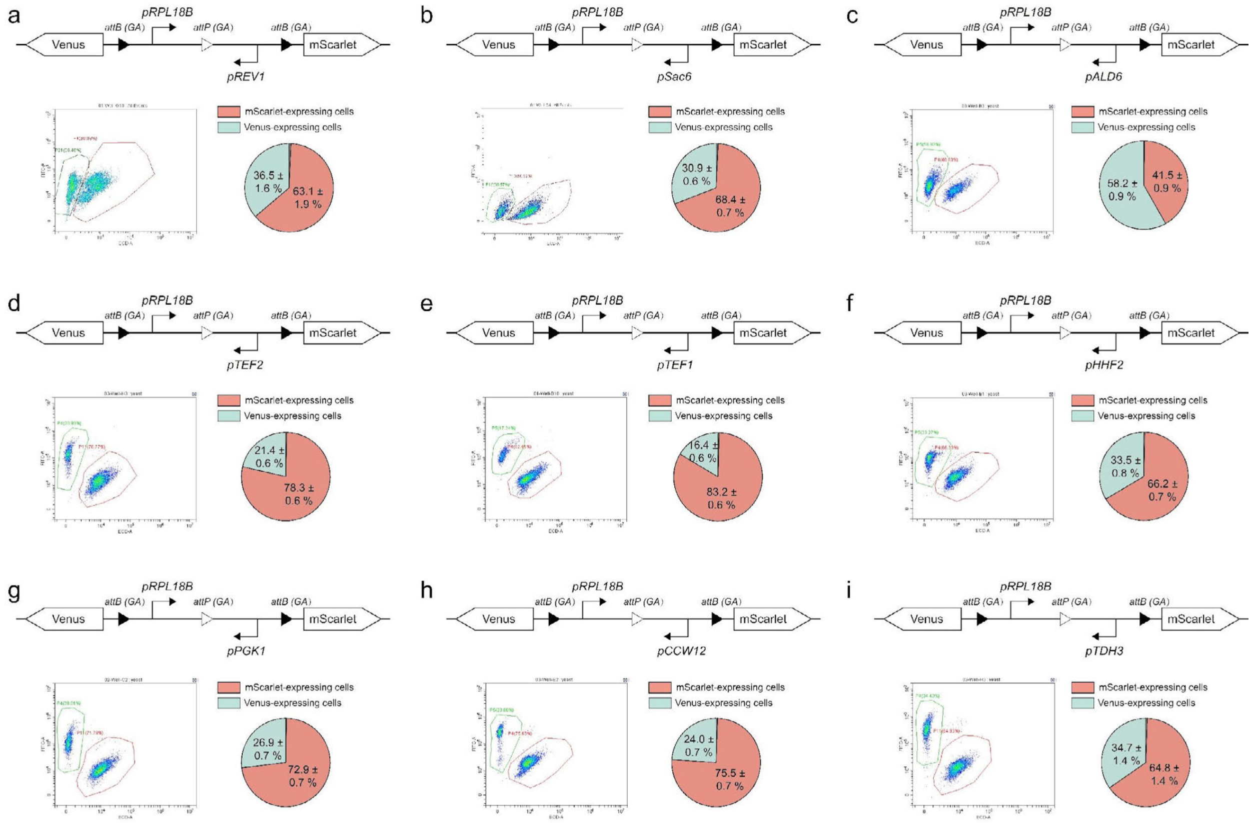

**Fig. S14 | Circuit design with a fixed *pRPL18B* promoter on the left arm and varying right arms.** Each panel shows the gene circuit design, flow cytometry results, and data on progeny cell distributions (data are mean ± s.d. of $n = 8$ replicates).

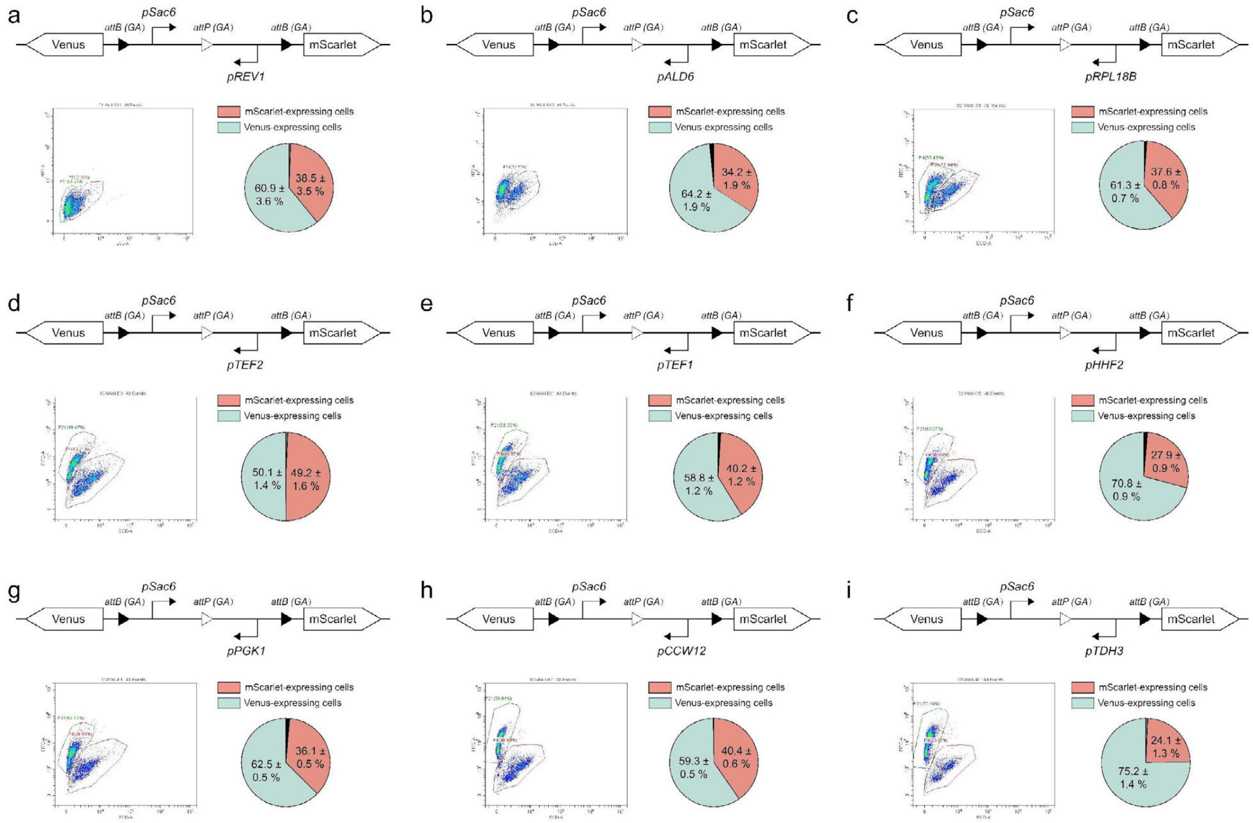

**Fig. S15 | Circuit design with a fixed *pSac6* promoter on the left arm and varying right arms.** Each panel shows the gene circuit design, flow cytometry results, and data on progeny cell distributions (data are mean ± s.d. of $n = 8$ replicates).

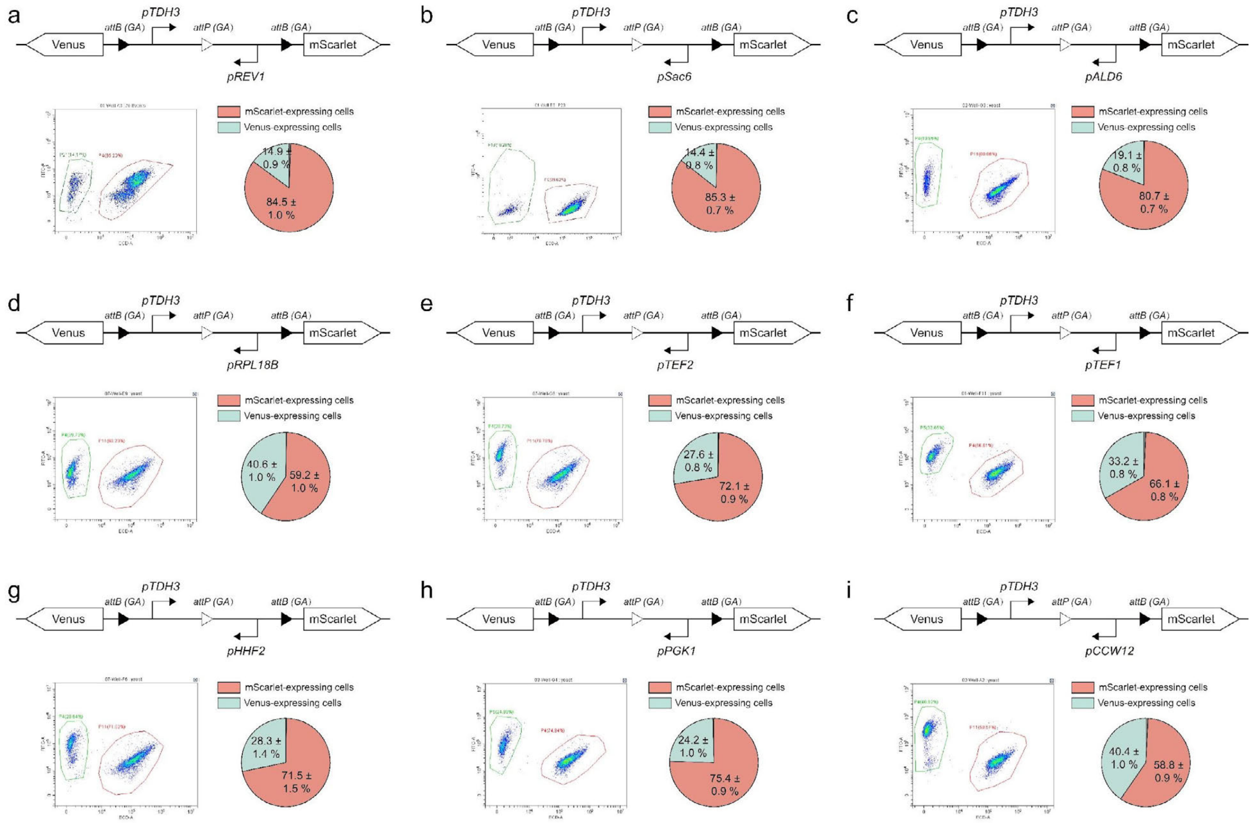

**Fig. S16 | Circuit design with a fixed *pTDH3* promoter on the left arm and varying right arms.** Each panel shows the gene circuit design, flow cytometry results, and data on progeny cell distributions (data are mean ± s.d. of $n = 8$ replicates).

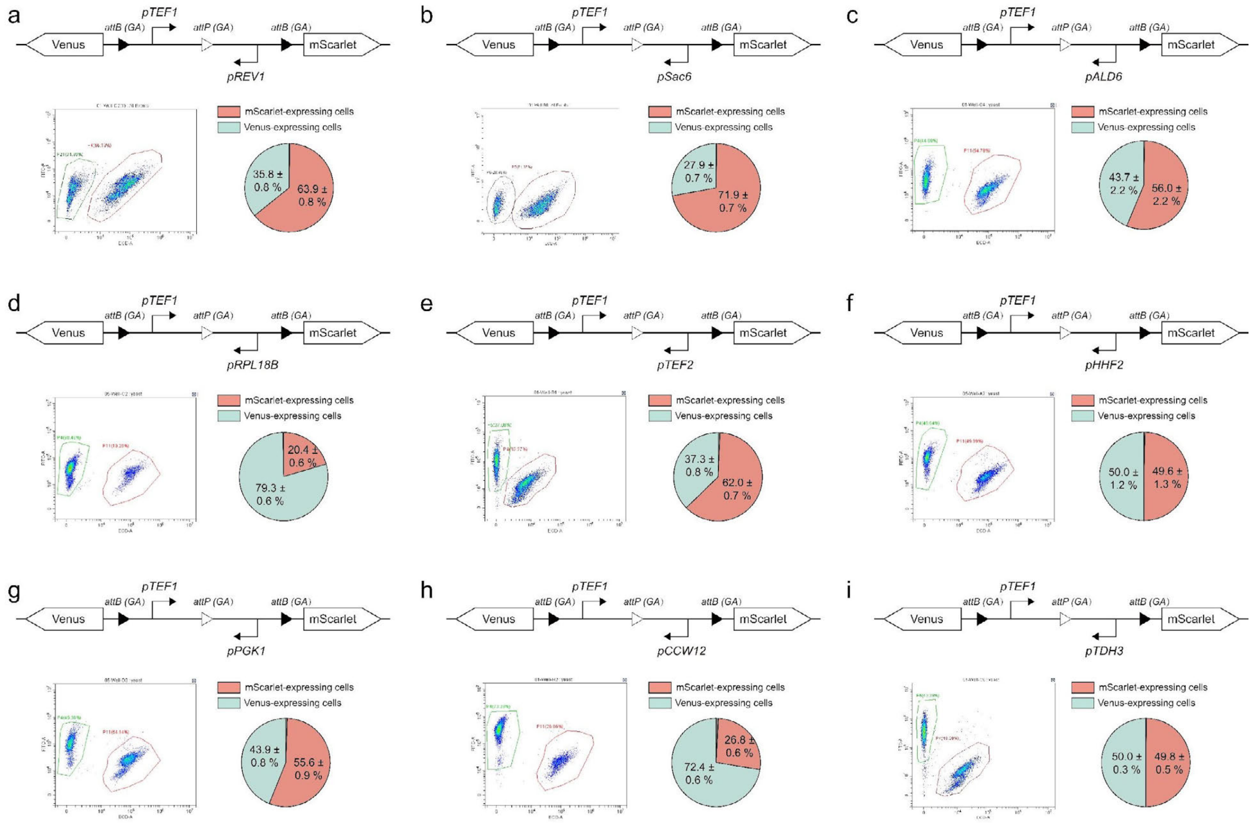

**Fig. S17 | Circuit design with a fixed *pTEF1* promoter on the left arm and varying right arms.** Each panel shows the gene circuit design, flow cytometry results, and data on progeny cell distributions (data are mean ± s.d. of $n$ = 8 replicates).

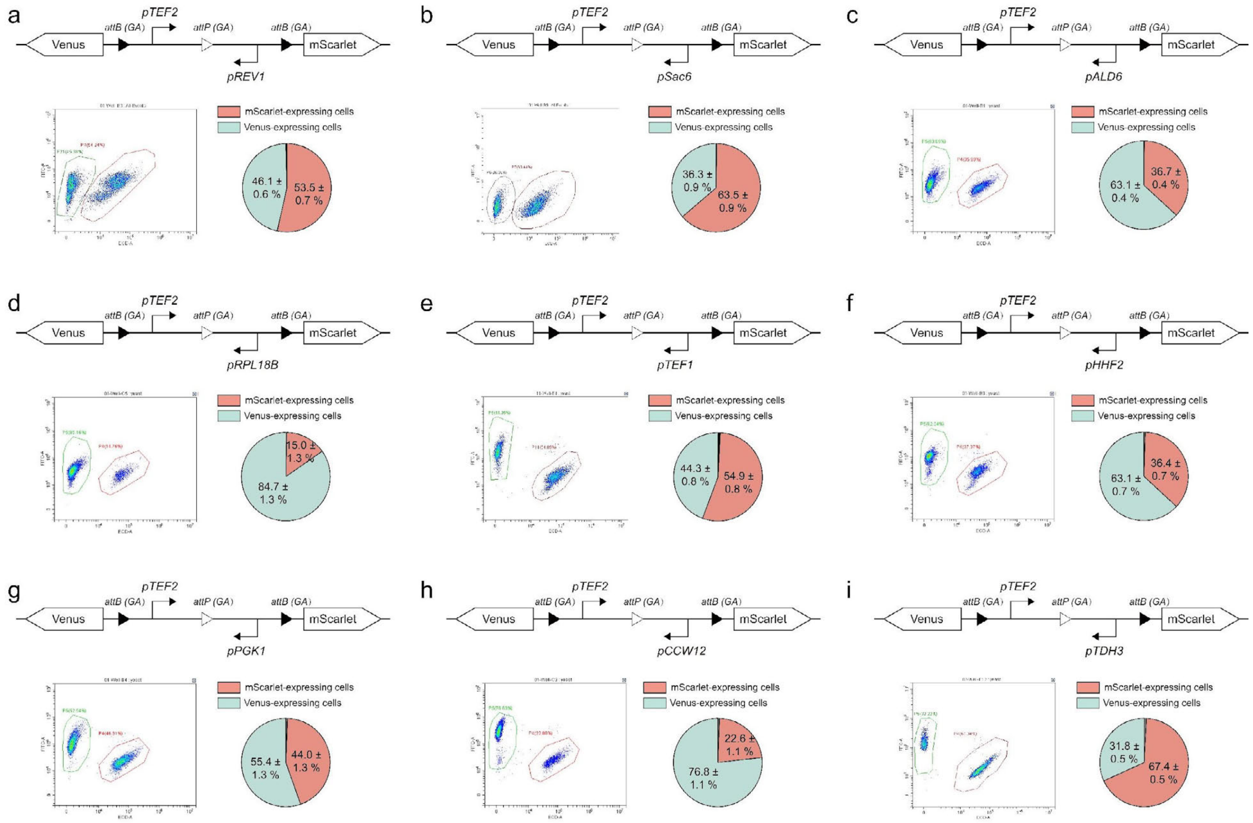

**Fig. S18 | Circuit design with a fixed *pTEF2* promoter on the left arm and varying right arms.** Each panel shows the gene circuit design, flow cytometry results, and data on progeny cell distributions (data are mean ± s.d. of $n = 8$ replicates).

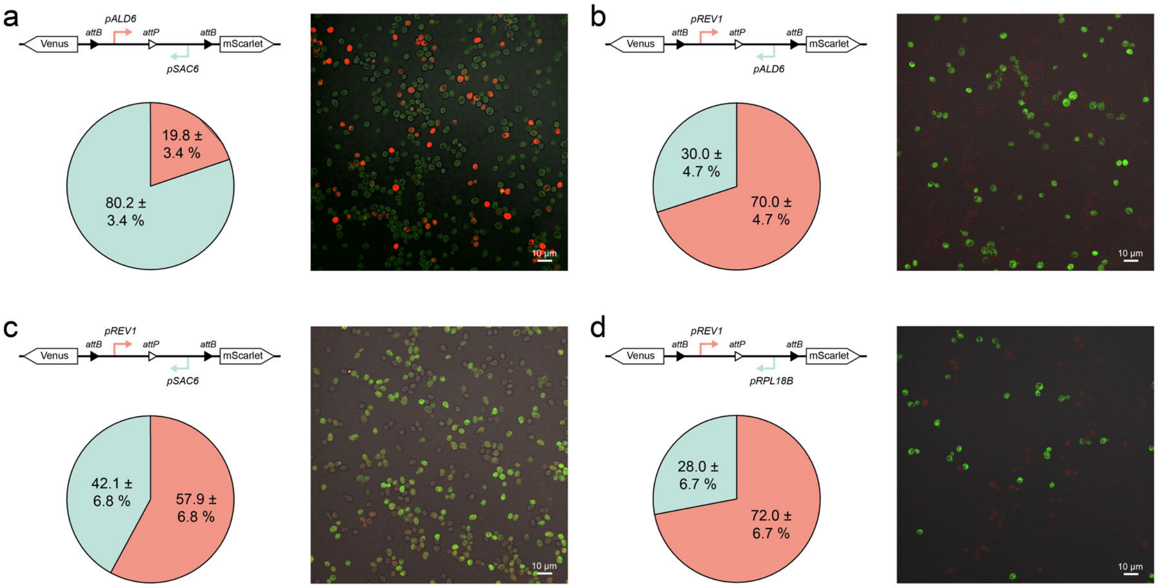

**Fig. S19 | Fluorescence microscopy-based quantification of cell differentiation in branching devices using weak promoters. a-d,** Schematic representations of four recombinase-based differentiation circuits, each containing two relatively weak constitutive promoters flanking the Bxb1 recombination sites. Each construct was transformed into yeast and induced for recombination. The resulting cell populations were imaged by fluorescence microscopy to capture expression of Venus (green) and mScarlet (red). Data were mean ± s.d. of $n = 6$ replicates.

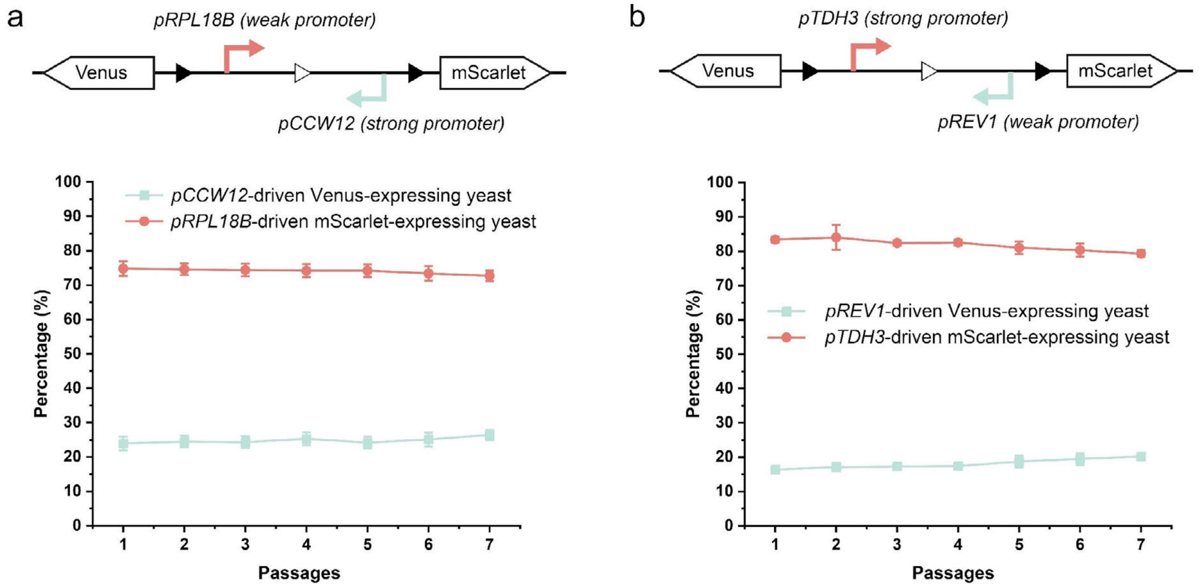

**Fig. S20 | Fluorescent protein expression from strong or weak promoters does not significantly affect the composition of yeast consortia over seven passages.** Results represent mean ± s.d. from $n$ = 4 independent experiments.

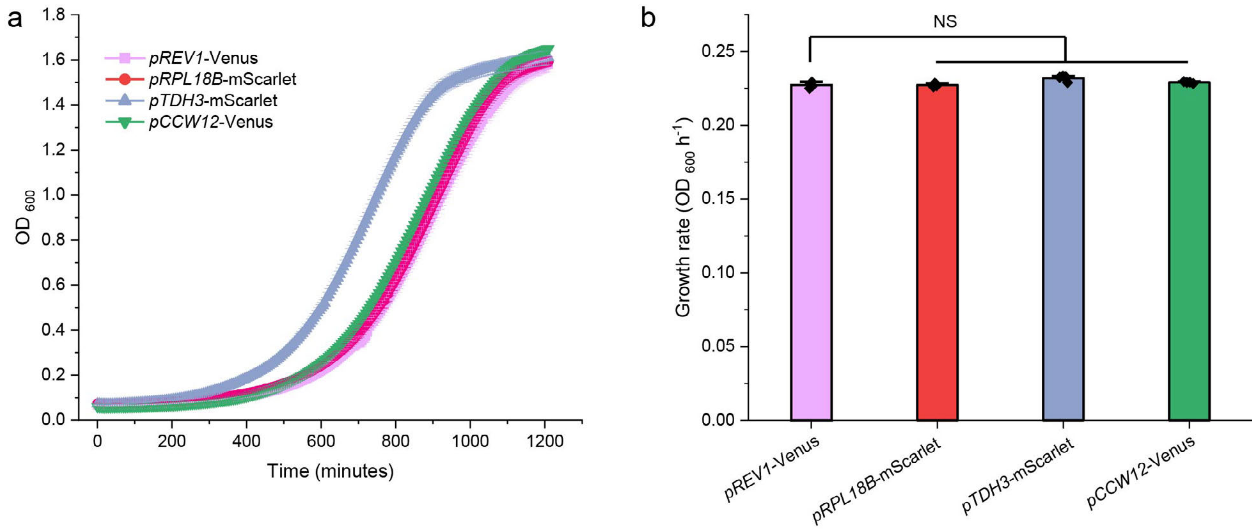

**Fig. S21 | Growth rate comparison between yeast expressing fluorescent proteins from weak and strong promoters. a**. Representative growth curves of 4 progeny cells generated from the circuit shown in **Figure S20**. **b**. All four progeny cells exhibited similar growth rates of approximately 0.23 $OD_{600}$ $h^{-1}$. Results represent the mean ± s.d. from $n$ = 4 independent experiments.

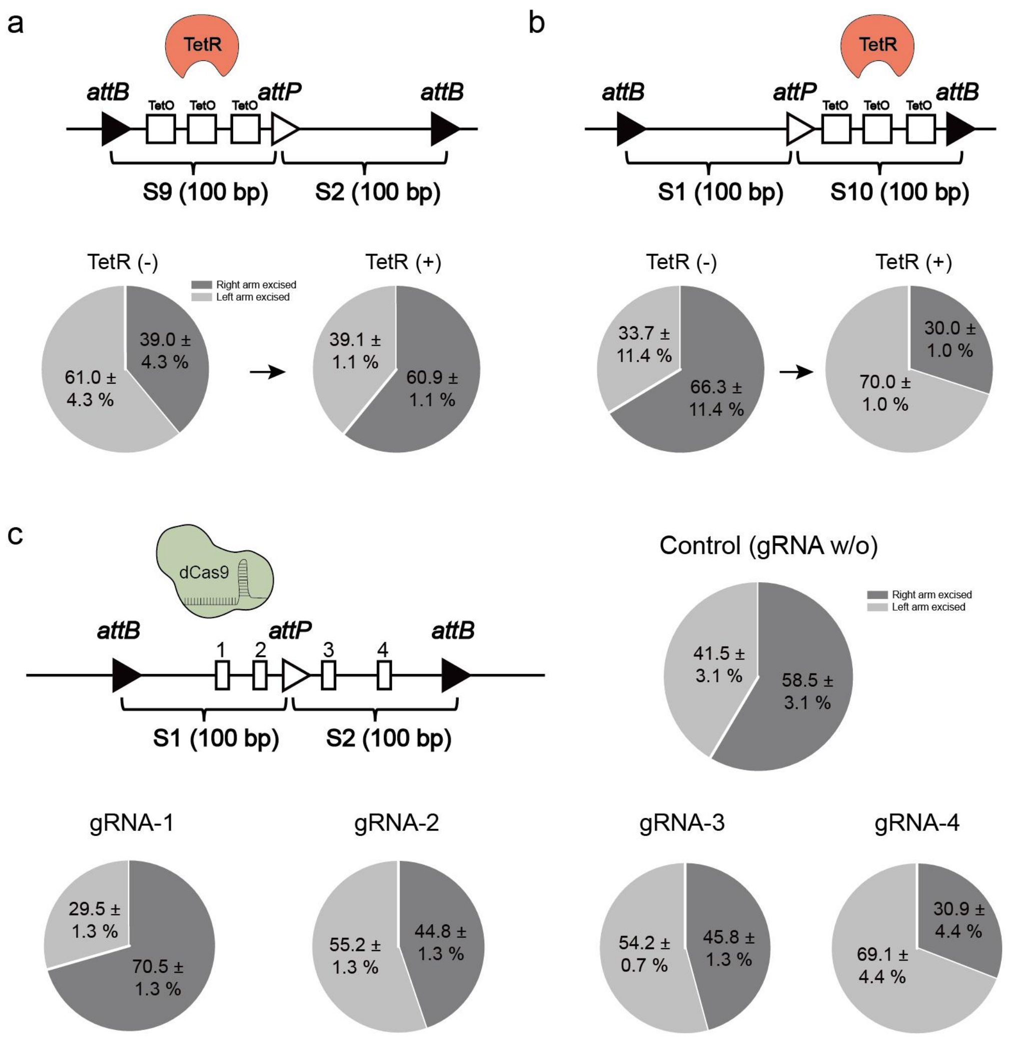

**Fig. S22 | Effects of steric hindrance on Bxb1 recombinase activity assessed using NGS.** **a**. Integration of three TetO operator sequences into the left arm of the branching device resulted in a Bxb1 excision probability of 61.0 ± 4.3% without TetR repressor. When TetR was present, binding TetO sequences reduced the excision probability to 39.1 ± 1.1% (data are mean ± s.d. of $n = 3$ replicates). **b.** The inclusion of three TetO sites between *attP* and *attB* also affected recombinase activity (data are mean ± s.d. of $n = 3$ replicates). **c.** The impact of dCas9 binding on Bxb1 activity depended on the positioning of the gRNA guiding dCas9 binding (data are mean ± s.d. of $n = 3$ replicates). The sequences of S1-S10 are listed in the **Table. S4**.

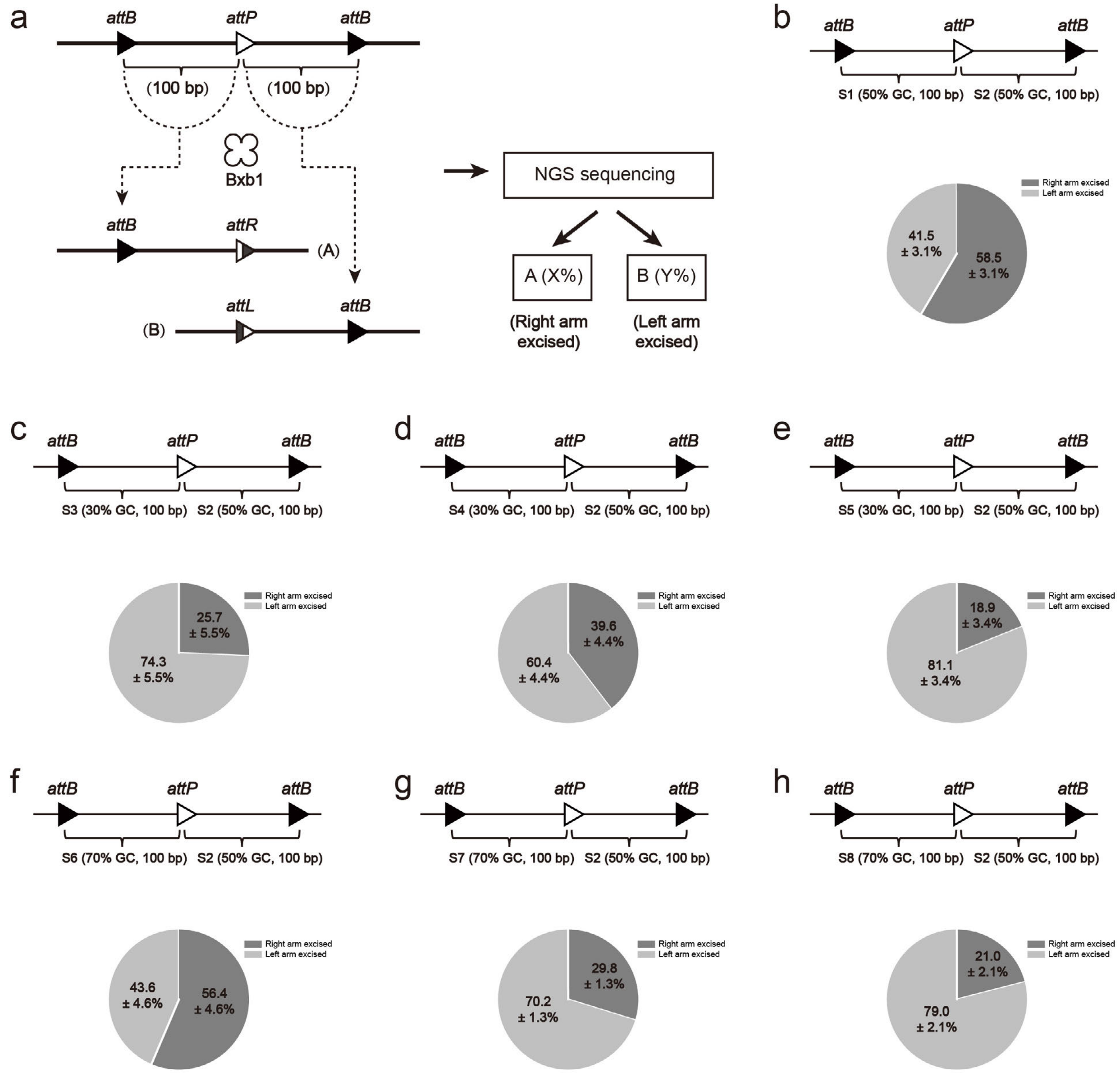

**Fig. S23 | Characterization of branching circuits via NGS**. **a.** Schematic of circuit design featuring two 100-bp random DNA sequences between *attB-attP* and *attP-attB* sites. NGS was used to analyze progeny distributions following cellular differentiation. **b-h.** Eight 100-bp sequences (S1 to S8) with varying GC content were incorporated into the circuit design: S1 and S2 (50% GC), S3-S5 (30% GC), and S6-S8 (70% GC). GC content influenced Bxb1 catalytic preferences, resulting in varied progeny distributions (data are mean ± s.d. of $n = 3$ replicates). This underscores the critical role of DNA sequences in modulating Bxb1 activity. The sequences of S1-S10 are listed in the **Table. S4**.

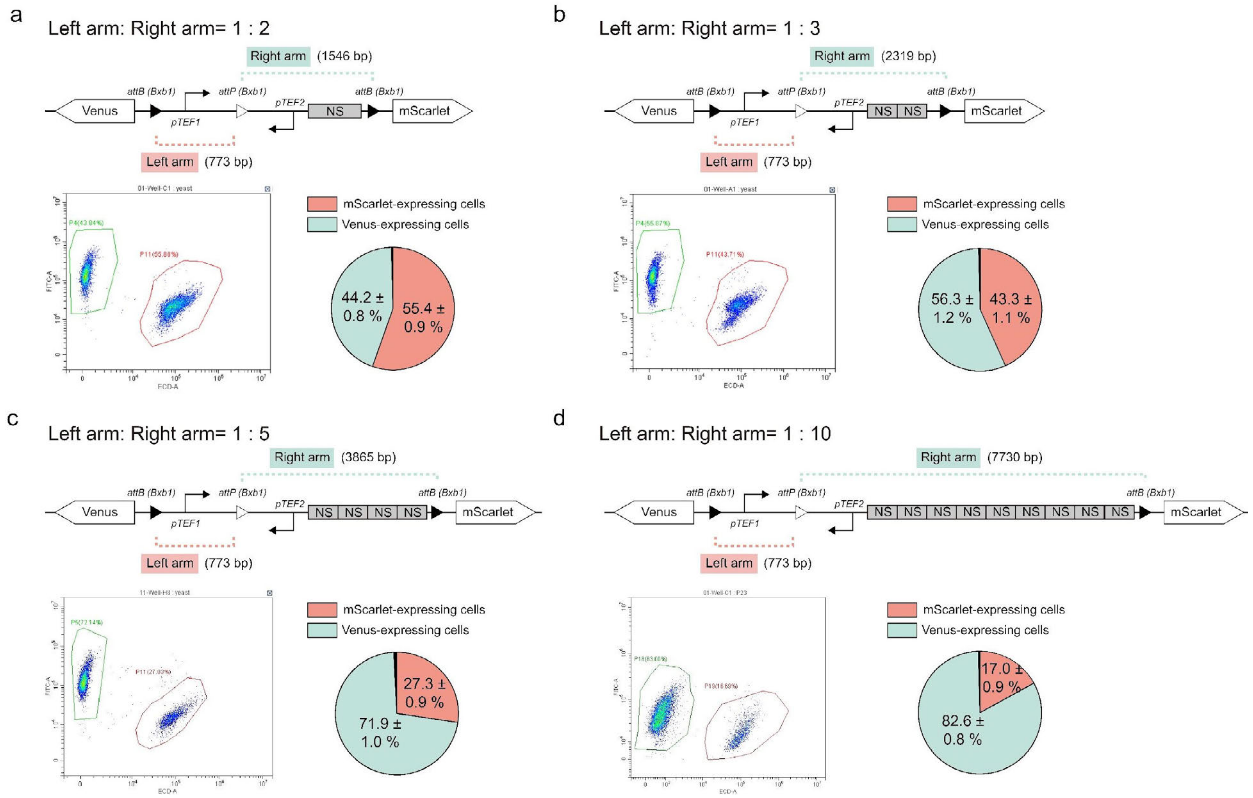

**Fig. S24 | Quantitative characterization of progeny differentiation as DNA length ratios between the left and right arms of engineered branching devices progressively shift from 1:2 to 1:10 (a-d).** Data are mean ± s.d. of *n* = 8 replicates.

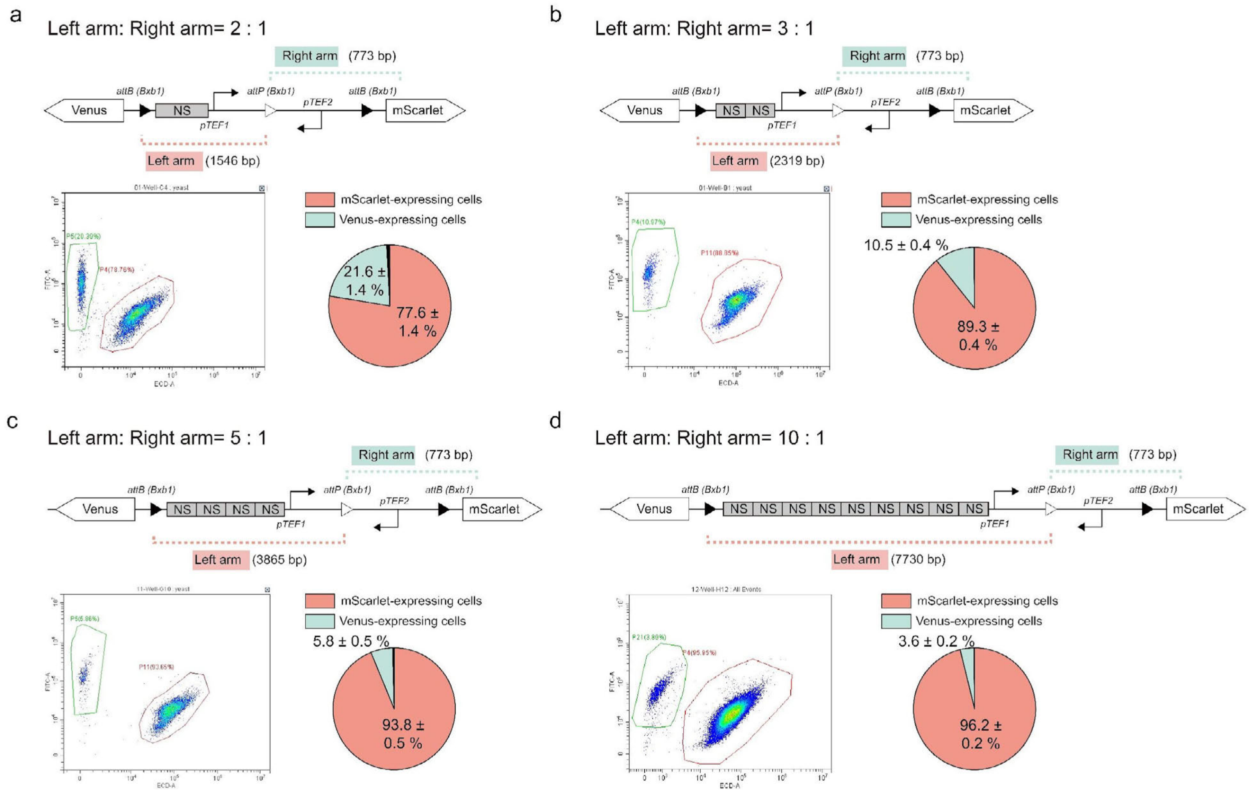

**Fig. S25 | Quantitative characterization of progeny differentiation as DNA length ratios between the left and right arms of engineered branching devices progressively shift from 2:1 to 10:1 (a-d).** Data are mean ± s.d. of $n = 8$ replicates. The sequences of NS are listed in the **Table. S4**.

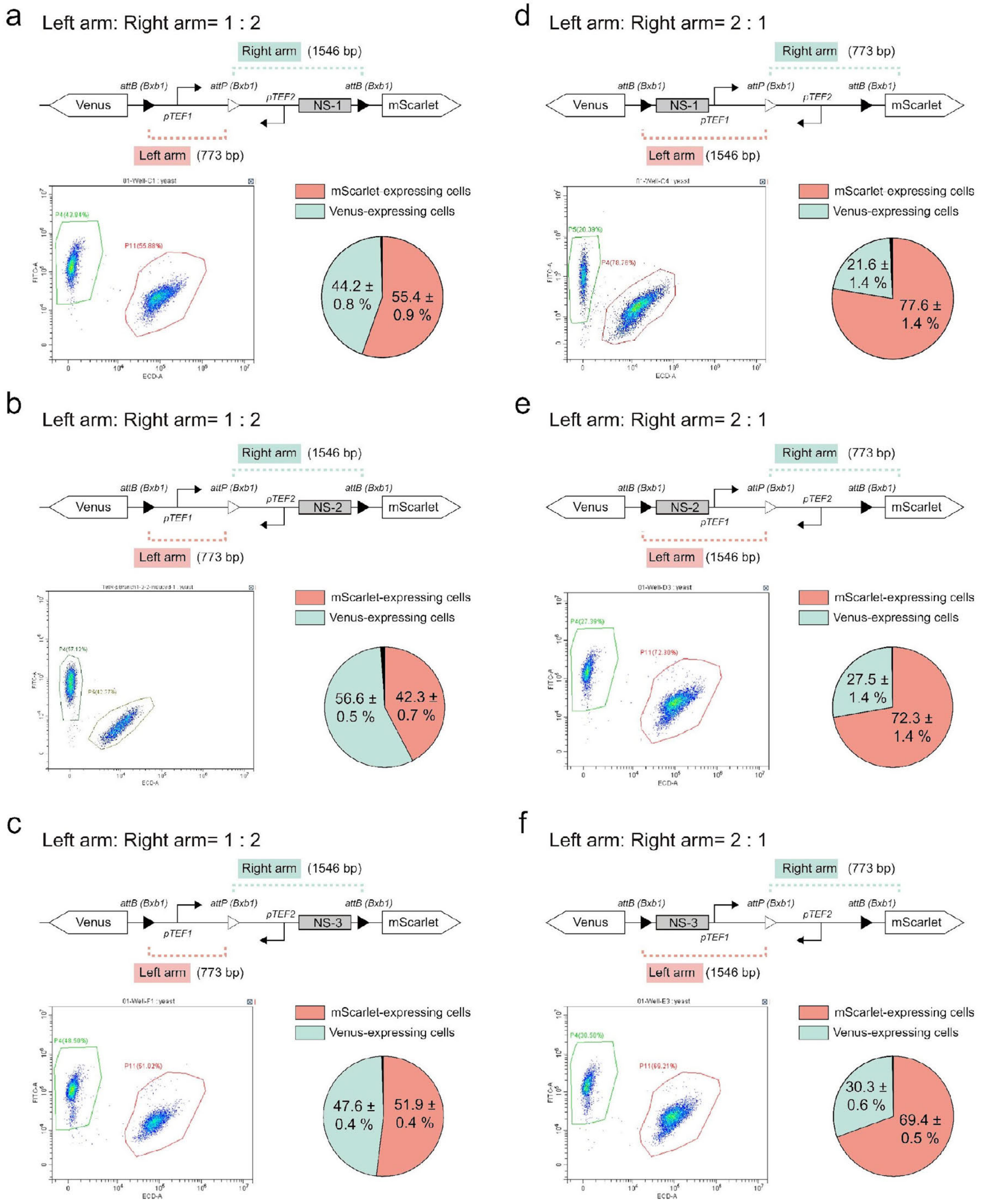

**Fig. S26 | Quantitative analysis of progeny differentiation in yeast with varying DNA length ratios and nonsense sequences (NS). a-c**. Quantitative analysis of yeast progeny differentiation with a 1:2 DNA length ratio between left and right arms, incorporating different 773-bp nonsense sequences (NS-1, NS-2, NS-3). **d-f.** Quantitative analysis of yeast progeny differentiation with a 2:1 DNA length ratio between the left and right arms. The sequences of NS are listed in the **Table. S4**. Data are mean ± s.d. of $n$ = 8 replicates.

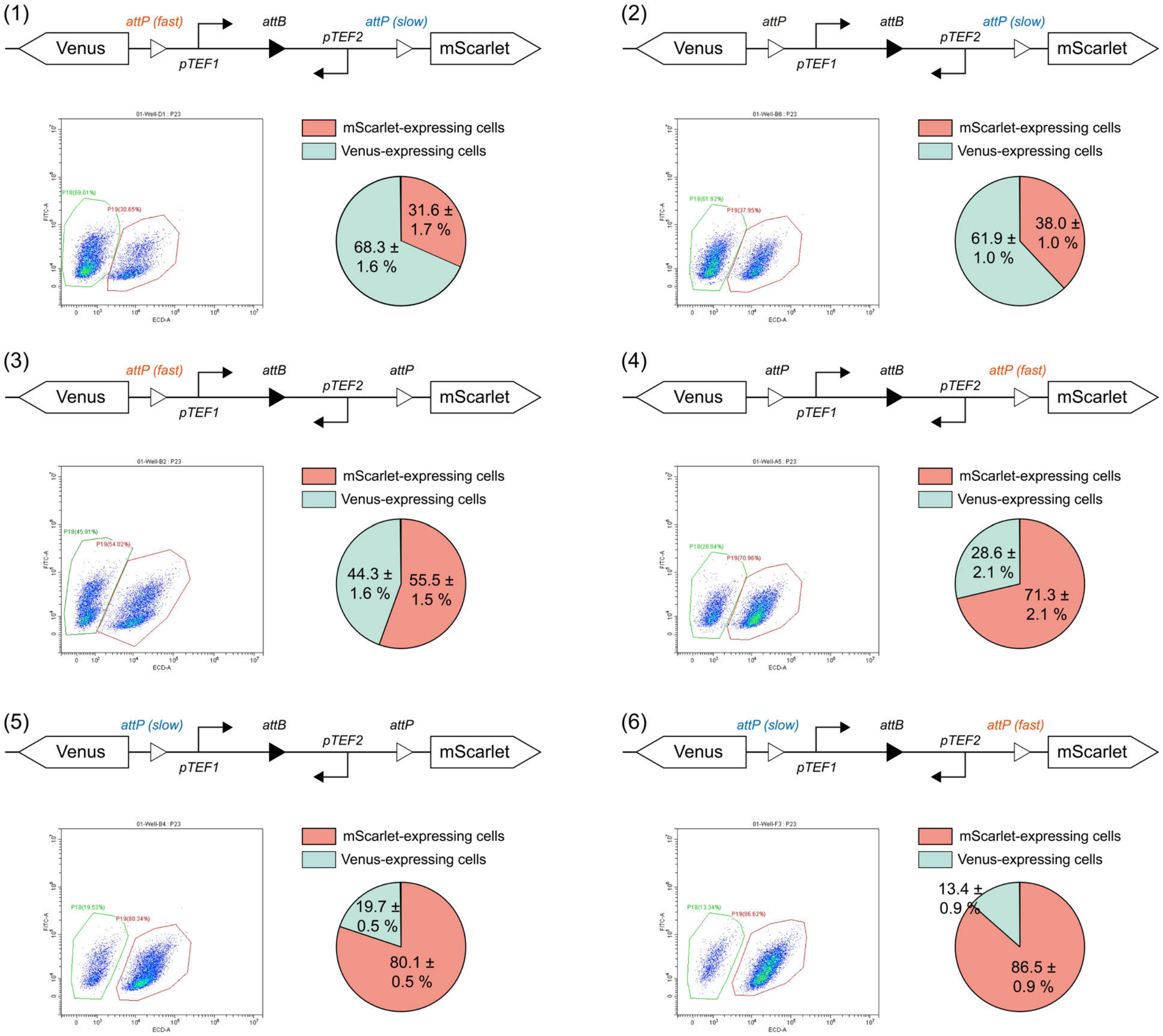

**Fig. S27 | Tuning cellular differentiation using branching circuits with *attP* variants exhibiting predictable catalytic rates.** Representative genetic circuits, flow cytometry data, and progeny distribution statistics. The sequences of *attP* variants are listed in the **Table. S4**. Data are mean ± s.d. of $n = 8$ replicates.

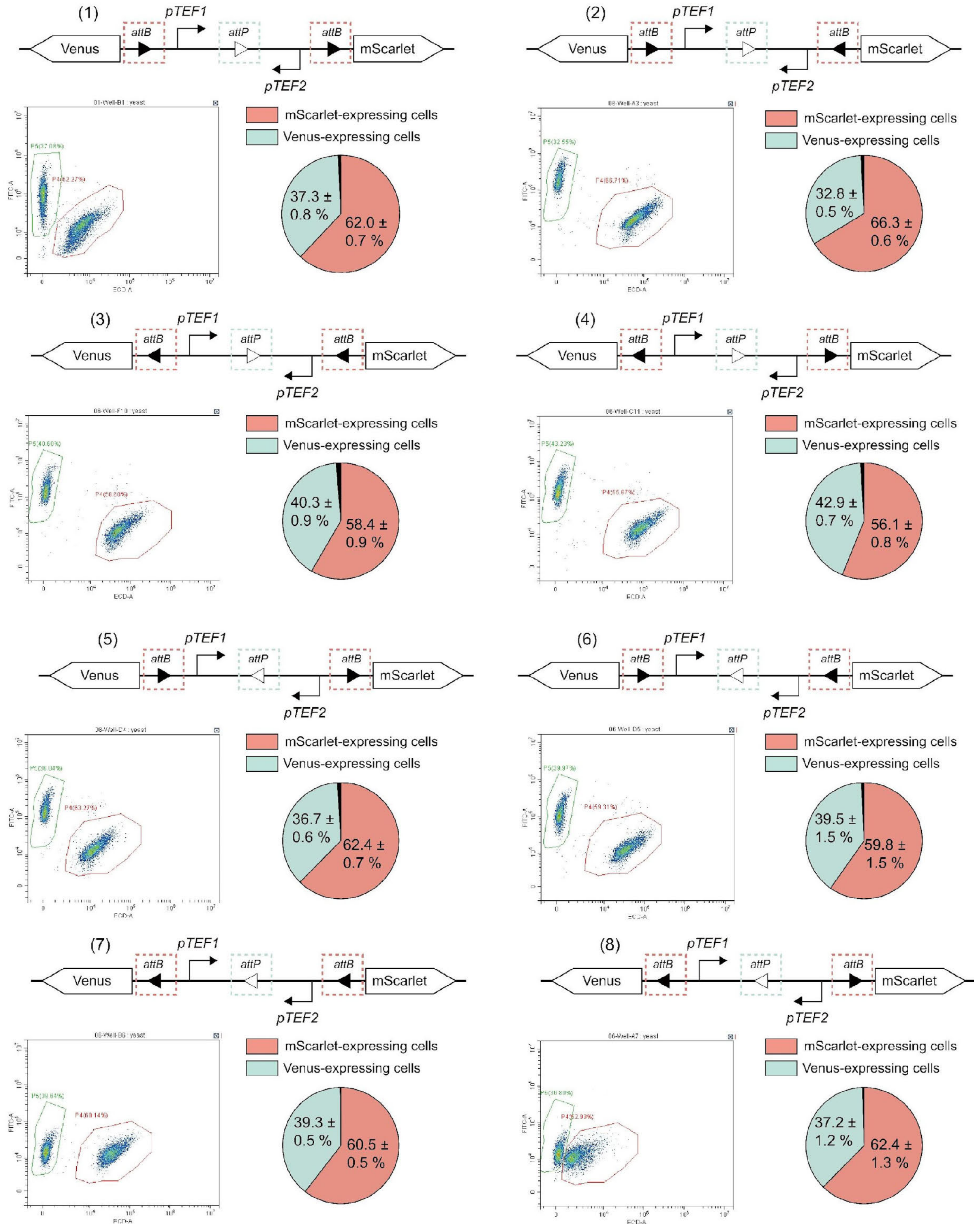

**Fig. S28 | Effect of *att* site orientation on cellular differentiation.** Representative circuit designs, flow cytometry data, and progeny distribution statistics. Data are mean ± s.d. of $n = 8$ replicates.

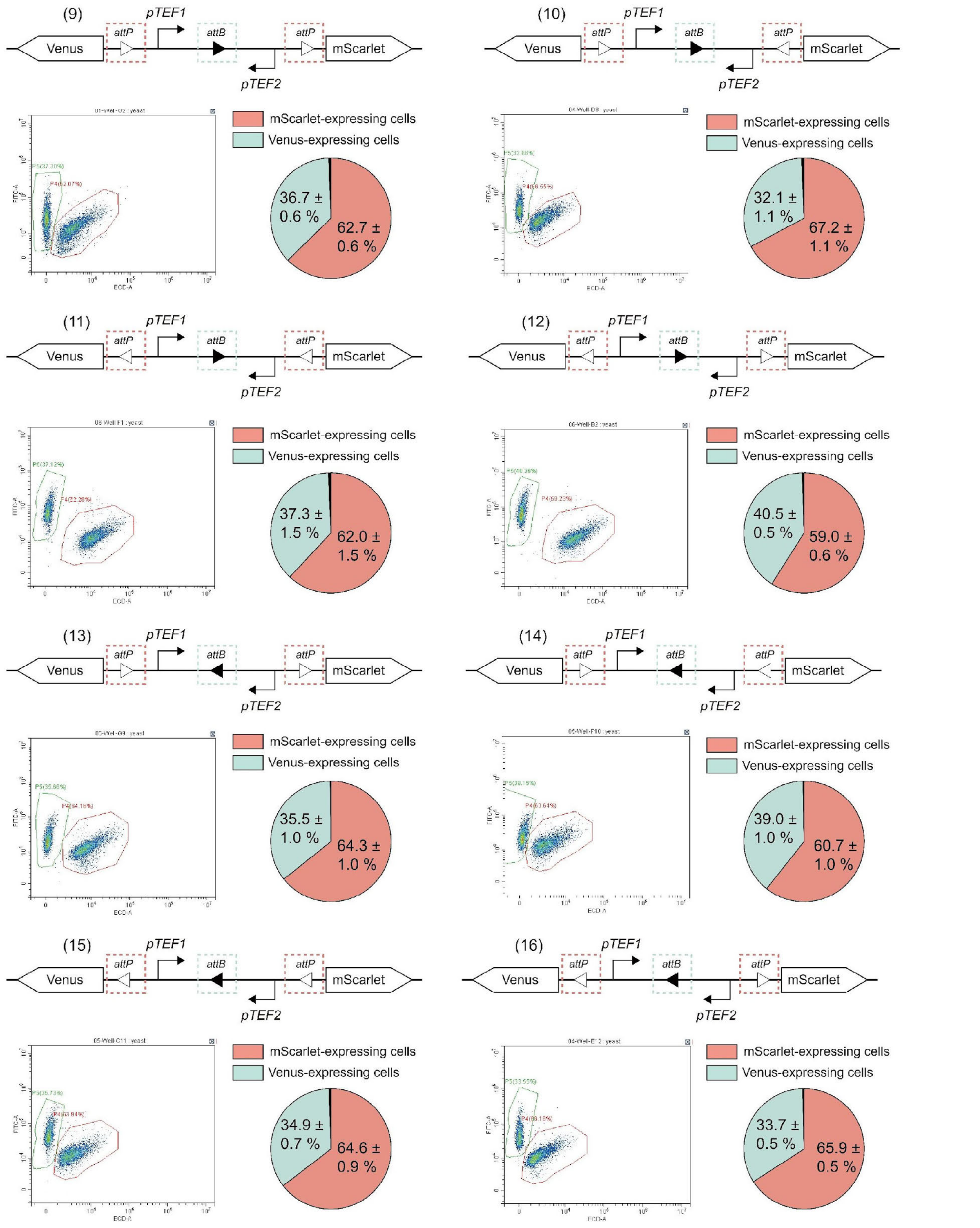

**Fig. S29 | Impact of *att* site orientation on cellular differentiation. Circuit design, flow cytometry data, and progeny distribution statistics.** Data are mean ± s.d. of $n = 8$ replicates.

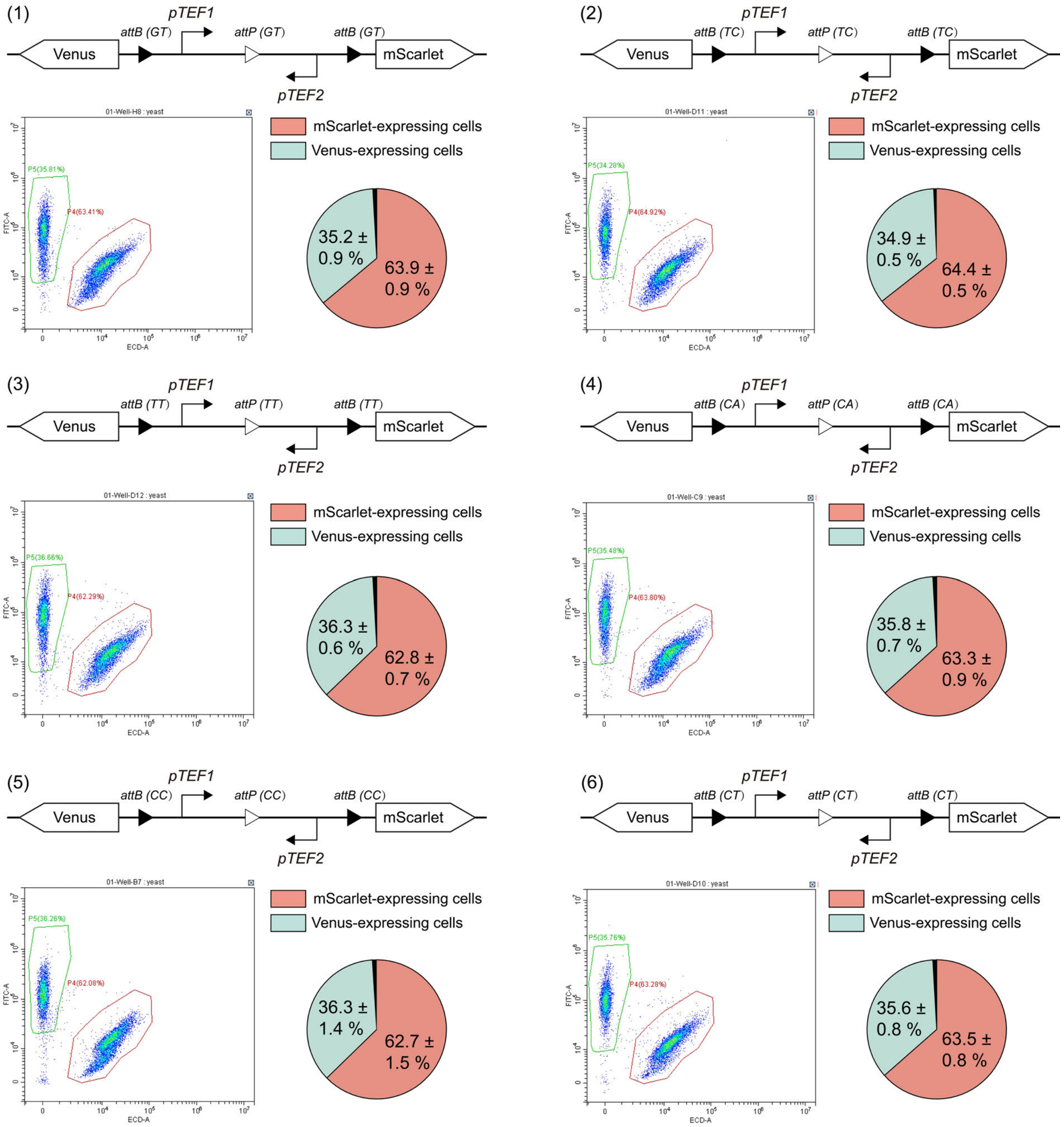

**Fig. S30 | Impact of orthogonal *att* variants on cellular differentiation.** Circuit designs, flow cytometry, and progeny distribution statistics. Data are mean ± s.d. of $n = 8$ replicates.

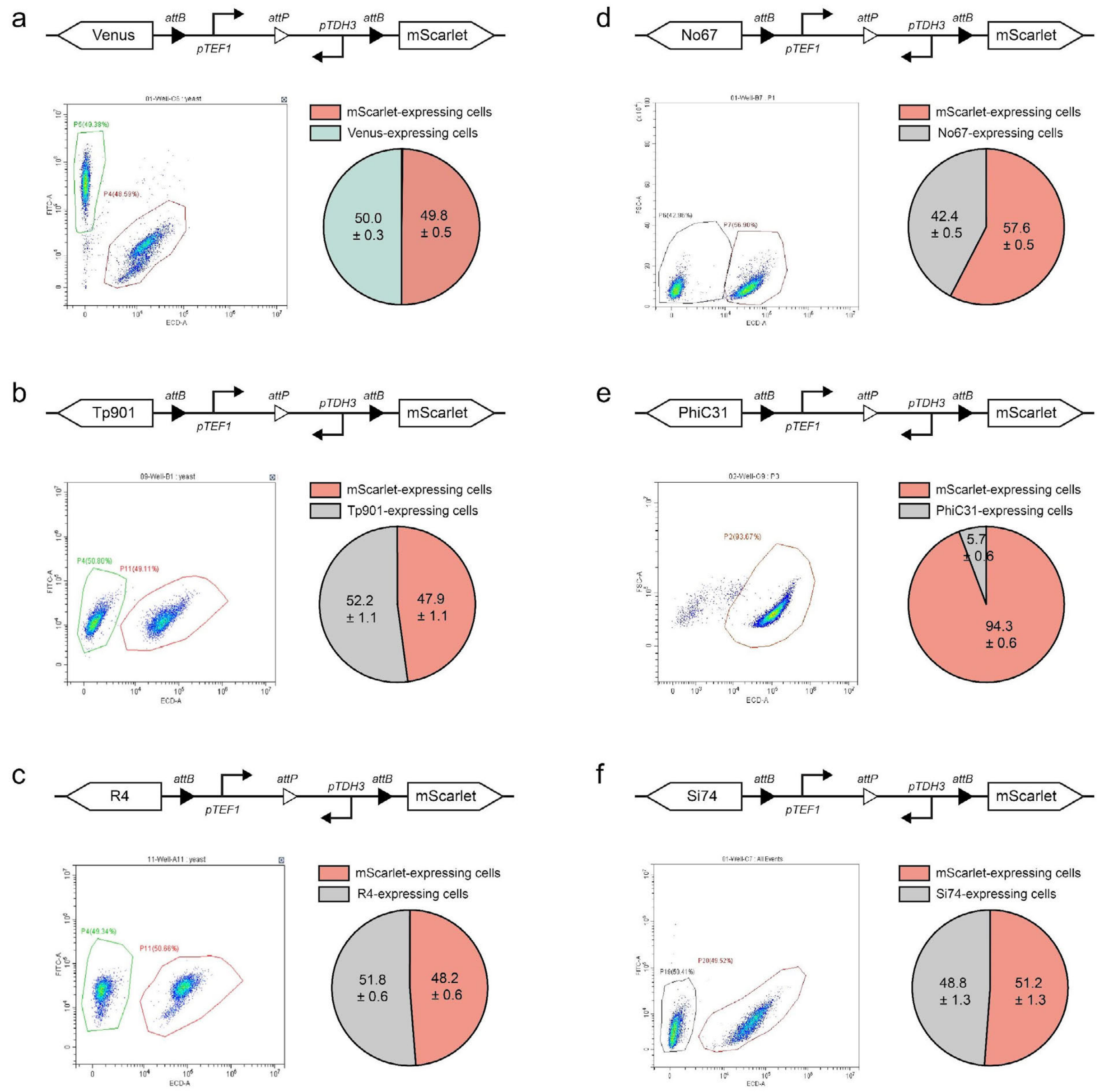

**Fig. S31 | Impact of varying outputs on progeny distribution in branching device designs**. **a-f**. Substitution of the left-arm reporter protein Venus (**a**) with alternative recombinases—Tp901 (**b**), No67 (**c**), R4 (**d**), PhiC31 (**e**), and Si74 (**f**)—altered progeny ratio distributions. The sequences of the used recombinases are listed in the **Table. S5**. Data are mean ± s.d. of *n* = 4 replicates.

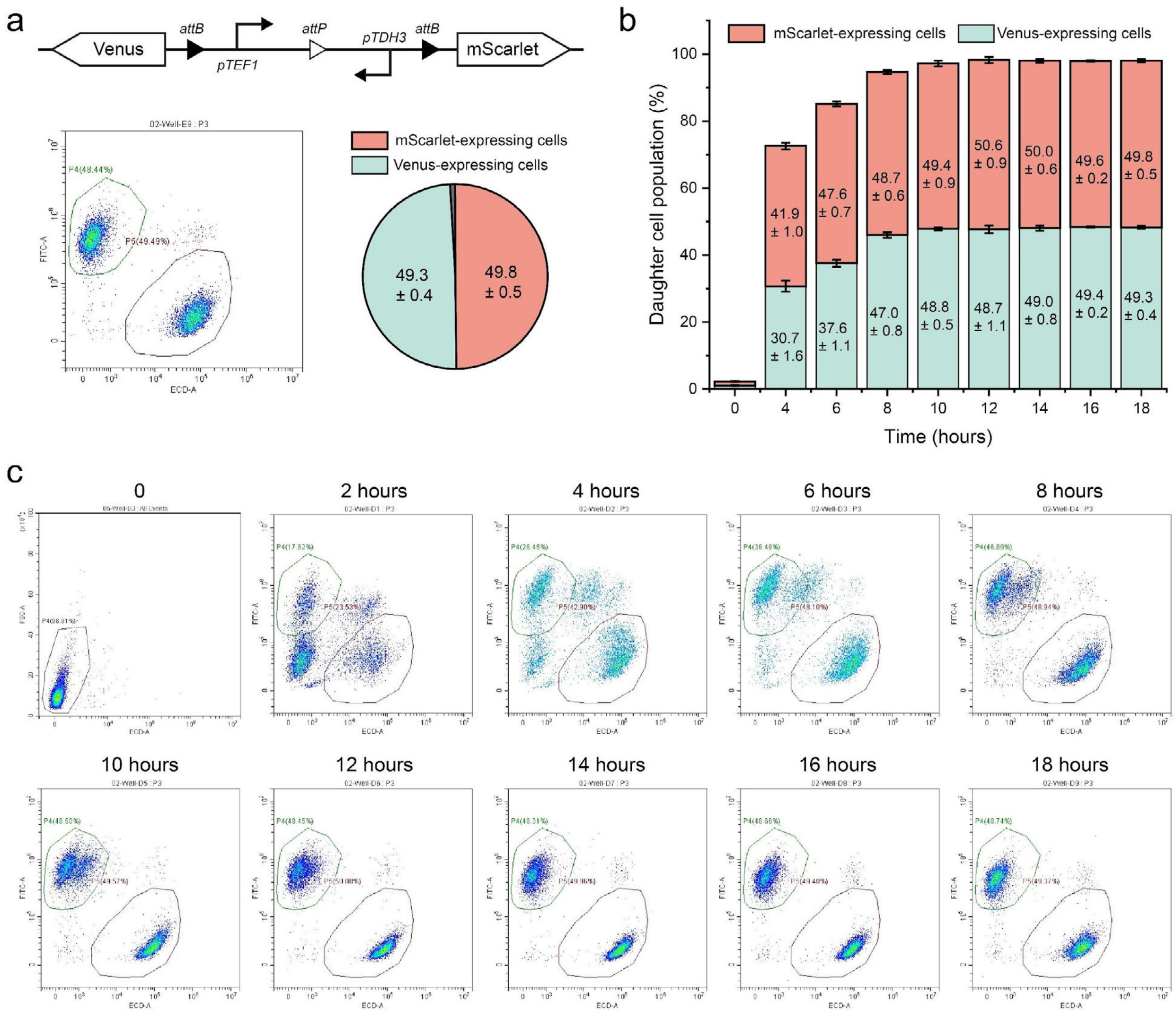

**Fig. S32 | Temporal dynamics of progeny populations during 18 hours of culturing. a.** Representative flow cytometry data and progeny distribution statistics for the genetic circuit following utilization of Venus and mScarlet as output proteins. **b.** Statistical data showing progeny dynamics over time with a final aTc inducer concentration of 1 mM (data are mean ± s.d. of *n* = 4 replicates). **c.** Flow cytometry results were measured at 2-hour intervals from 0 to 18 hours.

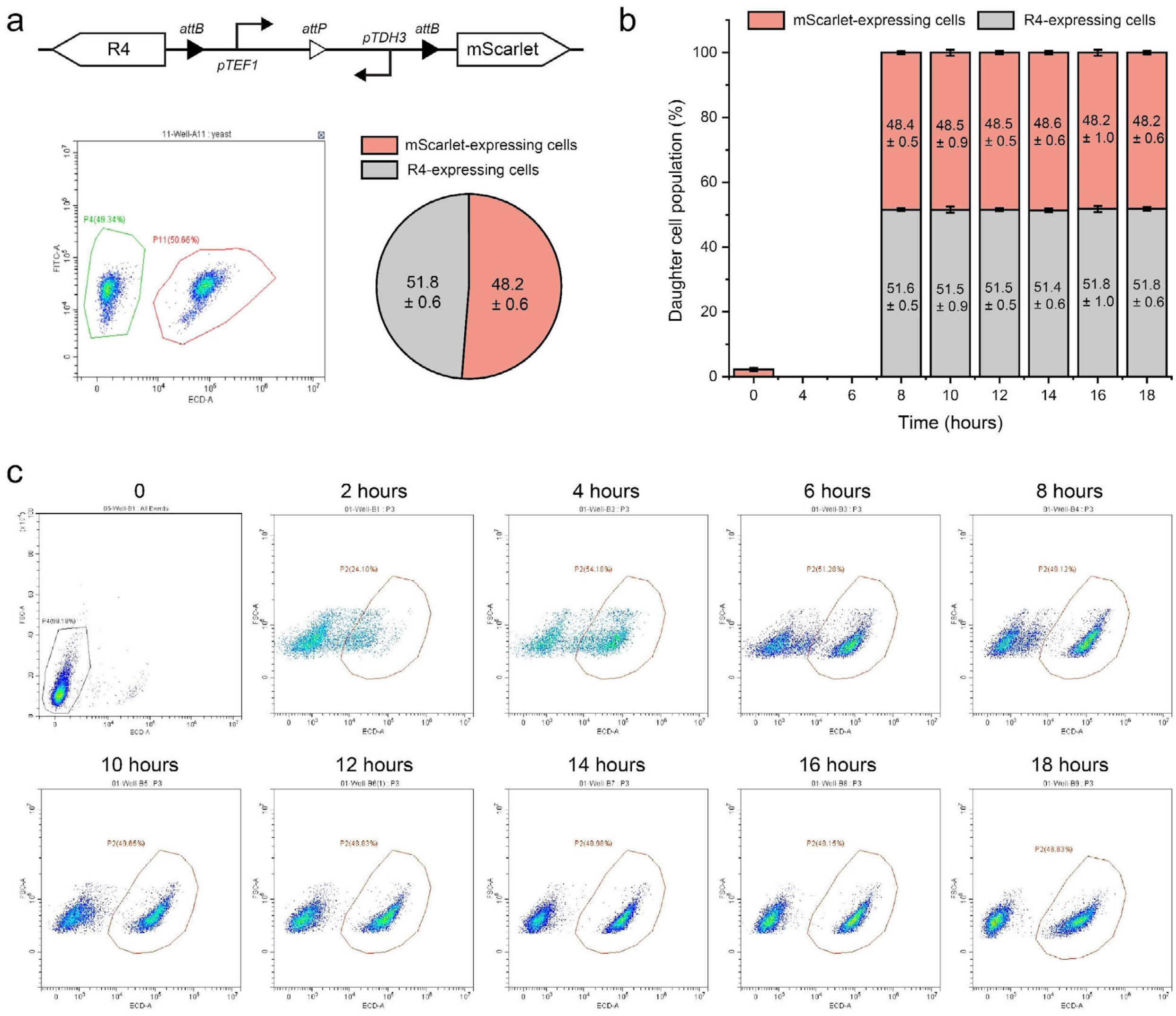

**Fig. S33 | Temporal dynamics of progeny populations during 18 hours of culturing. a.** Representative flow cytometry data and progeny distribution statistics for the genetic circuit using recombinase R4 and mScarlet as output proteins. **b.** Data showing progeny dynamics over time with a final aTc inducer concentration of 1 mM (data are mean ± s.d. of *n* = 4 replicates). **c.** Flow cytometry results were measured at 2-hour intervals from 0 to 18 hours.

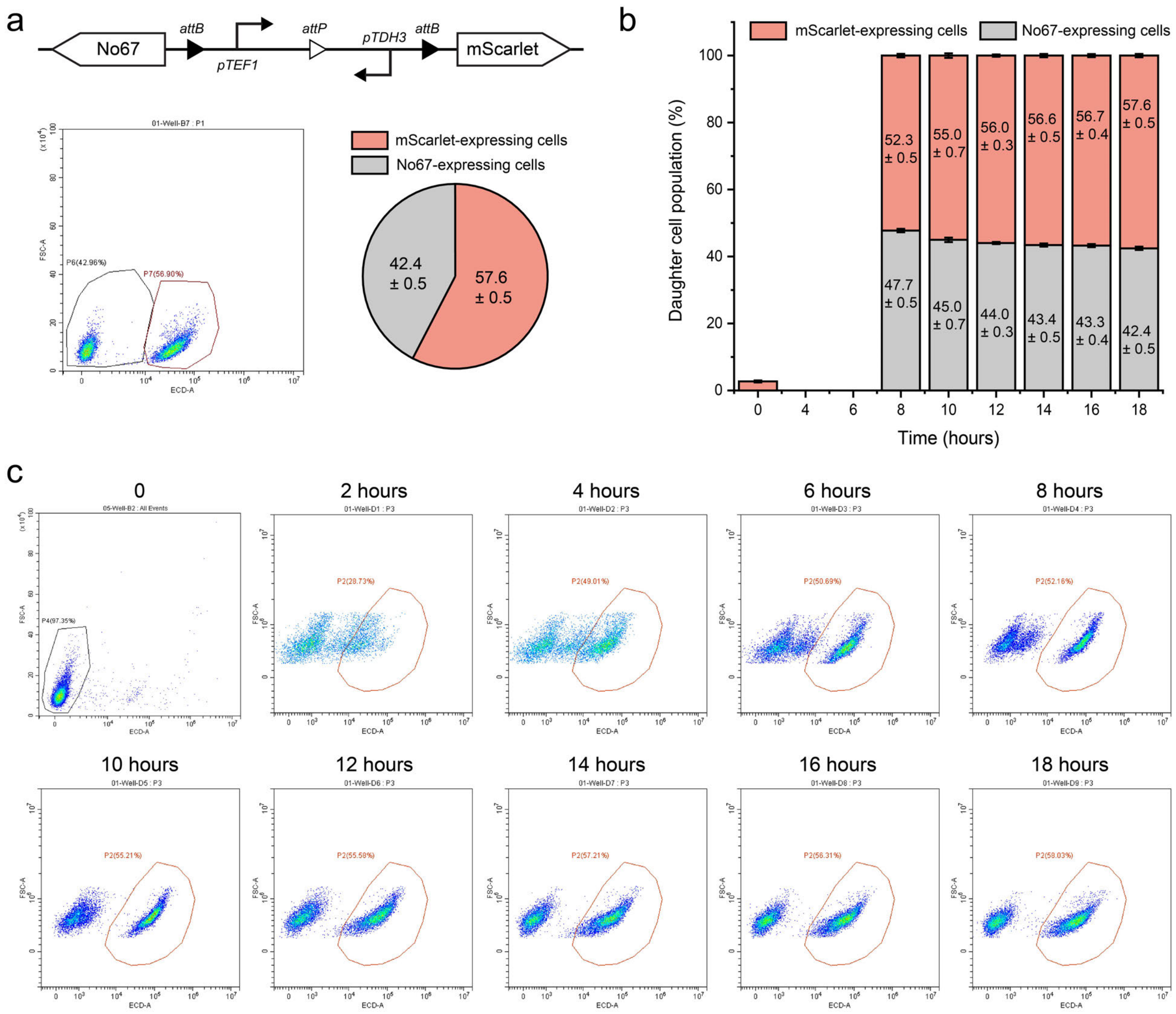

**Fig. S34 | Temporal dynamics of yeast progeny populations during 18 hours of culturing.** **a.** Representative flow cytometry data and progeny distribution statistics for the genetic circuit utilizing recombinase No67 and mScarlet as output proteins. **b.** Data showing progeny dynamics over time with a final aTc inducer concentration of 1 mM (data are mean ± s.d. of n = 4 replicates). **c.** Flow cytometry results were measured at 2-hour intervals from 0 to 18 hours.

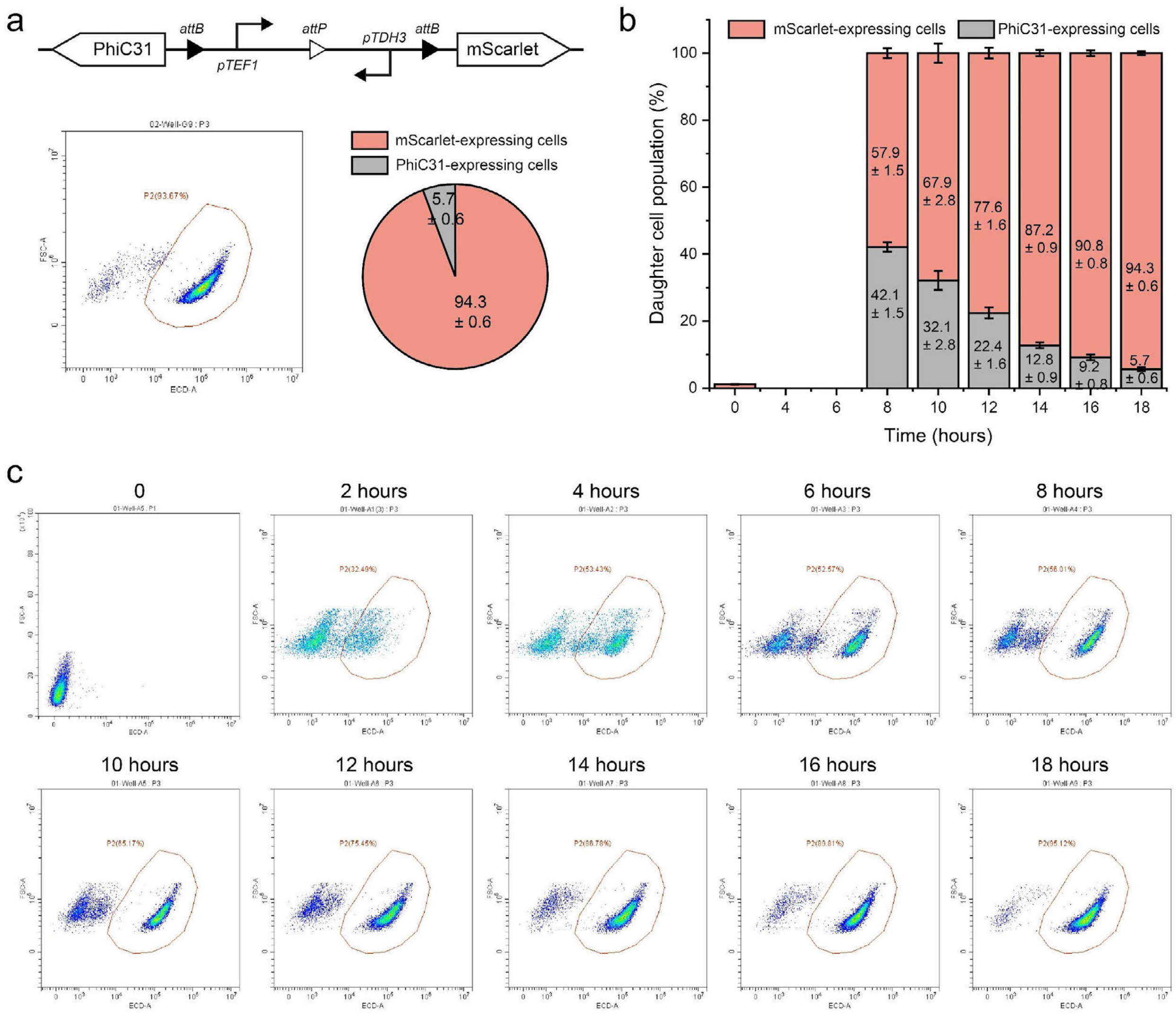

**Fig. S35 | Temporal dynamics of yeast progeny populations during 18 hours of culturing. a.** Representative flow cytometry data and progeny distribution statistics for the genetic circuit utilizing recombinase PhiC31 and mScarlet as output proteins. **b.** Statistical data showing progeny dynamics over time with a final aTc inducer concentration of 1 mM (data are mean ± s.d. of n = 4 replicates). **c.** Flow cytometry results were measured at 2-hour intervals from 0 to 18 hours.

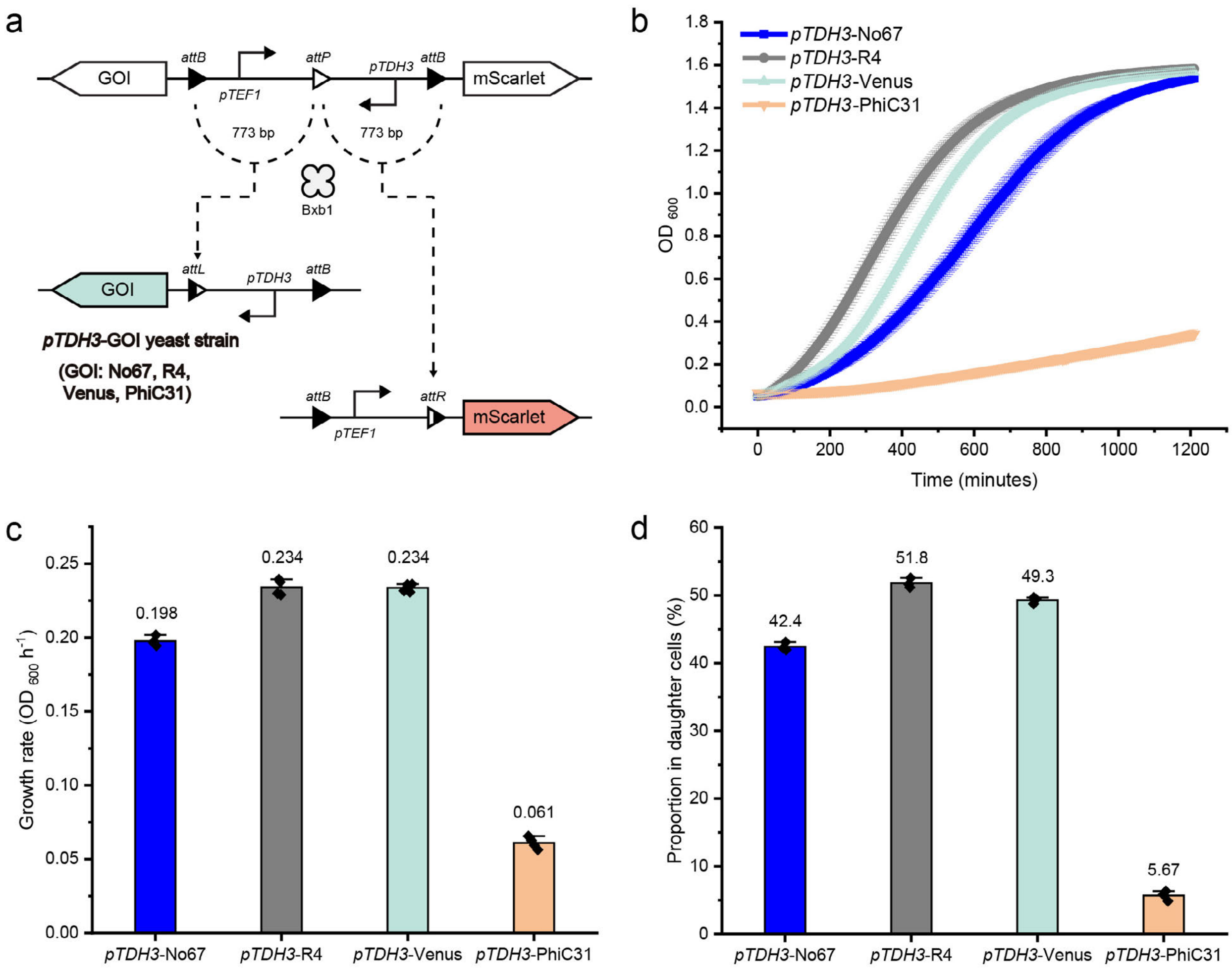

**Fig. S36 | Influence of progeny growth rates on consortia proportions. a.** Genetic circuits used. Founder cells differentiate into two subgroups: one expressing mScarlet via the *pTEF1* promoter and the other expressing a gene of interest (Venus, No67, R4, or PhiC31) via the *pTDH3* promoter. **b.** Growth curves of the four types of progeny yeast, with *pTDH3*-driven expression of Venus, No67, R4, or PhiC31 (data are mean ± s.d. of $n = 4$ replicates). **c.** Average growth rates of the four yeast types (data are mean ± s.d. of n = *4* replicates). **d.** Progeny distribution in yeast consortia generated by the circuit as depicted in **Fig. S36a** (data are mean ± s.d. of $n = 4$ replicates).

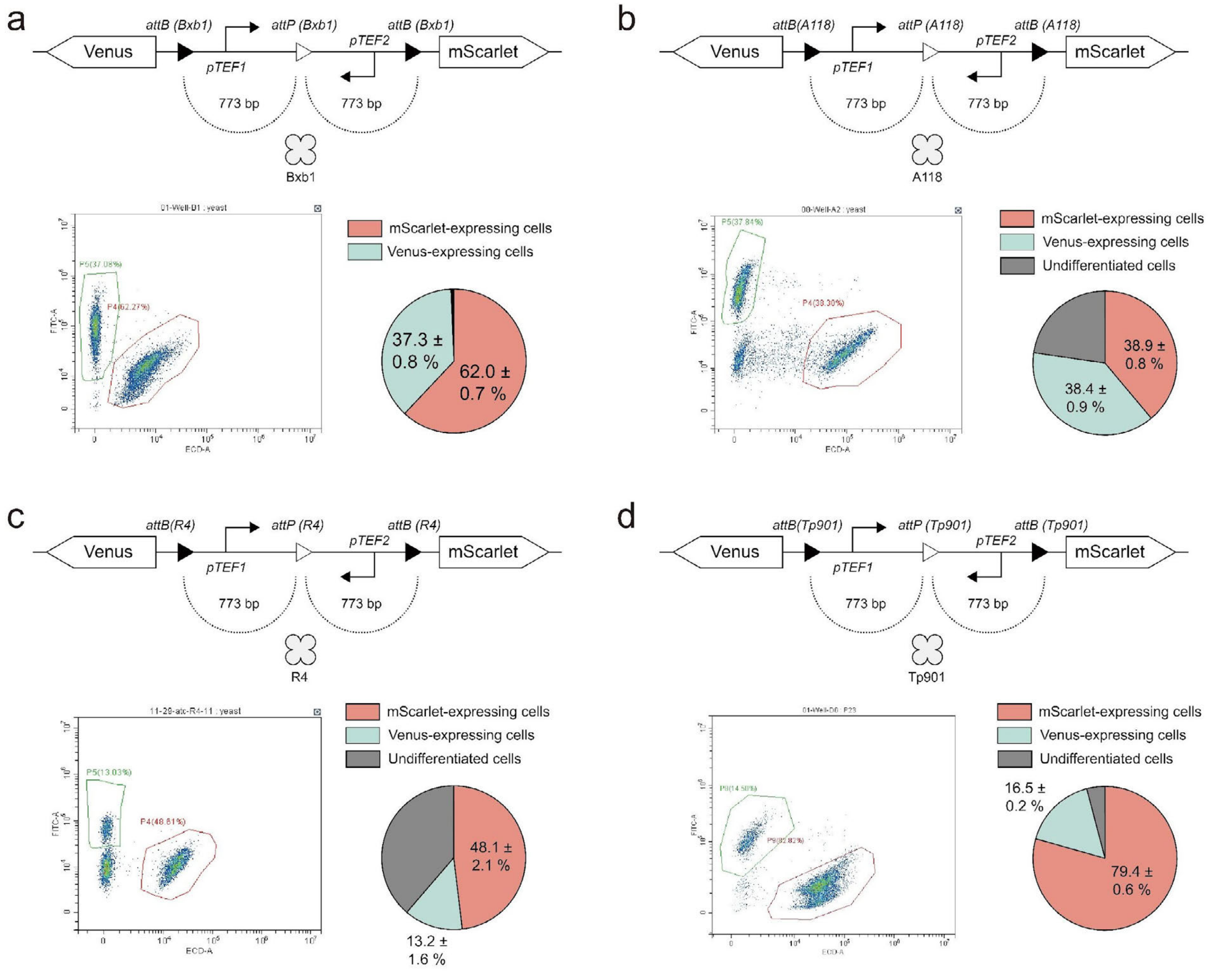

**Fig. S37 | Comparison of the recombinase performance in branching devices. a-d**. Schematics of circuit designs catalyzed by Bxb1, A118, R4, and Tp901, with representative flow cytometry data and progeny cell distribution statistics. Data are mean ± s.d. of $n = 8$ replicates.

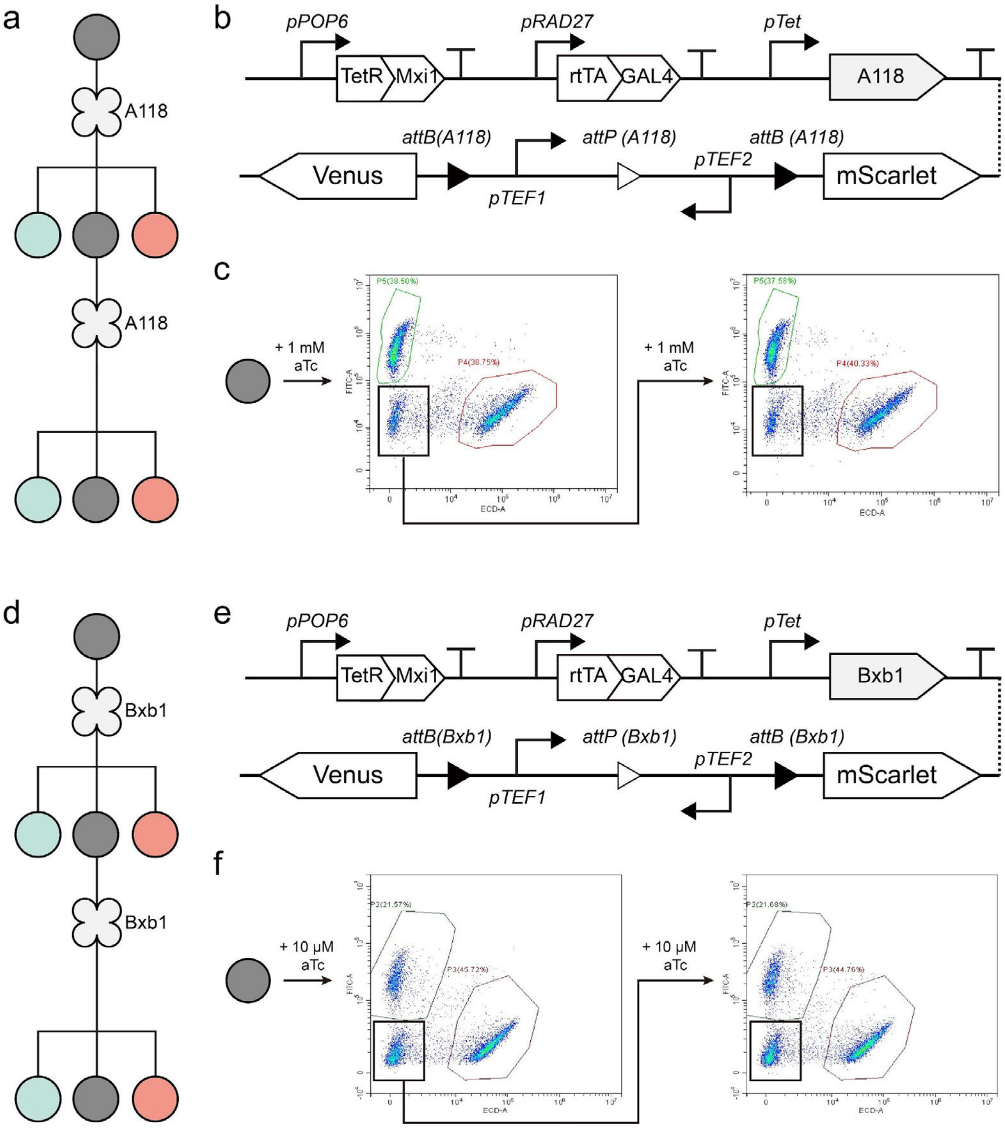

**Fig. S38 | Differentiation potential of partially differentiated yeast progeny. a**. Construction of an asymmetric differentiation system using A118 recombinase with low catalytic activity in yeast cells. **b.** Schematic of the genetic circuit. **c**. Induction with 1 mM aTc generates three types of progeny cells: red fluorescent, green fluorescent, and non-fluorescent populations. Re-induction of non-fluorescent cells with 1 mM aTc produces the same three progeny types. **d.** Partial induction of Bxb1 expression also enables asymmetric differentiation. **e**. Schematic of the genetic circuit used in the Bxb1 experiment. **f.** Representative flow cytometry data of the asymmetric differentiation system.

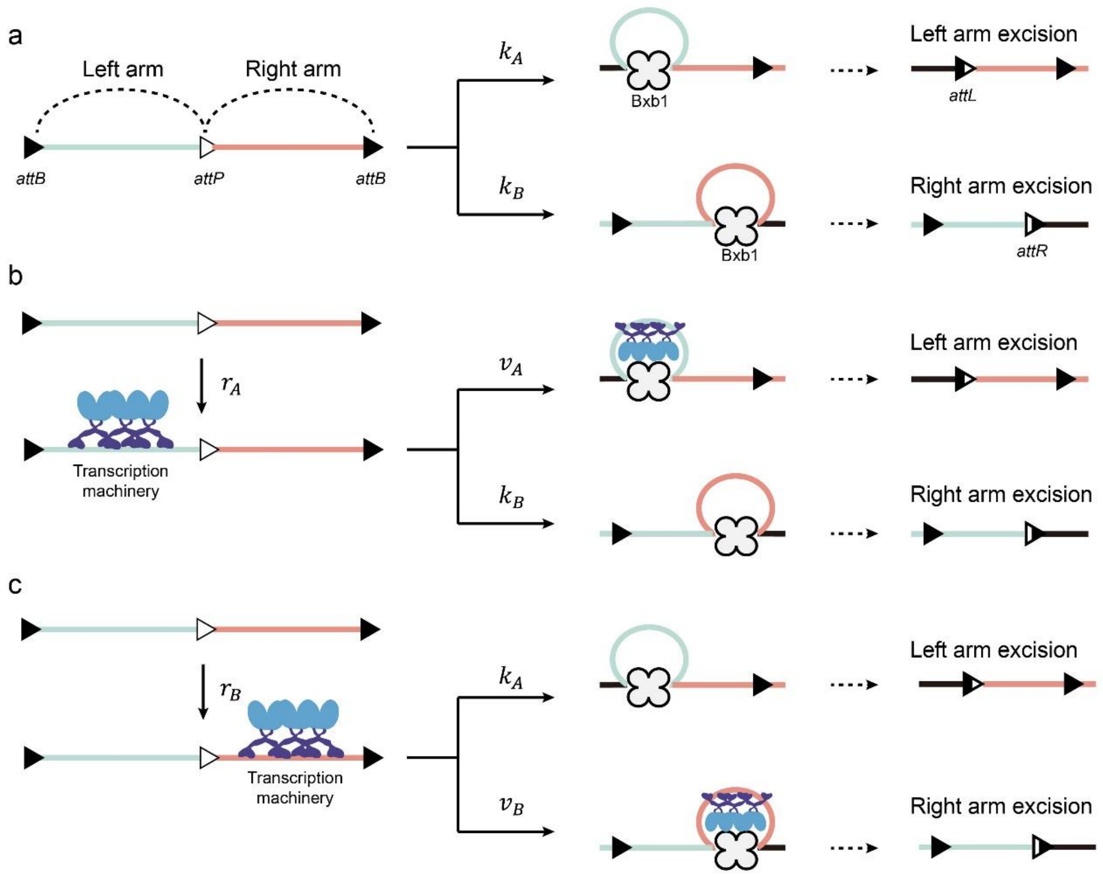

**Fig. S39 | Mathematical model predicting cell differentiation outcomes. a.** Schematic of genetic device with left and right arms cleaved by Bxb1 at rates of $k_A$ and $k_B$influenced by the *att* sites and DNA arm lengths. **b-c**. Promoters on the left or right arm recruit transcription machinery, introducing steric hindrance that reduces recombinase efficiency. The recruitment of more transcription machinery on the left arm reduces its cleavage rate from $k_A$ to $v_A$. Similarly, increased transcription machinery recruitment on the right arm changes its cleavage rate from $k_B$ to $v_B$. $r_A$ and $r_B$ are the binding rates of the transcription machinery. A variable $\alpha$ was used to describe the strength of the steric hindrance: $\alpha = \frac{v_A}{k_A} = \frac{v_B}{k_B}$.

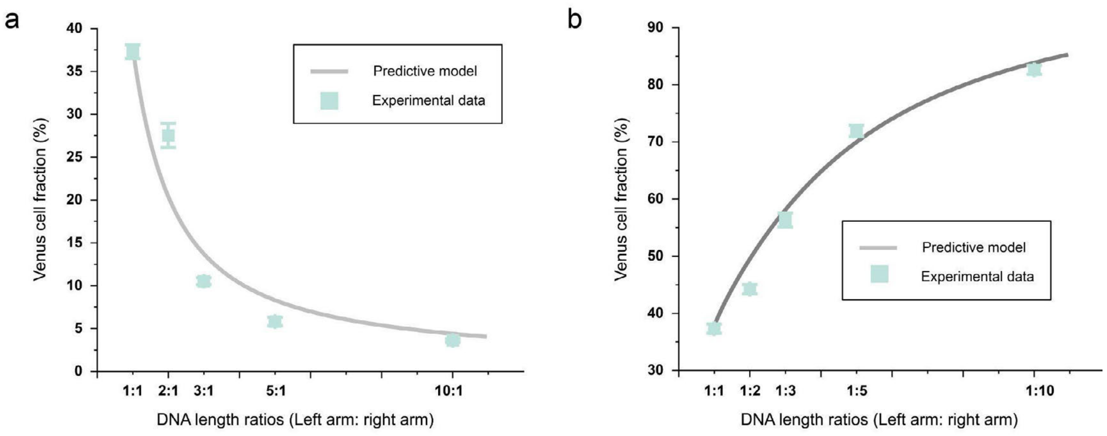

**Fig. S40 | Predictive modeling of arm length effects on differentiation. a.** Comparison of model predictions with experimental data for Venus-expressing cell ratios across DNA length ratios (left to right arm) from 1:1 to 10:1, based on circuits in **Fig. S25**. **b.** A similar comparison for DNA length ratios ranging from 1:1 to 1:10 based on circuits in **Fig. S24**. Gray lines represent model predictions, and green dots indicate experimental data (data are mean ± s.d. of $n = 8$ replicates).

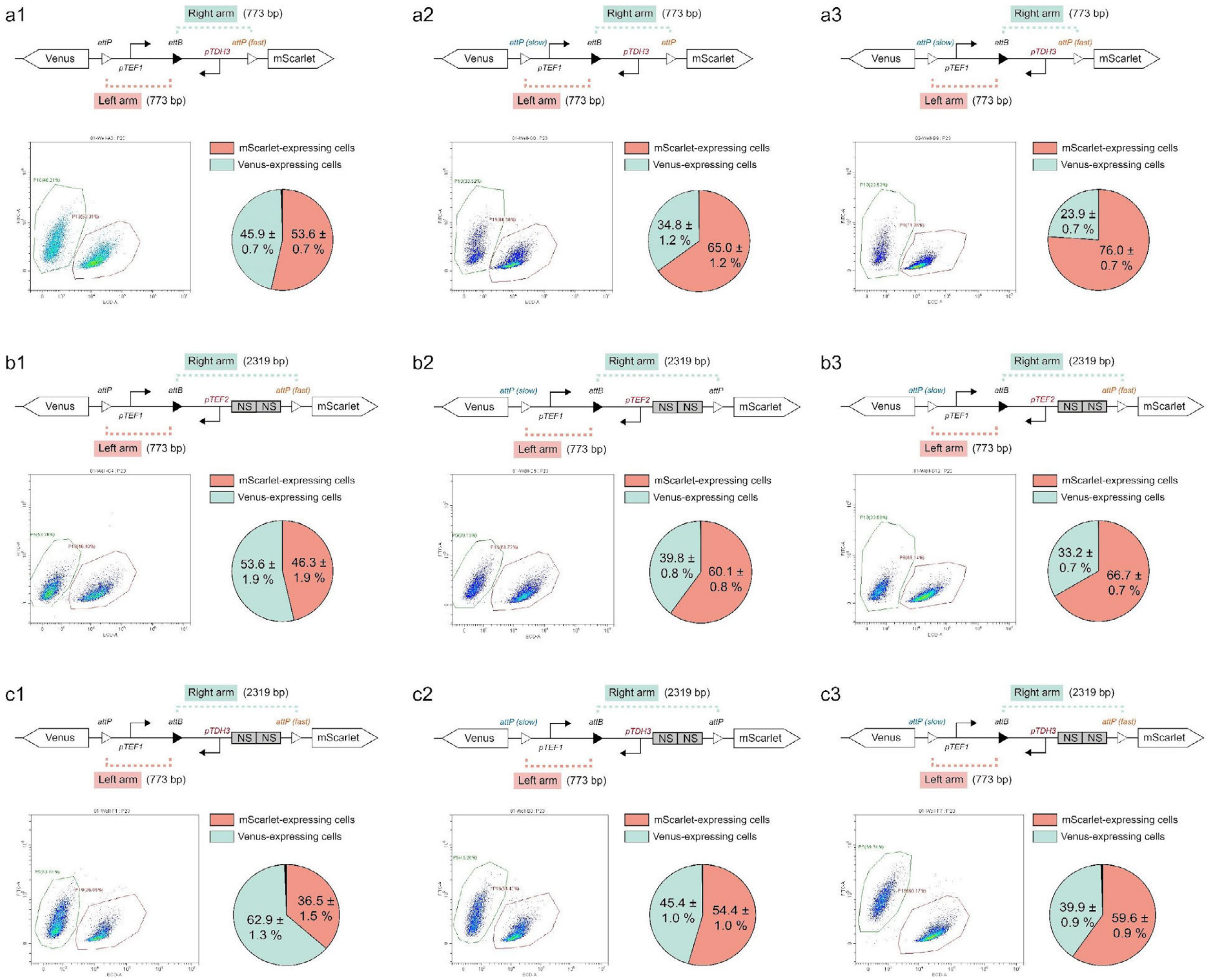

**Fig. S41 | Design of nine circuits to validate predictive modeling accuracy.** All designs incorporate variables such as arm length, *attP* variants, and promoter types. Data are mean ± s.d. of $n = 8$ replicates. The a1-c3 arrangement corresponds to the design presented in **Figure 2i** of the main text.

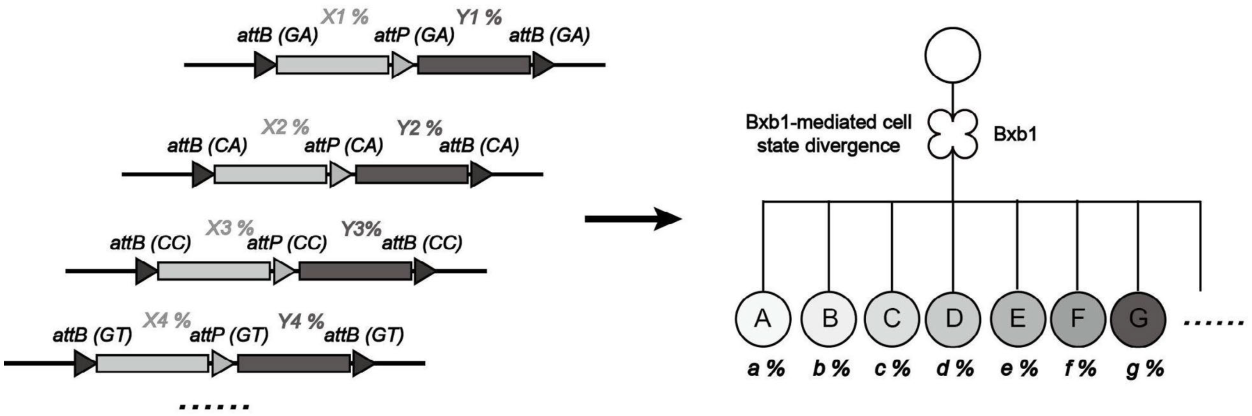

**Fig. S42 | Design principles for generating yeast multicellularity with parallel circuits.** Integration of multiple orthogonal recombinase circuits allows precise control over progeny phenotypes. With "n" orthogonal circuits, $2^n$ distinct progeny phenotypes can be generated.

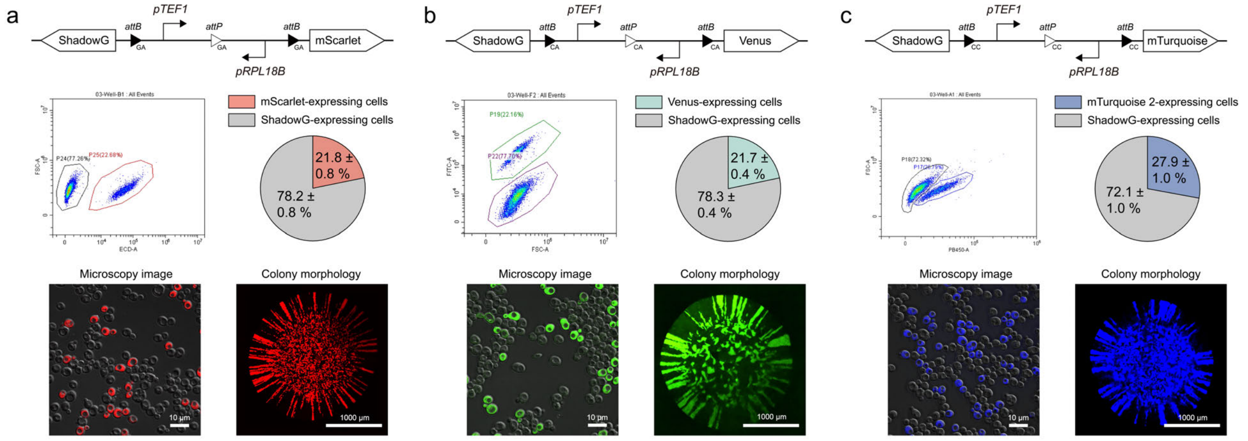

**Fig. S43 | Characterization of representative 1-layer orthogonal circuits.** Designs incorporate *pTEF1* and *pRPL18B* promoters, with reporters mScarlet (**a**), Venus (**b**), or mTurquoise 2 (**c**). Data are mean ± s.d. of $n = 8$ replicates.

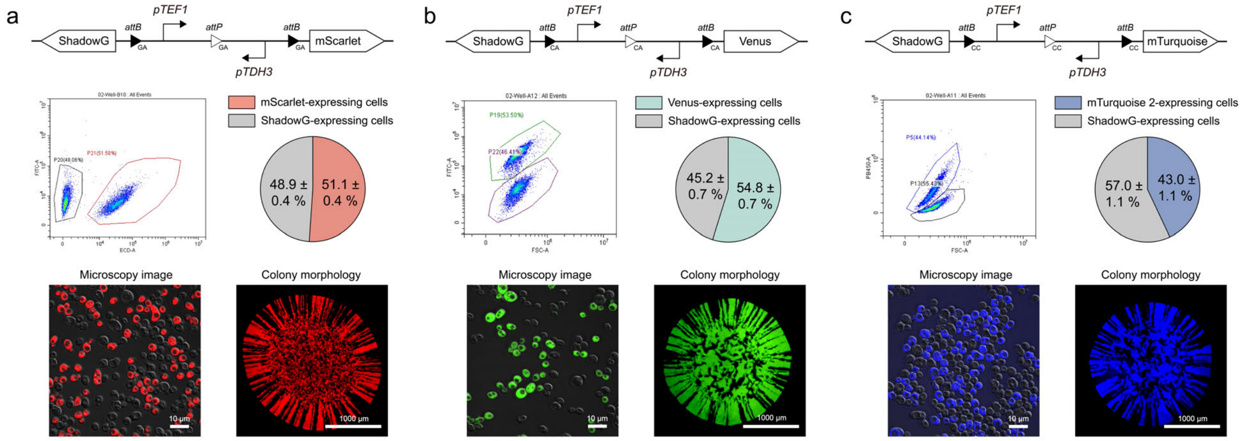

**Fig. S44 | Characterization of representative 1-layer orthogonal circuits using *pTEF1* and *pTDH3* as promoters.** Data are mean ± s.d. of *n* = 8 replicates.

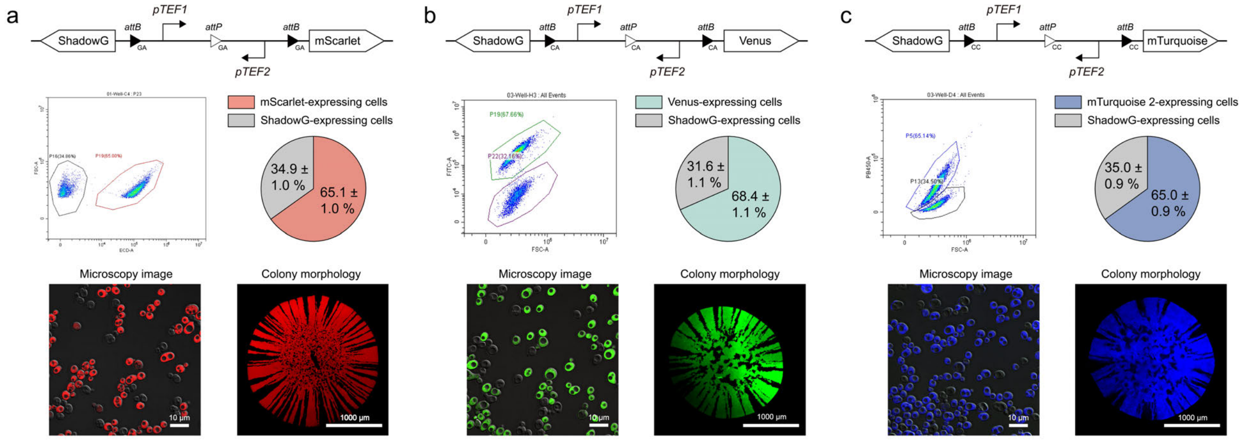

**Fig. S45 | Characterization of representative 1-layer orthogonal circuits using *pTEF1* and *pTEF2* as promoters.** Data are mean ± s.d. of *n* = 8 replicates.

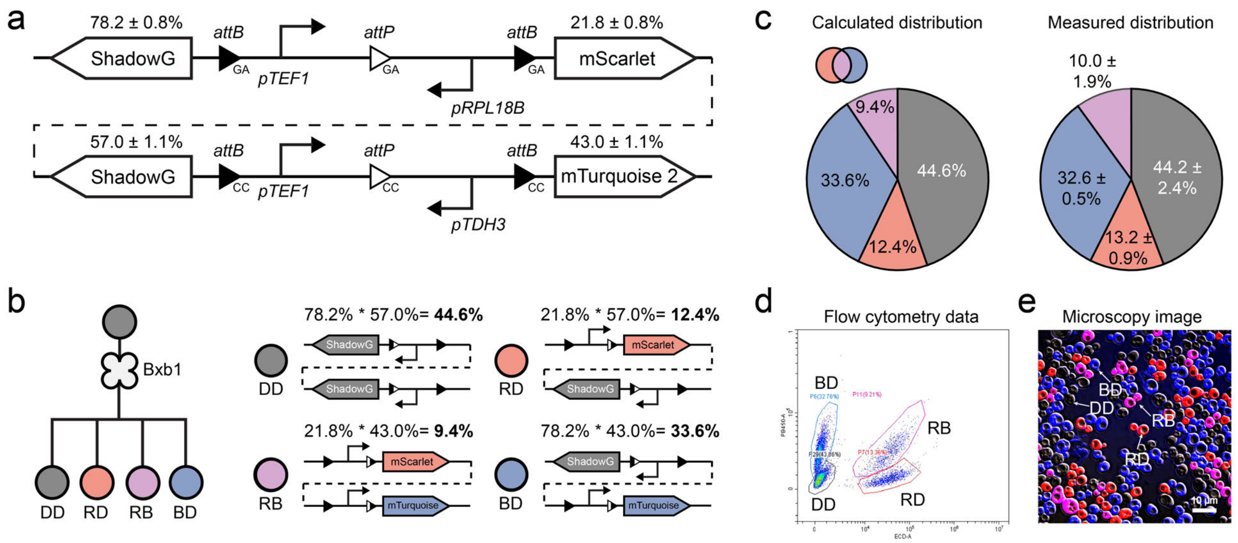

**Fig. S46 | Design and characterization of a representative 2× orthogonal circuit. a.** Schematic of a 2× orthogonal genetic device. **b.** Theoretical progeny types and proportions generated by the circuits. **c.** Comparison of theoretical progeny proportions with flow cytometry data. **d.** Flow cytometry identifies 4 distinct cell populations. **e.** Fluorescence microscopy confirms the presence of 4 distinct progeny types. Data are mean ± s.d. of $n = 8$ replicates.

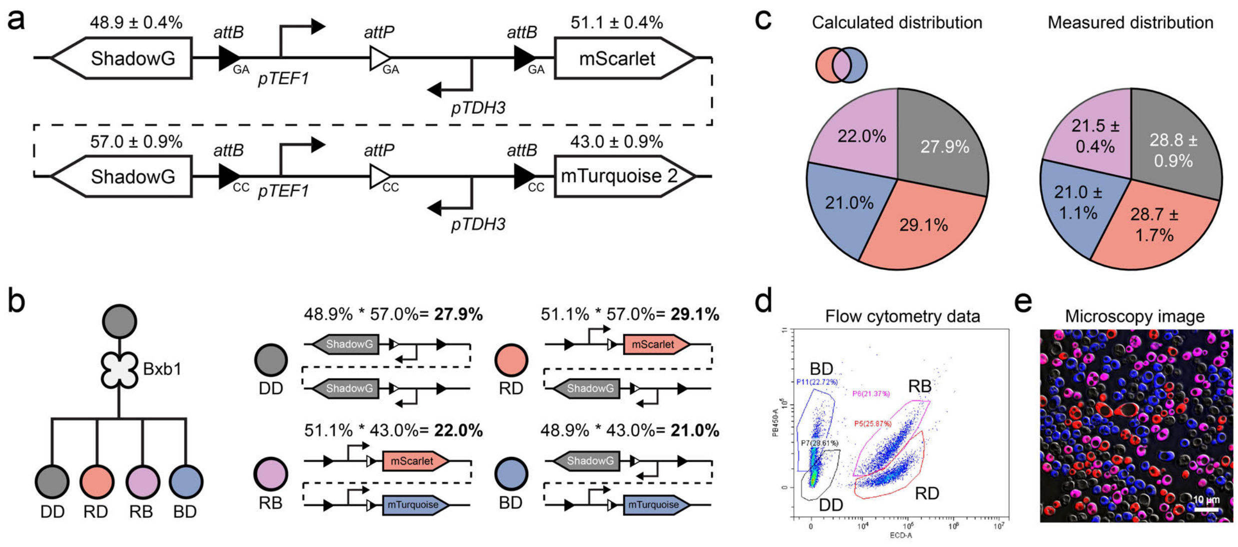

**Fig. S47 | Characterization of a 2× orthogonal circuit with modified promoter usage.** Compared to **Fig. S46**, changes in promoter design altered progeny distributions. Data are mean ± s.d. of $n = 8$ replicates.

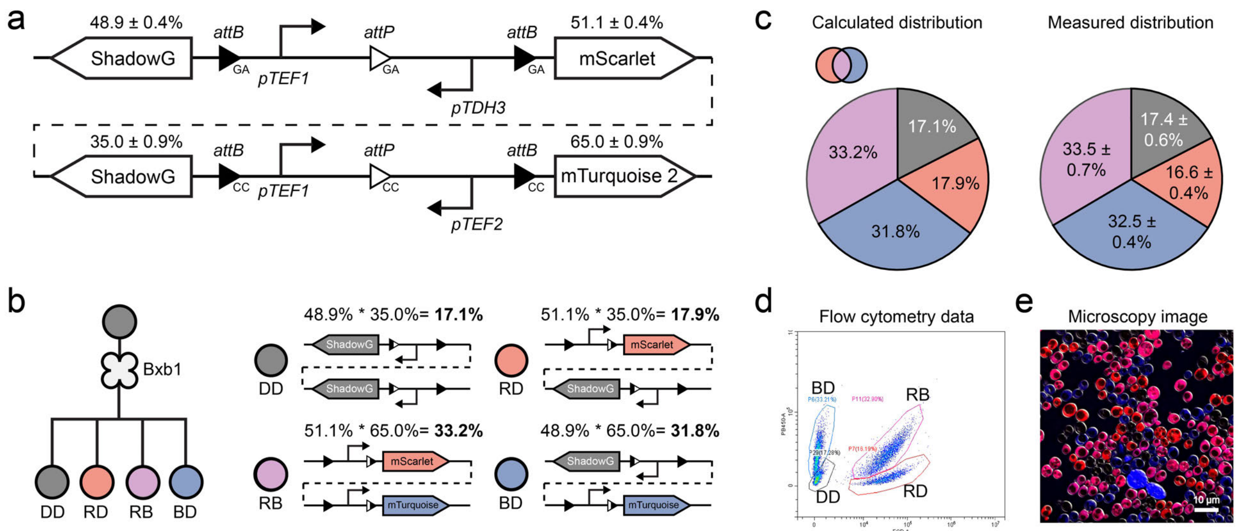

**Fig. S48 | Characterization of a 2× orthogonal circuit with modified promoter usage.** Compared to **Fig. S46**, changes in promoter design altered progeny distributions. Data are mean ± s.d. of $n = 8$ replicates.

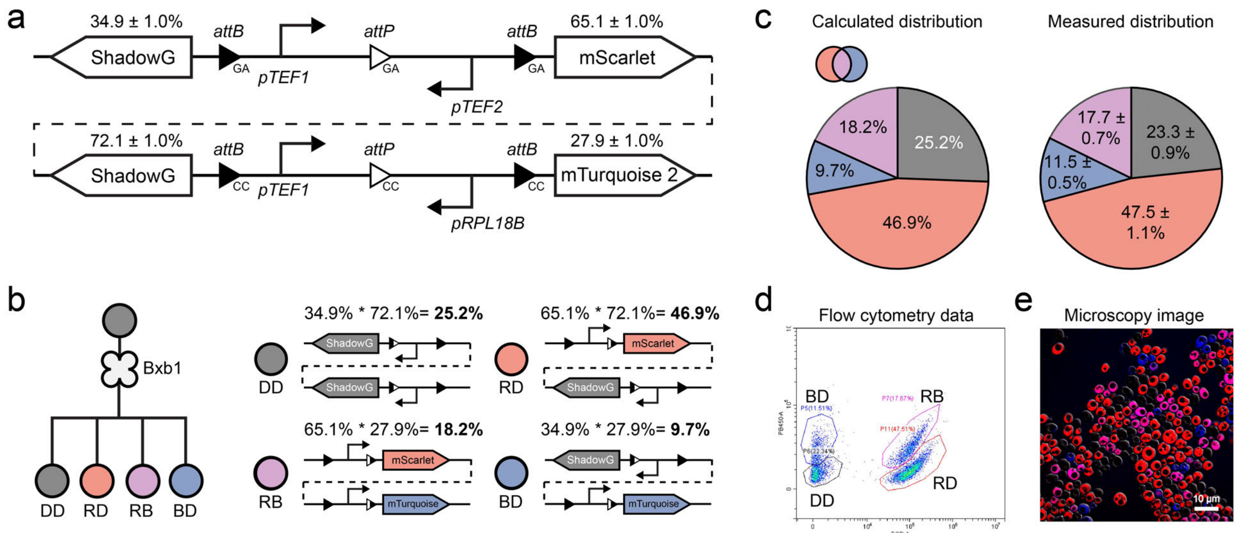

**Fig. S49 | Characterization of a 2× orthogonal circuit with modified promoter usage.** Compared to **Fig. S46**, changes in promoter design altered progeny distributions. Data are mean ± s.d. of $n = 8$ replicates.

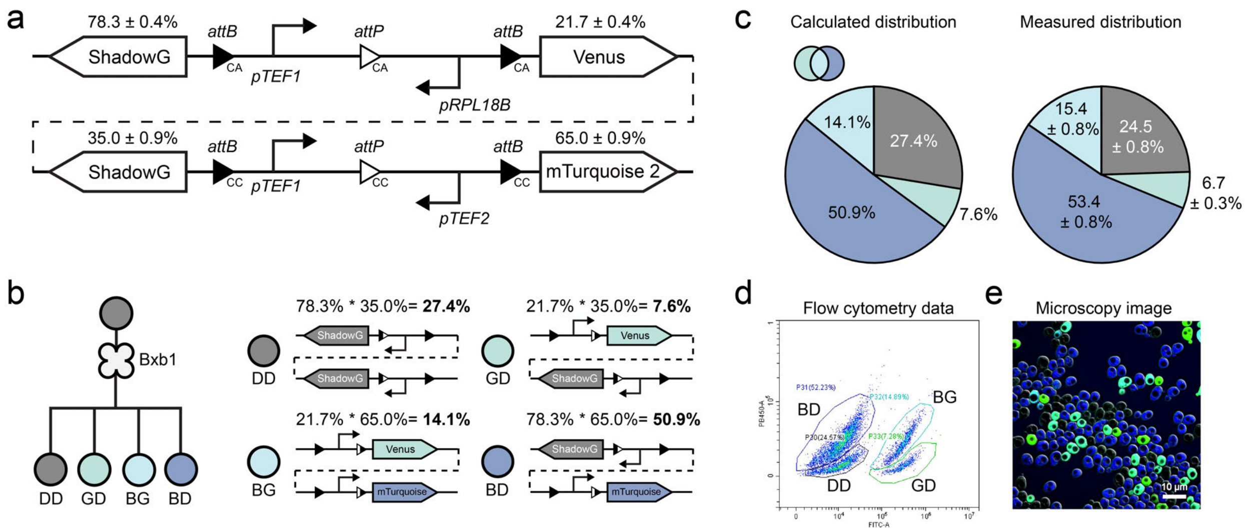

**Fig. S50 | Characterization of a 2× orthogonal circuit using three reporters.** The circuit was designed with ShadowG, Venus, and mTurquoise 2 reporters. Data are mean ± s.d. of $n = 8$ replicates.

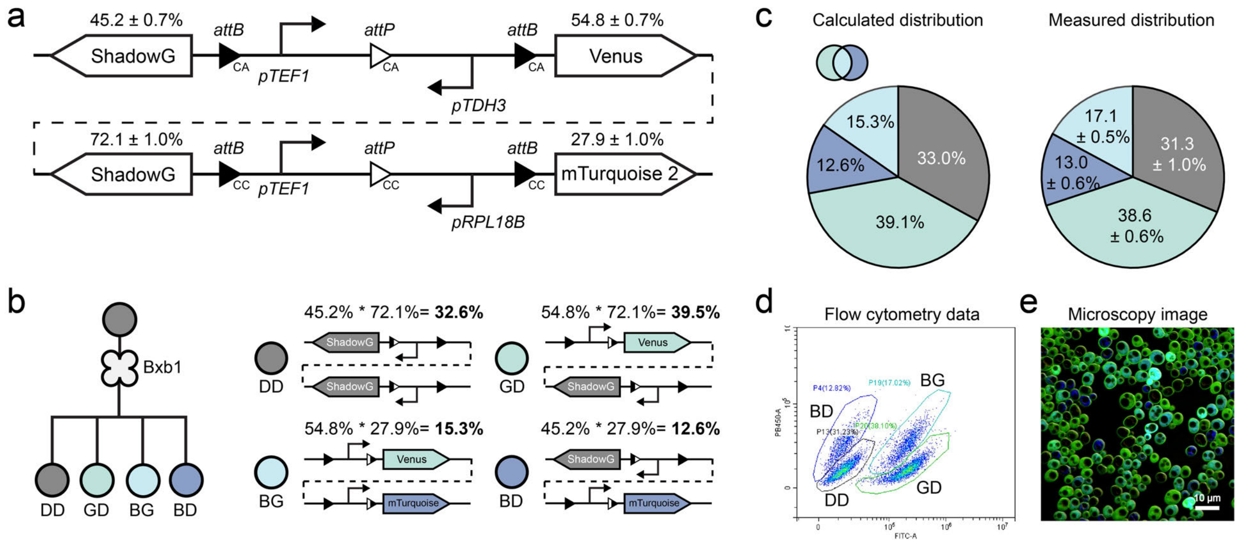

**Fig. S51 | Characterization of a 2× orthogonal circuit with modified promoter usage.** Compared to **Fig. S50**, changes in promoter design altered progeny distributions. Data are mean ± s.d. of $n = 8$ replicates.

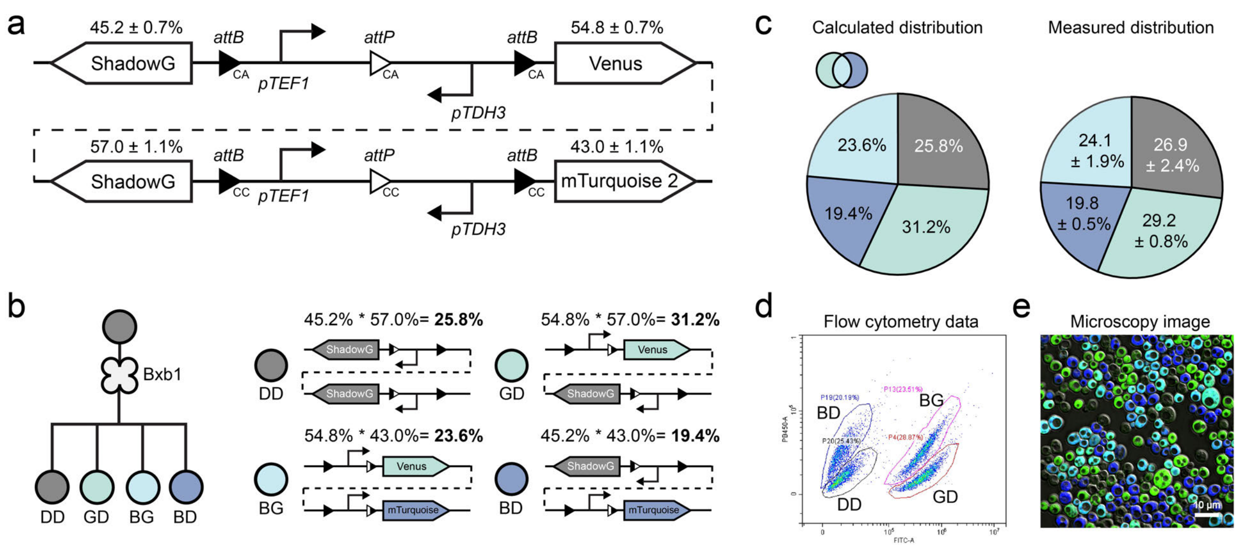

**Fig. S52 | Characterization of a 2× orthogonal circuit with alternative promoter configurations.** Compared to **Fig. S50**, changes in promoter design altered progeny distributions. Data are mean ± s.d. of $n = 8$ replicates.

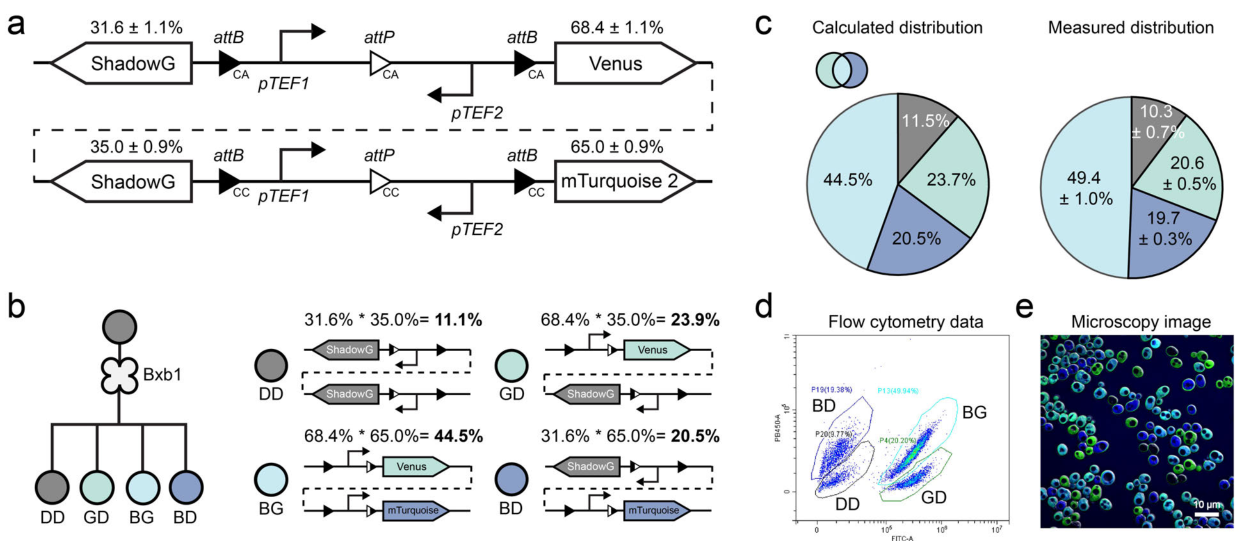

**Fig. S53 | Characterization of a 2× orthogonal circuit with alternative promoter configurations.** Compared to **Fig. S50**, changes in promoter design altered progeny distributions. Data are mean ± s.d. of $n = 8$ replicates.

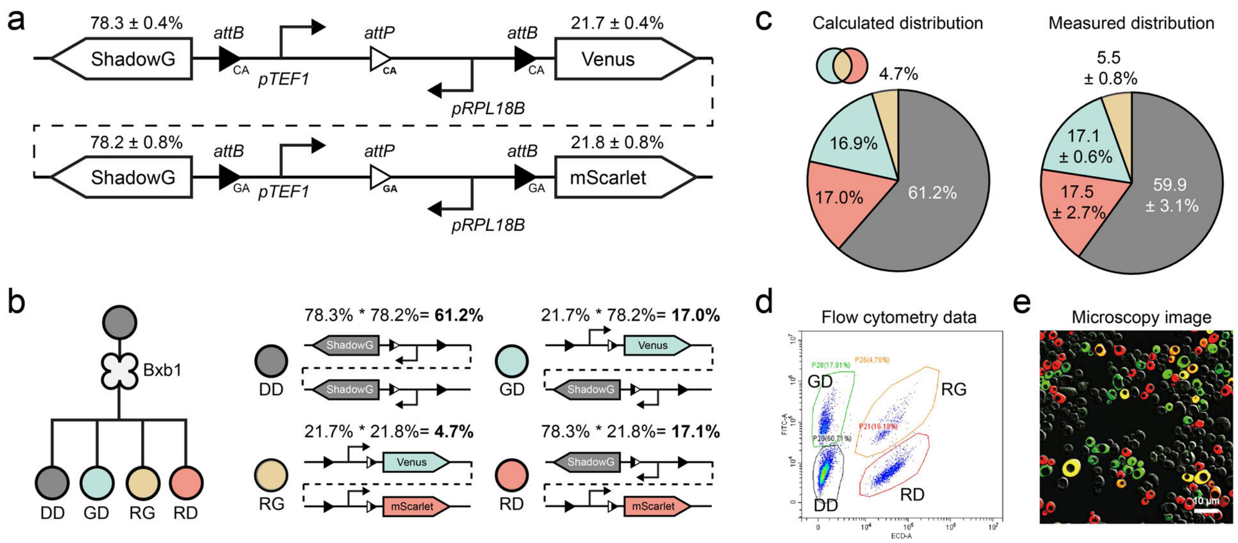

**Fig. S54 | Characterization of a 2× orthogonal circuit using three reporters.** The circuit was designed with ShadowG, Venus, and mScarlet reporters. Data are mean ± s.d. of $n = 8$ replicates.

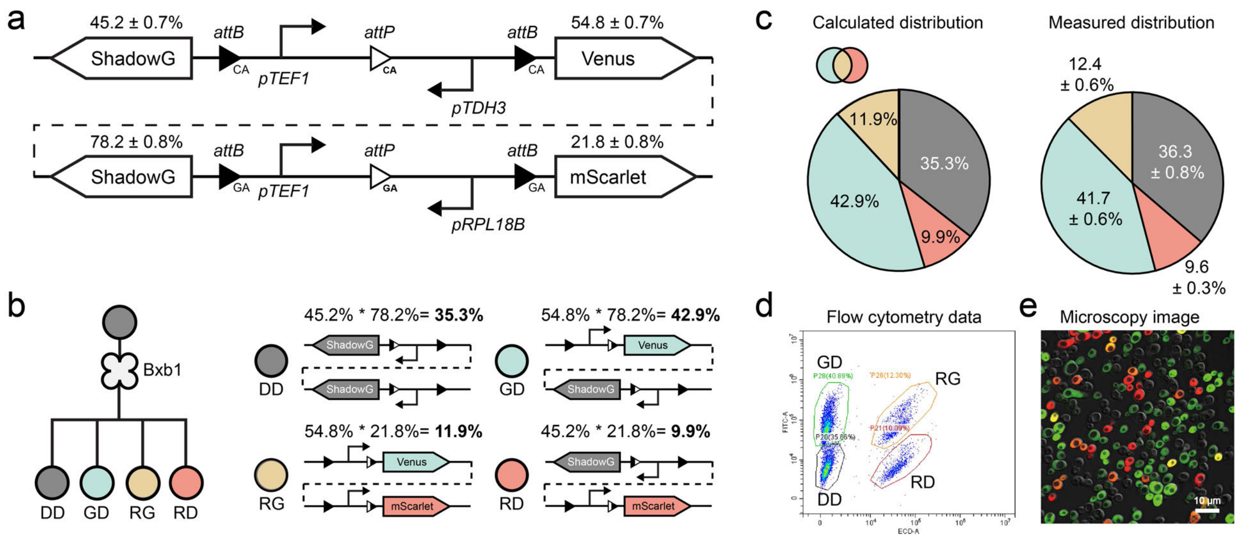

**Fig. S55 | Characterization of a 2× orthogonal circuit with modified promoter configurations.** Compared to **Fig. S54**, changes in promoter design altered progeny distributions. Data are mean ± s.d. of $n = 8$ replicates.

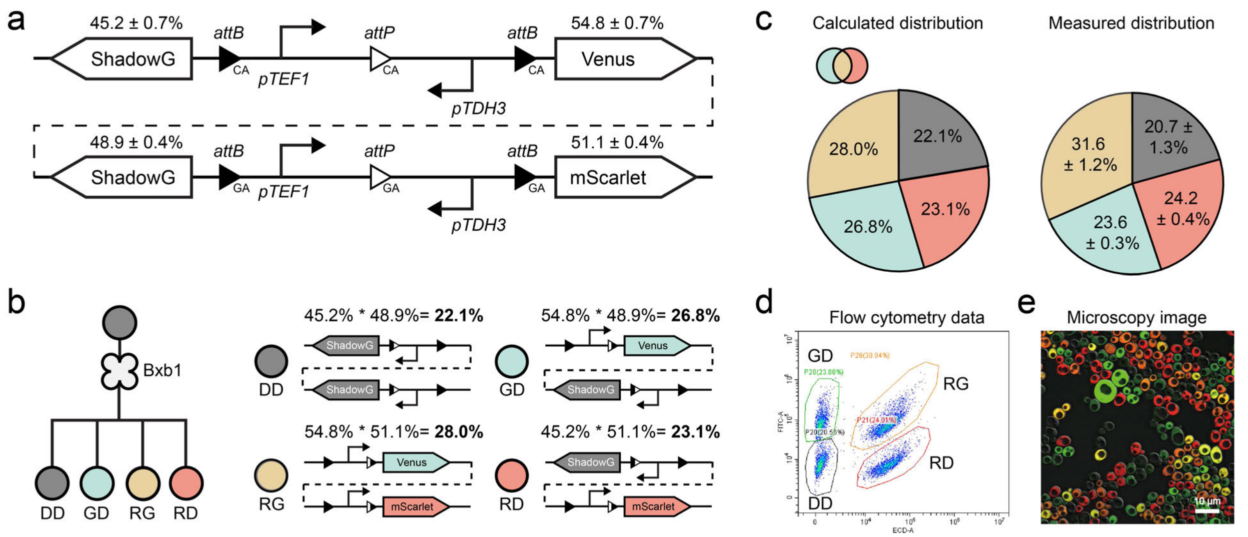

**Fig. S56 | Characterization of a representative 2× orthogonal circuit with modified promoter configuration.** Compared to **Fig. S54**, changes in promoter design altered progeny distributions. Data are mean ± s.d. of *n* = 8 replicates.

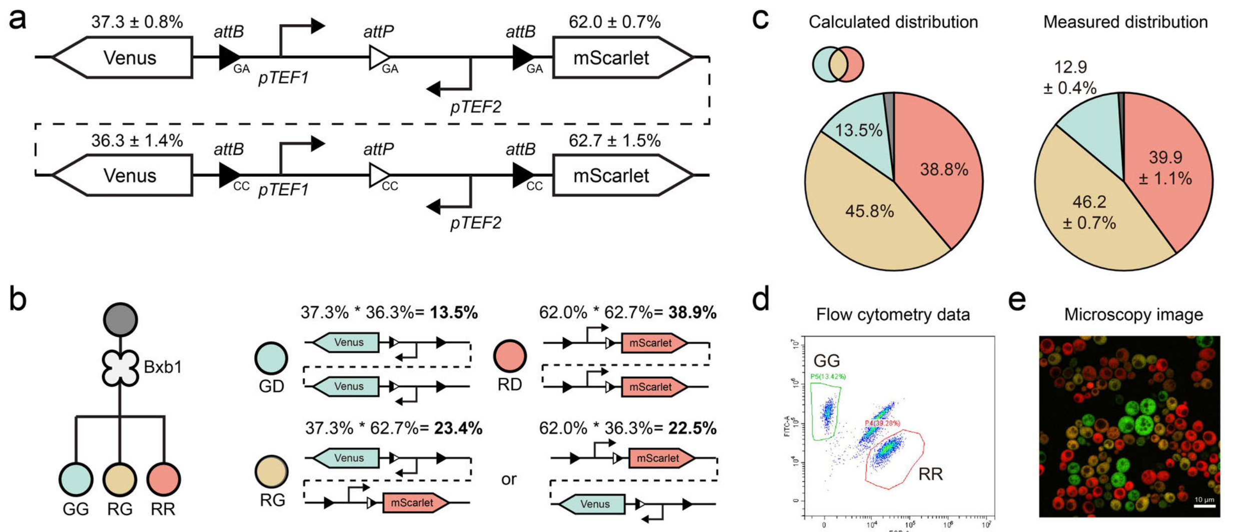

**Fig. S57 | Characterization of a 2× orthogonal circuit using two reporters.** The circuit was designed with Venus and mScarlet reporters. Data are mean ± s.d. of *n* = 8 replicates.

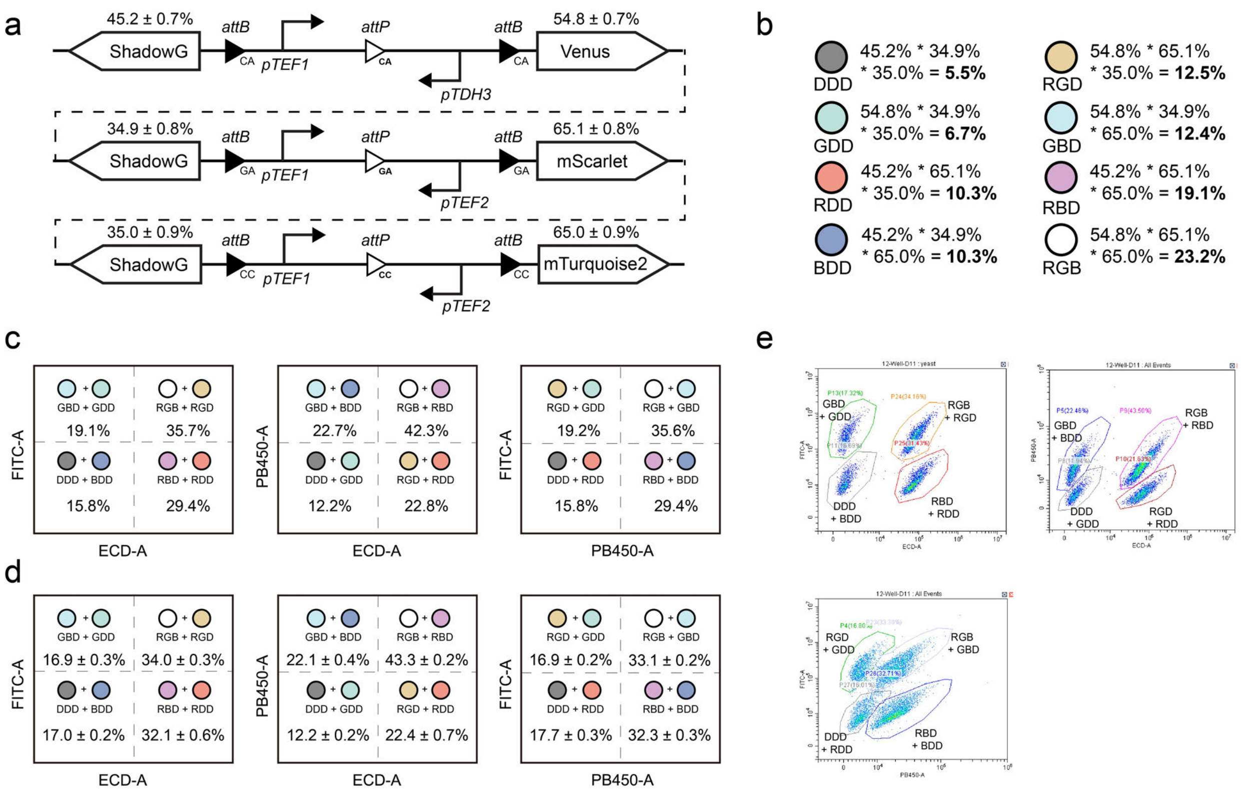

**Fig. S58 | Design and characterization of a representative 3× orthogonal circuit. a.** The 3× orthogonal gene circuits were designed, with the first circuit using CA versions of *att* sites, and *pTEF1* and *pTDH3* promoters to express ShadowG and Venus reporter genes. The second circuit employed GA *att* site versions and *pTEF1* and *pTEF2* as promoters, with ShadowG and mScarlet as reporters. The third circuit used CC *att* site versions and *pTEF1* and *pTEF2* as promoters, expressing ShadowG and mTurquoise 2. The progeny cell proportions represented the probabilities of reporter gene expression in the branching device. **b.** Theoretical proportions of progeny cell types generated by the designed branching device. **c.** Theoretical yeast consortia distributions as detected by flow cytometry. **d**. Experimentally measured progeny cell distributions using flow cytometry. **e.** Representative flow cytometry data showing progeny cell distributions from the 3× orthogonal circuit. Data are mean ± s.d. of *n* = 8 replicates.

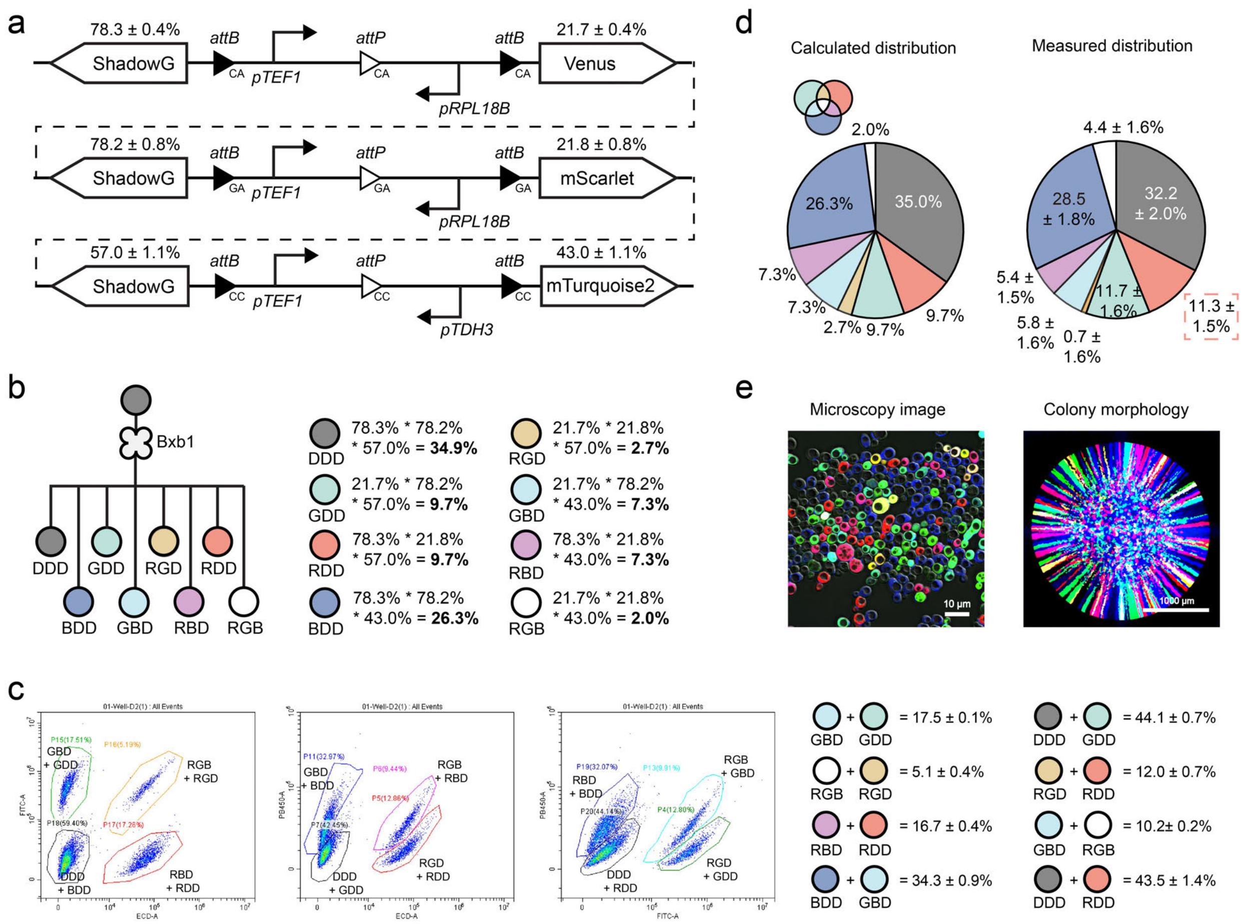

**Fig. S59 | Design and characterization of a representative 3× orthogonal circuit**. **a.** Design of the circuit. **b.** Theoretical types and proportions of progeny cells generated by the 3× orthogonal circuits. **c.** Representative flow cytometry images and statistical data showing progeny cell distributions. **d.** Theoretical yeast progeny proportions were compared to experimental values. Fluorescence imaging was used to determine the proportion of RDD cells, and these values were combined with flow cytometry data to estimate detailed progeny percentages. **e.** Fluorescence microscopy images display eight distinct yeast populations. Data are mean ± s.d. of $n = 8$ replicates.

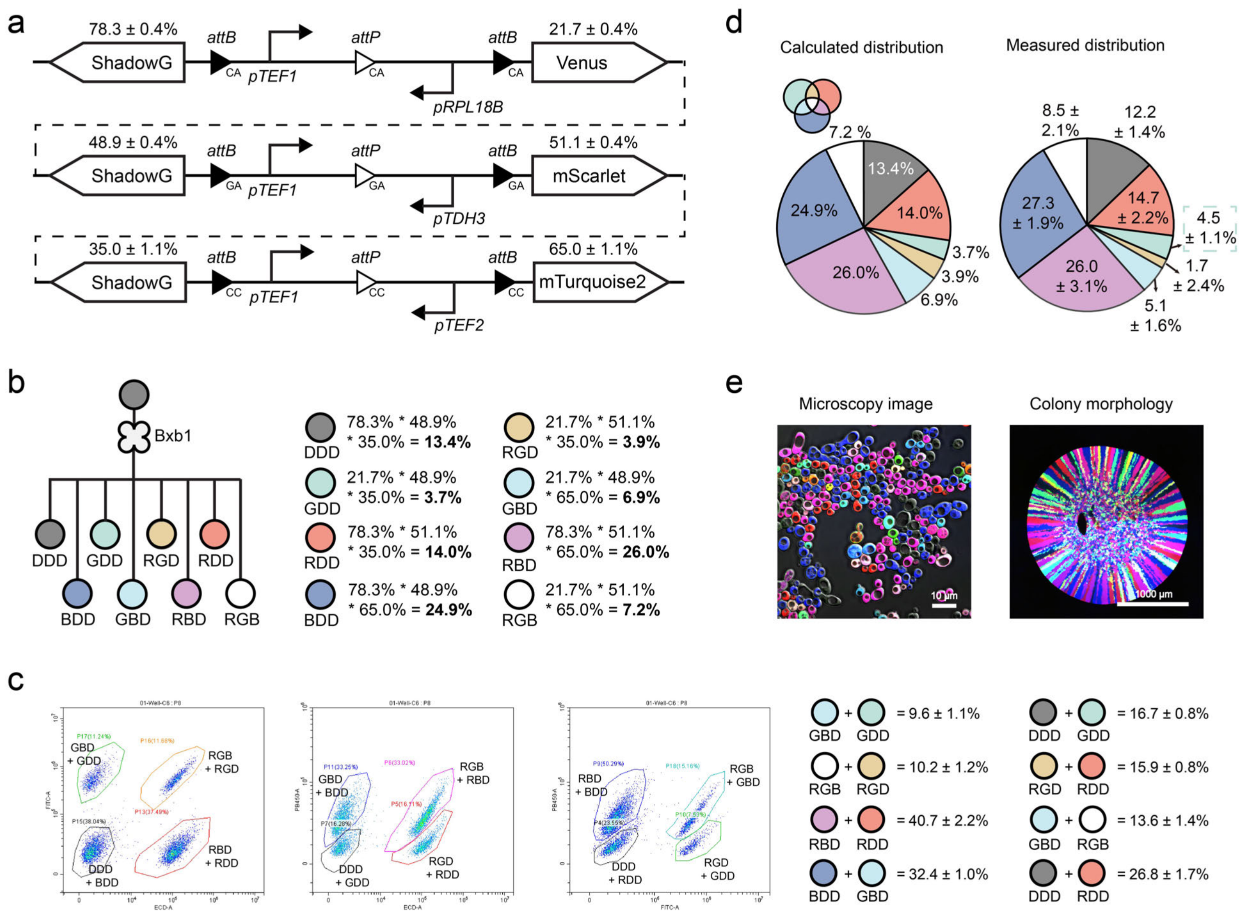

**Fig. S60 | Design and characterization of a representative 3× orthogonal circuit. a.** Design of the circuit. **b.** Theoretical types and proportions of progeny cells generated by the circuits. **c.** Representative flow cytometry images and statistical data showing progeny cell distributions. **d.** Theoretical yeast progeny proportions were compared to experimental values. Fluorescence imaging was used to estimate the proportion of GDD cells, and these values were combined with equations presented in **Fig. S60c** to calculate the distribution of all progeny strains. **e**. Fluorescence microscopy images display eight distinct yeast populations. Data are mean ± s.d. of *n* = 8 replicates.

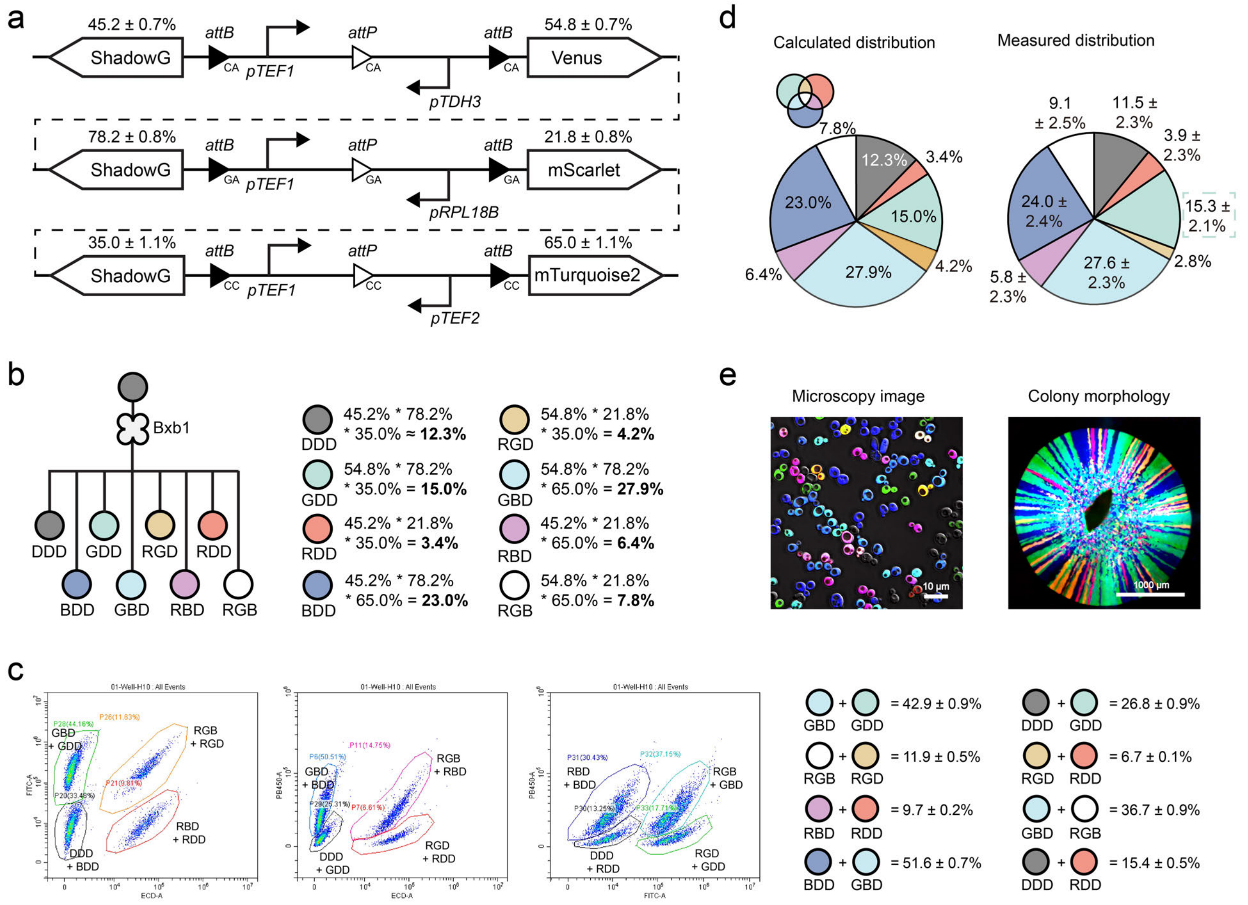

**Fig. S61 | Design and characterization of a representative 3× orthogonal circuit. a.** Design of the circuit. **b.** Theoretical types and proportions of progeny cells generated by the circuits. **c.** Representative flow cytometry images and statistical data showing progeny cell distributions. **d.** Theoretical yeast progeny proportions were compared to experimental values. Fluorescence imaging was used to estimate the proportion of GDD cells, and these values were combined with the equations presented in **Fig. S61c** to calculate the distribution of all progeny strains. **e**. Fluorescence microscopy images display eight distinct yeast populations. Data are mean ± s.d. of $n = 8$ replicates.

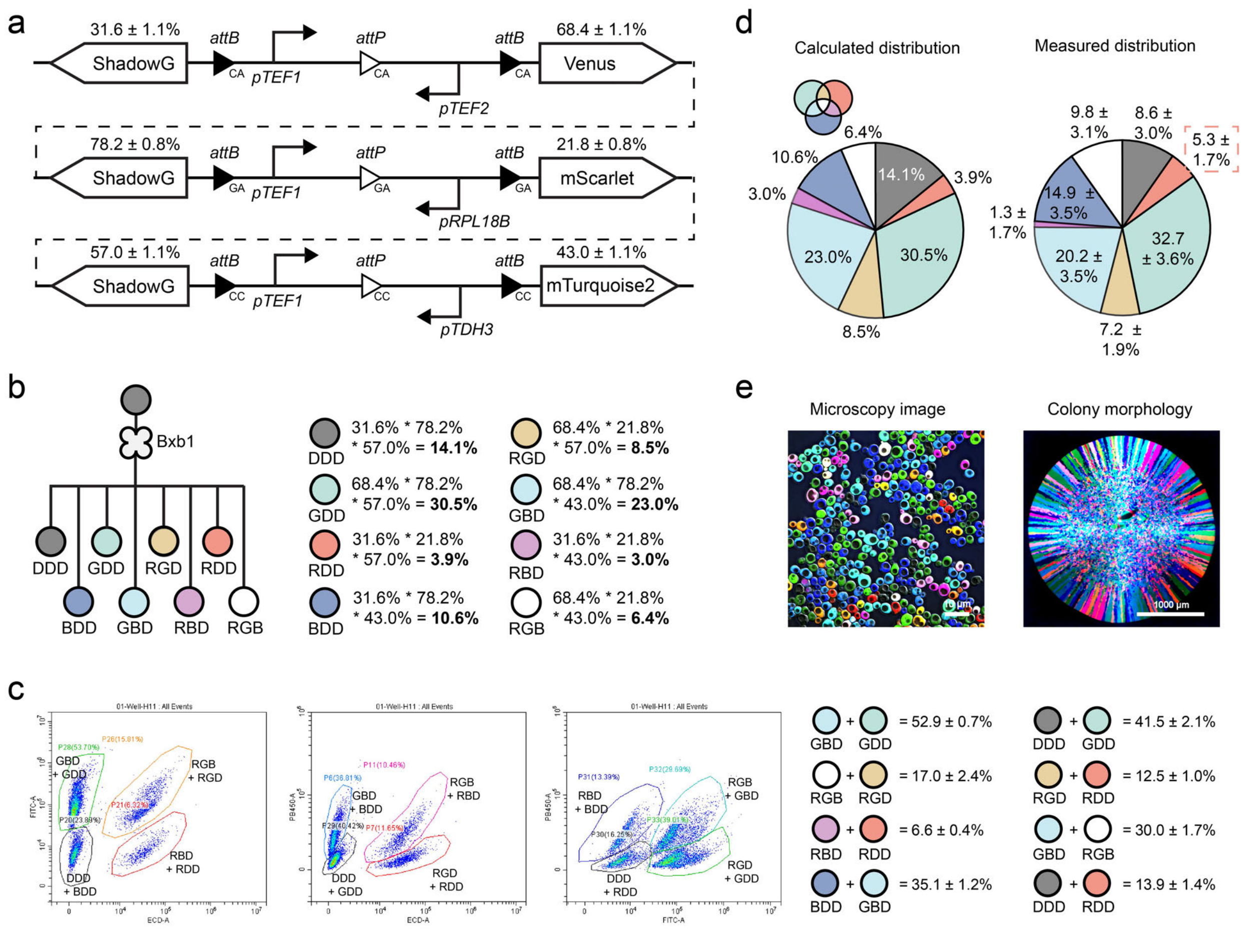

**Fig. S62 | Design and characterization of a representative 3× orthogonal circuit**. **a.** Design of the circuits. **b.** Theoretical types and proportions of progeny cells generated by the circuits. **c.** Representative flow cytometry images and statistical data showing progeny cell distributions. **d.** Theoretical yeast progeny proportions were compared to experimental values. Fluorescence imaging was used to estimate the proportion of RDD cells, and these values were combined with equations presented in **Fig. S62c** to calculate the distribution of all progeny strains. **e**. Fluorescence microscopy images display eight distinct yeast populations. Data are mean ± s.d. of *n* = 8 replicates.

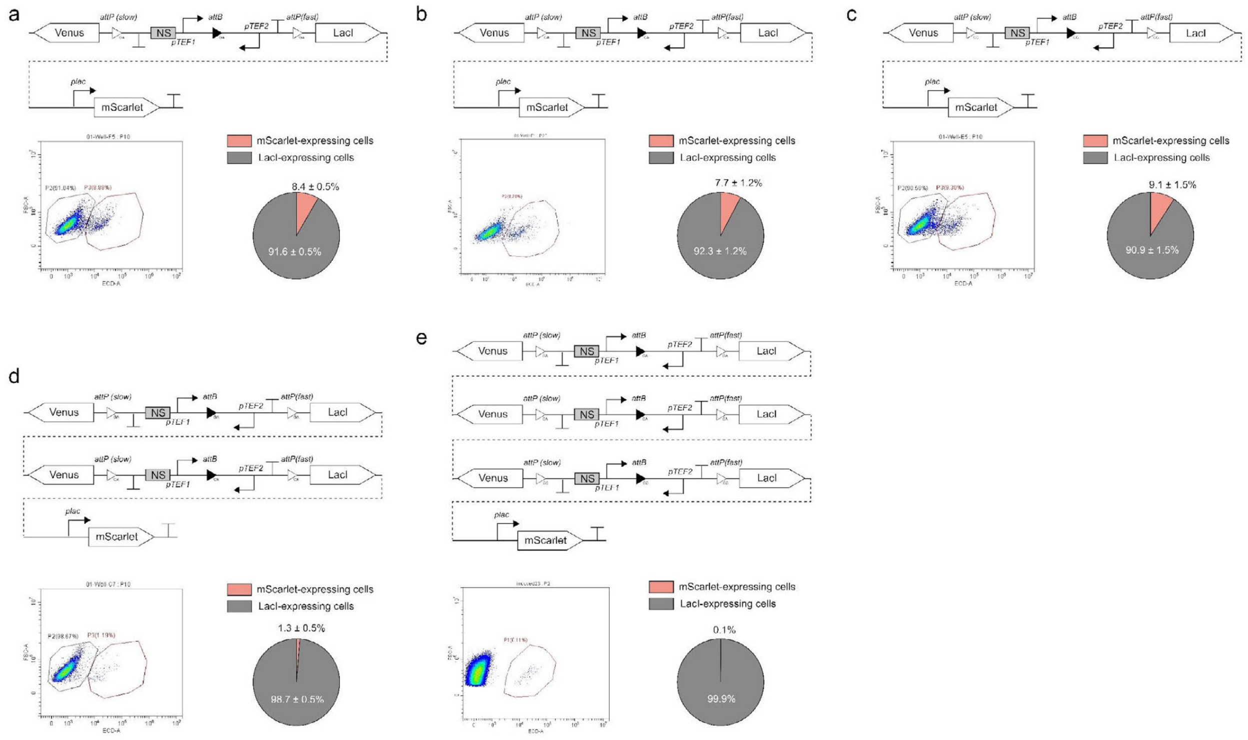

**Fig. S63 | Design and characterization of parallel branching devices for achieving extreme ratios. a-c.** Three gene circuits were designed using orthogonal *att* site variants (GA, CA, and CC), each using *pTEF1* and *pTEF2* promoters with Venus and LacI as reporter genes. LacI expression inhibits the *pLac* promoter, enabling only progeny yeast with left-arm excision to produce mScarlet. The probabilities of generating mScarlet-producing strains across these circuits vary from 7.7 ± 1.2% to 9.1 ± 1.5%. **d.** With mScarlet expression probabilities of 8.4 ± 0.5% and 7.7 ± 1.2%, the calculated probability for mScarlet expression in strains with the 2× orthogonal circuits is approximately 0.6%, closely aligning with the observed experimental value of 1.3 ± 0.5%. **e.** For strains with 3× orthogonal gene circuits, the probabilities of mScarlet expression in each were 8.4 ± 0.5%, 7.7 ± 1.2%, and 9.1 ± 1.5%. This results in a combined mScarlet expression probability of around 0.1% in theory, consistent with the observed experimental value. Data are mean ± s.d. of *n* = 6 replicates.

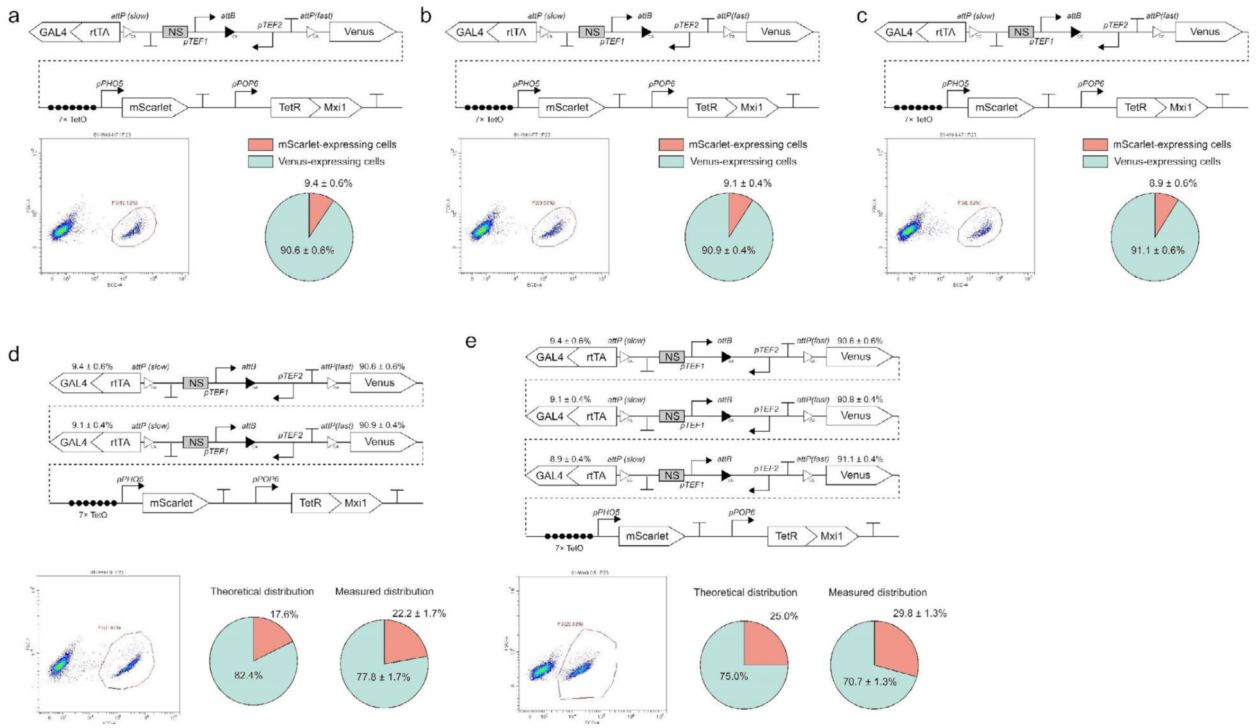

**Fig. S64 | Design and characterization of parallel branching devices for controlling progeny cell compositions. a-c**. Gene circuits were designed using three orthogonal *att* site variants (GA, CA, and CC), respectively. Due to the presence of repressor TetR-Mxi1, the expression of mScarlet was inhibited. In each circuit, there is approximately a 9% probability of producing rtTA-GAL4, which interacts with aTc molecules to trigger mScarlet expression, producing mScarlet and emitting red fluorescence. **d.** The progeny cell composition is generated from the founder cell encoding two orthogonal gene circuits. The likelihood of not expressing mScarlet is the chance that neither circuit produces rtTA-GAL4. Thus, in these two orthogonal circuits, the probability of mScarlet expression in progeny cells is $(1 - 0.906 \times 0.909) \times 100\% \approx 17.6\%$. The statistical data from flow cytometry revealed that the proportion of mScarlet-expressing yeast was 22.2 ± 1.7%, closely aligning with the theoretical values. **e.** The progeny cell composition arises from mother cells encoding three orthologous gene circuits. The likelihood of not expressing mScarlet is the chance that none of these three circuits produces rtTA-GAL4. The probability of mScarlet expression in progeny cells is $(1 - 0.906 \times 0.909 \times 0.911) \times 100\% \approx 25.0\%$. The statistical data from flow cytometry revealed that the proportion of mScarlet-expressing yeast was 29.8 ± 1.3%, closely aligning with the theoretical values. Data are mean ± s.d. of $n = 6$ replicates.

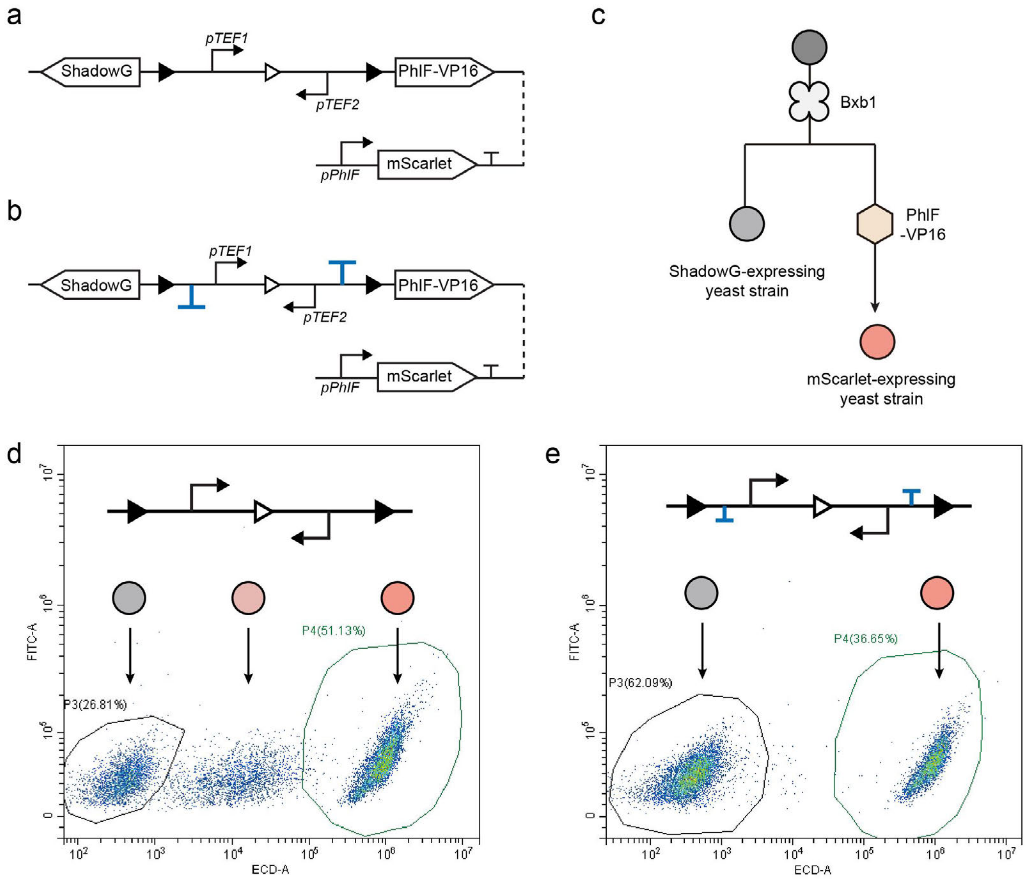

**Fig. S65 | Enhancing gene circuit performance by incorporating terminators between *att* sites. a.** The branching circuit in this experiment was designed to produce yeast progeny expressing either ShadowG or PhIF-VP16, a transcriptional factor that induces mScarlet expression via the *pPhIF* promoter. Leaky expression of PhIF-VP16 could convert non-fluorescent ShadowG-expressing yeast into red fluorescent cells. **b**. To prevent leaky expression of the PhIF-VP16, two yeast terminators were inserted between the *att* sites in the genetic circuit. **c**. Founder cells gradually differentiate into strains expressing either ShadowG or mScarlet. **d**. Representative flow cytometry data show that some cells in the original circuit (**a**) exhibit unintended red fluorescence. **e**. Flow cytometry data show that incorporating terminators into the circuit (**b**) successfully reduces cells with unintended red fluorescence. Experiments were performed in 6 replicates.

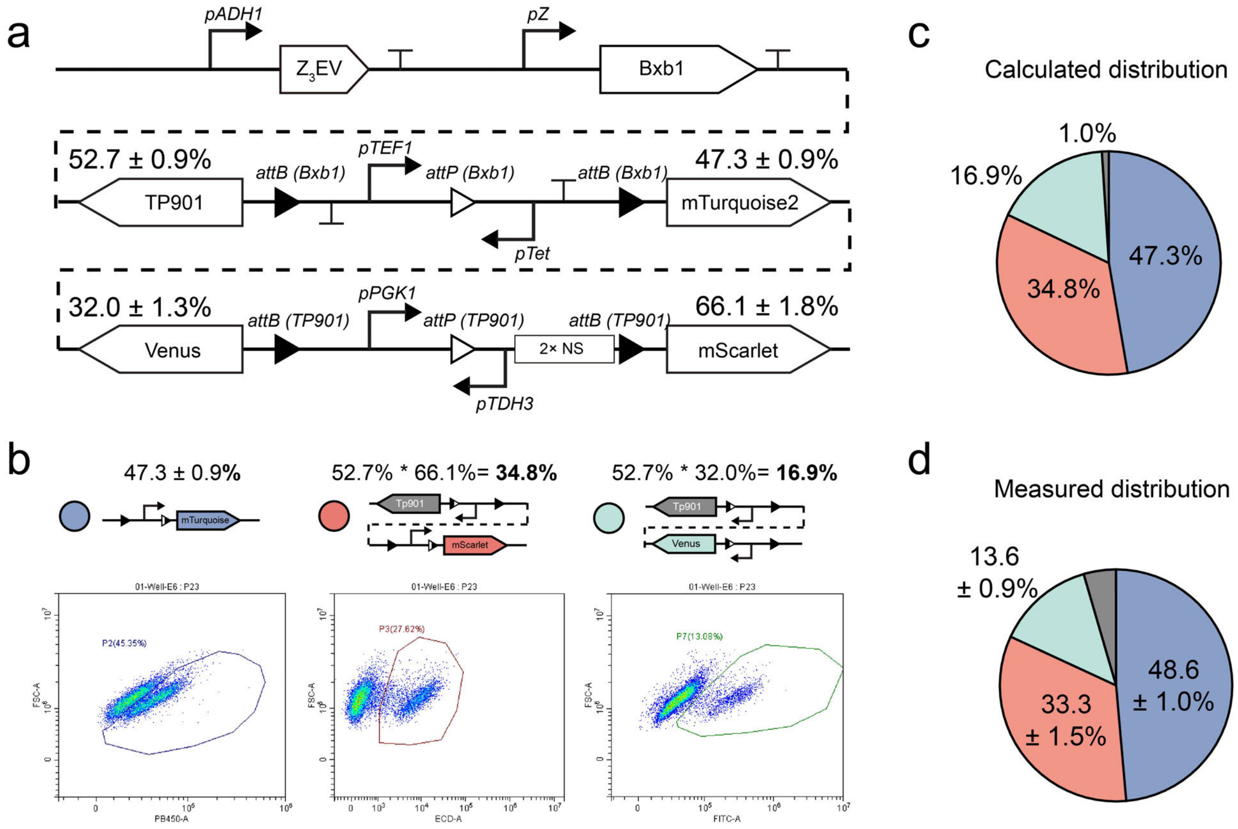

**Fig. S66 | Characterization of a representative cascading branching device. a.** Schematic of the 2-layer branching circuit design. **b**. Top: The theoretical proportions of progeny cell outcomes were obtained by multiplying the expression probabilities of the relevant genes within each circuit. Bottom: Representative flow cytometry measurements. **c.** Calculated distribution of progeny cell proportions. The 1.0% of non-fluorescent cells is likely due to the inefficient catalysis of the Tp901-recognized circuit, as shown in **Fig. S37**. **d.** Experimental measurements of progeny proportions align well with theoretical predictions. Data are mean ± s.d. of *n* = 8 replicates.

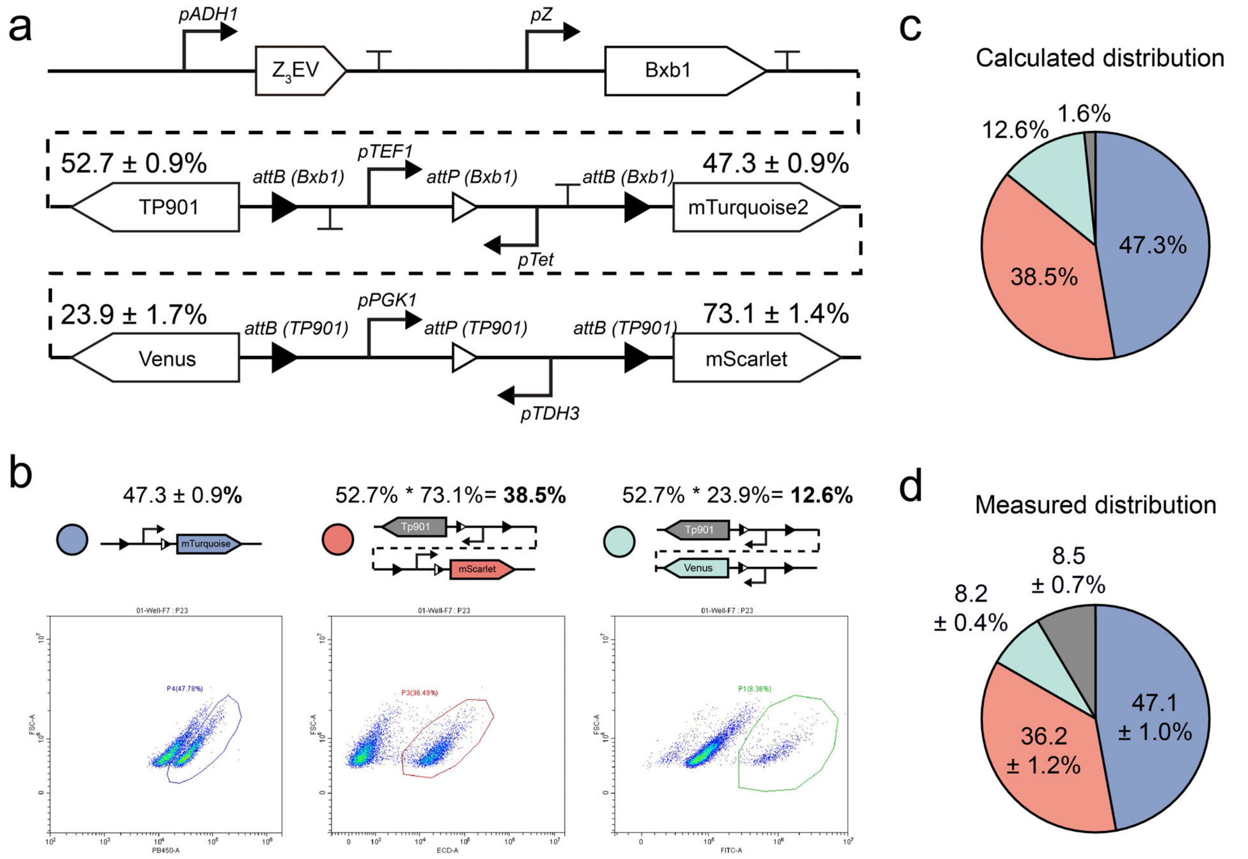

**Fig. S67 | Characterization of a representative cascading branching device. a.** Schematic of the 2-layer branching circuit design. **b**. Top: The theoretical proportions of progeny cell outcomes were obtained by multiplying the expression probabilities of the relevant genes within each circuit. Bottom: Representative flow cytometry measurements. **c**. Calculated distribution of progeny cell proportions. **d**. Experimental measurements of progeny proportions. Data are mean ± s.d. of *n* = 8 replicates.

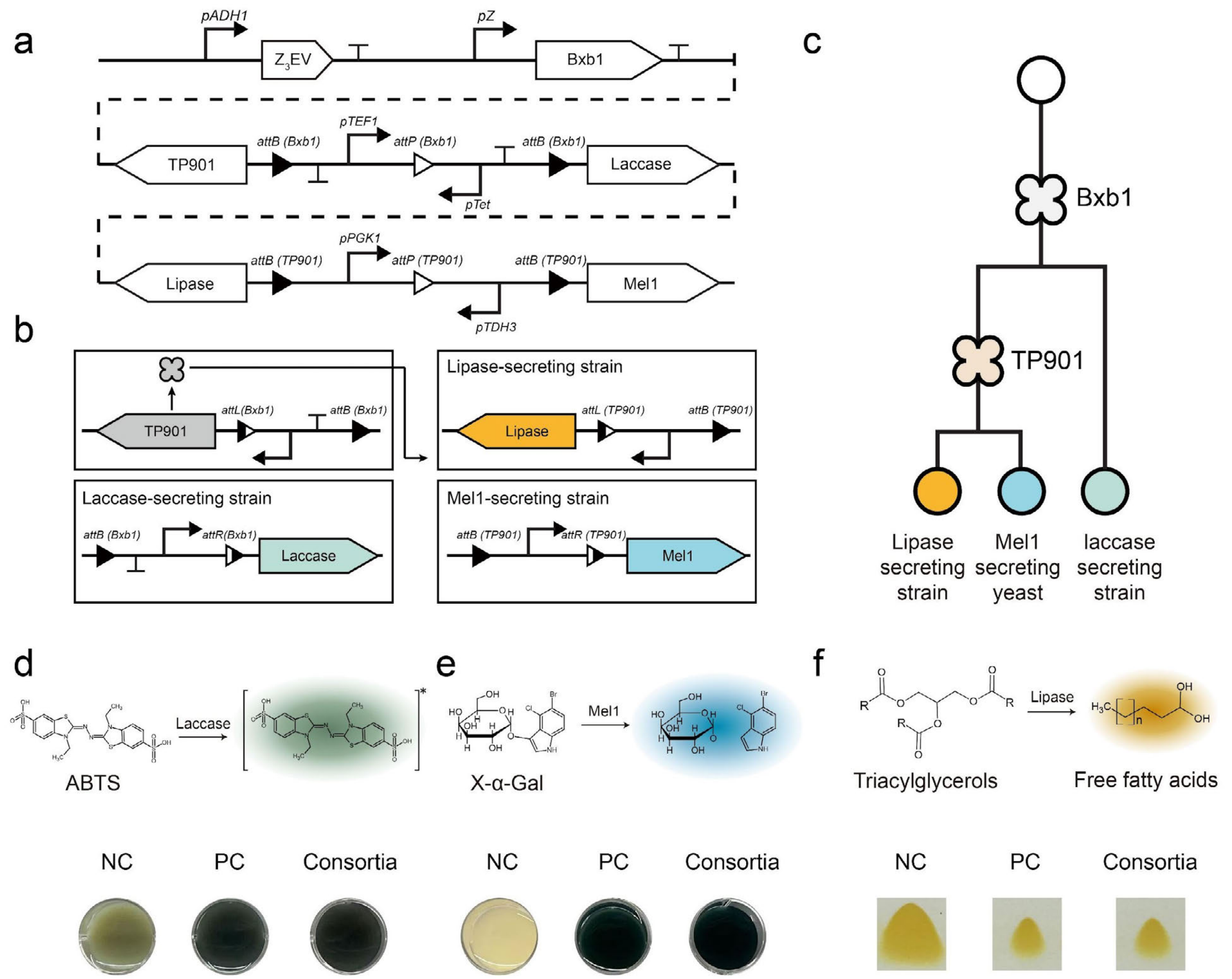

**Fig. S68 | Generating multifunctional microbial consortia using the designed cascading branching device**. **a**. Gene circuit design for a synthetic microbial consortium capable of degrading multiple pollutants. **b**. The first gene circuit, catalyzed by the Bxb1 recombinase, produces progeny yeast that express TP901 and secrete laccase. The yeast expressing TP901 further catalyzes the recombination of a second gene circuit, resulting in progeny that secrete lipase and Mel1. **c**. Schematic showing a single type of cell undergoing successive differentiation processes to produce a yeast consortium containing three enzyme-secreting strains. **d**. Degradation of ABTS by the yeast consortia. 'NC' represents the negative control (wild-type BY4741 yeast without laccase secretion), and 'PC' represents the positive control (engineered yeast constitutively secreting laccase). **e**. Degradation of X-α-Gal by the yeast consortia. **f**. Transformation of triacylglycerols into free fatty acids by the yeast consortia. Experiments were performed in triplicate.

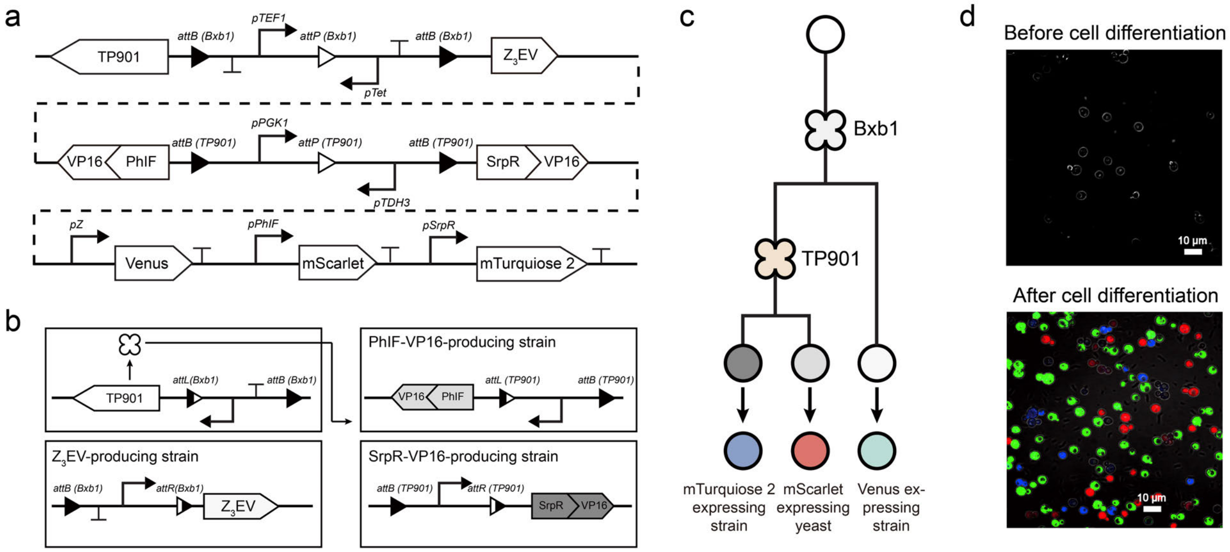

**Fig. S69 | Incorporating transcription factors (TFs) into a designed cascading branching device for precise cell fate regulation. a.** Design of a cascading branching device incorporating three TFs: $Z_3$EV, PhIF-VP16, and SrpR-VP16. **b.** The initial gene circuit, driven by Bxb1 recombinase, generates progeny cells expressing TP901 and $Z_3$EV. Yeast expressing TP901 then initiates recombination of a second gene circuit, producing two distinct progeny types that express proteins PhIF-VP16 and SrpR-VP16, respectively. A third circuit utilizes these proteins to activate a cassette containing Venus, mScarlet, and mTurquoise 2, controlled by specific promoters (*pZ, pPhIF, pSrpR*), activated by $Z_3$EV, PhIF-VP16, and SrpR-VP16. **c.** Schematic illustration showing sequential differentiation in a single cell, leading to the production of three fluorescent yeast cell types. **d.** Fluorescence microscopy images before and after cellular differentiation, highlighting the transitions driven by the gene circuits.

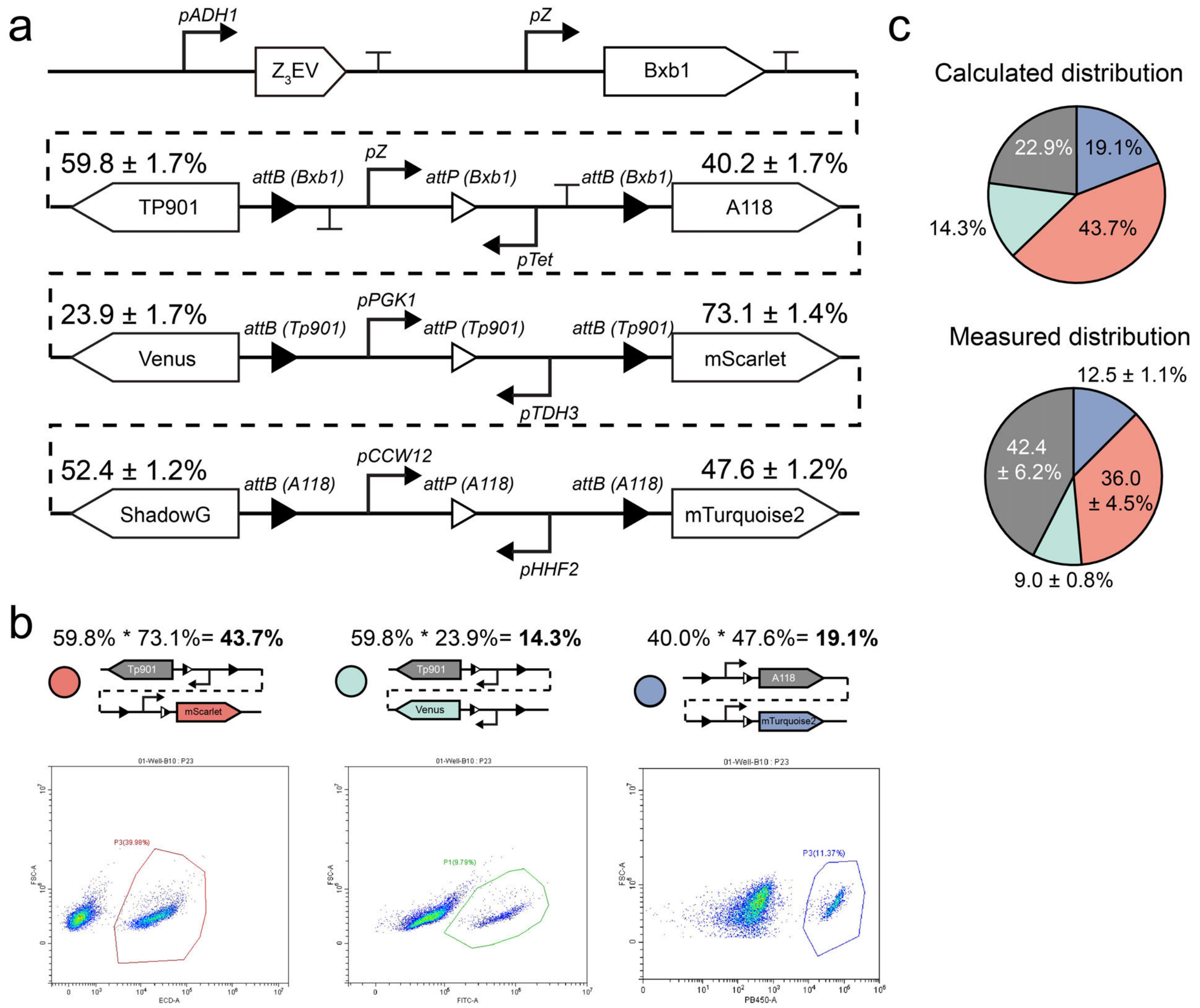

**Fig. S70 | Design and characterization of a 2-layer branching device recognized by three recombinases. a.** Schematic of the 2-layer branching circuit design. **b**. Theoretical probabilities of progeny cell outcomes and representative flow cytometry data of each progeny cell type. **c**. Comparison of theoretical and experimentally measured progeny cell proportions. Asymmetric differentiation due to inefficient catalysis of A118 in a population of undifferentiated (non-fluorescent) cells (**Fig. S37b**). This deviation is reflected in the distribution of blue and non-fluorescent cells. Data are mean ± s.d. of *n* = 8 replicates.

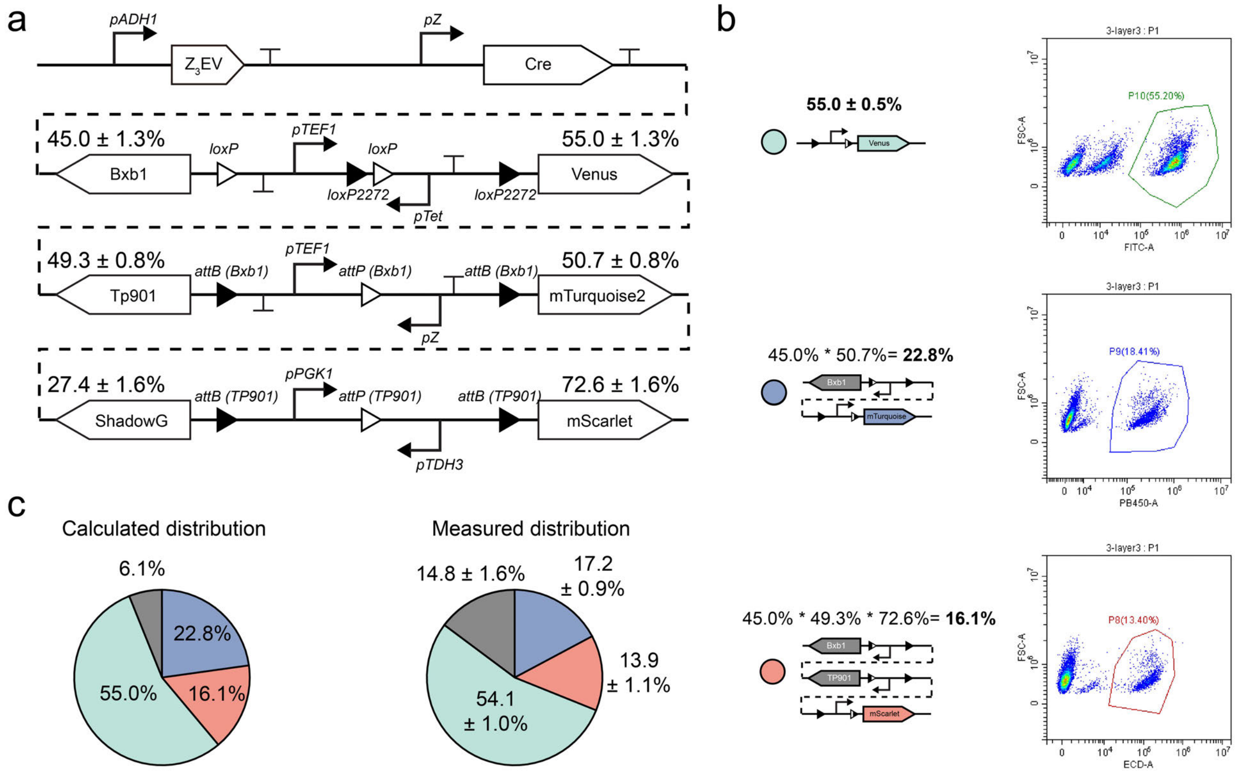

**Fig. S71 | Design and characterization of a 3-layer branching device recognized by three recombinases. a.** Design of a 3-layer sequential gene differentiation circuit. In the first layer, catalyzed by Cre recombinase, there is a 45.0 ± 1.3% probability of producing Bxb1-expressing strains and a 55.0 ± 1.3% chance of generating Venus-expressing yeast. In the second layer, catalyzed by Bxb1, there is a 49.3 ± 0.8% probability of producing Tp901-expressing progeny and a 50.7 ± 0.8% probability of generating mTurquoise 2-expressing strains. The third layer, catalyzed by Tp901, results in a 27.4 ± 1.6% probability of producing ShadowG-expressing strains and a 72.6 ± 1.6% chance of generating mScarlet-expressing yeast. **b**. Theoretical progeny distribution and representative flow cytometry data for each progeny cell type. **c**. Comparison of theoretical distribution and experimentally measured progeny distributions. Data are mean ± s.d. of $n$ = 8 replicates.

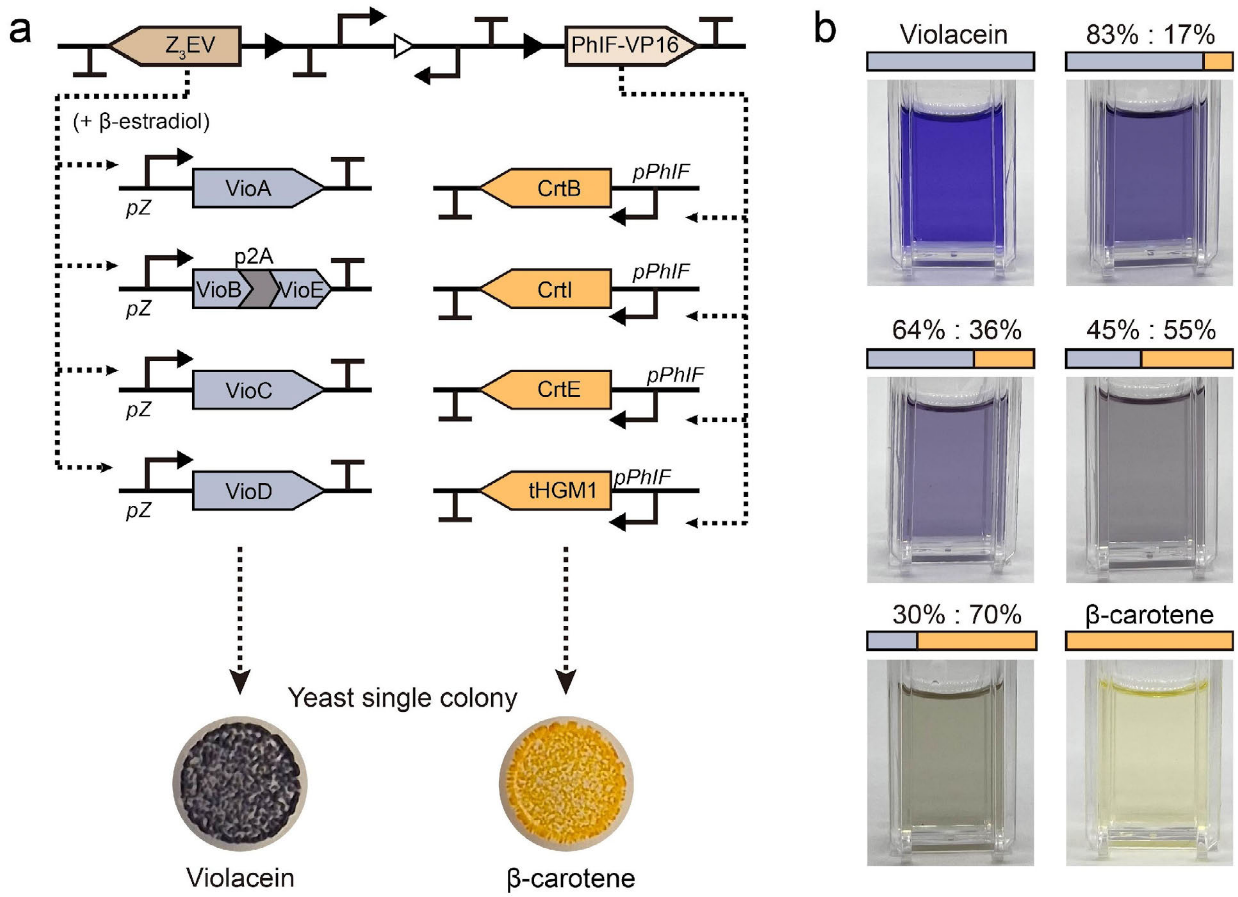

**Fig. S72 | Genetic circuits for violacein and β-carotene synthesis a.** Violacein synthesis, driven by the pZ promoter, integrates five enzymes (VioA, VioB, VioC, VioD, and VioE), into four yeast loci. Induction by $Z_3EV$ and β-estradiol triggers violacein production. β-carotene synthesis circuits integrate four enzymes (CrtE, CrtI, CrtB, and tHGM1) into four different loci, driven by *pPhlF* promoters. PhlF-VP16 production initiates β-carotene synthesis, producing yellow-to-orange pigments. **b.** Photograph of pigments extracted from yeast consortia using DMSO. Proportions of daughter cells were quantified by colony counting on agar plates after Bxb1 and β-estradiol induction. Genetic constructions are listed in **Fig. S73** and **Table. S7**. Experiments were conducted in triplicate.

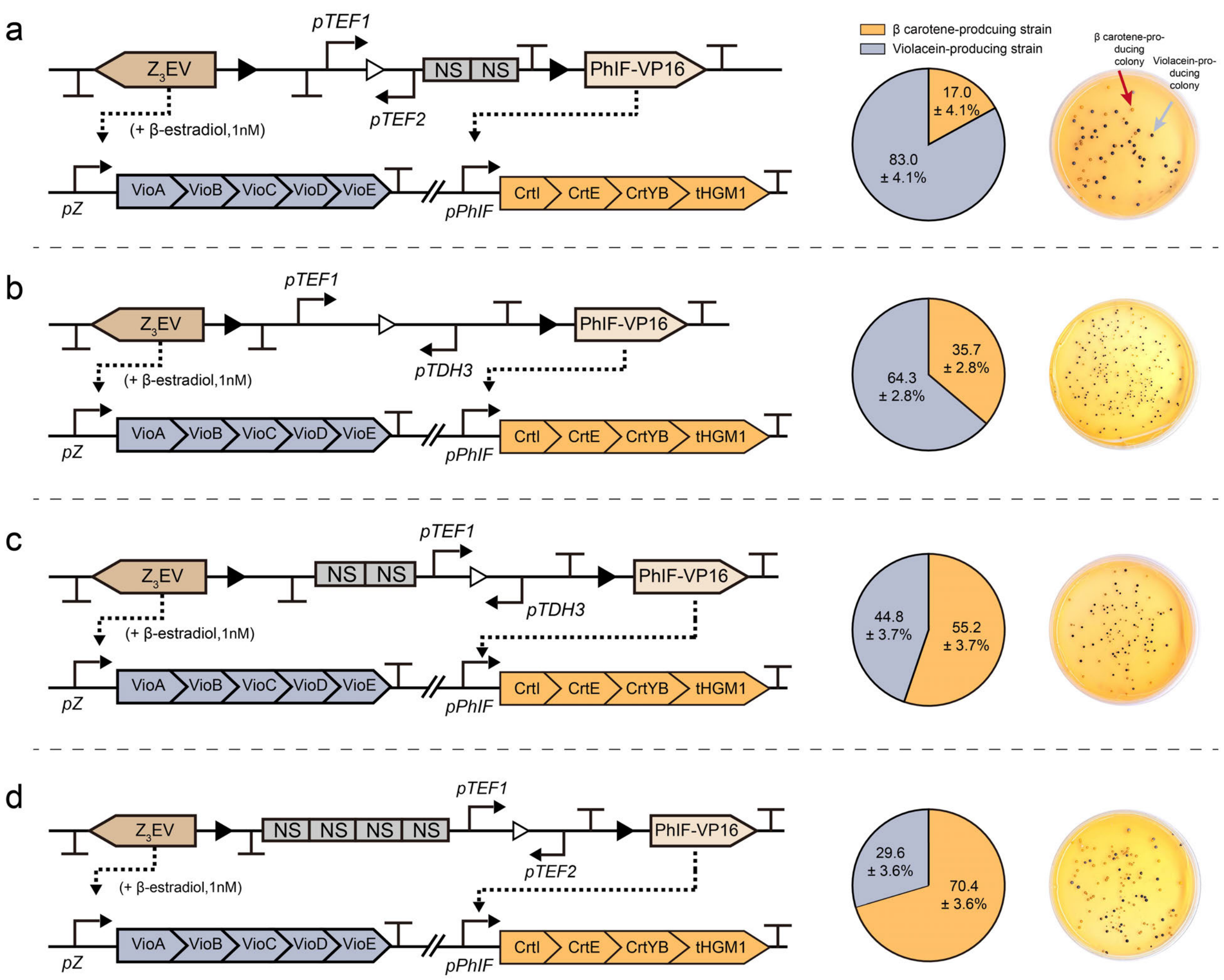

**Fig. S73 | Representative circuit configurations and quantification of differentiated yeast colonies producing violacein or β-carotene. a-d**, Schematic representations (left) of four synthetic gene circuits enabling binary fate branching in yeast. Each circuit integrates mutually exclusive biosynthetic modules encoding violacein (purple) or β-carotene (orange), controlled via recombinase-mediated DNA rearrangement. Circuit topologies differ in promoter identity, orientation, and insulator arrangement. Middle, Quantification of differentiated progeny based on colony pigmentation. Pie charts indicate the proportion of violacein- and β-carotene-producing colonies (data are mean ± s.d. of *n* = 3 replicates). Right, Representative colony images following induction with aTc (1 mM) and β-estradiol (1 nM).

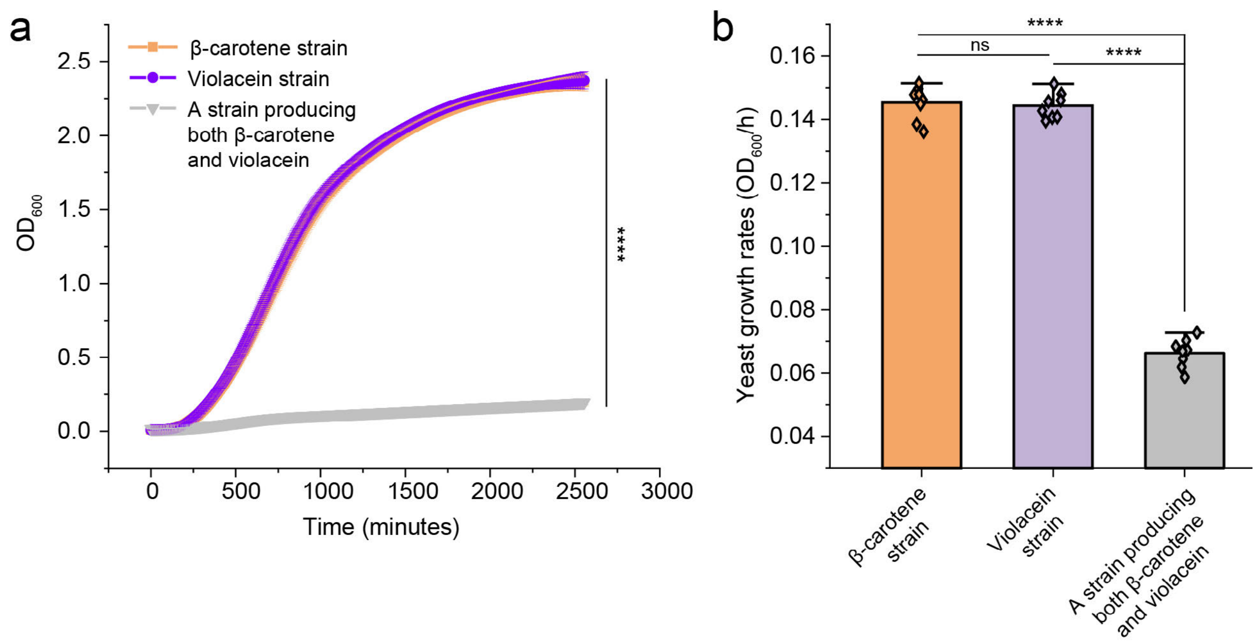

**Fig. S74 | Division of labor reduces metabolic burden in pigment-producing yeast strains.** **a**, Growth curves of three engineered yeast strains: a strain producing β-carotene only (orange), a strain producing violacein only (purple), and a strain co-producing both β-carotene and violacein (gray). The co-producing strain exhibited severely impaired growth compared to the single-pathway strains, indicating increased metabolic burden. **b**, Quantified growth rates ($OD_{600}$/hour) of the three strains. The β-carotene and violacein strains showed comparable growth rates (ns, not significant), while the co-producing strain exhibited a significantly lower growth rate (**** $p < 0.0001$, statistical analysis was performed using unpaired two-tailed Student's *t*-tests). Data are mean ± s.d. of $n = 8$ replicates.

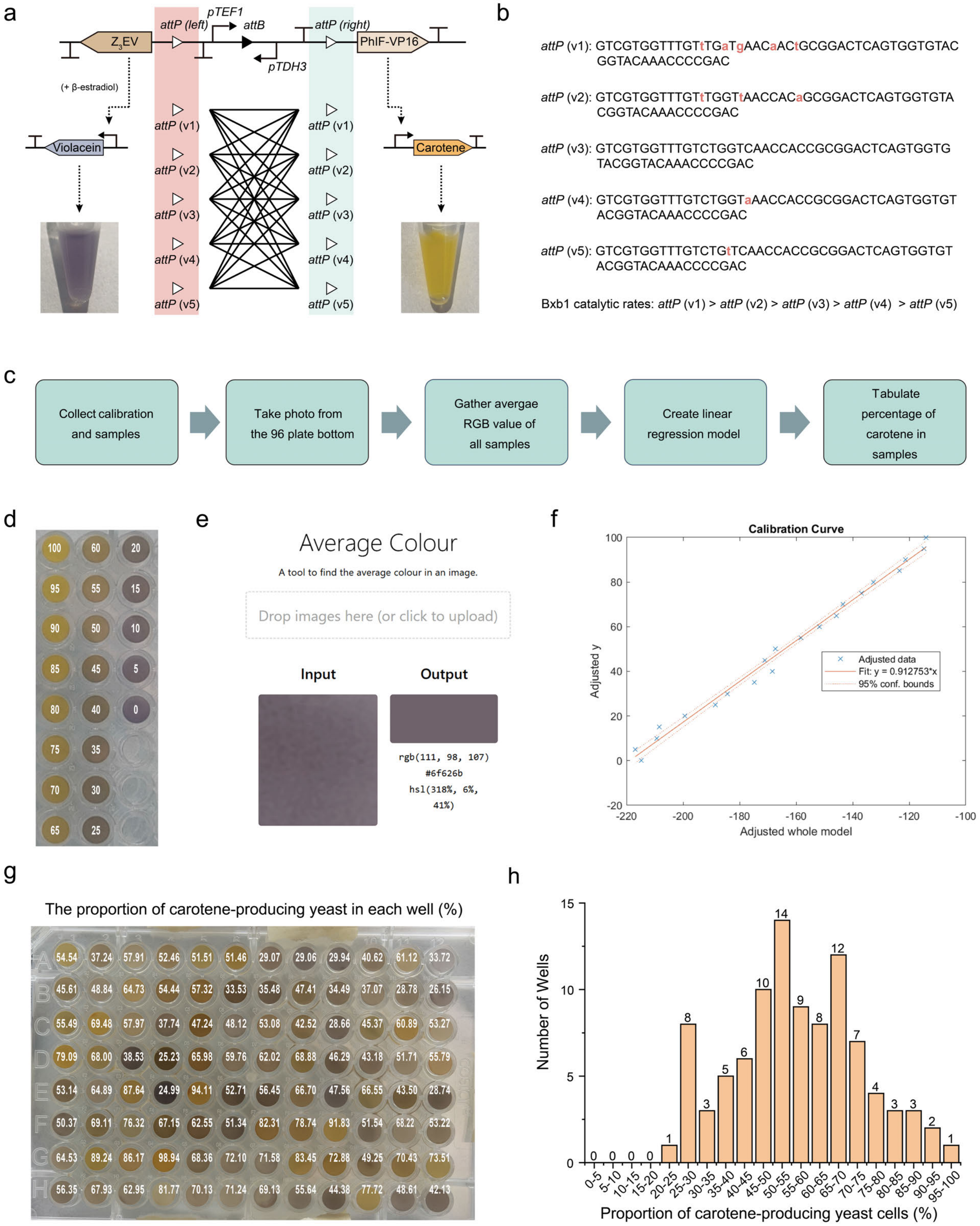

**Fig. S75 | Programmable pigment output profiling using recombination-guided yeast libraries and RGB image analysis. a**, Schematic of a recombinase-based circuit library designed to produce tunable ratios of violacein and carotene by using 5 *attP* variants with different recombination efficiencies. **b**, DNA sequences of selected *attP* variants (v1-v5) and their ranked recombination efficiencies with Bxb1 recombinase. **c**, Workflow for image-based quantification of pigment composition using color calibration and linear regression. **d**, Calibration plate containing 21 pigment mixtures with defined carotene-to-violacein ratios (from 0% to 100% carotene in 5% increments). **e**, Screenshot of the online tool (https://avg-colour.netlify.app/) used to extract average RGB values from well images. **f**, Calibration curve generated by MATLAB using linear regression (fit linear model) on RGB data from standard mixtures. The model demonstrated strong predictive performance ($R^2 = 0.9908$). **g**, Predicted proportion of carotene-producing cells in each well of the yeast differentiation library, visualized in a 96-well format. **h**, Histogram summarizing the distribution of carotene-producing cell ratios across the 96-well library. Each bar represents the number of wells falling into a given ratio bin.

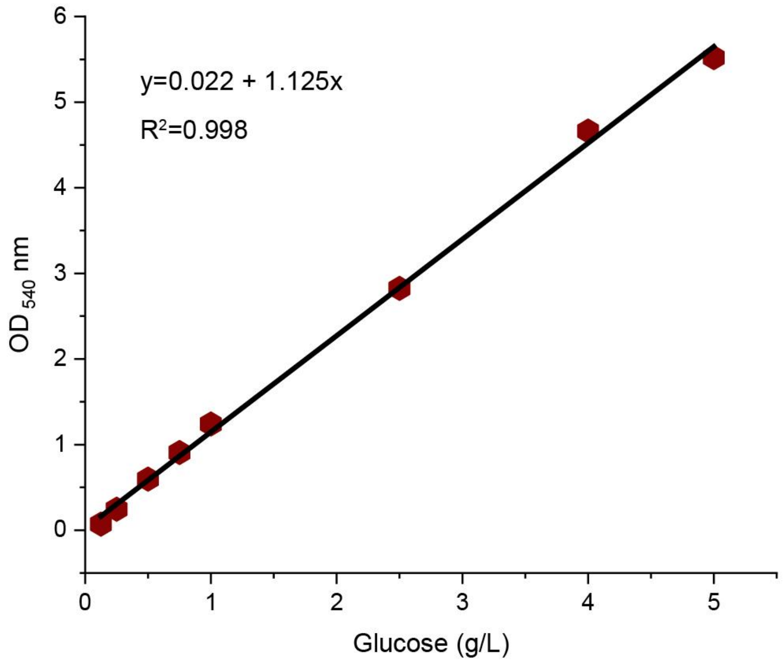

**Fig. S76 | Standard curve for glucose concentration measured by the DNS method.**

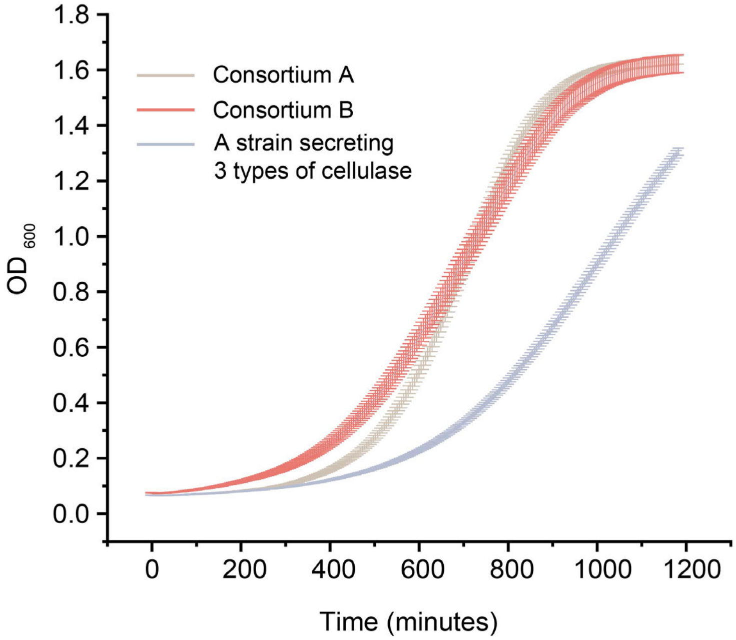

**Fig. S77 | Growth curves for cellulase-secreting yeast consortia.** Growth curves of three cellulase-secreting yeast systems. Growth rates are Consortium A (≈0.23 $OD_{600}$/h), Consortium B (≈0.24 $OD_{600}$/h), and the strain secreting all three cellulases (≈0.17 $OD_{600}$/h).

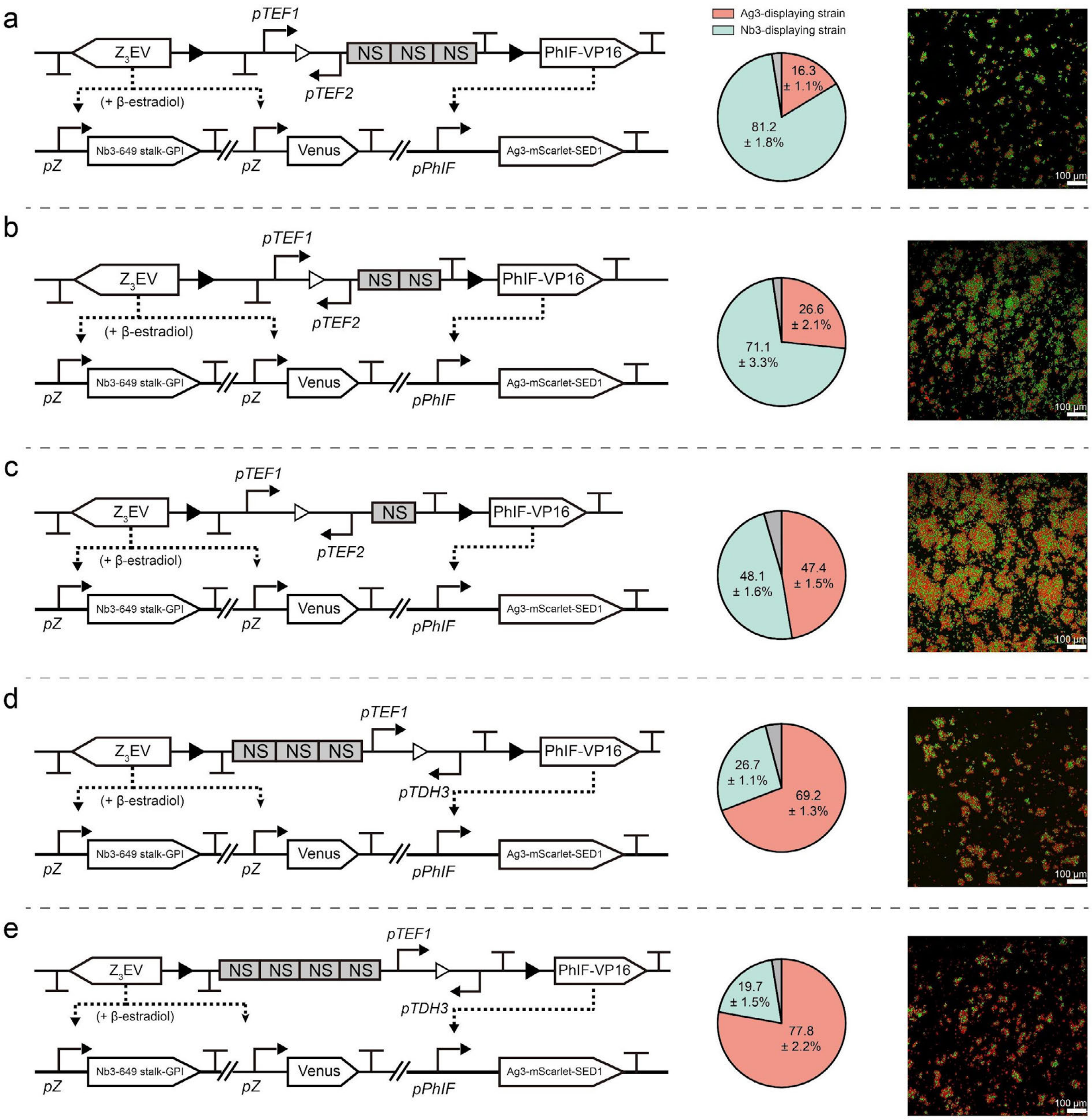

**Fig. S78 | Progeny cell distribution and fluorescence images of cellular morphogens. a-e.** Ratios of progeny cells (Nb3-expressing yeasts and Ag3-displaying strains) were adjusted to approximately 1:4 to 4:1. Data are mean ± s.d. of *n* = 8 replicates

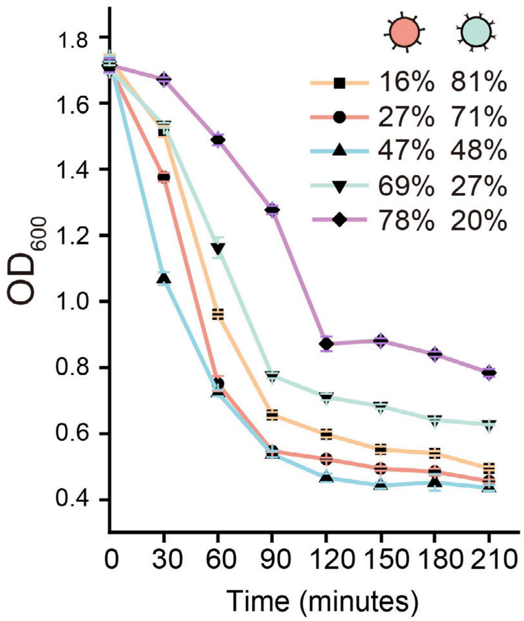

**Fig. S79 | Settling speeds of consortia with varying ratios of Nb3- and Ag3-displaying cells (data were mean ± s.d. of *n* = 3 replicates).**

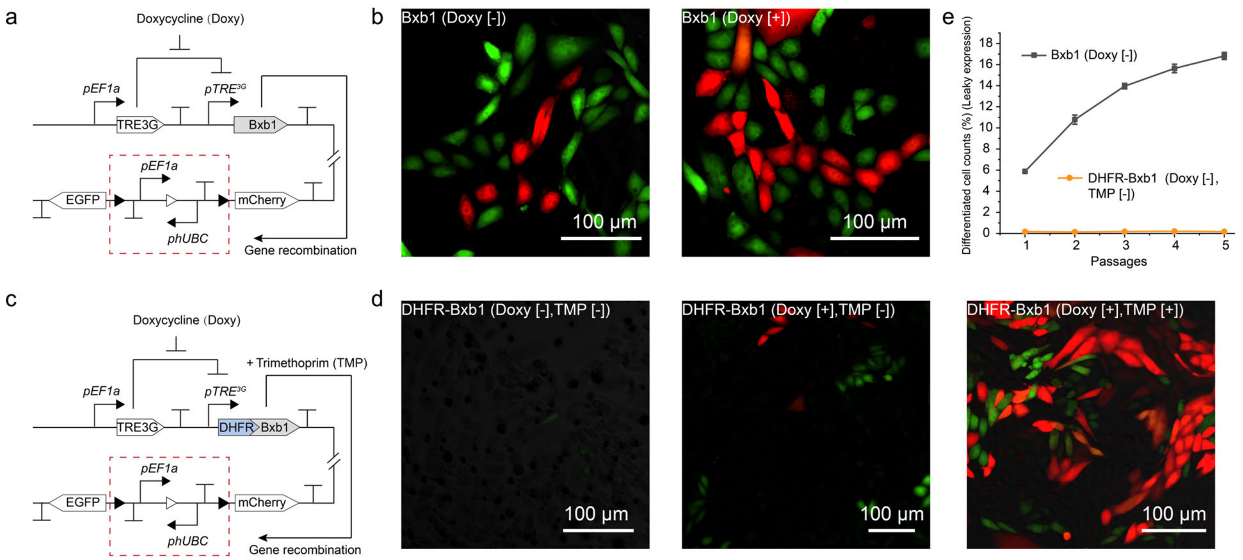

**Fig. S80 | Mitigating recombinase leakiness in mammalian cells using a DHFR-based degradation strategy. a**, Schematic of the doxycycline (Doxy)-inducible Bxb1 circuit in CHO-K1 cells. Bxb1 is expressed under the control of a *pTRE*$^{3G}$ promoter regulated by Doxy. **b,** Fluorescence microscopy images showing cellular differentiation under uninduced (Dox [-]) and induced (Dox [+]) conditions. Notably, spontaneous differentiation is observed even in the absence of Dox, indicating leaky recombinase expression. **c**, Modified genetic circuit in which Bxb1 is fused to a destabilizing DHFR (Dihydrofolate reductase) degron. In the absence of the stabilizing ligand trimethoprim (TMP), Bxb1 is rapidly degraded; TMP addition allows controlled stabilization and activation. **d**, Fluorescence microscopy images of CHO-K1 cells expressing DHFR-Bxb1 under different conditions. Minimal background fluorescence is observed in uninduced cells (Dox [-], TMP [-]), with a small number of differentiated cells upon Dox induction alone (Dox [+], TMP [-]). Robust and uniform differentiation is restored upon combined Dox and TMP induction (Dox [+], TMP [+]). **e**, Quantification of leaky differentiation over five serial passages. Wild-type Bxb1 shows a gradual increase in differentiated cells under uninduced conditions (gray line), whereas DHFR-Bxb1 effectively suppresses background recombination (orange line). Data are mean ± s.d. of $n = 3$ replicates.

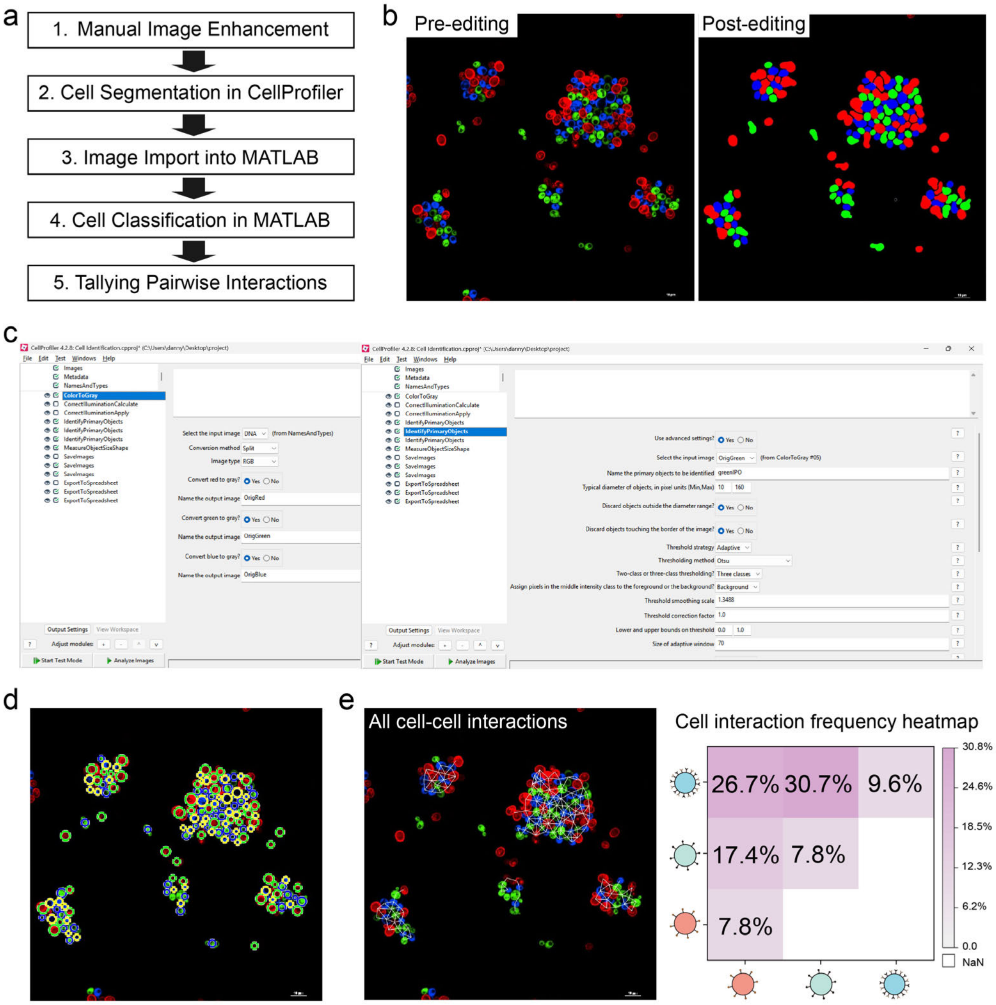

**Fig. S81 | Image-processing workflow for quantifying cell-cell interaction frequencies in engineered living systems. a,** Overview of a five-step computational pipeline used to quantify cell-cell interactions from fluorescence images of yeast populations: (1) manual image enhancement to resolve touching cells; (2) segmentation using CellProfiler; (3) image import into MATLAB; (4) cell classification based on RGB intensity; and (5) pairwise interaction analysis based on spatial proximity. **b**, Representative fluorescence images before and after manual editing. Post-editing improves boundary clarity between adjacent or dividing yeast cells to enable accurate segmentation and downstream analysis. **c**, Example CellProfiler interface illustrating the modules used for image processing. Images were separated by color channel and analyzed using the "Identify Primary Objects" module with typical object diameter settings (10-160 pixels) and adaptive thresholding (Otsu, 3-class mode). **d**, Representative segmentation result generated in MATLAB. Individual yeast cells are identified, classified by fluorescence profile, and outlined with corresponding color masks based on their dominant RGB intensity values. **e**, left: Representative map of all detected cell-cell interactions, visualized as connecting lines between neighboring cells located within a 35-pixel threshold. Right: Corresponding interaction frequency heatmap generated from this dataset. Each matrix element represents the normalized frequency of contact between each pair of cell types, allowing visualization of preferential self- or heterotypic interactions.

**Table. S1** | The predicted $k_A^0$ and $k_B^0$ for various *att* site mutations.

| Scheme of genetic circuit | Measured Venus cell fraction | $k_A^0$ | $k_B^0$ | Predicted Venus cell fraction |
|---|---|---|---|---|
| 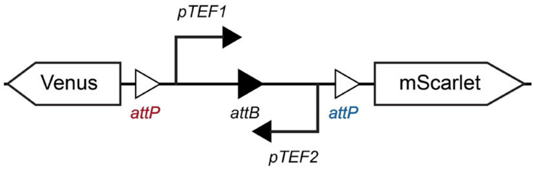 | 36.7 ± 0.6% | 0.38 | 0.14 | 37.8% |
| 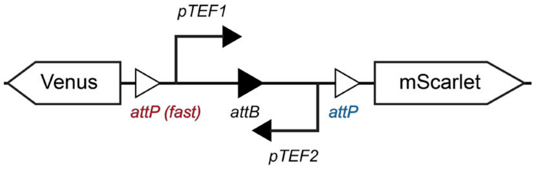 | 44.3 ± 1.6% | 0.50 | 0.14 | 44.4% |
| 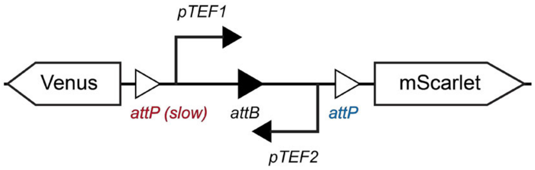 | 19.7 ± 0.5% | 0.15 | 0.14 | 19.7% |
| 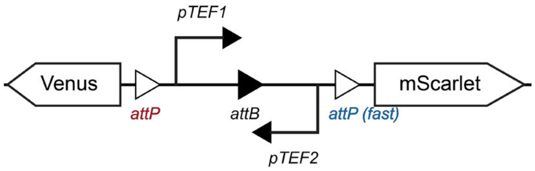 | 28.6 ± 2.1% | 0.38 | 0.43 | 28.6% |
| 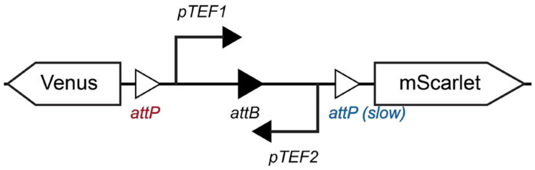 | 61.9 ± 1.0% | 0.38 | 0.03 | 58.7% |

**Table. S2** | Predicted $r_A$ and $r_B$ values for different promoter types.

| Scheme of genetic circuit | Measured Venus cell fraction | $k_A^0$ | $k_B^0$ | $r_A$ | $r_B$ | Predicted Venus cell fraction |
|---|---|---|---|---|---|---|
| 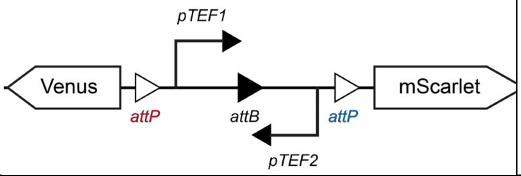 | 36.7 ± 0.6% | 0.38 | 0.14 | 0.56 | 0.01 | 37.8% |
| 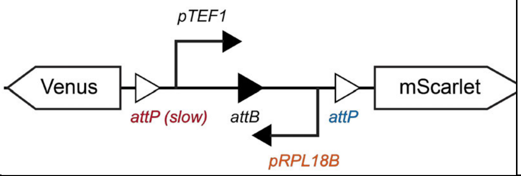 | 68.3 ± 1.2% | 0.15 | 0.14 | 0.56 | 1.37 | 68.2% |
| 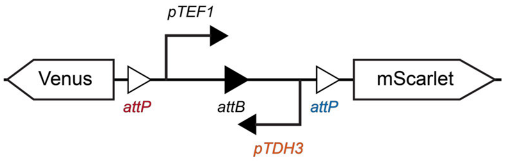 | 56.6 ± 0.6% | 0.38 | 0.14 | 0.56 | 0.14 | 44.5% |

**Table. S3** | Mathematical model predictions of green-to-red progeny cell ratios.

| Scheme of Genetic Circuit | Measured Venus cell fraction | $k_A^0$ | $k_B^0$ | $r_A$ | $r_B$ | Predicted Venus cell fraction |
|---|---|---|---|---|---|---|
| 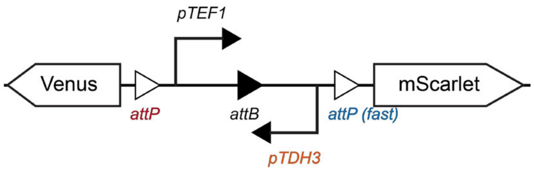 | 45.9 ± 0.7% | 0.38 | 0.25 | 0.56 | 0.14 | 39.7% |
| 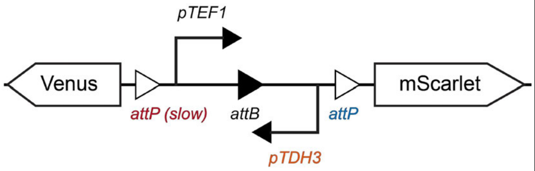 | 34.8 ± 1.2% | 0.15 | 0.14 | 0.56 | 0.14 | 30.1% |
| 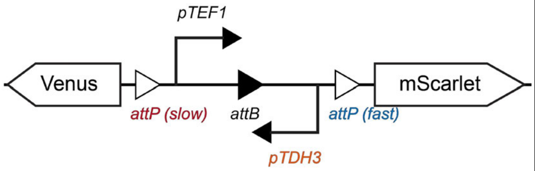 | 23.9 ± 0.7% | 0.15 | 0.25 | 0.56 | 0.14 | 26.2% |
| 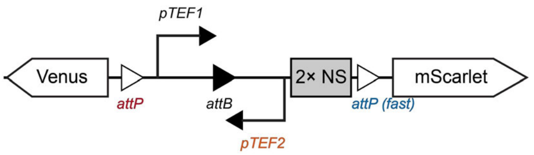 | 53.6 ± 1.9% | 0.38 | 0.25 | 0.56 | 0.01 | 47.2% |
| 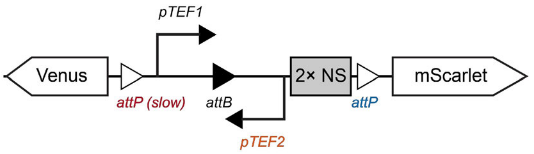 | 39.8 ± 0.8% | 0.15 | 0.14 | 0.56 | 0.01 | 34.5% |
| 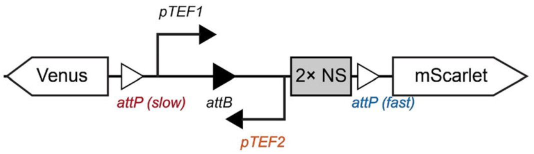 | 33.2 ± 0.7% | 0.15 | 0.25 | 0.56 | 0.01 | 26.5% |
| 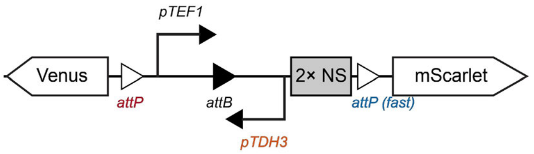 | 62.9 ± 1.3% | 0.38 | 0.25 | 0.56 | 0.14 | 54.0% |
| 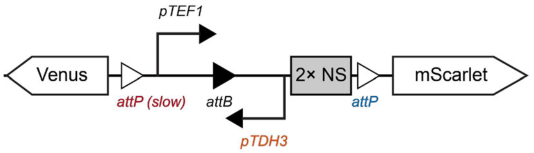 | 45.4 ± 1.0% | 0.15 | 0.14 | 0.56 | 0.14 | 47.5% |

| | | | | | | |
|---|---|---|---|---|---|---|
| 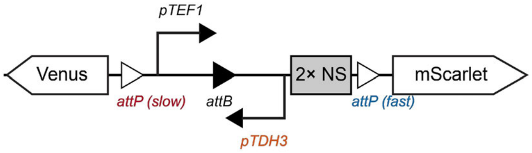 | 39.9 ± 0.9% | 0.1 5 | 0.25 | 0.56 | 0.14 | 38.4% |

**Table. S4** | DNA sequences of genetic parts used in this study.

| Type | Genetic parts | Description |
|---|---|---|
| Promoter | *pTEF1* | Yeast constitutive promoters |
| | *pTEF2* | |
| | *pALD6* | |
| | *pHHF2* | |
| | *pPGK1* | |
| | *pREV1* | |
| | *pRPL18B* | |
| | *pSac6* | |
| | *pTDH3* | |
| | *pCCW12* | |
| | *pPOP6* | |
| | *pADH1* | |
| | *pRAD27* | |
| | *pTet* | Yeast promoter induced by aTc |
| | *pZ* | Yeast promoter induced by β-estradiol |
| | *pLac* | Yeast promoter impressed by LacI |
| | *pSrpR* | Yeast promoter induced by SrpR-VP16 |
| | *pPhlF* | Yeast promoter induced by PhlF-VP16 |
| | *pJ72163* | *E. coli* constitutive promoters |
| | *pProD* | |
| | *pEF1* | HEK293FT/CHO-K1 mammalian cells' constitutive promoters |
| | *phUBC* | |
| | *pPGK* | |
| | *pSV40* | |
| | *pCMV* | |
| | *pTRE$^{3G}$* | HEK293FT/CHO-K1 mammalian cells' inducible promoters |
| | *p5UAS* | |
| *att* sites | *attB* (Bxb1_GA) | *att* sites for Bxb1 |
| | *attP* (Bxb1_GA) | |
| | *attB* (TP901) | *att* sites for TP901 |
| | *attP* (TP901) | |
| | *attB* (A118) | *att* sites for A118 |
| | *attP* (A118) | |
| | *attB* (R4) | *att* sites for R4 |
| | *attP* (R4) | |
| | *Cre* (loxP) | *att* sites for Cre |
| | *Cre* (loxP2272) | |

| | | |
|---|---|---|
| | *attB* (PhiC31) | *att* sites for PhiC31 |
| | *attP* (PhiC31) | |
| | *attP* (Bxb1_fast) | Bxb1 *attP* sits with predictive recombination rates |
| | *attP* (Bxb1_slow) | |
| Non-sense DNA sequence | 1× NS | Non-sense sequence (773 bp) |
| | 2× NS | Non-sense sequence (2 × 773 bp) |
| | 3× NS | Non-sense sequence (3 × 773 bp) |
| | 4× NS | Non-sense sequence (4 × 773 bp) |
| | 9× NS | Non-sense sequence (9 × 773 bp) |
| | S1 | Designed 100 bp sequence (50% GC contents) for NGS sequencing |
| | S2 | Designed 100 bp sequence (50% GC contents) for NGS sequencing |
| | S3 | Designed 100 bp sequence (30% GC contents) for NGS sequencing |
| | S4 | Designed 100 bp sequence (30% GC contents) for NGS sequencing |
| | S5 | Designed 100 bp sequence (30% GC contents) for NGS sequencing |
| | S6 | Designed 100 bp sequence (70% GC contents) for NGS sequencing |
| | S7 | Designed 100 bp sequence (70% GC contents) for NGS sequencing |
| | S8 | Designed 100 bp sequence (70% GC contents) for NGS sequencing |
| | S9/S10 | Designed 100 bp sequence with 3 TetO inserted for NGS sequencing |

**Table. S5** | Sequences of proteins in this study.

| Type | Protein | Description |
|---|---|---|
| Fluorescent proteins | Venus | Green fluorescent protein |
| | EGFP | Green fluorescent protein |
| | mScarlet | Red fluorescent protein |
| | mCherry | Red fluorescent protein |
| | mTurquoise2 | Blue fluorescent protein |
| | tagBFP | Blue fluorescent protein |
| | Crimson | NIR light-activated fluorescent protein |
| | iRFP680 | NIR light-activated fluorescent protein |
| | ShadowG | GFP mutant without fluorescence |
| Recombinases | Bxb1 | Serine recombinase |
| | Si74 | Serine recombinase |
| | PhiC31 | Serine recombinase |

| | | |
|---|---|---|
| | R4 | Serine recombinase |
| | Tp901 | Serine recombinase |
| | Cre-EBD | Tyrosine recombinase |
| | A118 | Serine recombinase |
| | No67 | Serine recombinase |
| Transcription factors (TFs) | $Z_3$EV | TF for *pZ* promoter |
| | LacI | TF for *pLac* promoter |
| | SrpR-VP16 | TF for *pSrpR* promoter |
| | PhlF-VP16 | TF for *pPhlF* promoter |
| | TetR-Mxi1 | TF for *pTet* promoter |
| | rtTA-GAL4 | TF for *pTet* promoter |
| | GAL4-VP64 | TF for *p5UAS* promoter |
| | Tet-On® $^{3G}$ | TF for *pTRE*$^{3G}$ promoter |
| Enzyme | Lipase | Lipase catalyzes the hydrolysis of fats (lipids) into glycerol and free fatty acids |
| | Mel1 | α-galactosidase hydrolyzes α-galactosidic |
| | Laccase | Laccase catalyzes the oxidation of phenolic compounds |
| | VioA | Violacein synthesis-related enzyme |
| | VioB | Violacein synthesis-related enzyme |
| | VioC | Violacein synthesis-related enzyme |
| | VioD | Violacein synthesis-related enzyme |
| | VioE | Violacein synthesis-related enzyme |
| | CrtI | Carotene synthesis-related enzyme |
| | CrtE | Carotene synthesis-related enzyme |
| | CrtYB | Carotene synthesis-related enzyme |
| | tHGM1 | Carotene synthesis-related enzyme |
| | BGL1 | β-glucosidase that catalyzes the hydrolysis of glycosidic bonds in β-glucosides |
| | EG2 | Endoglucanase that breaks down the internal β-1,4-glycosidic bonds in cellulose |
| | CBH2 | Cellobiohydrolase 2 catalyzes the hydrolysis of cellulose by cleaving off cellobiose units from the non-reducing ends of cellulose chains |
| Degradation tags | UbiM | Protein degradation tags for yeast |
| | UbiY | |
| | UbiR | |
| | DHFR | Protein degradation tags for mammalian cells |
| Signal peptides | $SP_{sed1}$ | Yeast signal peptide |
| | $SP_{werid}$ | Yeast signal peptide |
| | $SP_{Mf\alpha}$ | Yeast signal peptide |
| | $SP_{a\ factor}$ | Yeast signal peptide |
| | $SP_{albumin}$ | Yeast signal peptide |
| | $SP_{amylase}$ | Yeast signal peptide |
| | $SP_{Ig\kappa\ leader}$ | Mammalian cell signal peptide |
| | $SP_{CD8\alpha}$ | Mammalian cell signal peptide |
| Anchor proteins | SED1 | Surface-anchoring protein for yeast display |
| | 649 stalk-GPI | Surface-anchoring protein for yeast display |

| | | |
|---|---|---|
| | ICAM-1 | Surface-anchoring protein for mammalian cell display |
| | PDGFRβ TM domain | Surface-anchoring protein for mammalian cell display |
| Antigen-antibody | Ag1 | Ag1 specifically interacts with Nb1 |
| | Ag2 | Ag2 specifically interacts with Nb2 |
| | Ag3 | Ag3 specifically interacts with Nb3 |
| | Nb1 | Nb1 specifically interacts with Ag1 |
| | Nb2 | Nb2 specifically interacts with Ag2 |
| | Nb3 | Nb3 specifically interacts with Ag3 |
| | Z17 | Z17 specifically interacts with Z18 |
| | Z18 | Z18 specifically interacts with Z17 |
| | Sg30 | Sg30 specifically interacts with Sg61 |
| | Sg61 | Sg61 specifically interacts with Sg30 |
| | Dockerin | Dockerin specifically interacts with cohesion |
| | Cohesion | Cohesion specifically interacts with dockerin |
| | Spycatcher | Spycatcher specifically interacts with Spytag |
| | Spytag | Spytag specifically interacts with Spycatcher |
| | LaG17 | LaG17 specifically interacts with EGFP |
| | LaG16 | LaG16 specifically interacts with EGFP |
| | LaM4 | LaM4 specifically interacts with mCherry |
| Antigen or antibody fused with anchor proteins for surface display | Ag3-mScarlet-SED1 | Protein that was displayed on the yeast surface |
| | Nb3-649 stalk-GPI | |
| | Ag1-649 stalk-GPI | |
| | Nb1-649 stalk-GPI | |
| | Dockerin -649 stalk-GPI | |
| | Cohesion -649 stalk-GPI | |
| | Spycatcher -649 stalk-GPI | |
| | Spytag -649 stalk-GPI | |
| | LaG17-Notch-Gal4 VP64 | Protein that was displayed on the CHO-K1 surface |
| | AntiCD19-Notch-Gal4 VP64 | |
| | CD19-Tm | |
| | LaG16-ICAM-1 | |
| | EGFP-ICAM-1 | |
| | LaM4-ICAM-1 | |
| | mCherry-ICAM-1 | |
| | EGFP-Tm | |

**Table. S6** | Background plasmids employed in this study. Links to annotated plasmid sequences are provided for all constructs.

| Plasmid | Construct details | Source |
|---|---|---|
| BAC | A bacterial artificial chromosome (BAC) vector | Roquet et al[3] |
| pYTK001 | Entry vector designed for cloning new DNA fragments via BsmBI Golden Gate reactions | Lee et al[4] |
| pYTK096 | Pre-assembled plasmid equipped with genetic elements for cloning in *E. coli* and subsequent integrative transformation into the ***URA3*** locus of *S. cerevisiae*. DNA fragments can be inserted into this plasmid using BsaI Golden Gate reactions. | |
| pYTK097 | Pre-assembled plasmid equipped with genetic elements for cloning in *E. coli* and subsequent integrative transformation into the ***LEU2*** locus of *S. cerevisiae*. DNA fragments can be inserted into this plasmid using BsaI Golden Gate reactions. | This study |
| pYTK098 | Pre-assembled plasmid equipped with genetic elements for cloning in *E. coli* and subsequent integrative transformation into the ***HO*** locus of *S. cerevisiae*. DNA fragments can be inserted into this plasmid using BsaI Golden Gate reactions. | |
| pYTK099 | Pre-assembled plasmid equipped with genetic elements for cloning in *E. coli* and subsequent integrative transformation into the ***HIS3*** locus of *S. cerevisiae*. DNA fragments can be inserted into this plasmid using BsaI Golden Gate reactions. | |
| pYTK100 | Pre-assembled plasmid equipped with genetic elements for cloning in *E. coli* and subsequent integrative transformation into the ***MET15*** locus of *S. cerevisiae*. DNA fragments can be inserted into this plasmid using BsaI Golden Gate reactions. | |
| pMYT095 | Pre-assembled plasmid equipped with genetic elements for cloning in *E. coli* and Cas9 protein for genome editing in *S. cerevisiae.* gRNA cassettes can be inserted into this plasmid using BsaI Golden Gate reactions. | Shaw et al[5] |
| pMYT076 | Pre-assembled plasmid equipped with genetic elements for cloning in *E. coli* and subsequent integrative transformation into *S. cerevisiae* genome (**Int.2 site**) with the assistance of Cas9. DNA fragments can be inserted into this plasmid using BsaI Golden Gate reactions. | |
| pMYT078 | Pre-assembled plasmid equipped with genetic elements for cloning in *E. coli* and subsequent integrative transformation into *S. cerevisiae* genome (**Int.4 site**) with the assistance of Cas9. DNA fragments can be inserted into this plasmid using BsaI Golden Gate reactions. | |
| pMYT079 | Pre-assembled plasmid equipped with genetic elements for cloning in *E. coli* and subsequent integrative transformation into *S. cerevisiae* genome (**Int.5** site) with the assistance of Cas9. DNA fragments can be inserted into this plasmid using BsaI Golden Gate reactions. | |
| pMYT080 | Pre-assembled plasmid equipped with genetic elements for cloning in *E. coli* and subsequent integrative transformation into *S. cerevisiae* genome (**Int.6** site) with the assistance of Cas9. DNA | |

| | |
|---|---|
| | fragments can be inserted into this plasmid using BsaI Golden Gate reactions. |
| pMYT081 | Pre-assembled plasmid equipped with genetic elements for cloning in *E. coli* and subsequent integrative transformation into *S. cerevisiae* genome (**Int.7** site) with the assistance of Cas9. DNA fragments can be inserted into this plasmid using BsaI Golden Gate reactions. |
| pMYT082 | Pre-assembled plasmid equipped with genetic elements for cloning in *E. coli* and subsequent integrative transformation into *S. cerevisiae* genome (**Int.8** site) with the assistance of Cas9. DNA fragments can be inserted into this plasmid using BsaI Golden Gate reactions. |
| pMYT083 | Pre-assembled plasmid equipped with genetic elements for cloning in *E. coli* and subsequent integrative transformation into *S. cerevisiae* genome (**Int.9** site) with the assistance of Cas9. DNA fragments can be inserted into this plasmid using BsaI Golden Gate reactions. |
| pMYT084 | Pre-assembled plasmid equipped with genetic elements for cloning in *E. coli* and subsequent integrative transformation into *S. cerevisiae* genome (**Int.10** site) with the assistance of Cas9. DNA fragments can be inserted into this plasmid using BsaI Golden Gate reactions. |

**Table. S7** | Representative plasmids used in *S. cerevisiae* in this study. Annotated plasmid sequences are available for all constructs.

| Plasmid | Construct details | Scheme |
|---|---|---|
| *pTet*-Bxb1 | Expression of Bxb1 induced by aTc. Construct was designed for propagation in *E. coli* and integration at the ***HIS3*** locus in *S. cerevisiae.* | 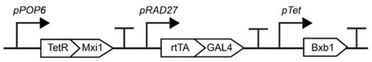 |
| *pZ*-Bxb1 | Expression of Bxb1 induced by β-estradiol. Construct was designed for propagation in *E. coli* and integration at the ***HIS3*** locus in *S. cerevisiae.* | 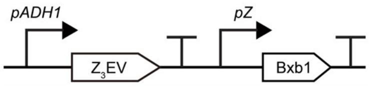 |
| pBranch 01 | Construct for controlled yeast cell differentiation. Designed for propagation in *E. coli* and integration at the ***URA3*** locus in *S. cerevisiae.* | 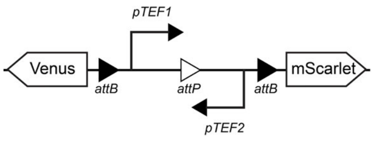 |

| | | |
|---|---|---|
| pBranch 02 | Construct for investigating the effects of *att* site arrangements on yeast cell differentiation. Designed for propagation in *E. coli* and integration at the ***URA3*** locus in *S. cerevisiae.* | 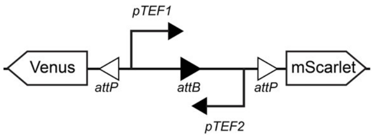 |
| pBranch 03 | Construct for investigating the effects of DNA length between *att* sites on yeast cell differentiation. Designed for propagation in *E. coli* and integration at the ***URA3*** locus in *S. cerevisiae.* | 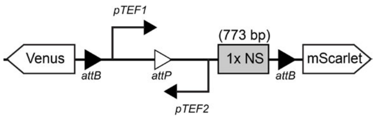 |
| pBranch 04 | Construct for investigating the effects of DNA length between *att* sites on yeast cell differentiation. Designed for propagation in *E. coli* and integration at the ***URA3*** locus in *S. cerevisiae.* | 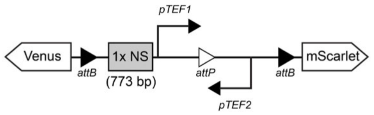 |
| pBranch 05 | Construct for investigating the effects of *att* sites with predictive recombination rates on yeast cell differentiation. Designed for propagation in *E. coli* and integration at the ***URA3*** locus in *S. cerevisiae.* | 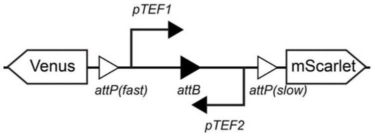 |
| pBranch 06 | Construct incorporating various factors to influence yeast cell differentiation. Designed for propagation in *E. coli* and integration at the ***URA3*** locus in *S. cerevisiae.* | 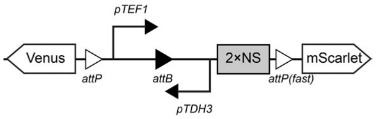 |
| pBranch 07 | Construct for designing multiple parallel and orthogonal circuits. Designed for propagation in *E. coli* and integration at the ***URA3*** locus in *S. cerevisiae.* | 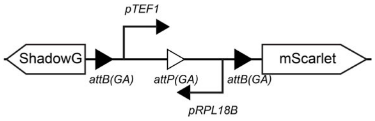 |
| pBranch 08 | Construct for designing multiple parallel and orthogonal circuits. Designed for propagation in *E. coli* and integration at the ***HO*** locus in *S. cerevisiae.* | 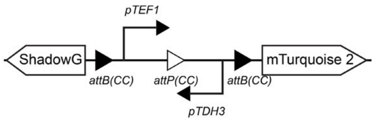 |

| | | |
|---|---|---|
| pBranch 09 | Construct for designing multiple parallel and orthogonal circuits. Designed for propagation in *E. coli* and integration at the ***LEU2*** locus in *S. cerevisiae.* | 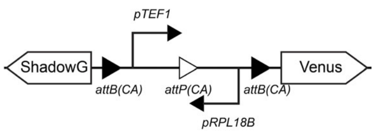 |
| pBranch 10 | Construct for designing multiple parallel and orthogonal circuits. Designed for propagation in *E. coli* and integration at the ***URA3*** locus in *S. cerevisiae.* | 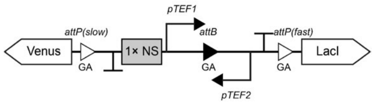 |
| pBranch 11 | Construct for designing multiple parallel and orthogonal circuits. Designed for propagation in *E. coli* and integration at the ***LEU2*** locus in *S. cerevisiae.* | 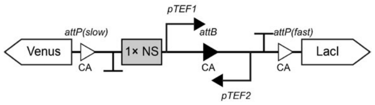 |
| pBranch 12 | Construct for designing multiple parallel and orthogonal circuits. Designed for propagation in *E. coli* and integration at the ***HO*** locus in *S. cerevisiae.* | 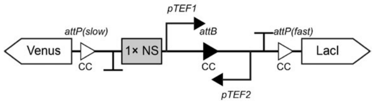 |
| *pLac*-mScarlet | Construct that drove mScarlet expression via promoter *pLac*. Designed for propagation in *E. coli* and integration at the ***MET15*** locus in *S. cerevisiae.* | 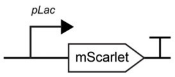 |
| pBranch 13 | Construct for designing multiple sequential circuits. Designed for propagation in *E. coli* and integration at the ***URA3*** locus in *S. cerevisiae.* | 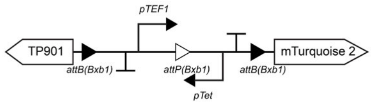 |
| pBranch 14 | Construct for designing multiple sequential circuits. Designed for propagation in *E. coli* and integration at the ***HO*** locus in *S. cerevisiae.* | 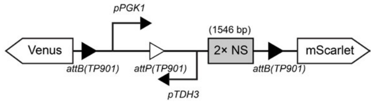 |
| pBranch 15 | Construct producing 2 progeny cells, each expressing a transcription factor. Designed for propagation in *E. coli* and integration at the ***URA3*** locus in *S. cerevisiae.* | 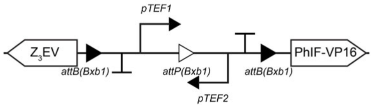 |

| | | |
|---|---|---|
| <u>pMYT076-*pPhlF*-CrtI</u> | Construct that drove *CrtE* expression via promoter *pPhlF*. Designed for propagation in *E. coli* and integration at locus 2 (**Int.2 site**) in *S. cerevisiae.* | 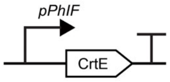  |
| <u>pMYT078-*pPhlF*-tHGM1</u> | Construct that drove *tHGM1* expression via promoter *pPhlF*. Designed for propagation in *E. coli* and integration at locus 4 (**Int.4 site)** in *S. cerevisiae.* | 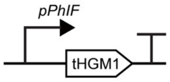  |
| <u>pMYT079-*pPhlF*-CrtYB</u> | Construct that drove *CrtYB* expression via promoter *pPhlF*. Designed for propagation in *E. coli* and integration at locus 5 (**Int.5 site**) in *S. cerevisiae.* | 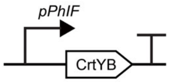  |
| <u>pMYT080-*pPhlF*-CrtE</u> | Construct that drove *CrtE* expression via promoter *pPhlF*. Designed for propagation in *E. coli* and integration at locus 6 (**Int.6 site**) in *S. cerevisiae.* | 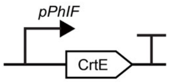  |
| <u>pMYT081-*pZ*-VioA</u> | Construct that drove *VioA* expression via promoter *pZ*. Designed for propagation in *E. coli* and integration at locus 7 (**Int.7 site**) in *S. cerevisiae.* | 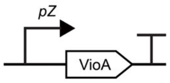  |
| <u>pMYT082-*pZ*-VioB-p2A-VioE</u> | Construct that drove *VioB* and *VioE* expression via promoter *pZ*. Designed for propagation in *E. coli* and integration at locus 8 (**Int.8 site**) in *S. cerevisiae.* | 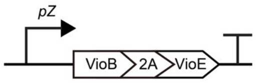  |
| <u>pMYT083-*pZ*-VioC</u> | Construct that drove *VioC* expression via promoter *pZ*. Designed for propagation in *E. coli* and integration at locus 9 (**Int.9 site**) in *S. cerevisiae.* | 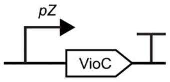  |
| <u>pMYT084-*pZ*-VioD</u> | Construct that drove *VioD* expression via promoter *pZ*. Designed for propagation in *E. coli* | 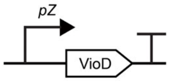  |

| | | |
|---|---|---|
| | and integration at locus 10 (**Int.10 site**) in *S. cerevisiae.* | |
| pMYT095-gRNA2456 | Construct encoding Cas9 and gRNAs for genome integration at loci 2, 4, 5, and 6. | 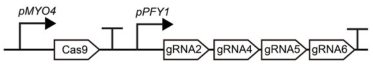 |
| pMYT095-gRNA78910 | Construct encoding Cas9 and gRNAs for genome integration at loci 7, 8, 9, and 10. | 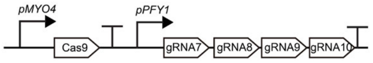 |
| *pZ*-Nb3-Venus | Construct that drove the surface-displaying of Nb3 via promoter *pZ*. Designed for propagation in *E. coli* and integration at the *LEU2* locus in *S. cerevisiae.* | 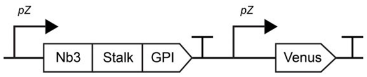 |
| *pPhlF*-Ag3 | Construct that drove the surface-displaying of Ag3 via promoter *pPhlF*. Designed for propagation in *E. coli* and integration at the *HO* locus in *S. cerevisiae.* | 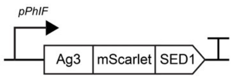 |

**Table. S8** | Representative plasmids used in CHO-K1 in this study. Annotated plasmid sequences are available for all constructs.

| Plasmid | Construct details | Scheme |
|---|---|---|
| LV-Doxy-Bxb1 | Construct expressing Bxb1 under the control of the *pTRE*$^{3G}$ promoter, designed for propagation in *E. coli* and lentiviral integration into HEK293FT or CHO-K1 cells. | 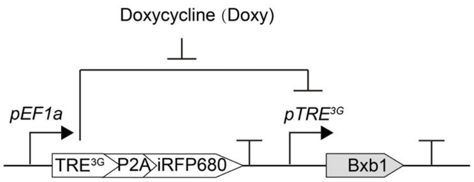 |
| LV-Doxy-Bxb1 (DHFR) | Construct expressing DHFR-Bxb1 under the control of the *pTRE*$^{3G}$ promoter, designed for propagation in *E. coli* | 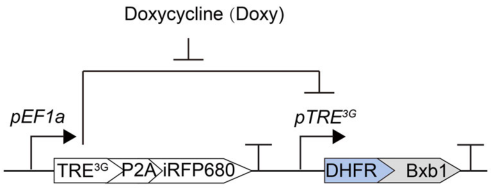 |

| | | |
|---|---|---|
| | and lentiviral integration into HEK293FT or CHO-K1 cells. | |
| LV-p5UAS-Cdh1-mCherry | Construct expressing Cdh1-T2A-mCherry under the control of the *p5UAS* promoter, designed for propagation in *E. coli* and lentiviral integration into HEK293FT or CHO-K1 cells. | 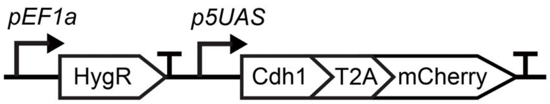 |
| pBranch-EGFP-pEF1-hUBC-mCherry | Branching circuit enabling mother cell differentiation into EGFP or mCherry lineages; integrated into HEK293FT or CHO-K1 via PhiC31. | 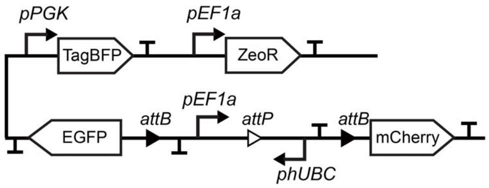 |
| pBranch-LaM4-ICAM1-pEF1-hUBC-mCherry-ICAM1 | Branching circuit enabling mother cell differentiation into LaM4- or mCherry-expressing lineages; integrated into HEK293FT or CHO-K1 cells via PhiC31-mediated recombination. | 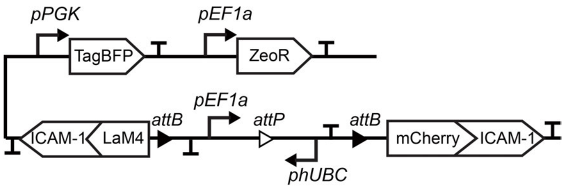 |
| pBranch-EGFP-ICAM1-pEF1-hUBC-LaG16-ICAM1 | Branching circuit enabling mother cell differentiation into EGFP- or LaG16-expressing lineages; integrated into HEK293FT or CHO-K1 cells via PhiC31-mediated recombination. | 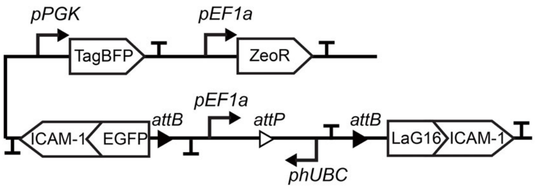 |
| pBranch-mKate2-Cdh6-pEF1-hUBC-Cdh1-mYFP | Branching circuit enabling mother cell differentiation into mKate2-Cdh6- or mYFP-Cdh1-expressing lineages; integrated into HEK293FT or CHO-K1 cells via PhiC31-mediated recombination. | 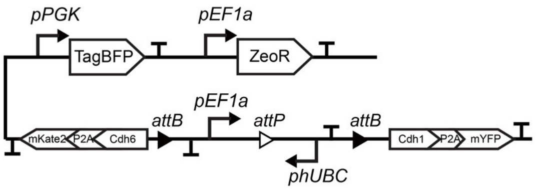 |
| pBranch-EGFP-TM-pEF1-hUBC-LaG17-GAL4-VP64 | Branching circuit enabling mother cell differentiation into EGFP-TM- or LaG17-Gal4 VP64-expressing | 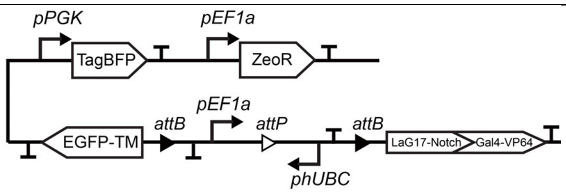 |

| | | |
|---|---|---|
| | lineages; integrated into HEK293FT or CHO-K1 cells via PhiC31-mediated recombination. | |
| pBranch-CD19-EGFP-TM-pEF1-hUBC-antiCD19-GAL4-VP64 | Branching circuit enabling mother cell differentiation into CD19-EGFP-TM- or antiCD19-Gal4 VP64-expressing lineages; integrated into HEK293FT or CHO-K1 cells via PhiC31-mediated recombination. | 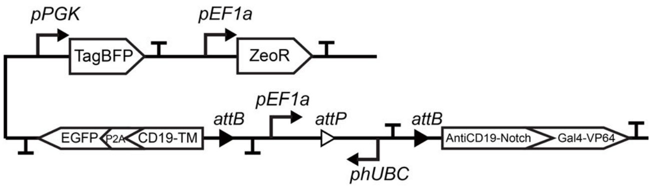  |
